# *mvh*: an R tool to assemble and organize virtual herbaria from openly available specimen images

**DOI:** 10.1101/2024.08.25.609600

**Authors:** Thais Vasconcelos, James D. Boyko

**Affiliations:** Department of Ecology and Evolutionary Biology, University of Michigan, Ann Arbor, Michigan 48109, USA; University of Michigan Herbarium, University of Michigan, Ann Arbor, Michigan 48108, USA; Michigan Institute of Data Science, University of Michigan, Ann Arbor, Michigan 48109, USA

**Keywords:** biodiversity, specimen digitization, GBIF, natural history collections

## Abstract

**Premise:** Recent efforts in digitizing and imaging herbarium specimens have enhanced their use in systematics, ecology, and evolutionary studies. However, there is a lack of user-friendly tools that facilitate the assembly and organizing of customized sets of herbarium specimen images on personal devices, i.e. a personal virtual herbarium.

**Methods:** Here we present the R package *mvh* (stands for “my virtual herbarium”), a software that includes functions designed to search and download metadata and openly available images associated with herbarium specimens based on taxon or geography. The download function also includes an argument to resize images according to a user-imputed quality preference.

**Results:** We tested the functionalities of *mvh* by searching metadata associated with five randomly sampled sets of ten vascular plant species (taxon-based search) and five sets of ten terrestrial coordinates (geography-based search). The main download function had a success rate of 99%, downloading 291 out of the 293 images found in the search. Possible reasons for download failure are also reported as part of the functions’ output.

**Conclusions:** As long as stable internet connection is available, the R package *mvh* makes the assembly and organizing of personal virtual herbaria an easy task that can help botanists to investigate novel empirical questions as well as trends in digitization efforts.

## Introduction

Herbarium collections, originally established to preserve dried plant specimens for pharmaceutical and taxonomic purposes, have evolved into irreplaceable resources of data that can be used to answer a broad range of scientific questions (Funk, 2003; Lavoie, 2013). Just in the last decade, these collections have been used to address topics such as historical changes in plant distribution (Feeley, 2012), extinction risk assessment (Nic Lughadha et al., 2019), tracking changes in phenological patterns associated with climate change (Park et al., 2019), phylogenomics (Maurin et al., 2021), and others, proving that their applications are difficult to predict but continue to grow in significance. Much of the novel applications of herbarium specimens in biodiversity research have been made possible due to the increase in efforts associated with specimen digitization and imaging. As herbarium collections enter the digital era, digitization projects have become a priority for many institutions, first to safeguard specimens, ensuring that this source of data is preserved should anything happen to the physical collections, but then also to allow botanists to find and use specimens deposited across many institutions (Nelson et al., 2013; Sweeney et al., 2018).

Image data, in particular, has become increasingly important in recent years as the accessibility of computer vision tools has expanded (Weinstein, 2018; Lürig et al., 2021). These new tools enable botanists to use large datasets of herbarium specimen images to automate the extraction of label information (Weaver and Smith, 2022), as well as the scoring of phenological stage (Davis et al., 2020) and measurement of phenotypic traits such as color (Boyko, 2024) and leaf size (Weaver et al., 2020). Of course, the quality of the image and associated metadata are critical for the successful application of these tools, and therefore these tools need to be supported by user-friendly applications that allow users to easily assemble well-curated datasets of specimen images.

Here we present the R package *mvh* (stands for “my virtual herbarium”) which includes a set of functions designed to facilitate building datasets of images of herbarium specimens and associated metadata in personal devices, based on taxon or geography. The package interfaces with GBIF by using rgbif (Chamberlain and Boettiger, 2017) functions to compile a list of specimens with images openly available for download. It then accesses the URLs linked to these images and downloads them into a user-determined directory, naming the files according to taxon name and GBIF ID, a unique identifier for each collection. R package *mvh* is currently submitted to CRAN and available through GitHub at https://github.com/tncvasconcelos/mvh.

### Functionalities

The main pipeline of *mvh* includes two functions: “search_specimen_metadata” and “download_specimen_images” (Table 1, Figure 1A). The first function, “search_specimen_metadata”, interacts with rgbif’s occ_search to search and find records of preserved specimens with images in the GBIF dataset. This search can be either taxon-based with the argument “taxon_name”, geography-based with the arguments “coordinates” and “buffer_distance”, or both. Note that, because the function “search_specimen_metadata” is essentially a wrapper of rgbif::occ_search, it will take any argument that rgbif::occ_search takes, and more experienced R users may want to use this functionality to customize their search in other ways. Taxon-based search will take any scientific name under GBIF’s taxonomic backbone for the search. Geography-based searches will take any latitude and longitude passed through the “coordinates” argument and create a square around that point of edge length in degrees determined by the “buffer_distance” argument. For instance, if the user inputs “coordinates = c(42.28, -83.74)” and “buffer_distance=1”, it will create a square of edge 1 degree with centroid in latitude 42.28° N and longitude 83.74° W. As a result, “search_specimen_metadata” will return a data.frame object containing all metadata associated with the specimens found under the taxon and/or geography criteria, including a column with the URL for the specimen image (column “media_url”) and the license regulating data usage.

**Table 1:** Main functionalities of R package *mvh*. Note that both search_specimen_metadata and download_specimen_images require internet connection to run.

| Function name | Short description | Main arguments | Output |
| --- | --- | --- | --- |
| `search_specimen_metadata` | Interacts with `rgbif` to search metadata associated with preserved specimens with images available. | “`taxon_name`” (for taxon-based searches), “`coordinates`”, and “`buffer_distance`” (for geography-based searches) | A data.frame including all metadata associated with specimens returned by the search. |
| `download_specimen_images` | Takes the output of `search_specimen_metadata` to download images of herbarium specimens through their URLs. | “`metadata`” (the output from `search_specimen_metadata`) | A data.frame with summarized metadata and the status of the download. |
| `plot_specimens_by_institution` | Takes the output of `search_specimen_metadata` to plot a barplot of the institutions where most specimens are deposited. | “`metadata`” (the output from `search_specimen_metadata`) | A barplot sorted in decreasing order of number of specimens per institution. |
| `plot_specimens_by_country` | Takes the output of `search_specimen_metadata` to plot a barplot of the countries where most | “`metadata`” (the output from `search_specimen_metadata`) | A barplot sorted in decreasing order of |
|  | specimens were collected. |  | number of specimens per country. |

**Figure 1:**
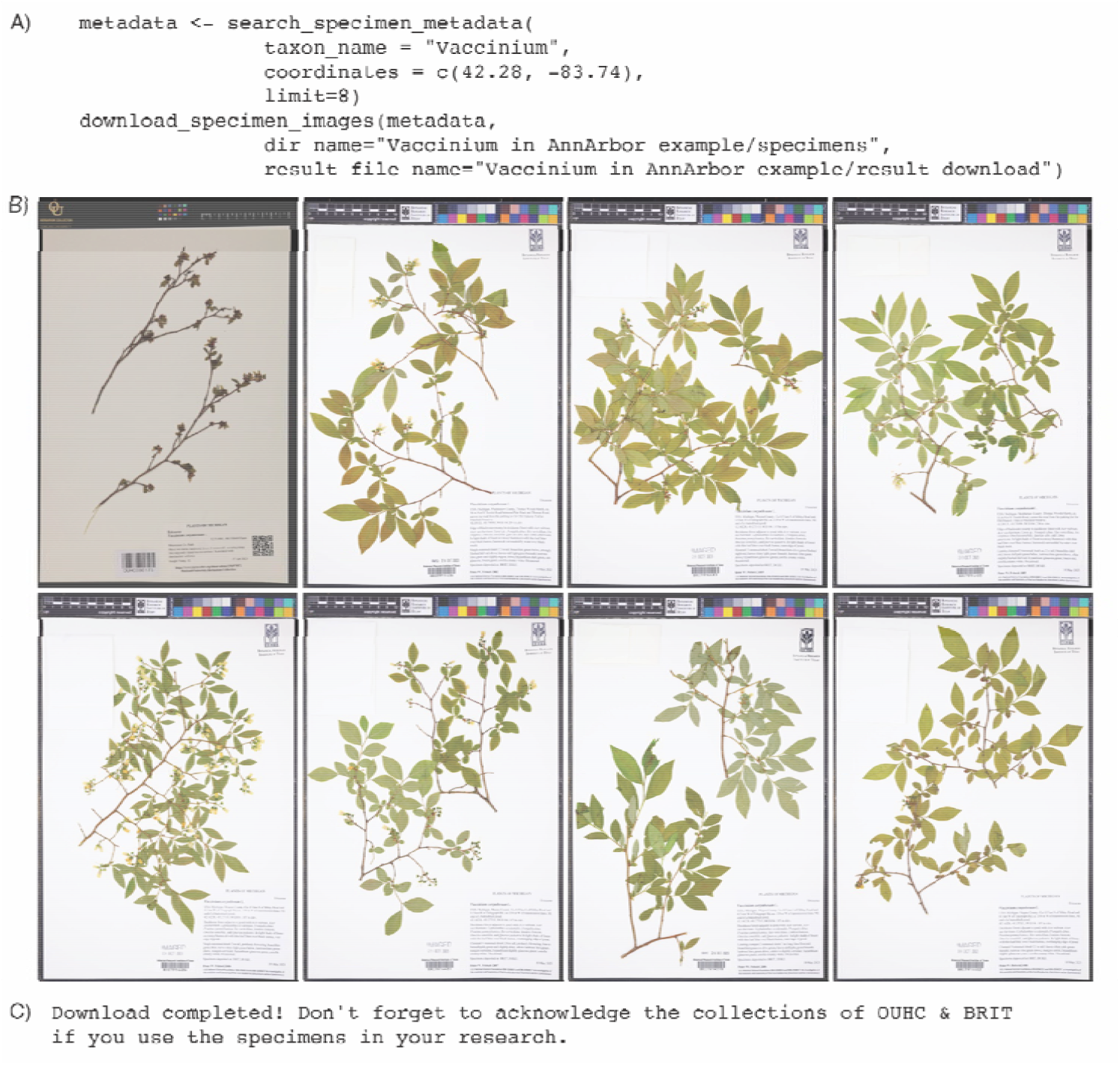
A) Example of a *mvh* pipeline to search and download up to eight specimens (“limit=8”) of the blueberry genus *Vaccinium* (Ericaceae) from the Ann Arbor (MI, USA) area (“coordinates = c(42.28, -83.74)”). B) Specimens’ images downloaded as a result of the pipeline. C) Message reminding the user to acknowledge collections where specimens are deposited if they are used in publications.

The second function in the pipeline, “download_specimen_images”, will take as the main argument the resulting data.frame from “search_specimen_metadata” to download all images linked to the URL in the media_url column in format .jpg (Figure 1B). Each image will be named as a combination of taxon name and gbifID, an unique number that will allow the user to connect the specimen to the metadata table (given that many GBIF observations lack collector name and number). Other arguments in the “download_specimen_images” function allow the user to customize and organize the download of the images. The argument “dir_name” will determine the name of the folder created in the working directory at the beginning of the downloading process where the images are going to be saved. The “result_file_name” argument allows the user to rename the result table to be written in the working directory with the status of the download. This result table is a spreadsheet created in the designated working directory with summarized metadata about the specimens returned by the search. As the function attempts to download each of the images returned by the search, it will write in the last two columns of the spreadsheet whether the download was successful or not (column “status”) and, if the download failed, the message associated with the error for further investigation (column “error_message”) (e.g. so the user knows if download failed due to an issue of broken URL or internet instability). The spreadsheet will also return the license regulating the use of the image, the holder of the rights to the image and image size in bytes.

The “resize” argument will specify whether the images should be resized after the download. The default of this argument is NULL, meaning that if nothing is changed the images will keep their original quality and size after download. However, if the user chooses to use this argument, it will take any value from 1 to 100 to determine how much should the size and quality of the original file be reduced to. For instance, a resize=75 will reduce the original image to 75% of its original size. Depending on the goal of the user, using the “resize” feature may be important because some high quality herbarium images are heavy, which can limit the assembly of large collections depending on the space available in the user’s personal device. High quality herbarium specimens that can be used for looking at tiny morphological structures (e.g. trichomes) are large in terms of file size and building a virtual herbarium of 100 specimen images of original quality may require over 1GB of storage space in the personal device.

However, this feature should be used with caution in certain situations, because the accuracy of newly available software to automate trait scoring from herbarium images (e.g. LeafMachine, Weaver et al., 2020) is often resolution dependent. The user will need to evaluate the use of the argument on a case-by-case basis. Finally, the argument “timeout_limit” will determine how long R should spend trying to download an image before the connection fails. The default for the “timeout_limit” is 300 seconds, meaning that each download will try to connect to the network for 5 minutes before crashing. Adjusting this argument can be helpful if the user has an unstable internet connection and requires a longer buffer time to complete a given download. At the end of the download, the function will also print on the console a message reminding the user to acknowledge the collections where the images come from if they are used in publications (Figure 1C).

The package *mvh* also includes two plotting functions to allow the user to visualize two important components of the metadata associated with the search. The first is in which countries the collections in their personal herbaria were collected (“plot_specimens_by_country”) and the second is in which institutions they are currently deposited (“plot_specimens_by_institution”), which often follow herbarium acronyms according to Thiers (cont. updated) (Table 1). Both functions will take as argument the data.frame resulting from the “search_specimen_metadata” function to plot a barplot in decreasing order of number of specimens in each category (Figure 2).

**Figure 2:**
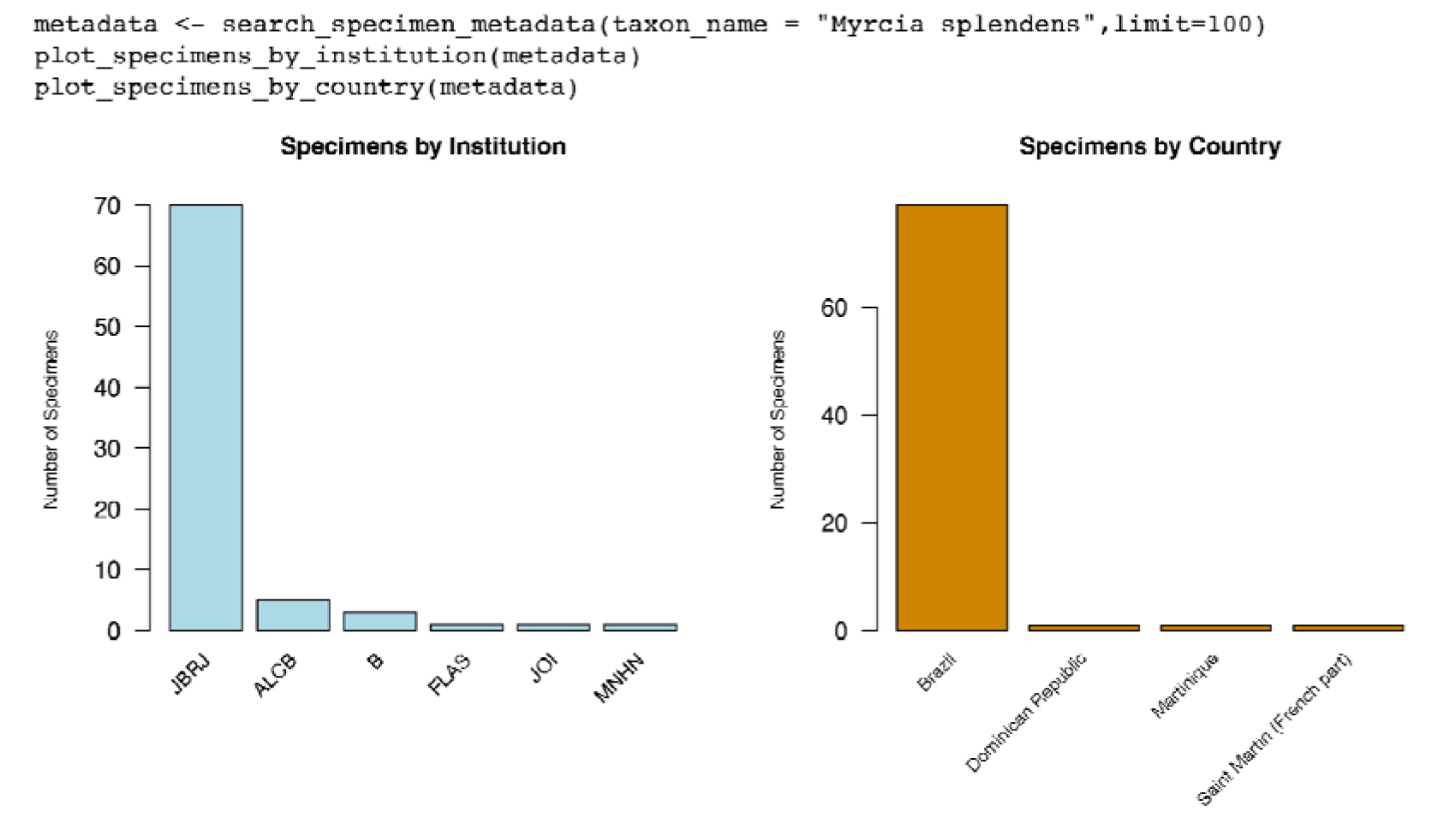
Example of a *mvh* pipeline to search up to 100 specimens (limit=“100”) of the widespread species *Myrcia splendens* (Myrtaceae) and plot the number of specimens per institution and country.

### Study case

We tested the functionality of the *mvh* package by running the main pipeline in two study cases, a taxon-based one and a geography-based one. In the first, we took five random samples of 10 species each from the list of accepted species of vascular plants from POWO (2024). In the second, we used as “coordinate” input the latitude and longitude of ten global national parks.

Because the intention was merely to test the success rate of the downloads, in both cases we set the “limit” argument of “search_specimen_metadata” to “limit=5” for speed, meaning the search will be limited to five specimens. The “resize” argument of “download_specimen_images” was set to “resize=5”, meaning that downloaded images will be resized to 5% of their original quality. The pipeline was run at an internet speed of 350 MBPS using a MacBook Pro macOS Venture 13.1. The full script for the search, including our seed number for the random sample of species, latitude and longitude of the ten global national parks, result tables, images, and code to summarize the results is available as Supplementary Material.

The pipeline found 242 specimen images in the taxon-based search, successfully downloading 240 of them in 22.32 minutes, which results in a success rate of 99% and an average of 5.58 seconds per specimen. In the geography-based search, the pipeline found 51 specimen images and successfully downloaded 51 of them in 6.42 minutes, for a 100% success rate and an average download speed of 7.55 seconds per image. The original size of the files at the time of download ranged from 4.34KB to 80.01MB, with a median of 226.4KB (Figure 3A), requiring 1.26GB of space in the personal device to complete the download at the original quality. After resizing images to 5% of their original quality, however, the 291 downloaded images occupied a total of 109.2 MB of space, with images ranging from 1.97KB to 7.83MB, with a median of 78.80KB (Figure 3B).

**Figure 3:**
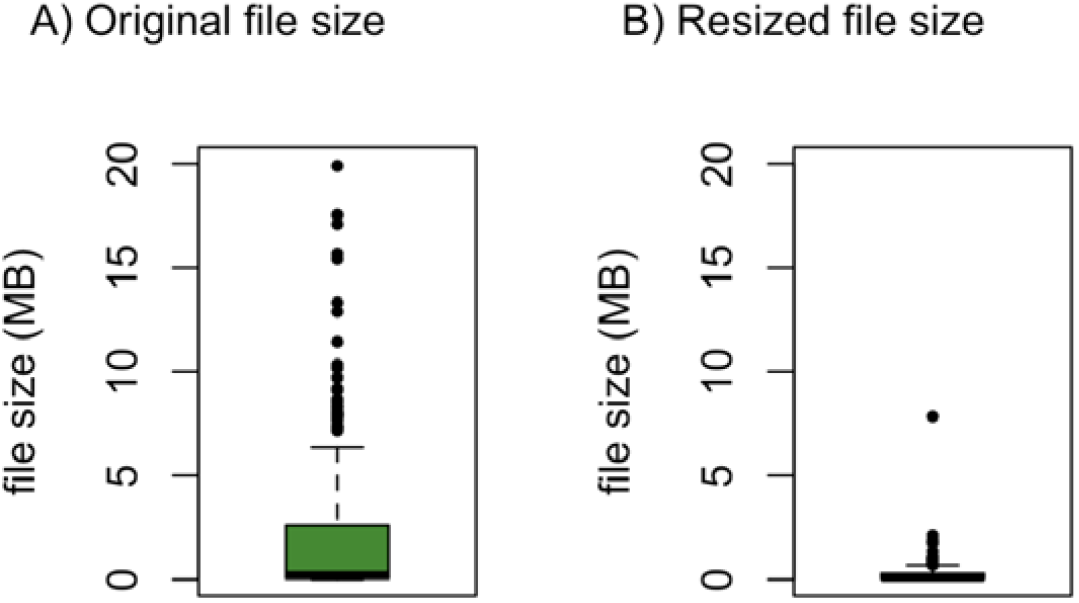
Comparison of file sizes before (“resize=100”) and after (“resize=5”) resizing.

## Discussion

### Comparison with other software and additional notes

To our knowledge, there is currently no other user-friendly pipeline to download and organize images of herbarium specimens available as an R package. Performing automated download of specimen images in R is possible through a combination of functions of R package rgbif and some programming, which is essentially what we developed and documented here.

An advantage of using GBIF as a base for our URL search is that it is not only the largest data aggregator available for biodiversity data, but it also allows fuzzing matches of taxon names that are not properly harmonized through the function rgbif::occ_search. It is important to note that the use of our pipeline does not generate a DOI linked to the metadata as it is the case for rgbif::occ_download. However, our pipeline still allows for reproducibility as long as users make the result tables available as supplementary material in their projects. We also advise users to acknowledge the collections where the specimens come from, as these are the entities responsible for making the data available.

### Limitations

As we worked on the development of this package and in improving the success rate of the downloads, we observed that, as long as the metadata has a valid URL in the identifier slot of the media file on GBIF, the download is very likely to be successful. However, two main types of errors in failed downloads were recurrent: (1) error of URL not existing or not linked with an image, which will appear as “cannot open URL [url]” in the “status” column of the result table; and (2) error of time limit for download reached, which will appear as “download from [url] failed” in the “status” column of the result table. The first error is difficult to tackle because it can be caused by many different issues, including human error during specimen digitization (URL in the wrong format or in the wrong data slot) and media data that used to exist but has since been excluded by the collections after being exported to GBIF. Different collections export their metadata in different formats, and we extensively tested the functionalities of *mvh* so that it is able to capture this variation for most of the large collections. The second error can usually be dealt with by increasing the “time.limit” arguments of the “download_specimen_images” function. In fact, we observed that the percentage of failed downloads in very large queries decreased from 98% to 99% once a “time.limit” of 300 seconds was set, the reason why we set this as the function’s default.

Another observed source of download failure is authentications such as CAPTCHAs that some collection websites require to access images through the URL available on GBIF. In those cases, we observed that once the conditions for accessing the image are accepted once, all subsequent attempts to download images using our R pipeline proceed normally when the same computer is used. In that way, we advise users to manually access the URLs printed in the “error_message” column of the result table and rerun the pipeline after verifying if this is the reason for failure downloads. A final limitation to discuss is that, because the function “download_specimen_images” works by accessing the URL of the specimen image as provided by the original institutions, it will occasionally download images that are not of herborized material (e.g. living plants) if their URL address is indistinguishable from that of the equivalent herbarium specimen. Currently, we found no way to avoid this to happen occasionally in large queries, but we anticipate that this will not be a big issue with most projects.

## Conclusion

The R package *mvh* provides a flexible and user-friendly pipeline to download images of herbarium specimens deposited across institutions. It can accelerate and facilitate the use of this information for taxonomy, floristics, phenotypic analyses and biodiversity studies in general. Currently, most herbarium collections are not completely digitized, so virtual herbaria are not perfect representations of the whole range of physical specimens deposited in these institutions. However, we do think that tools like this package will become increasingly useful as more collections are being digitized and imaged, and novel uses of herbarium specimen images are unlocked.

## Acknowledgements

We thank Will Weaver for conversations that improved this manuscript and the *mvh* pipeline. We also thank the herbaria where the images used here are deposited for making specimen images openly available for non-commercial research: A, B, BAYLU, BBM, BR, BRI, BRIT, BRLU, BRY, CHR, CM, COI, COLO, CR, DBG, DES, E, F, GH, GJO, HIFP, JBRJ, K, KYO, LBV, MEL, MICH, MNHN, MO, MW, NBF, NCU, NEBC, NEON, NGCPR, NHMUK, NMNZ, NSW, NY, P, PRC, RBGE, RSA, TASM, TRH, UCSB, UCR, US, W, WS, WU, Z (all acronyms following Thiers, cont. updated).

## References

Chamberlain, S. A., Boettiger, C. (2017) R Python, and Ruby clients for GBIF species occurrence data. PeerJ Preprints 5: e3304v1.

Davis, C. C. (2023). The herbarium of the future. Trends in Ecology & Evolution, 38(5), 412–423.

Davis, C. C., Champ, J., Park, D. S., et al. (2020). A new method for counting reproductive structures in digitized herbarium specimens using mask R-CNN. Frontiers in Plant Science, 11, 1129.

Feeley, K. J. (2012). Distributional migrations, expansions, and contractions of tropical plant species as revealed in dated herbarium records. Global Change Biology, 18(4), 1335–1341.

Funk, V.A., (2003) The importance of herbaria. Plant Science Bulletin. 49(3) 2003

Hussein, B. R., Malik, O. A., Ong, W. H., & Slik, J. W. F. (2022). Applications of computer vision and machine learning techniques for digitized herbarium specimens: A systematic literature review. Ecological Informatics, 69, 101641.

Lavoie, C. (2013) Biological collections in an ever changing world: Herbaria as tools for biogeographical and environmental studies. Perspectives in Plant Ecology, Evolution and Systematics, 15(1), 68–76.

Lürig, M. D., Donoughe, S., Svensson, E. I., Porto, A., & Tsuboi, M. (2021). Computer vision, machine learning, and the promise of phenomics in ecology and evolutionary biology. Frontiers in Ecology and Evolution, 9, 642774.

Maurin, O., Anest, A., Bellot, S., et al. (2021). A nuclear phylogenomic study of the angiosperm order Myrtales, exploring the potential and limitations of the universal Angiosperms353 probe set. American Journal of Botany, 108(7), 1087–1111.

Nelson, G., Paul, D., Riccardi, G. & Mast, A.R. (2012) Five task clusters that enable efficient and effective digitization of biological collections. ZooKeys, (209), p. 19.

Nic Lughadha, E., Walker, B. E., Canteiro, C., et al. (2019). The use and misuse of herbarium specimens in evaluating plant extinction risks. Philosophical transactions of the Royal Society B, 374(1763), 20170402.

Park, D. S., Breckheimer, I., Williams, A. C., Law, E., Ellison, A. M., & Davis, C. C. (2019). Herbarium specimens reveal substantial and unexpected variation in phenological sensitivity across the eastern United States. Philosophical Transactions of the Royal Society B, 374(1763), 20170394.

POWO (2024) Plants of the World Online Facilitated by the Royal Botanic Gardens, Kew. URL http://www.plantsoftheworldonline.org/ [Accessed August, 2024].

R Core Team (2024). R: A language and environment for statistical computing. R Foundation for Statistical Computing, Vienna, Austria. URL https://www.R-project.org/.

Sweeney, P.W., Starly, B., Morris, P.J., Xu, Y., Jones, A., Radhakrishnan, S., Grassa, C.J. & Davis, C.C. (2018) Large–scale digitization of herbarium specimens: Development and usage of an automated, high–throughput conveyor system. Taxon, 67(1), 165-178.

Thiers B.M. (Continuously updated) Index Herbariorum: A global directory of public herbaria and associated staff. New York Botanical Garden’s Virtual Herbarium. https://sweetgum.nybg.org/science/ih/ [Accessed August, 2024]

Weinstein, B.G. (2018) A computer vision for animal ecology. Journal of Animal Ecology, 87(3), 533–545.

