## Supplementary figures and images for "*mvh*: an R tool to assemble and organize virtual herbaria from openly available specimen images"

### Cousinia_resinosa_1799060282.jpeg

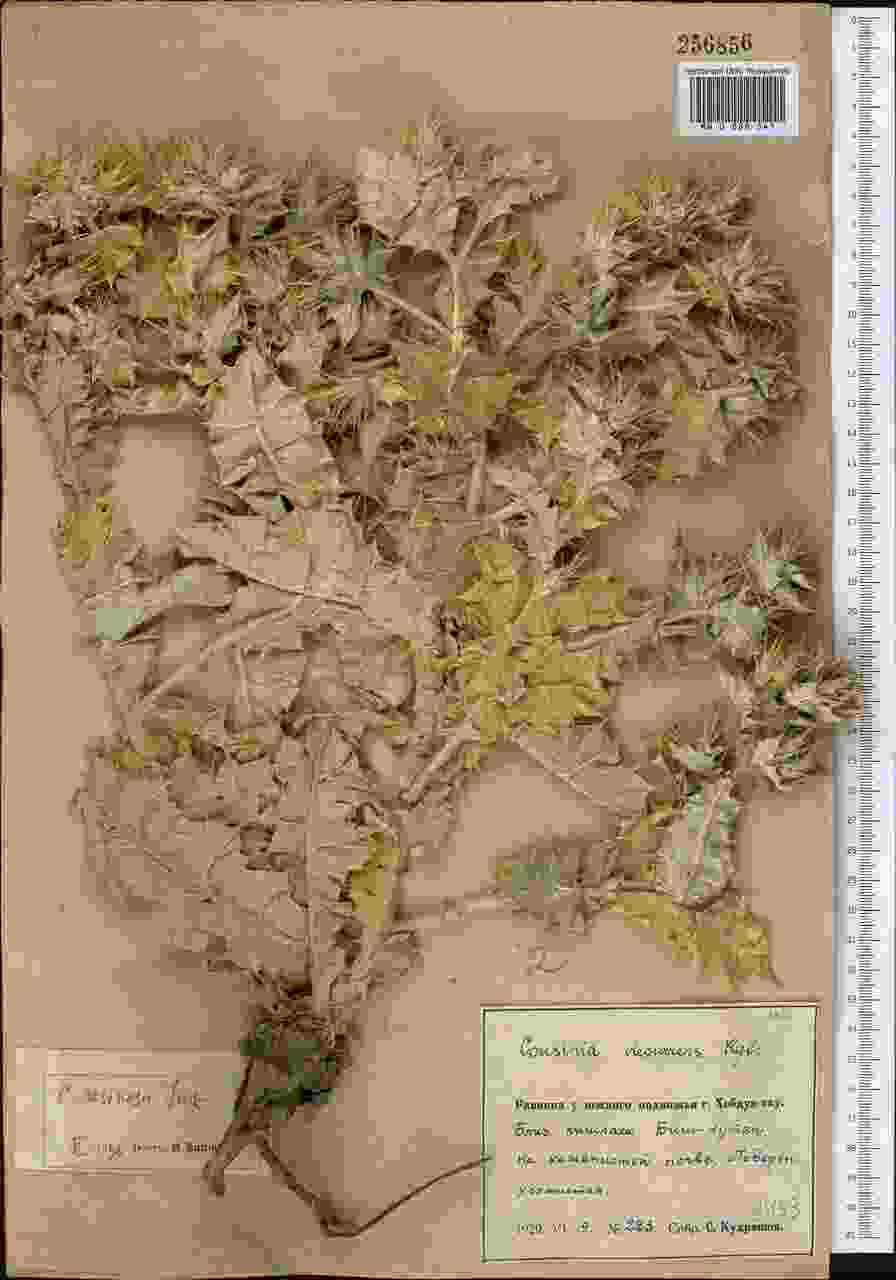

### Hermannia_erodioides_1840197175.jpeg

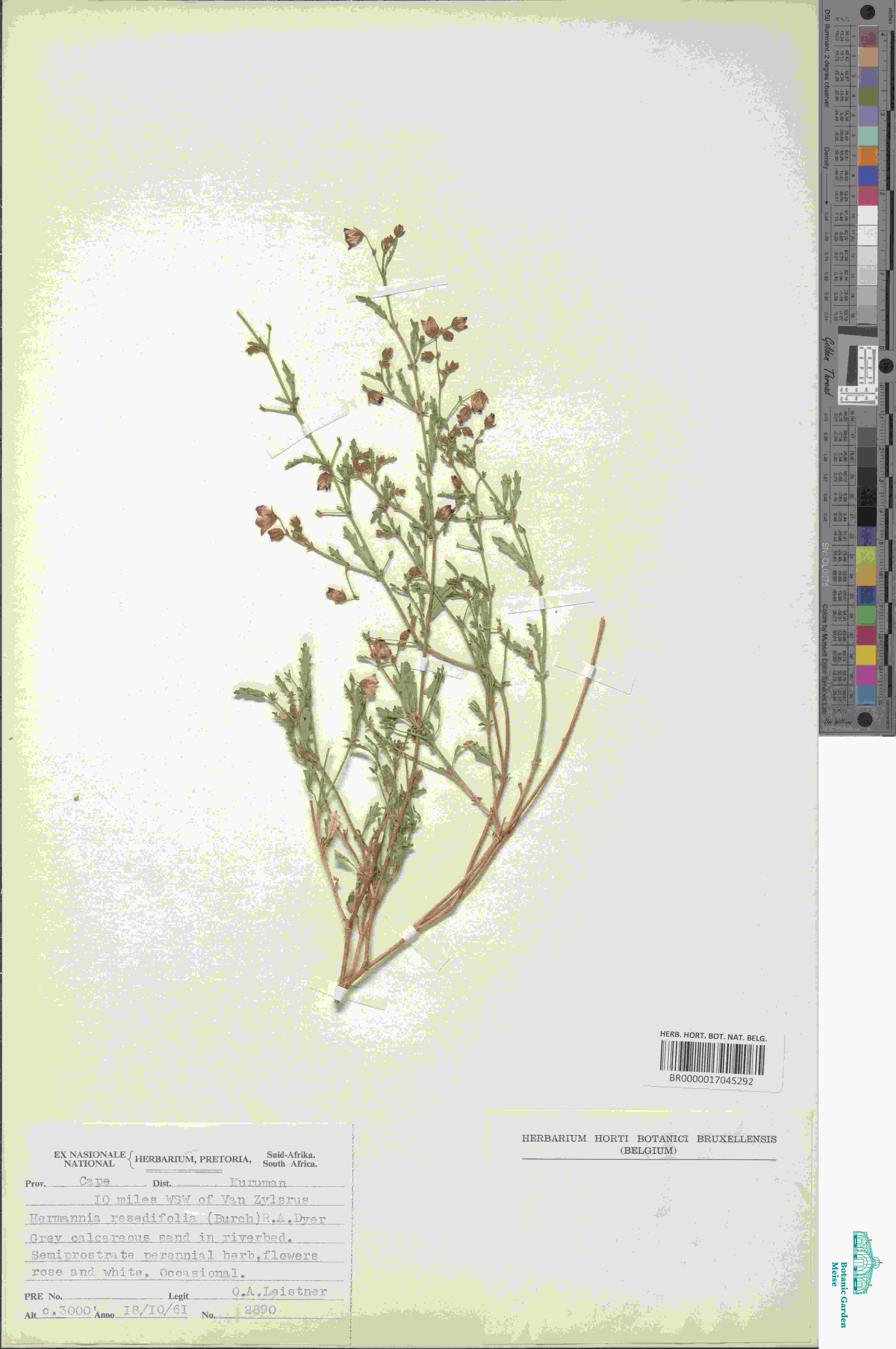

### Hermannia_erodioides_2515572126.jpeg

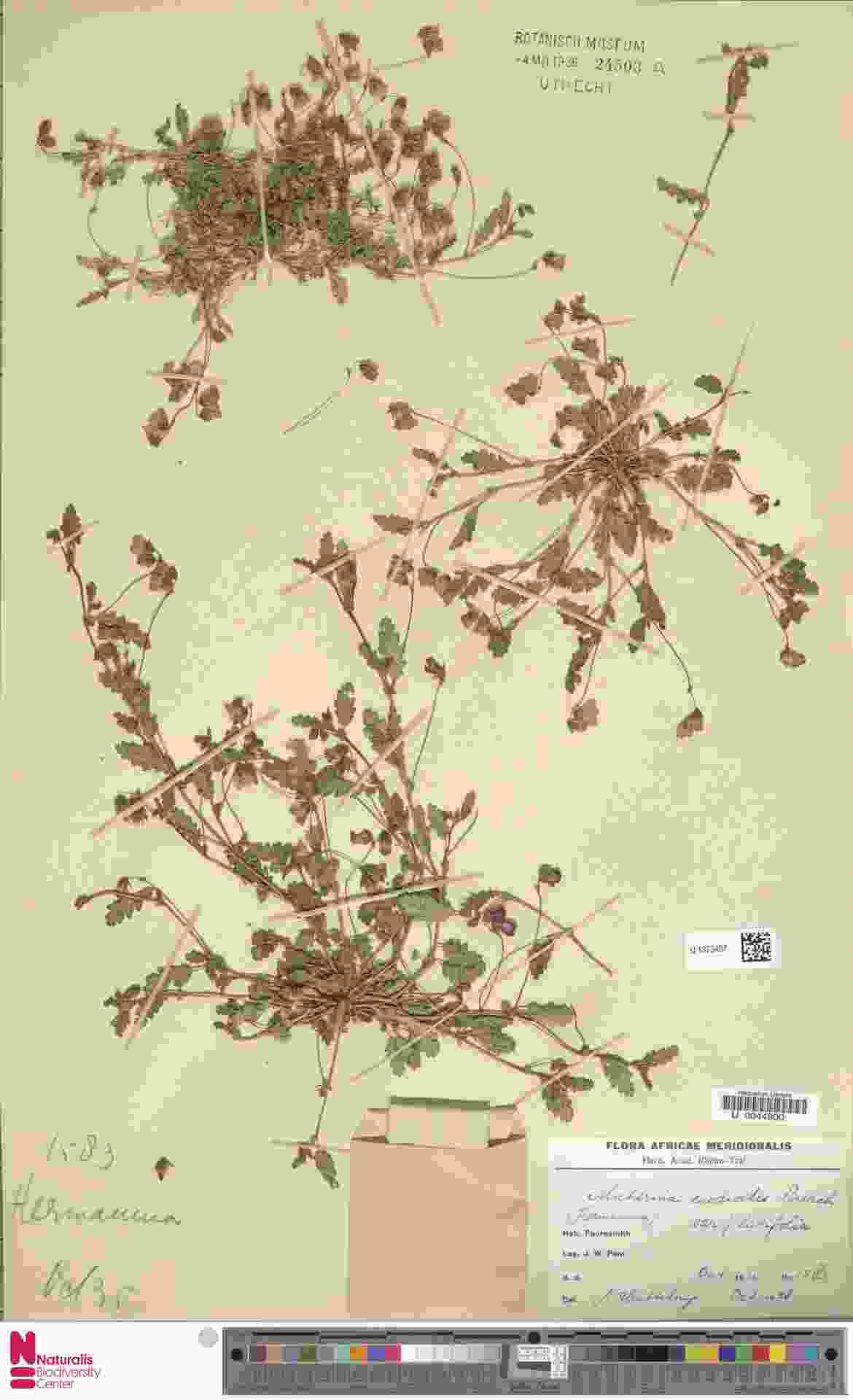

### Hermannia_erodioides_4876273741.jpeg

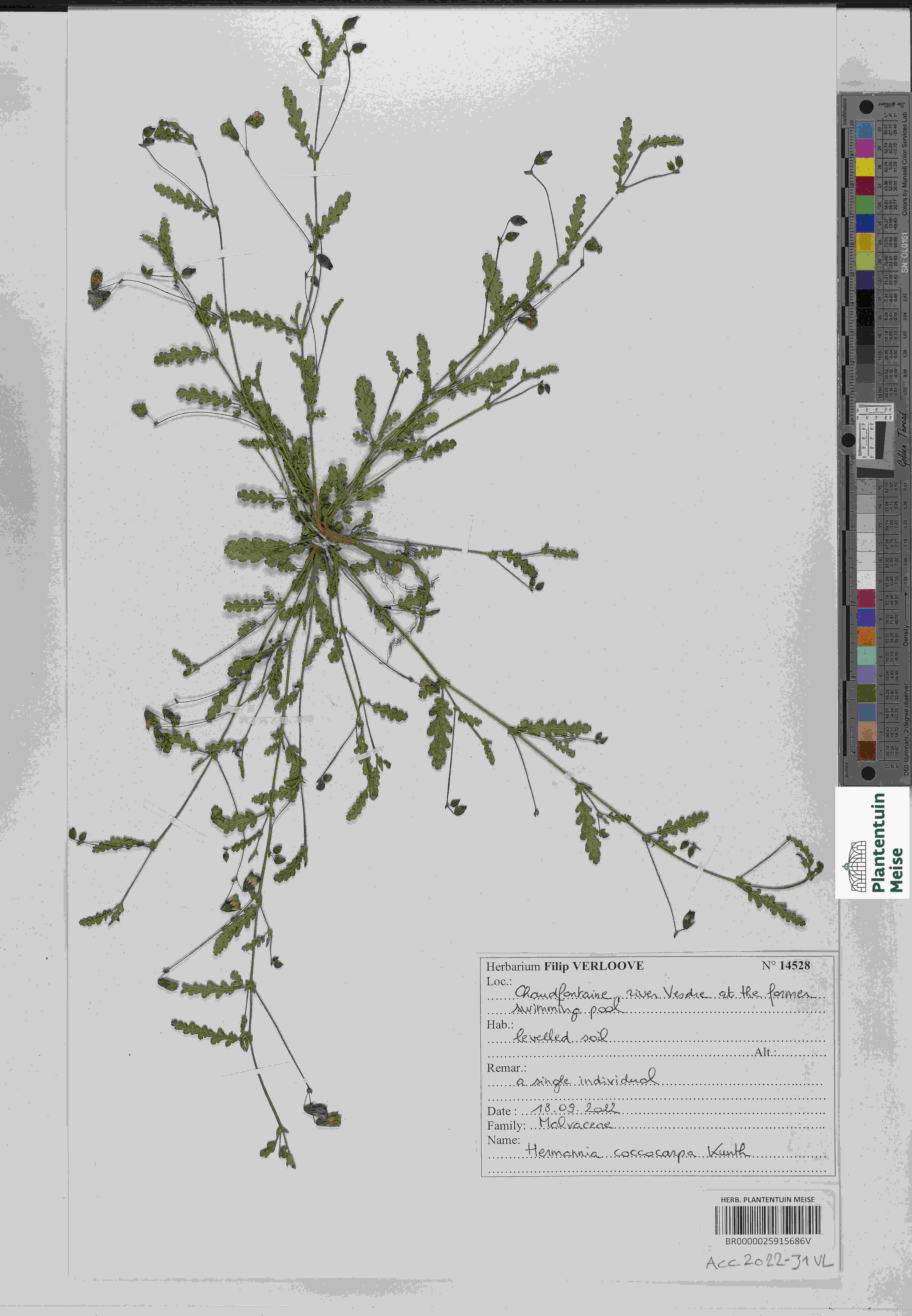

### Lipotriche_scandens_1455982488.jpeg

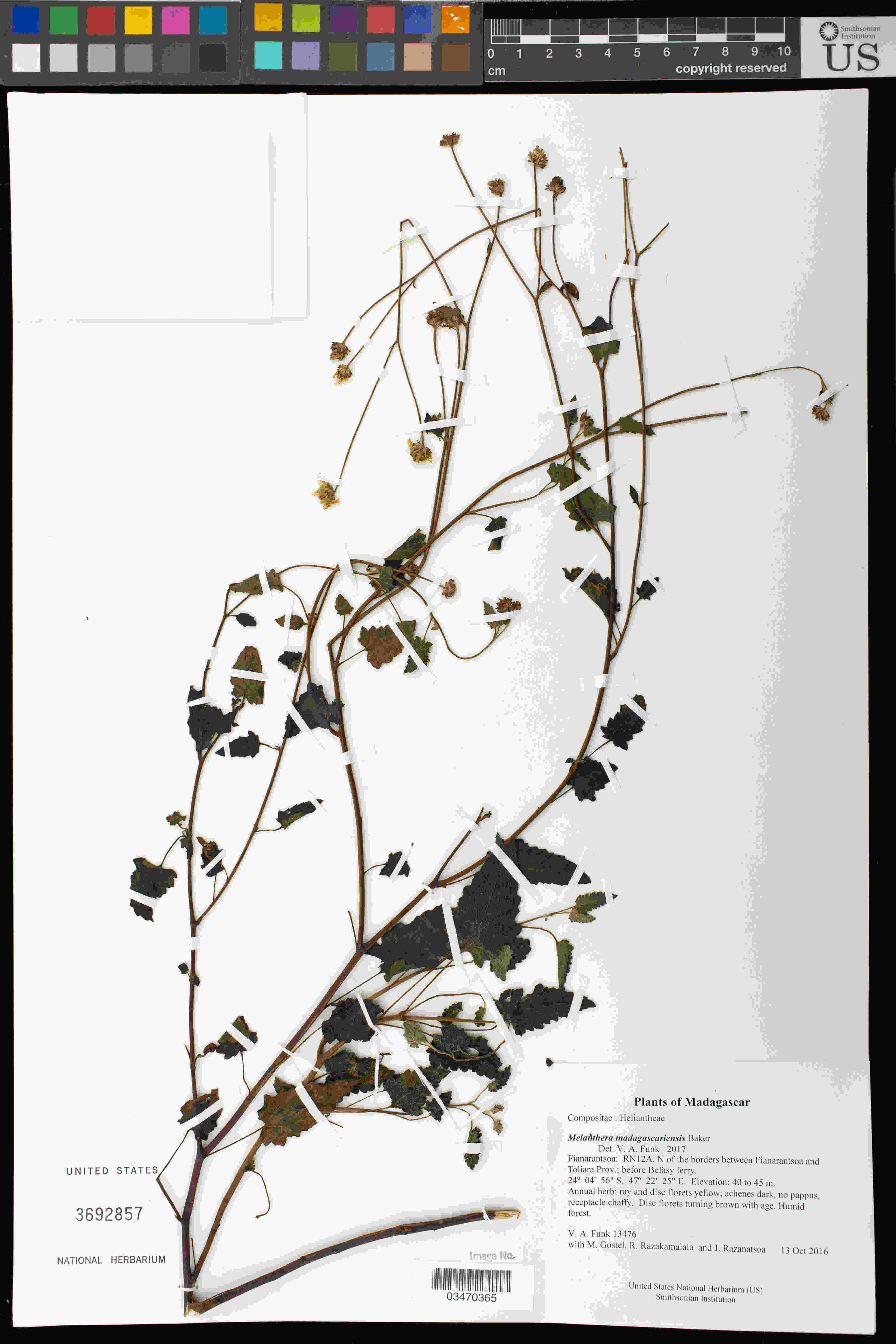

### Lipotriche_scandens_2268854208.jpeg

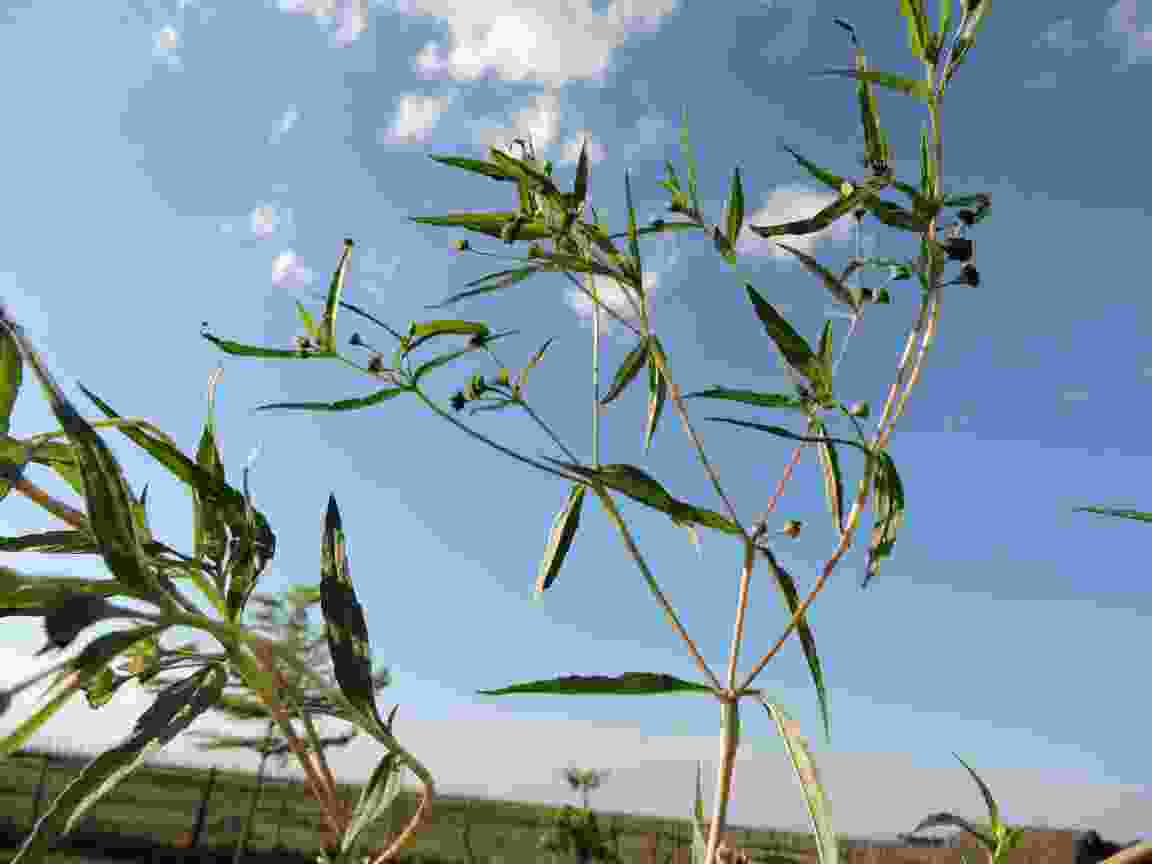

### Lipotriche_scandens_4519477085.jpeg

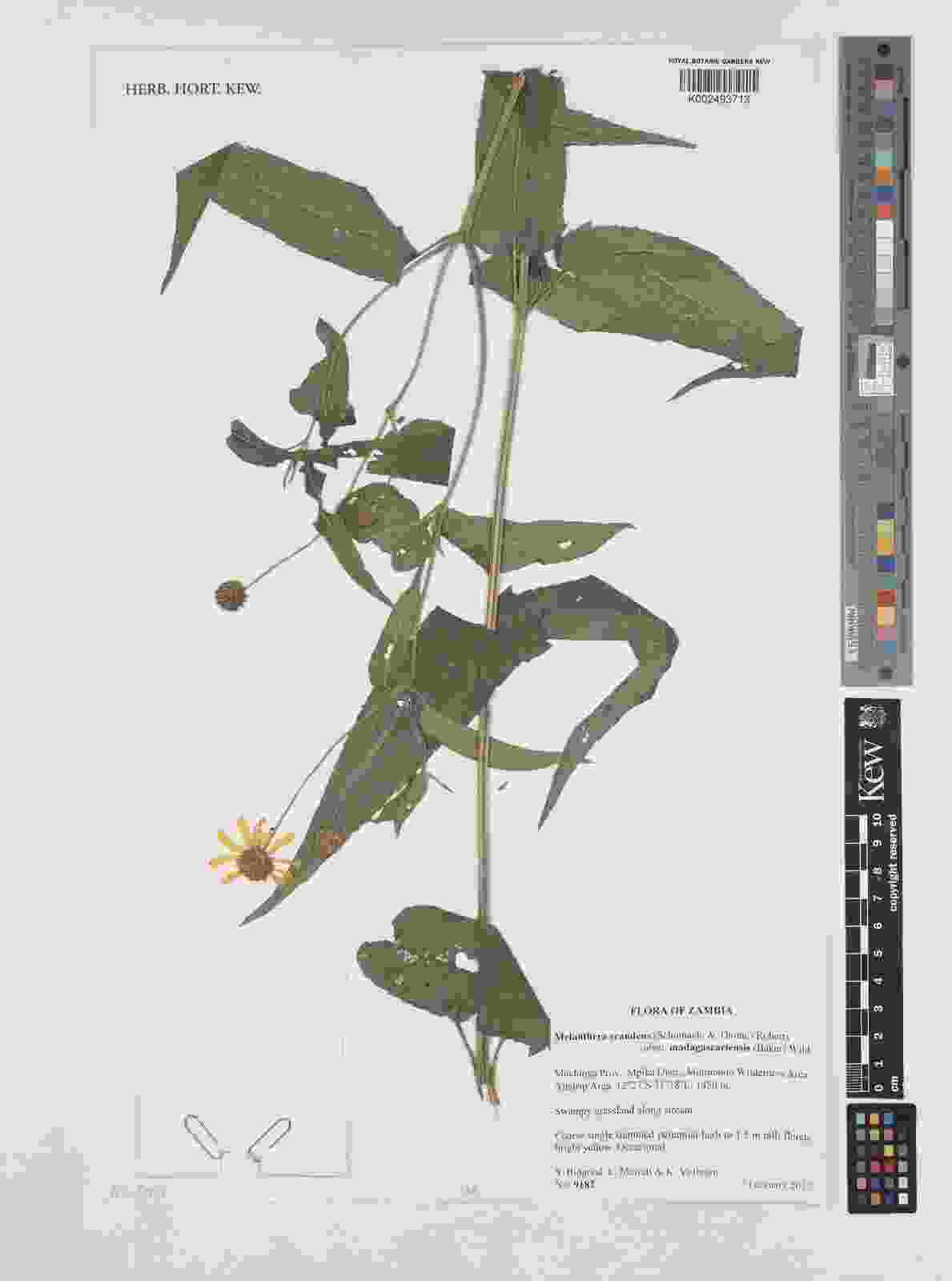

### Lipotriche_scandens_4519489127.jpeg

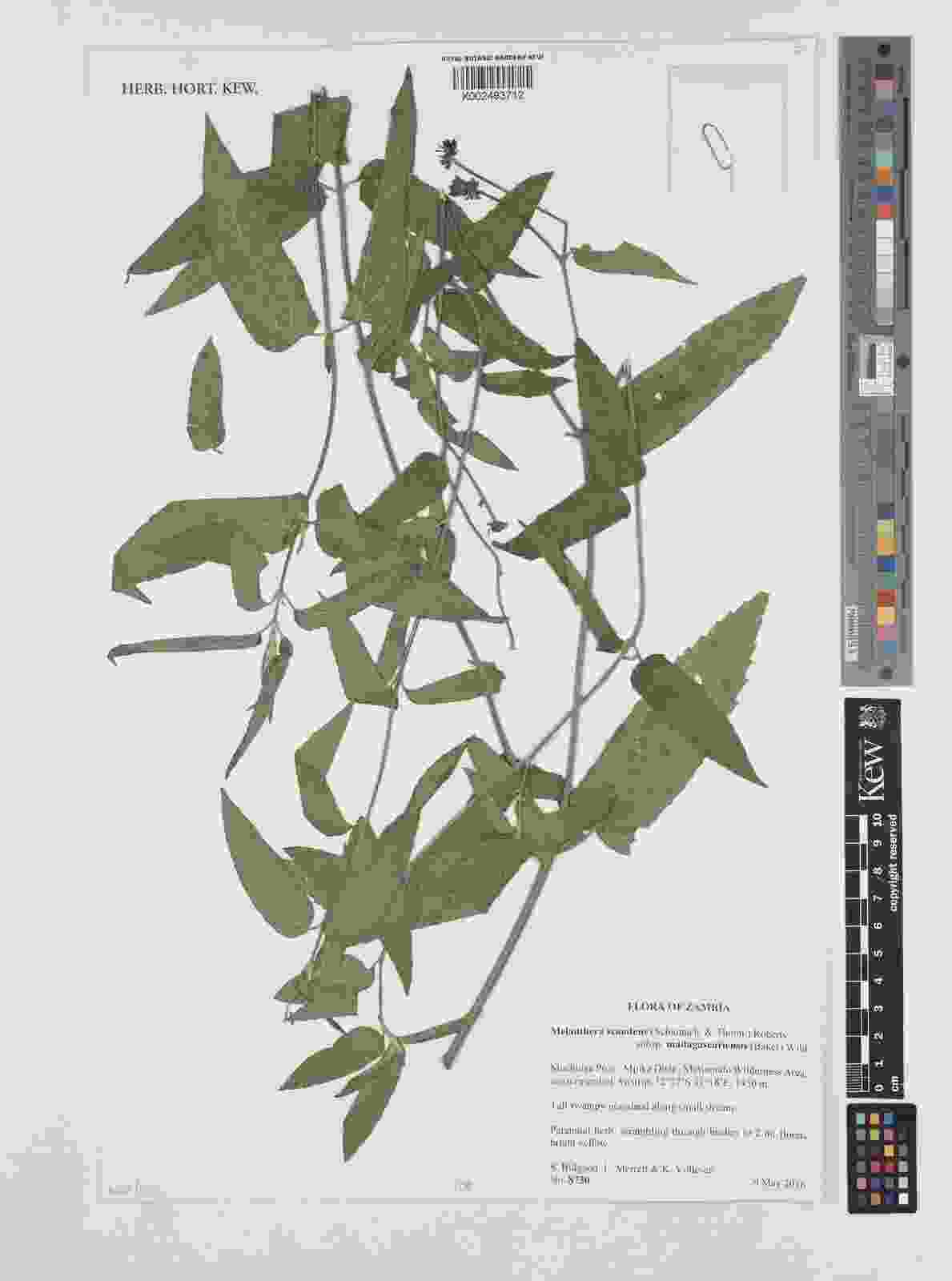

### Piper_staminodiferum_437530854.jpeg

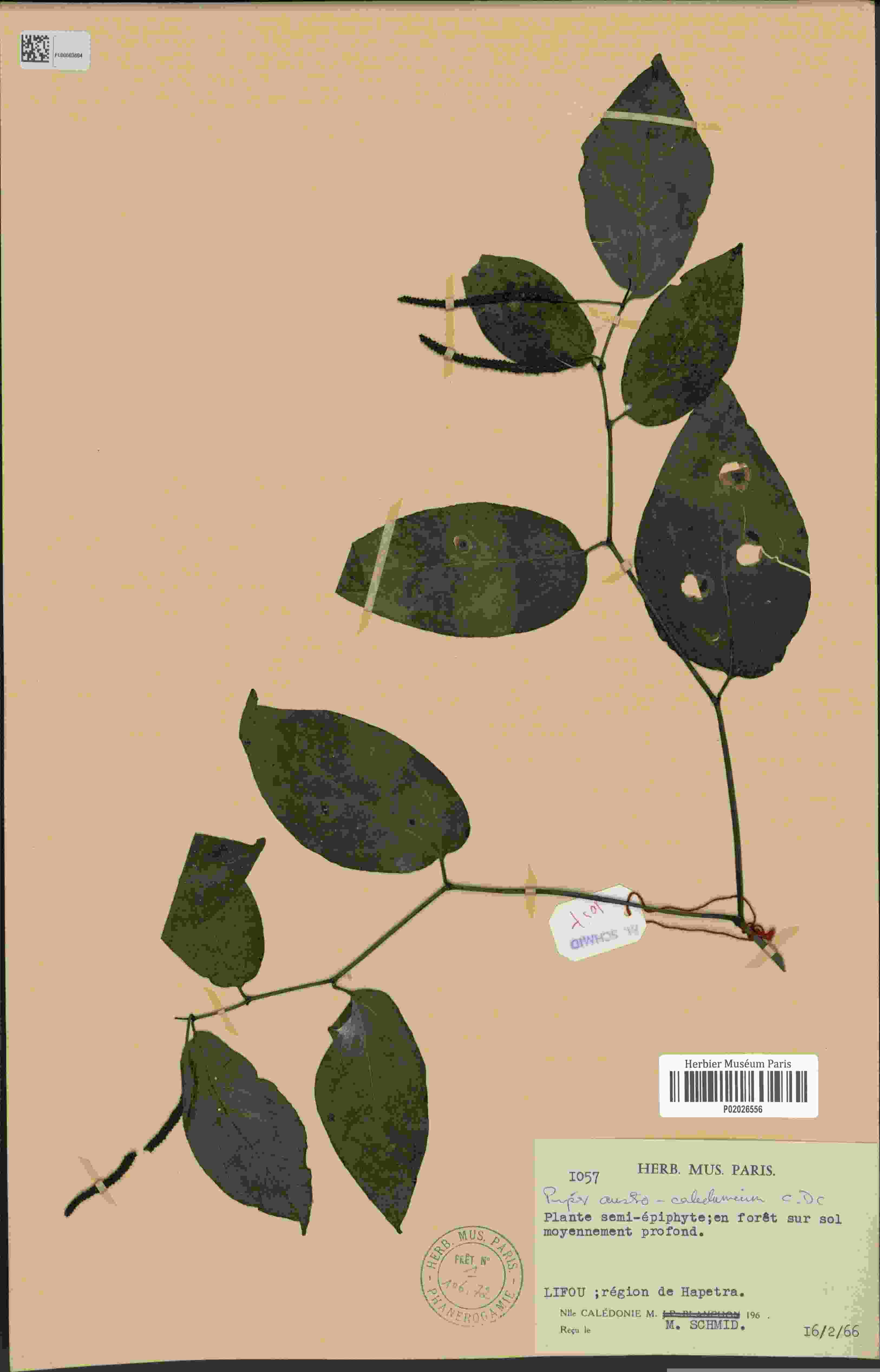

### Piper_staminodiferum_438834721.jpeg

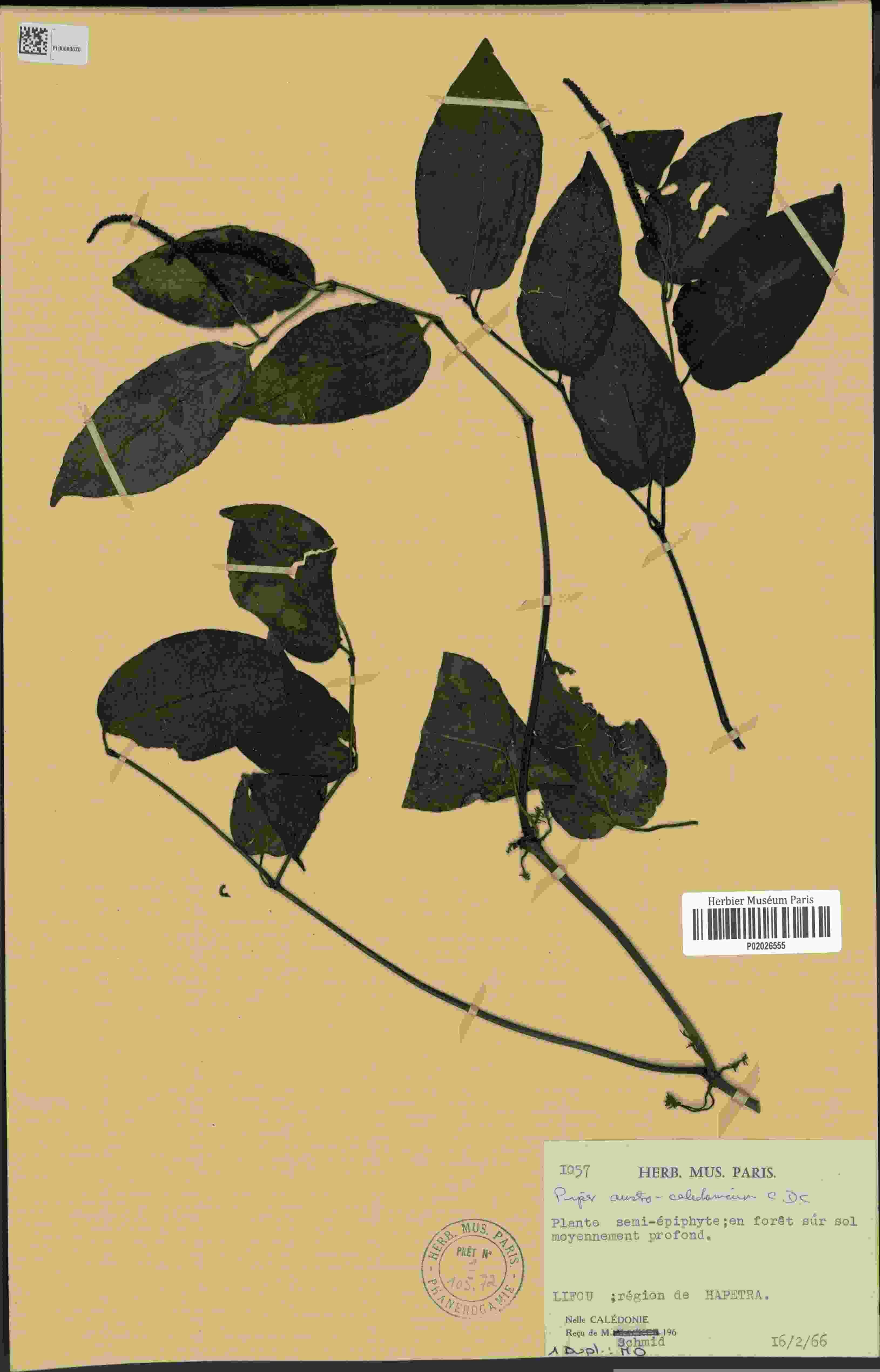

### Piper_staminodiferum_2575318271.jpeg

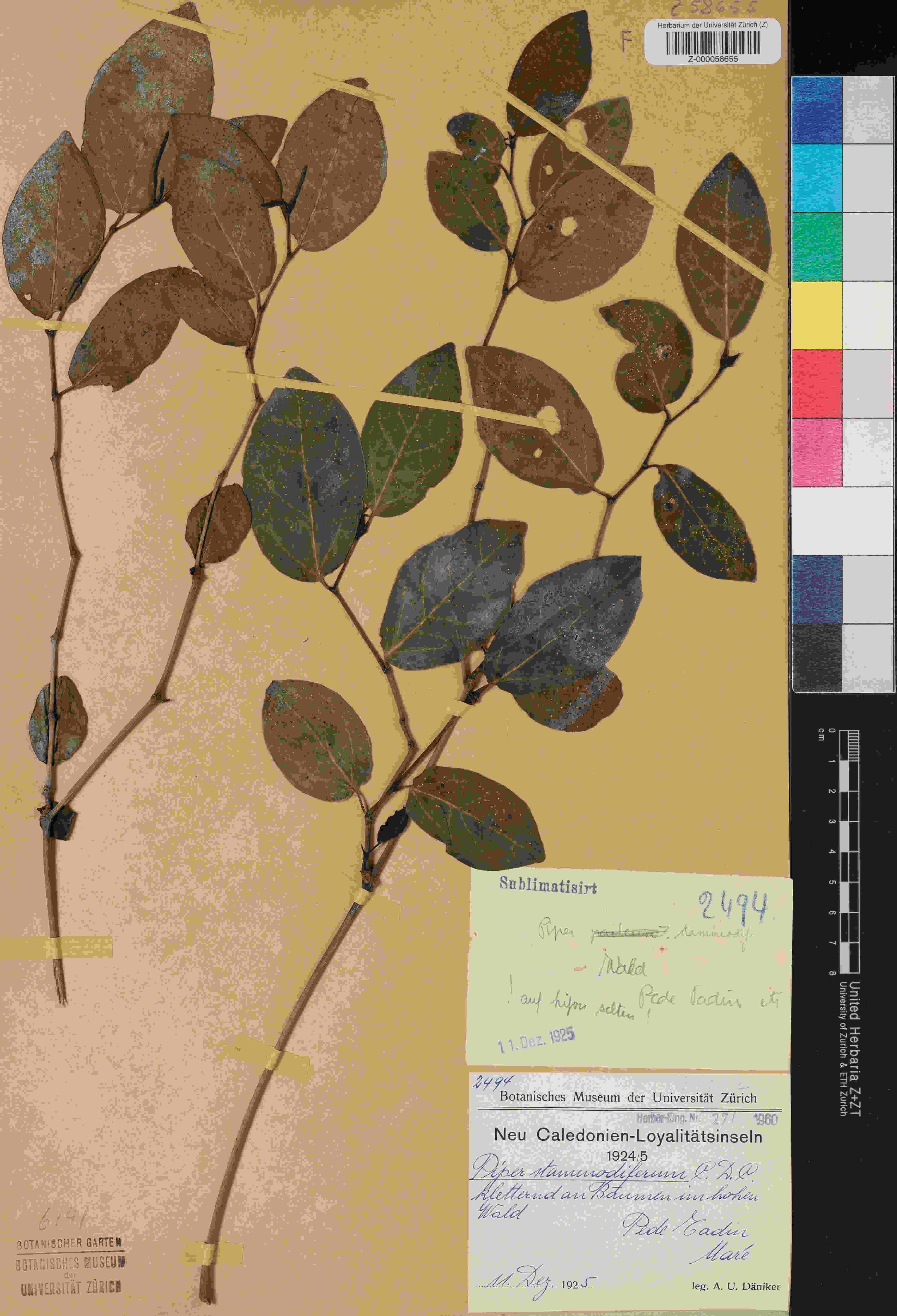

### Piper_staminodiferum_4031847283.jpeg

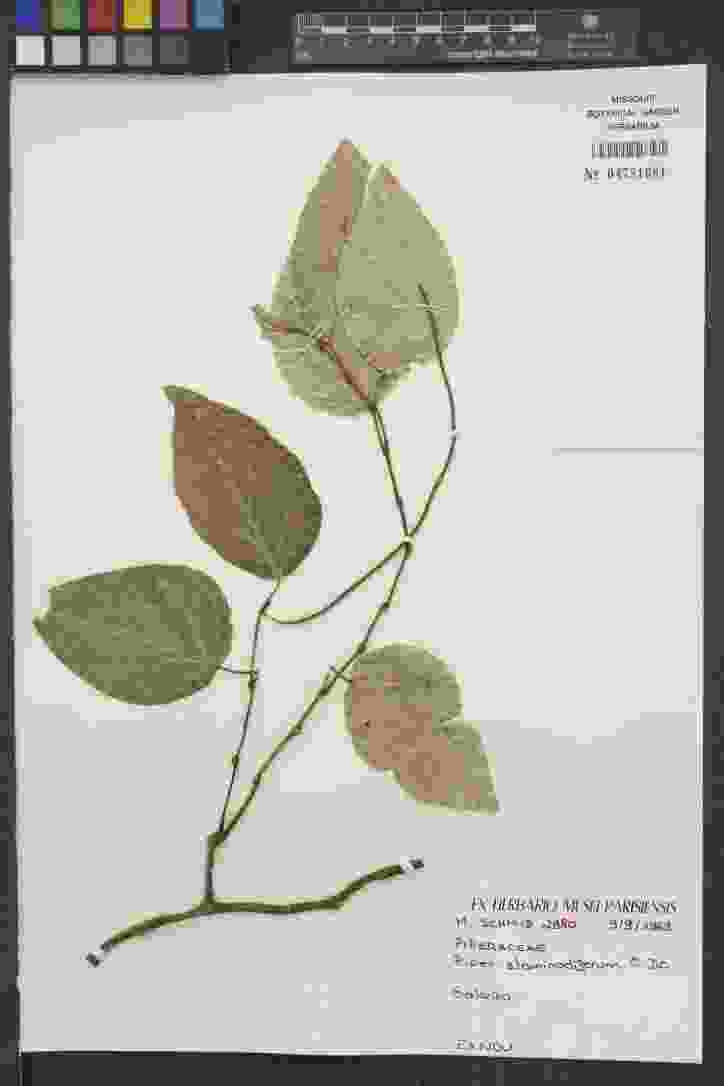

### Pithecellobium_lanceolatum_1258177064.jpeg

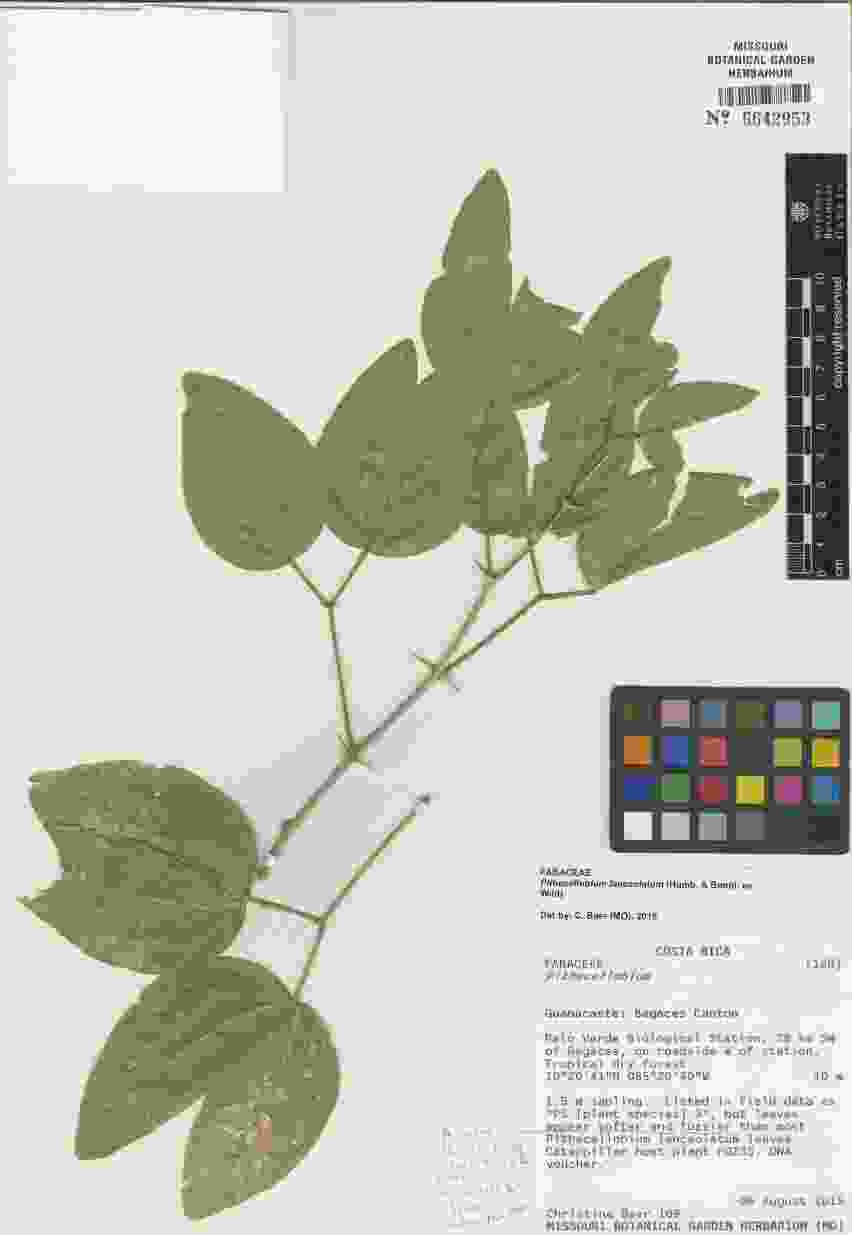

### Pithecellobium_lanceolatum_4441028404_2.jpeg

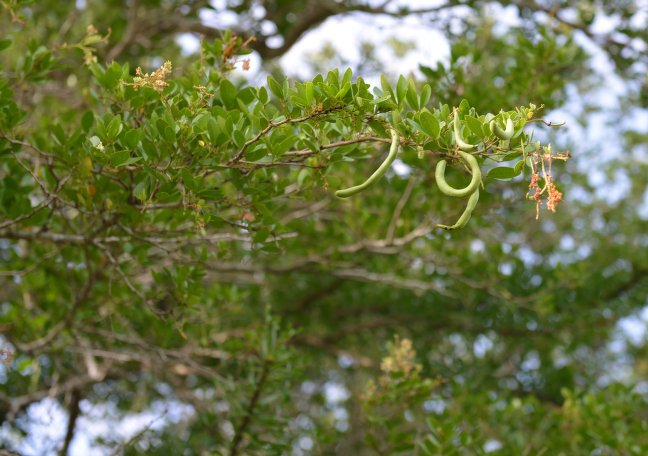

### Pithecellobium_lanceolatum_4441029617.jpeg

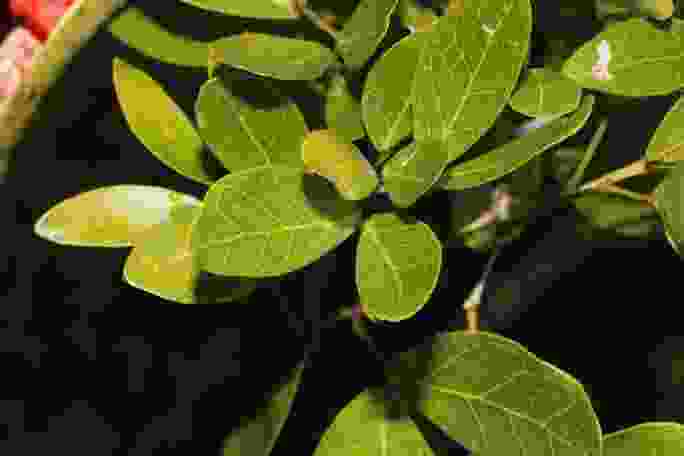

### Pithecellobium_lanceolatum_4441029617_2.jpeg

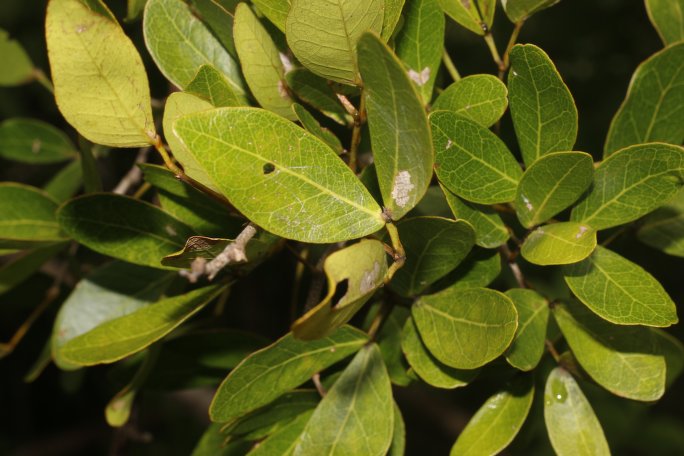

### Pithecellobium_lanceolatum_4441029617_3.jpeg

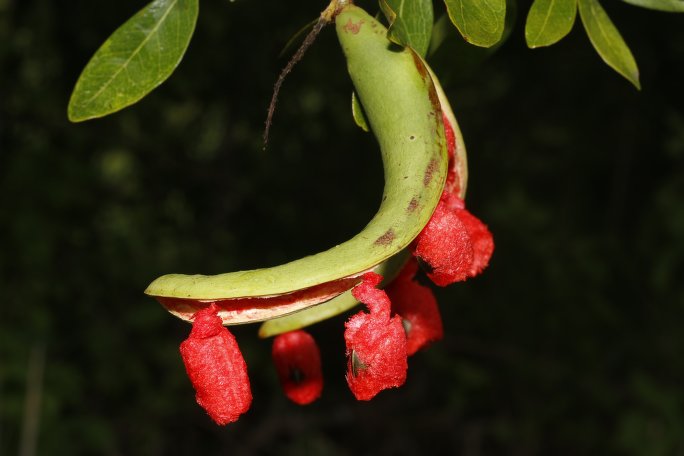

### Pithecellobium_lanceolatum_4441029617_5.jpeg

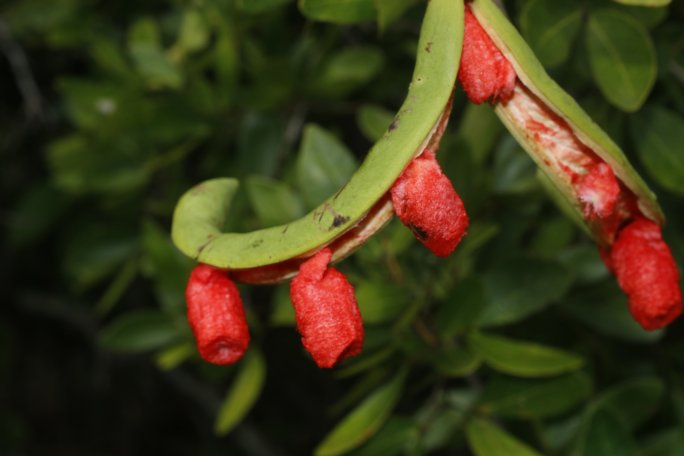

### Pithecellobium_lanceolatum_4441029617_8.jpeg

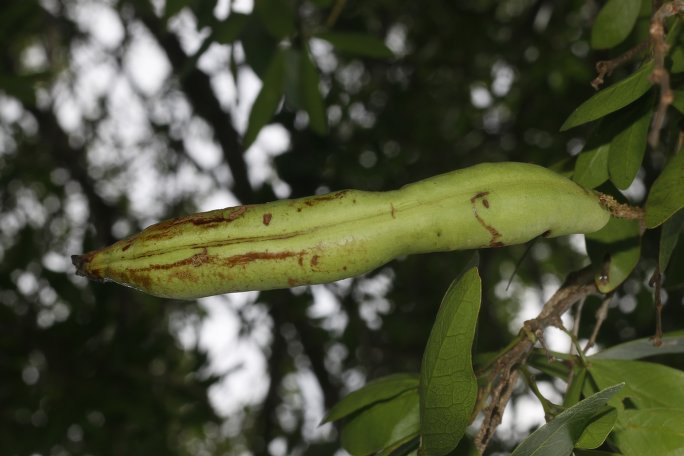

### Pithecellobium_lanceolatum_4441029617_9.jpeg

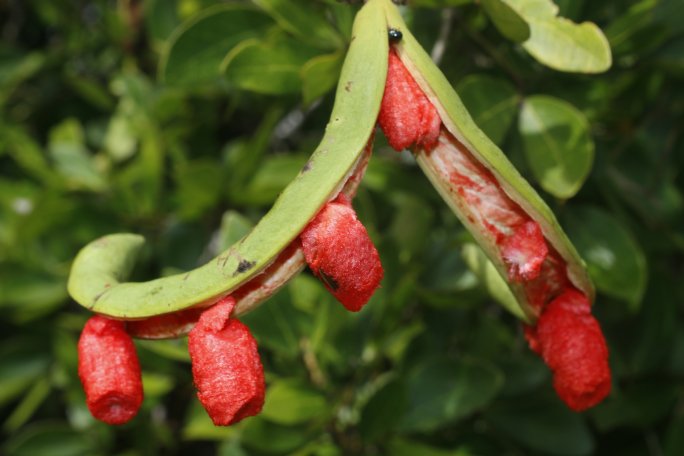

### Pithecellobium_lanceolatum_4441029617_12.jpeg

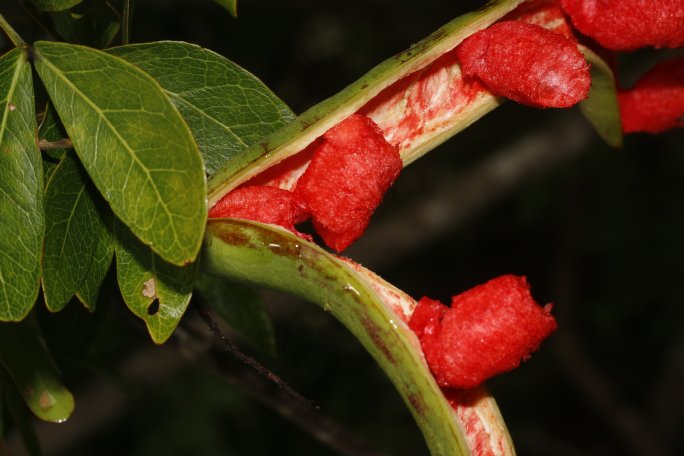

### Pithecellobium_lanceolatum_4441029617_13.jpeg

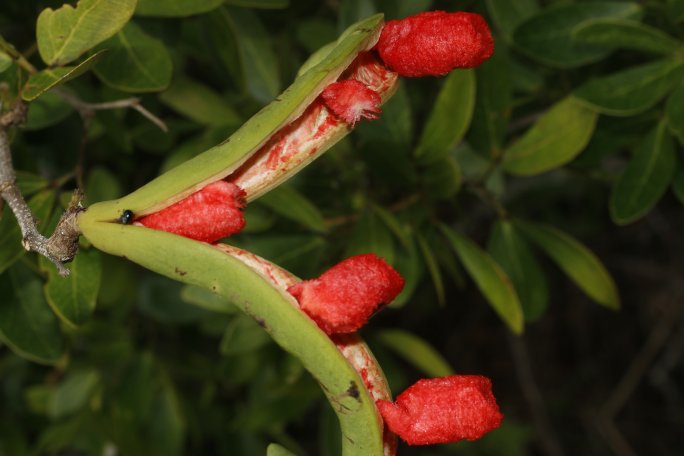

### Pithecellobium_lanceolatum_4441029617_14.jpeg

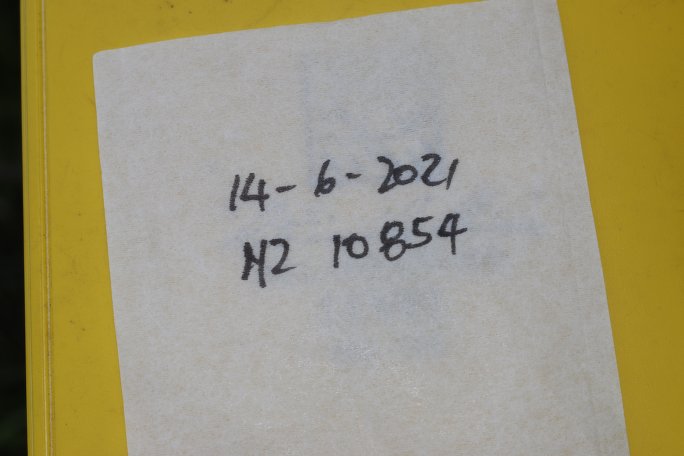

### Pithecellobium_lanceolatum_4441029617_15.jpeg

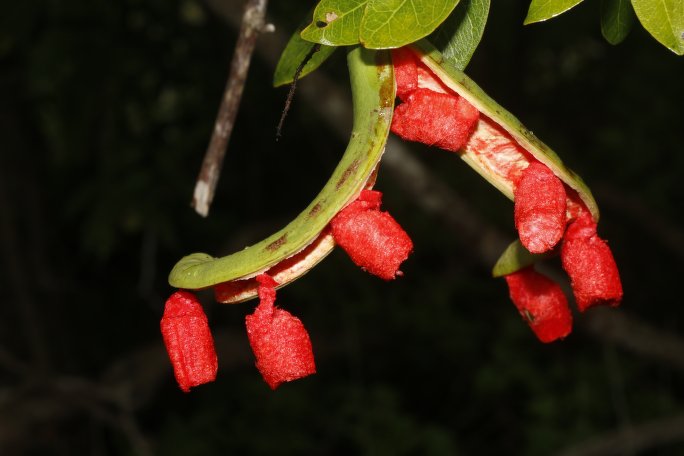

### Pithecellobium_lanceolatum_4441037521.jpeg

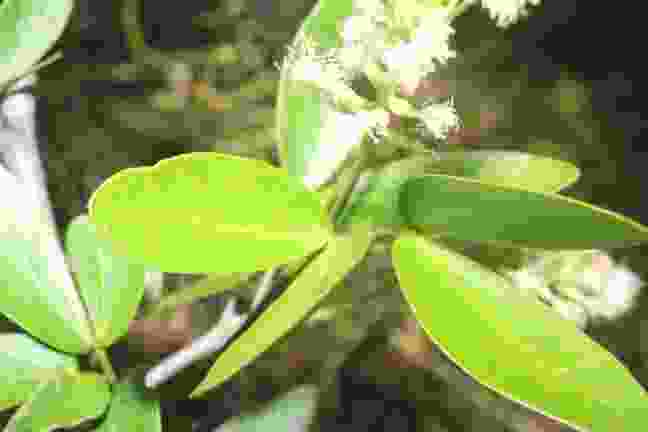

### Pithecellobium_lanceolatum_4441037521_1.jpeg

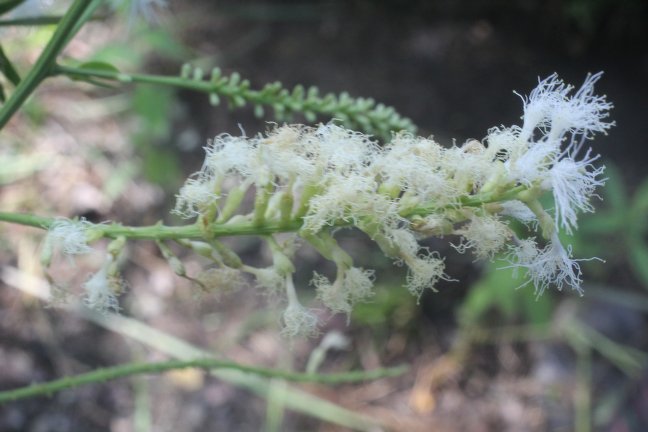

### Pithecellobium_lanceolatum_4441037521_2.jpeg

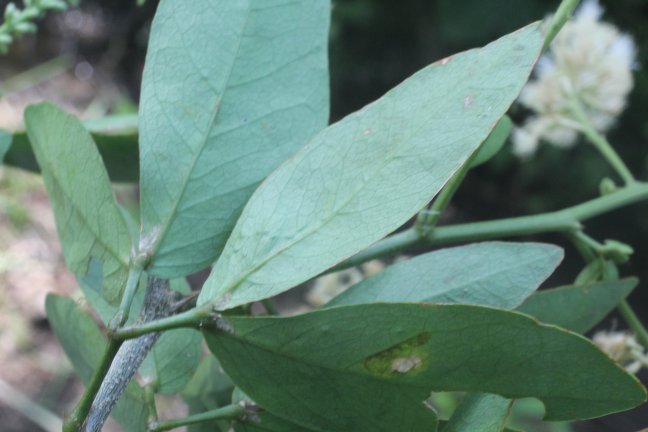

### Pithecellobium_lanceolatum_4441037521_6.jpeg

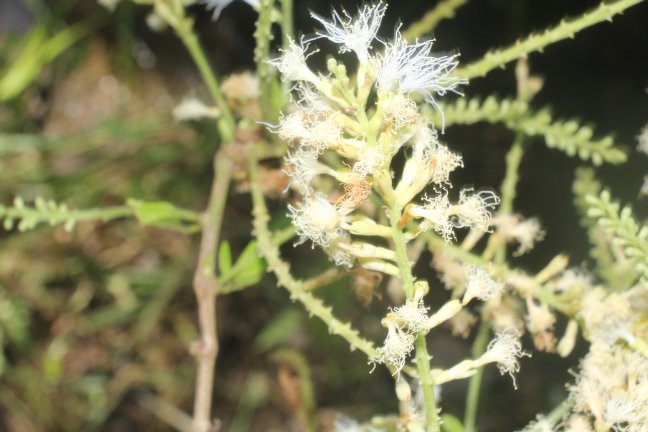

### Pithecellobium_lanceolatum_4441037521_7.jpeg

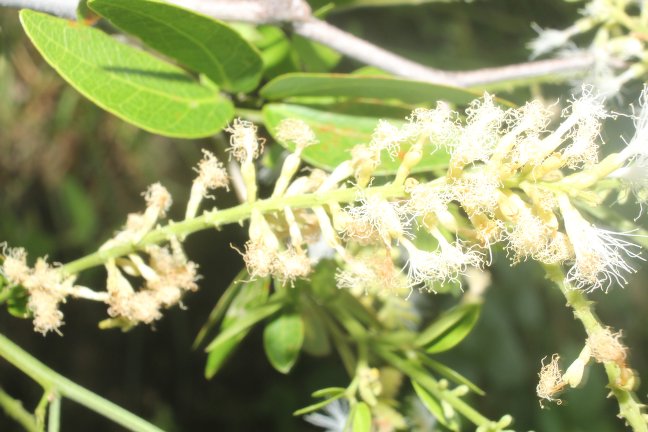

### Pithecellobium_lanceolatum_4441037521_10.jpeg

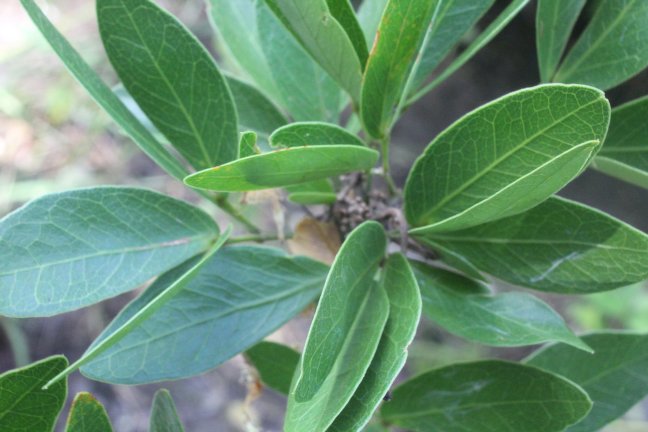

### Pithecellobium_lanceolatum_4441037521_11.jpeg

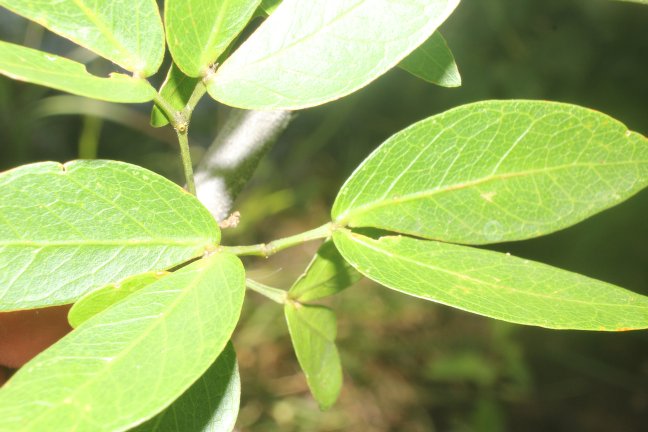

### Pithecellobium_lanceolatum_4441037521_13.jpeg

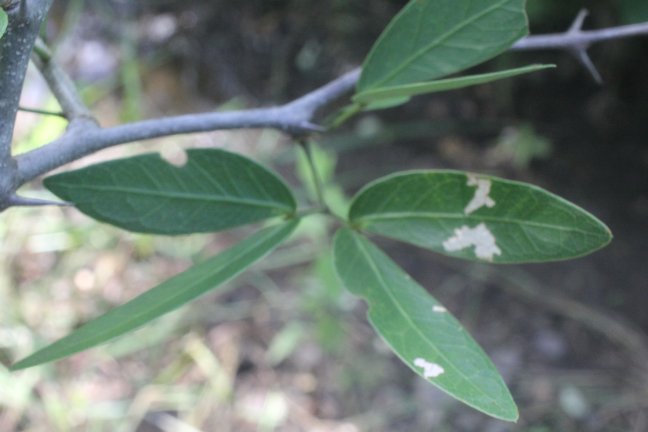

### Pithecellobium_lanceolatum_4441037521_16.jpeg

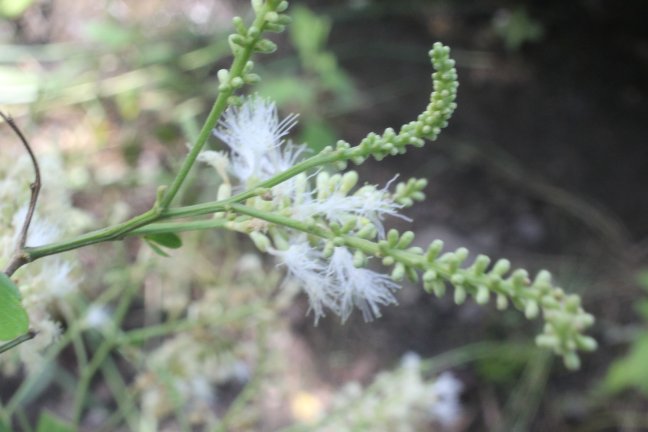

### Pithecellobium_lanceolatum_4441037521_17.jpeg

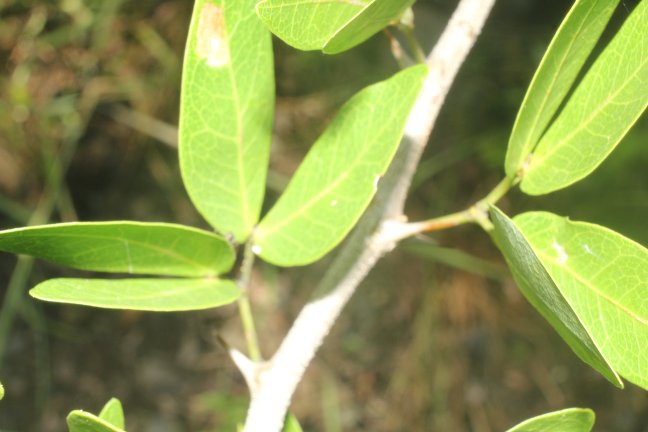

### Pithecellobium_lanceolatum_4441043496_1.jpeg

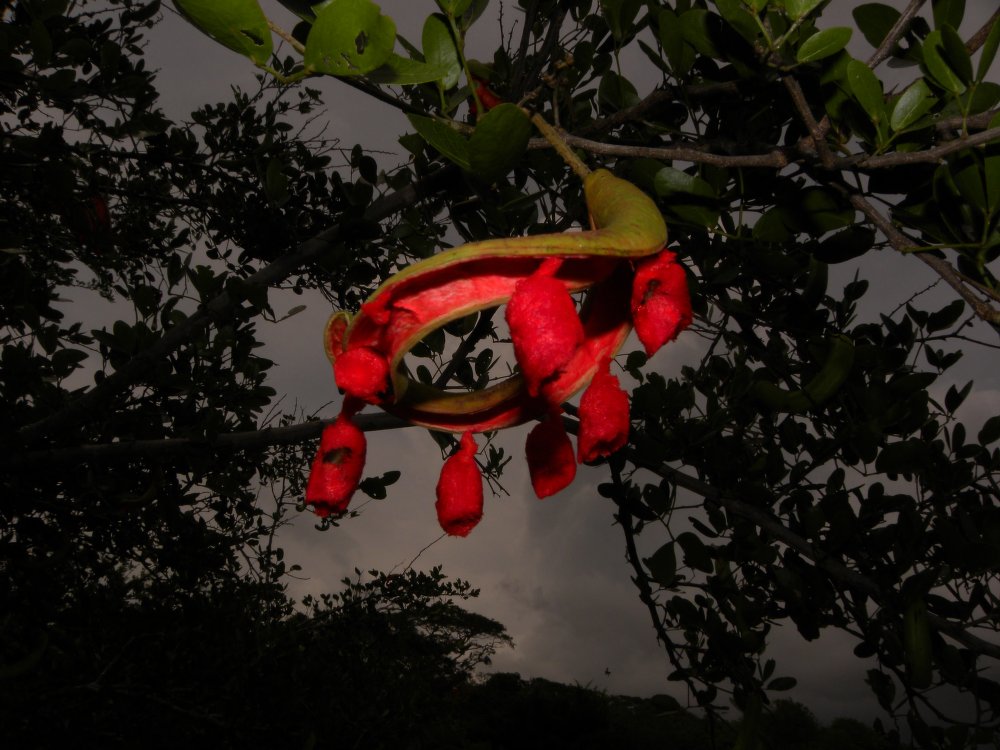

### Scilla_siberica_4065713402.jpeg

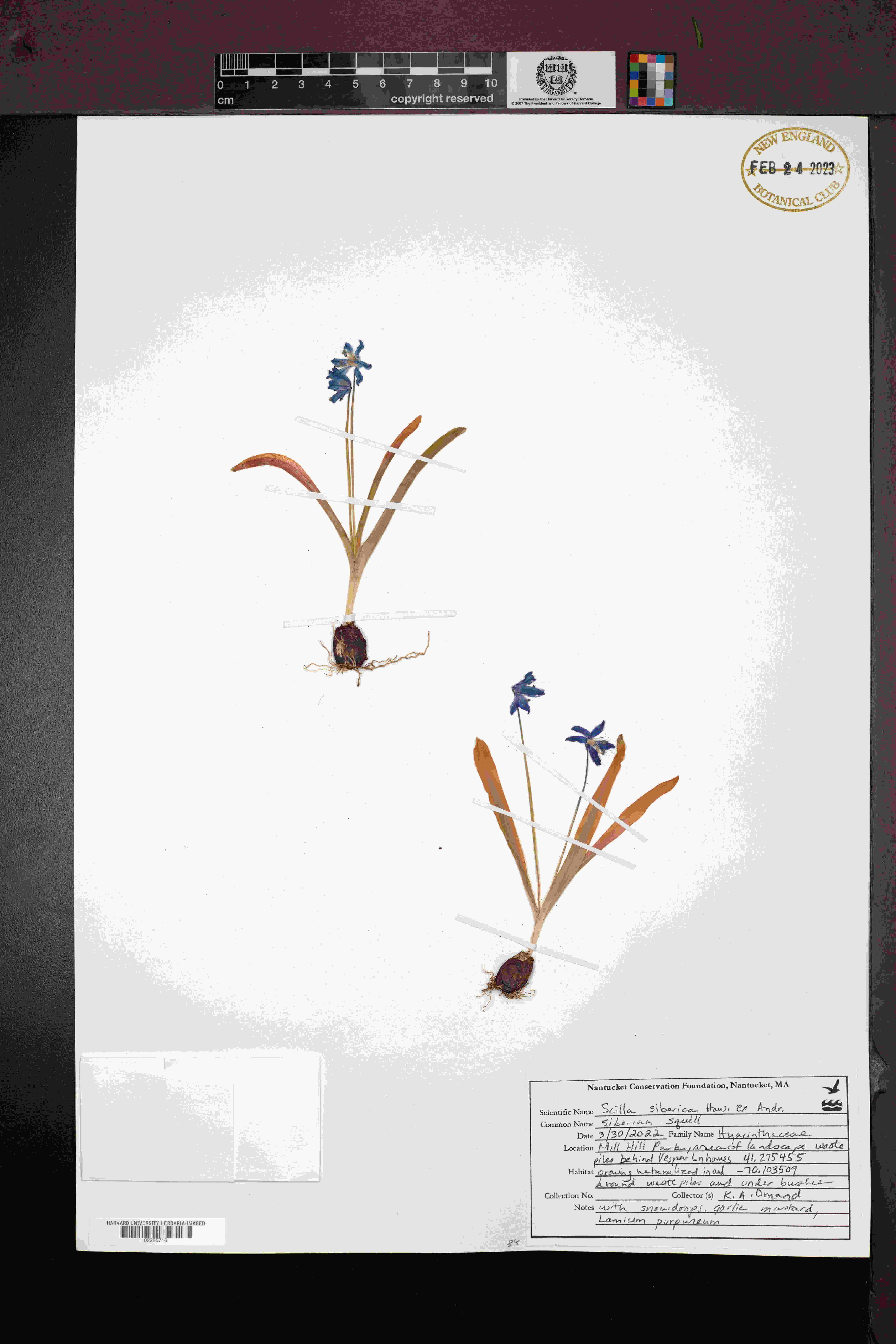

### Scilla_siberica_4849931752.jpeg

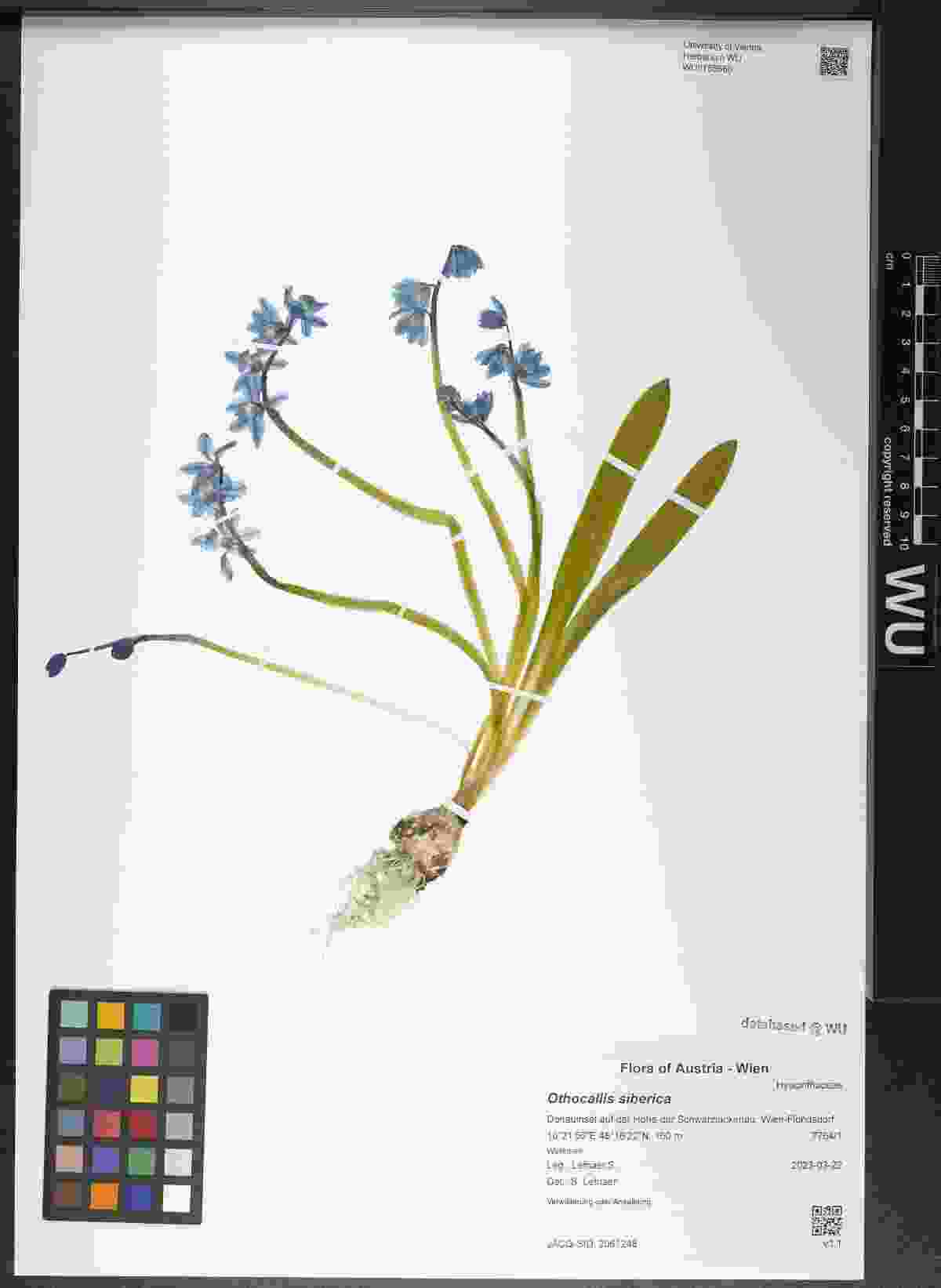

### Scilla_siberica_4856085645.jpeg

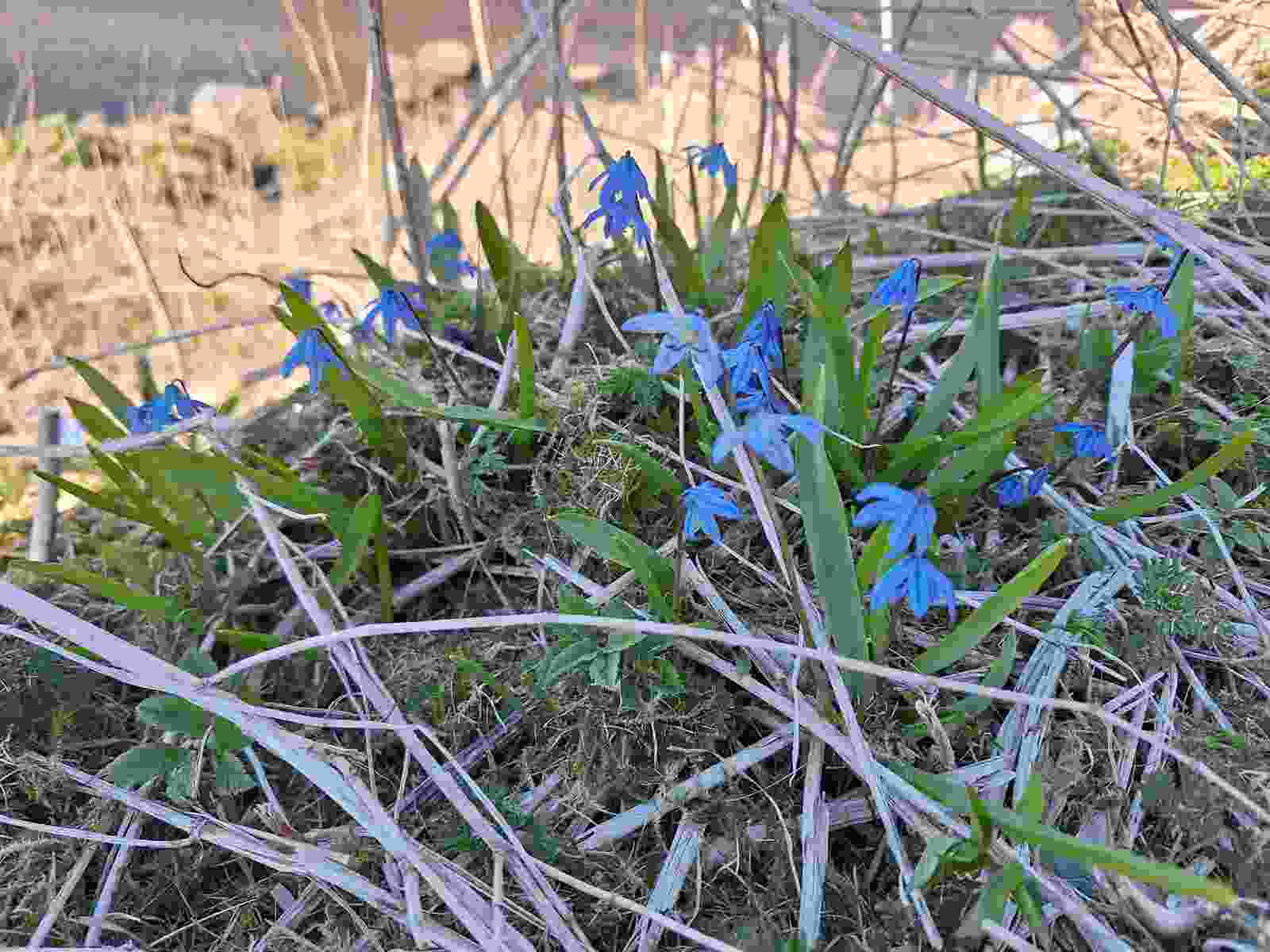

### Scilla_siberica_4856085645_1.jpeg

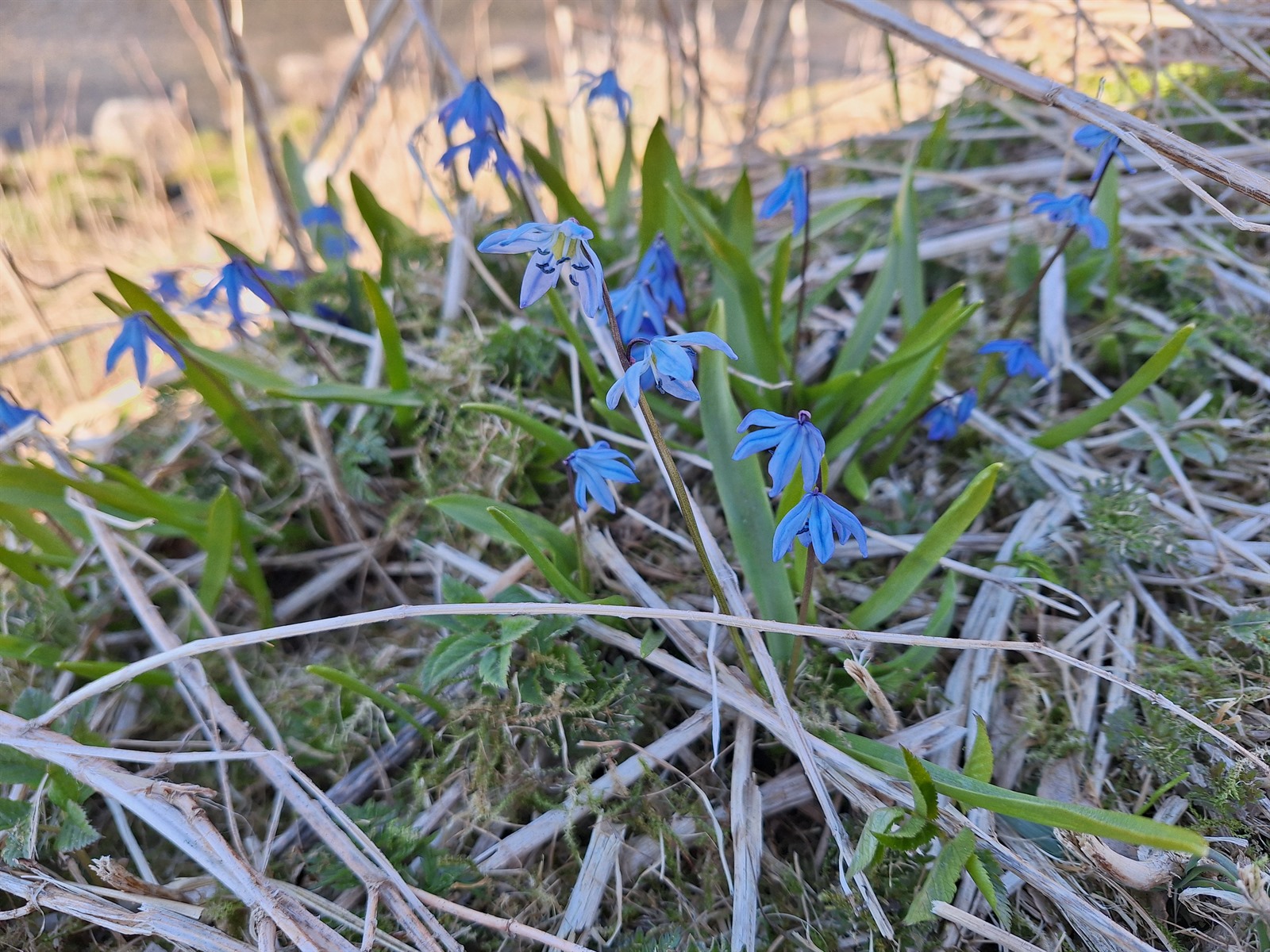

### Senecio_gallicus_2598787522.jpeg

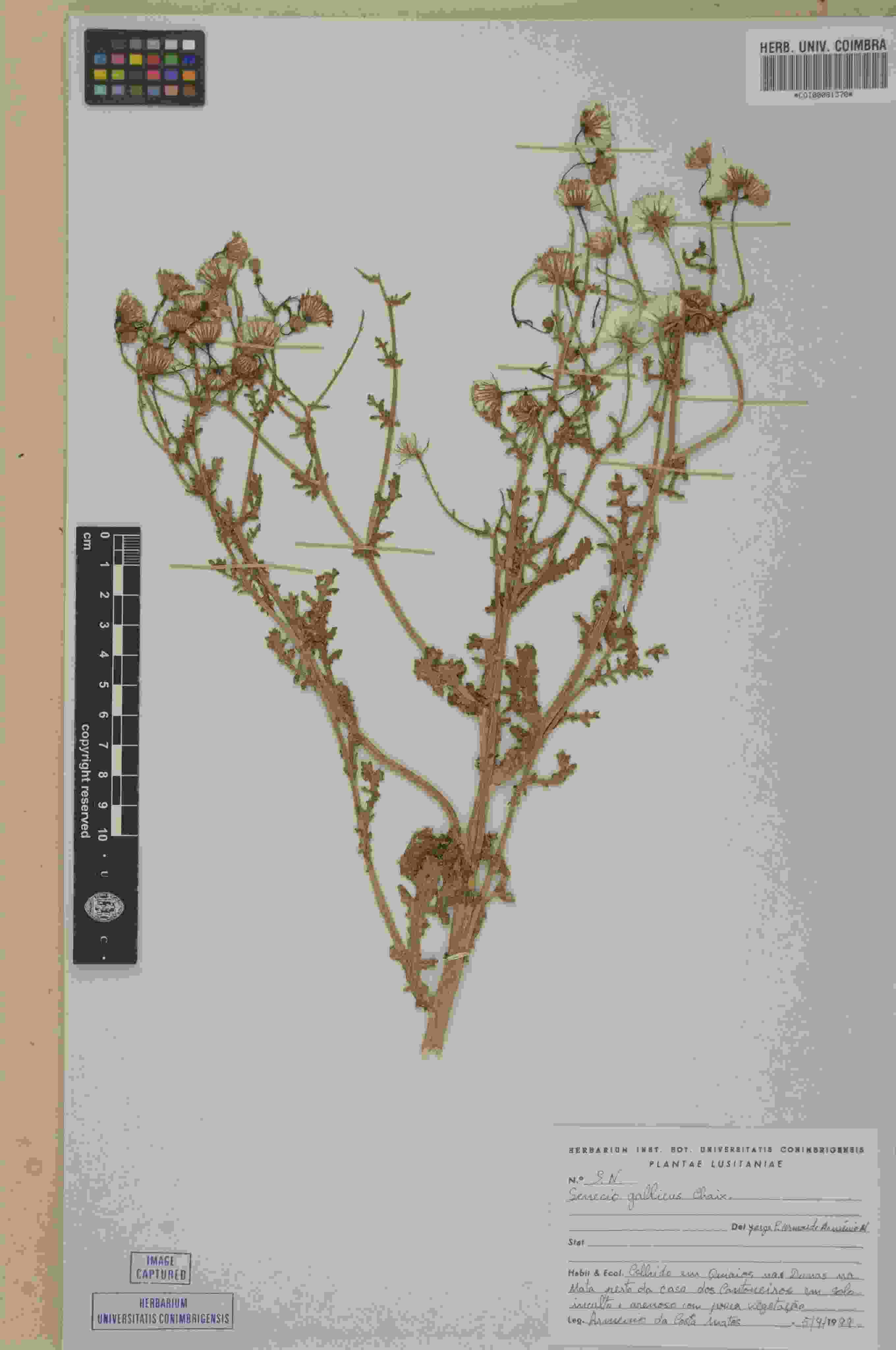

### Senecio_gallicus_2598867681.jpeg

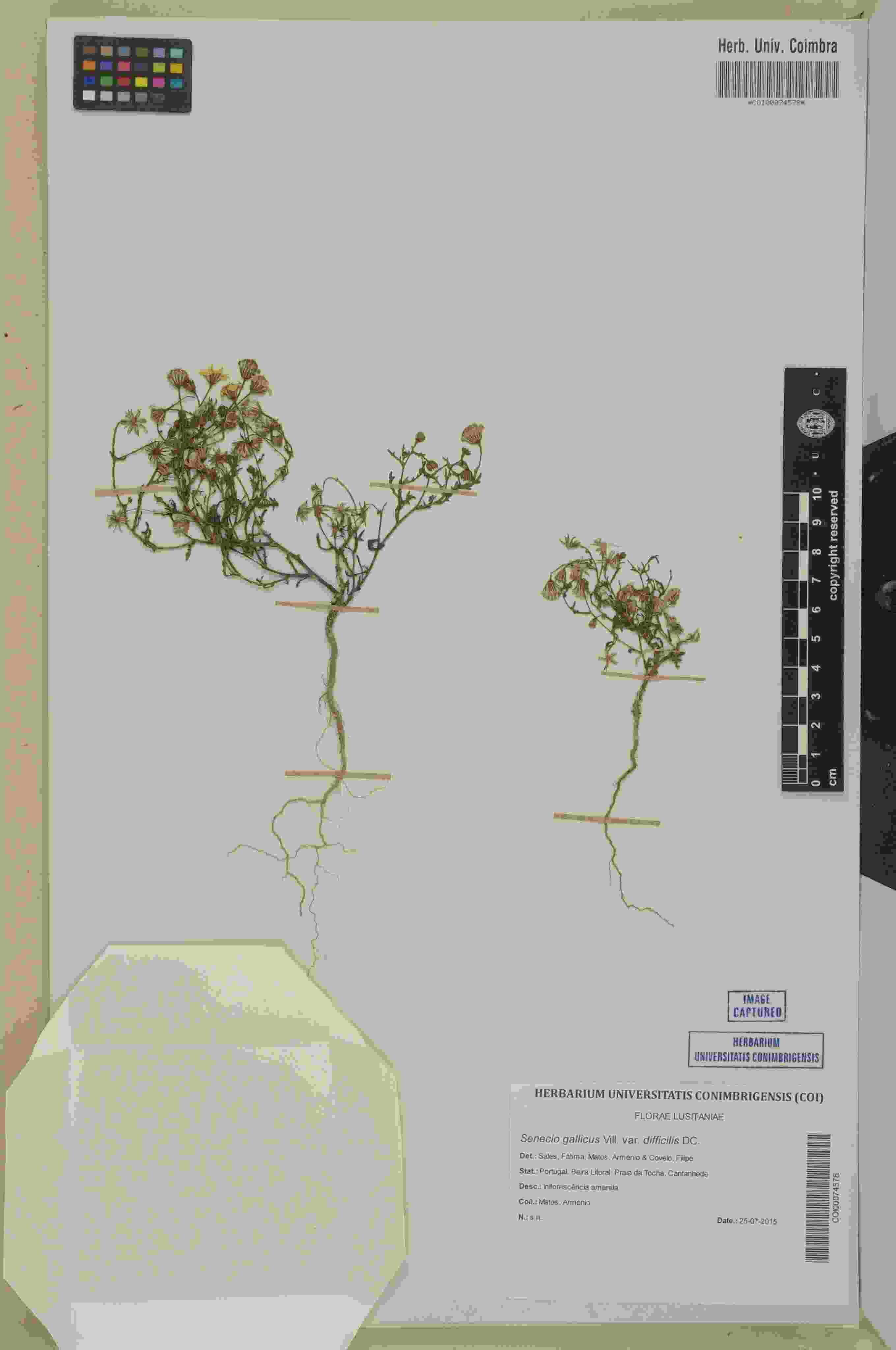

### Tridentopsis_buckleyana_2807501559.jpeg

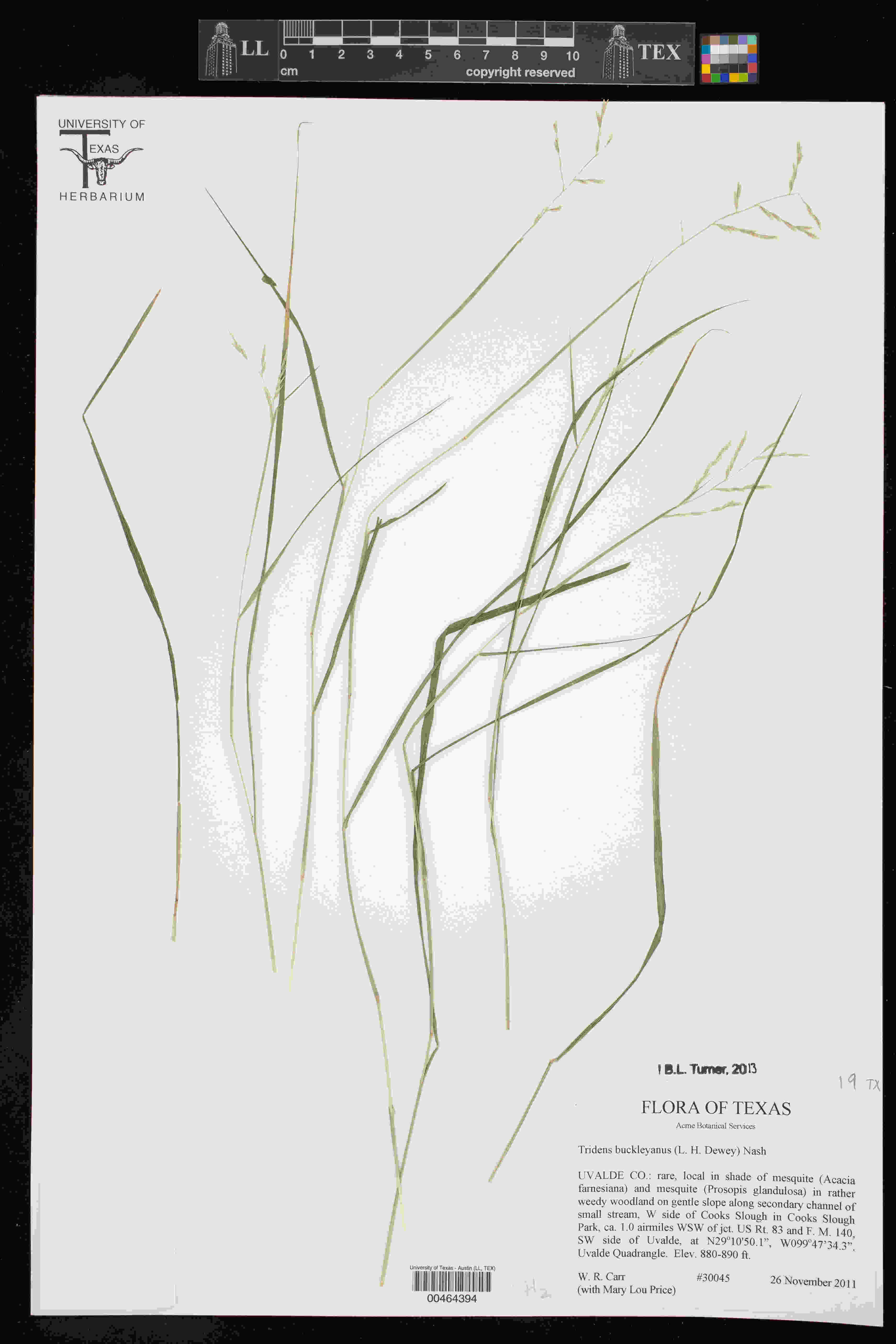

### Tridentopsis_buckleyana_3978705256.jpeg

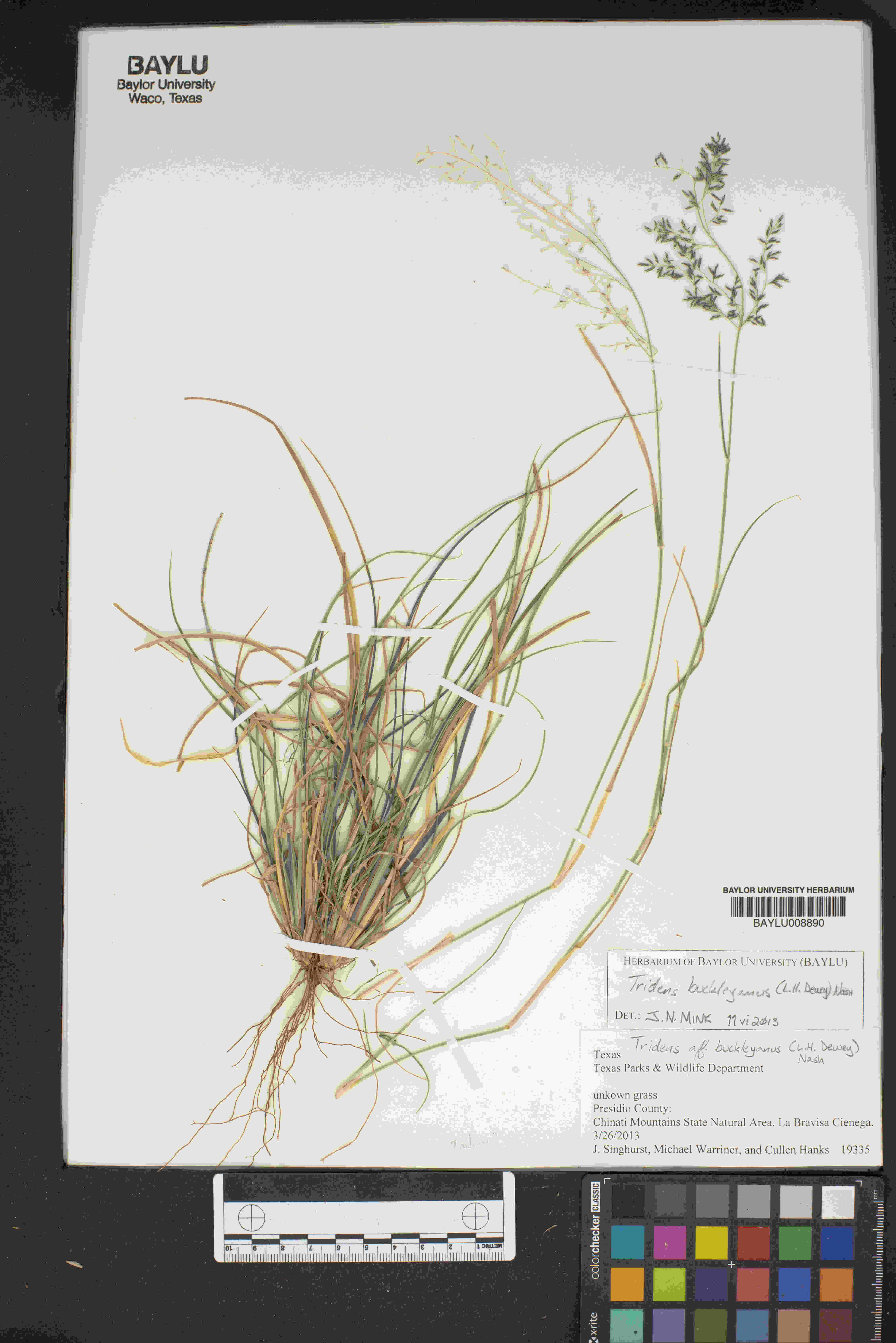

### Verbascum_austroiranicum_4524875316.jpeg

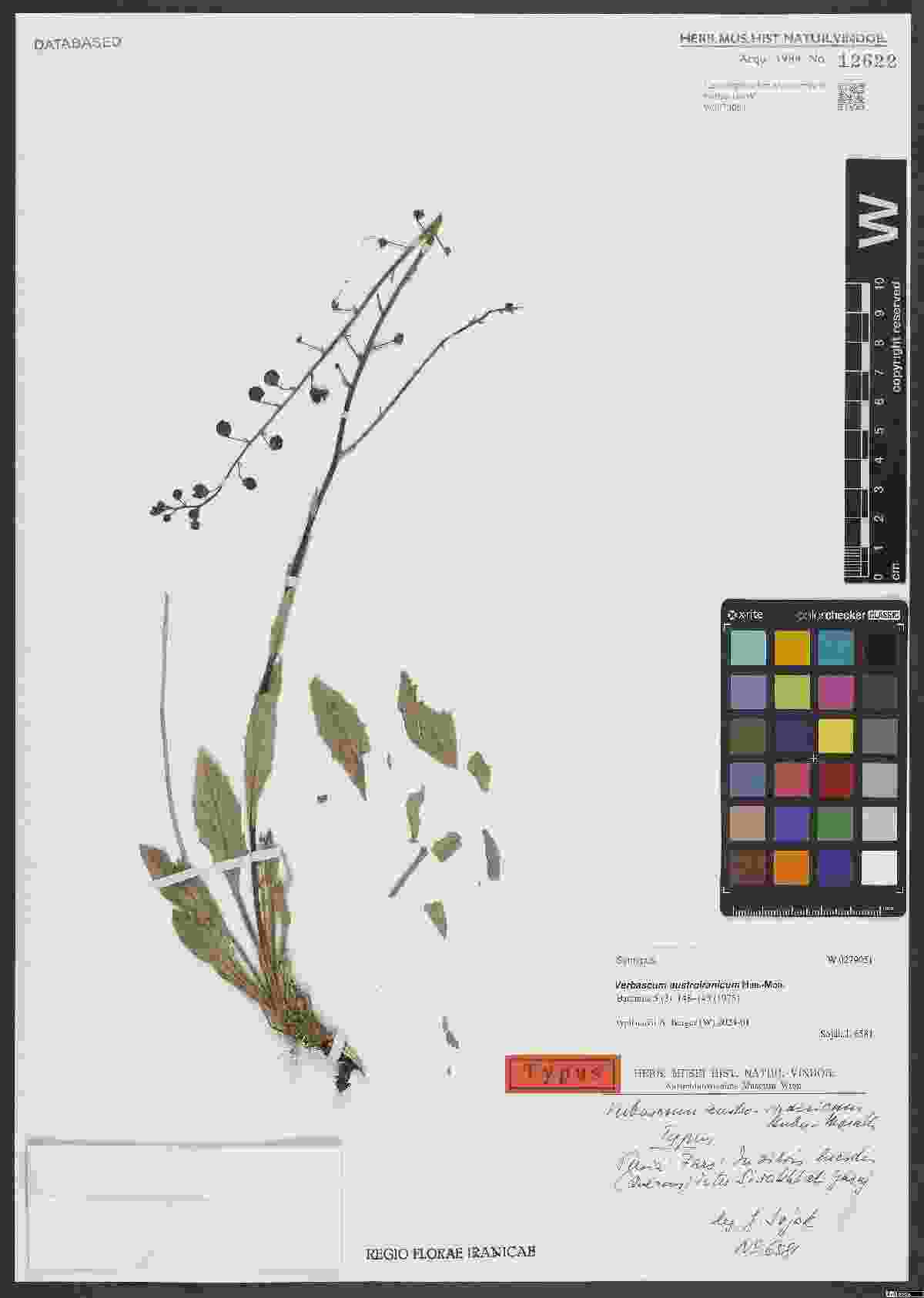

### Verbascum_austroiranicum_4524876321.jpeg

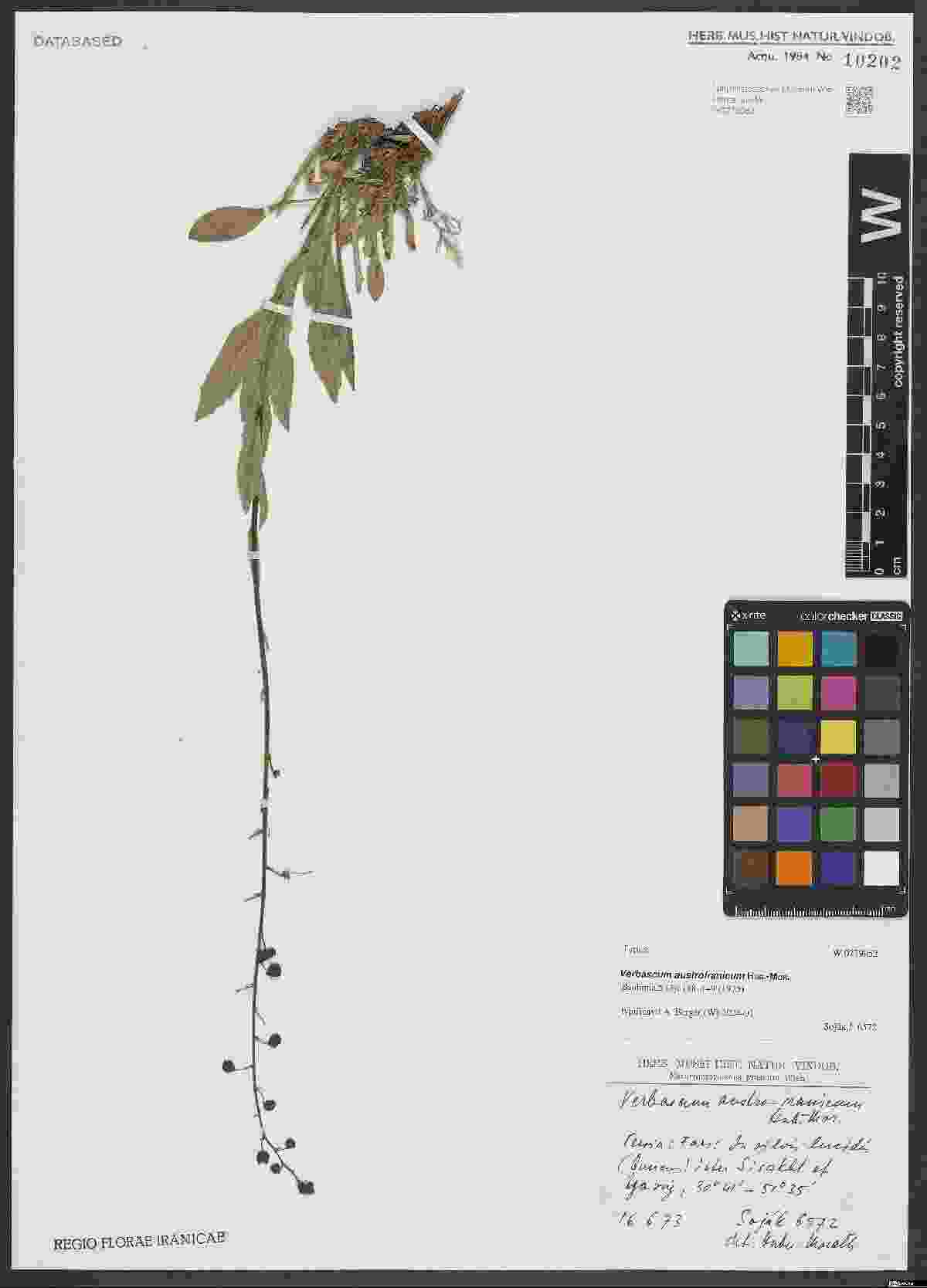

### Viscum_orientale_2515335796.jpeg

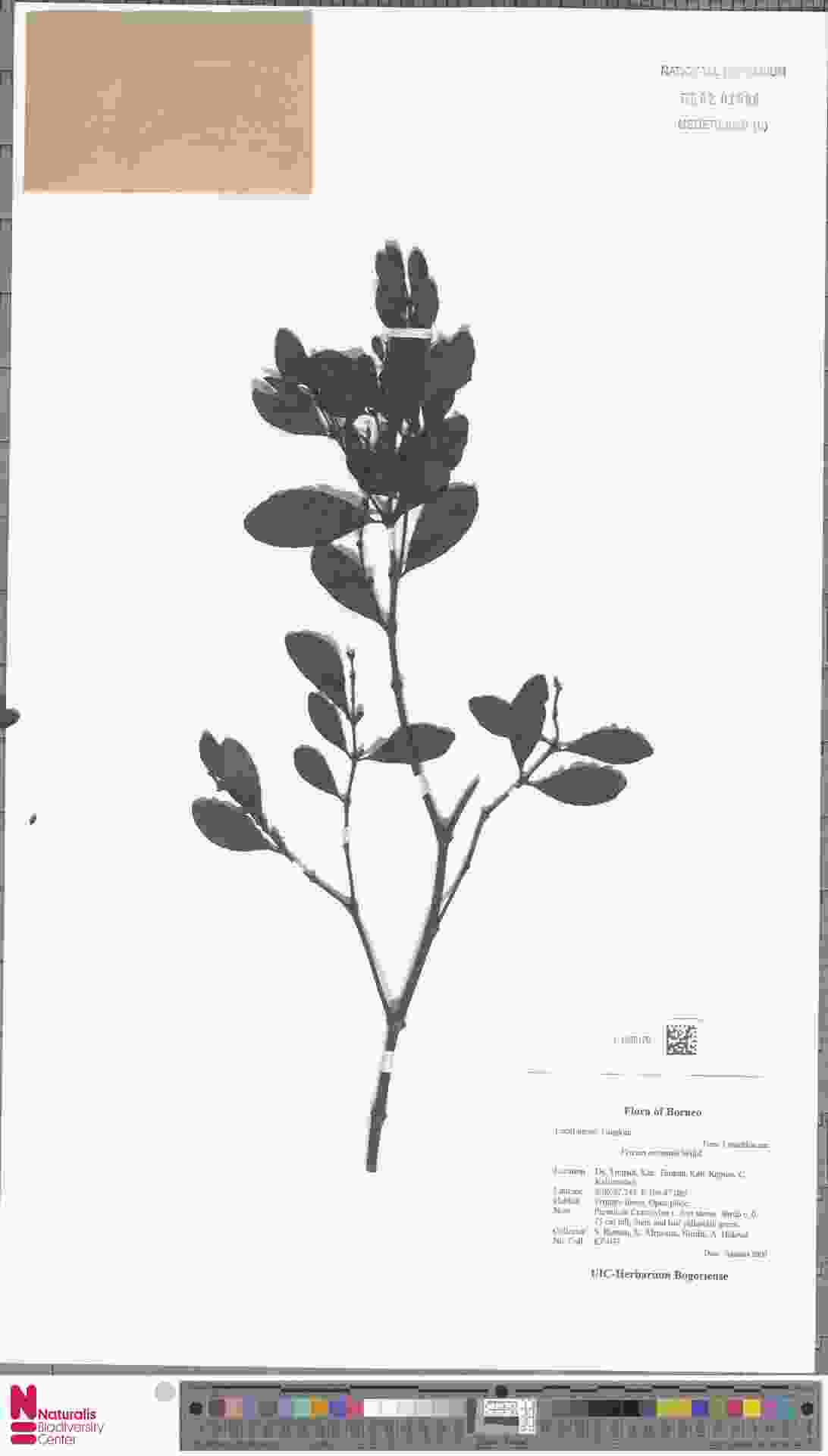
