## Supplementary figures and images for "*mvh*: an R tool to assemble and organize virtual herbaria from openly available specimen images"

### Aristida_kenyensis_1318076536.jpeg

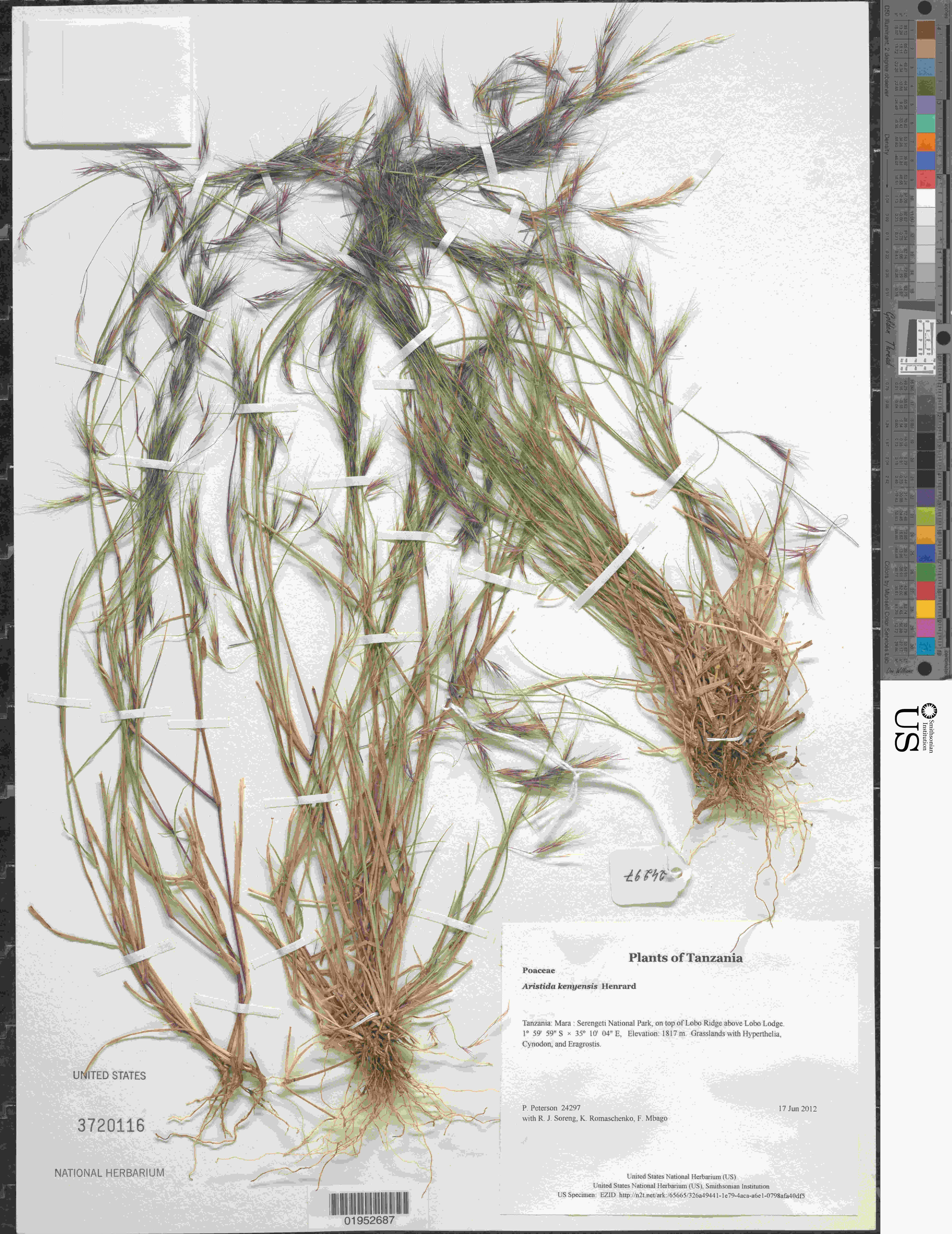

### Aristida_kenyensis_1319267269.jpeg

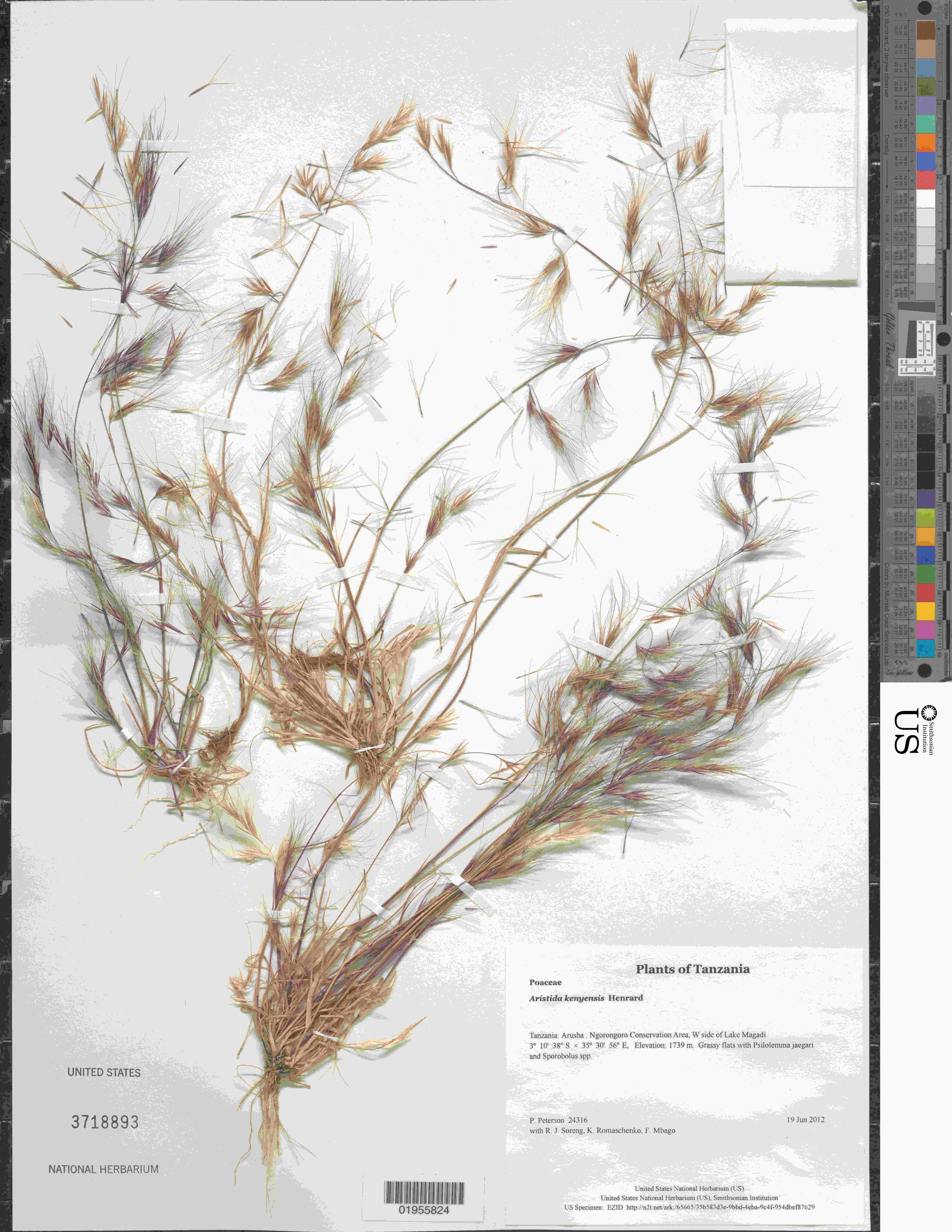

### Aristida_kenyensis_2268838537.jpeg

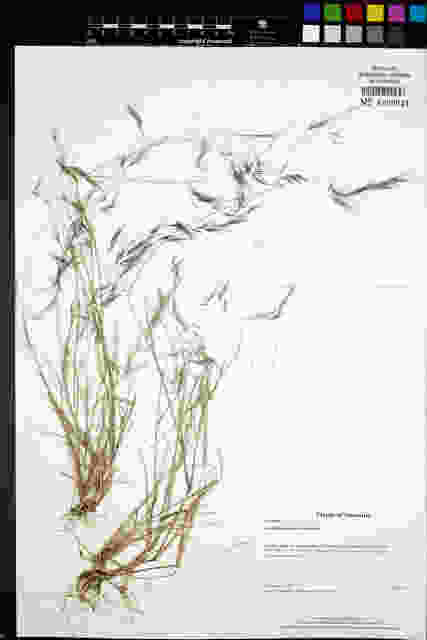

### Aristida_kenyensis_4849312049.jpeg

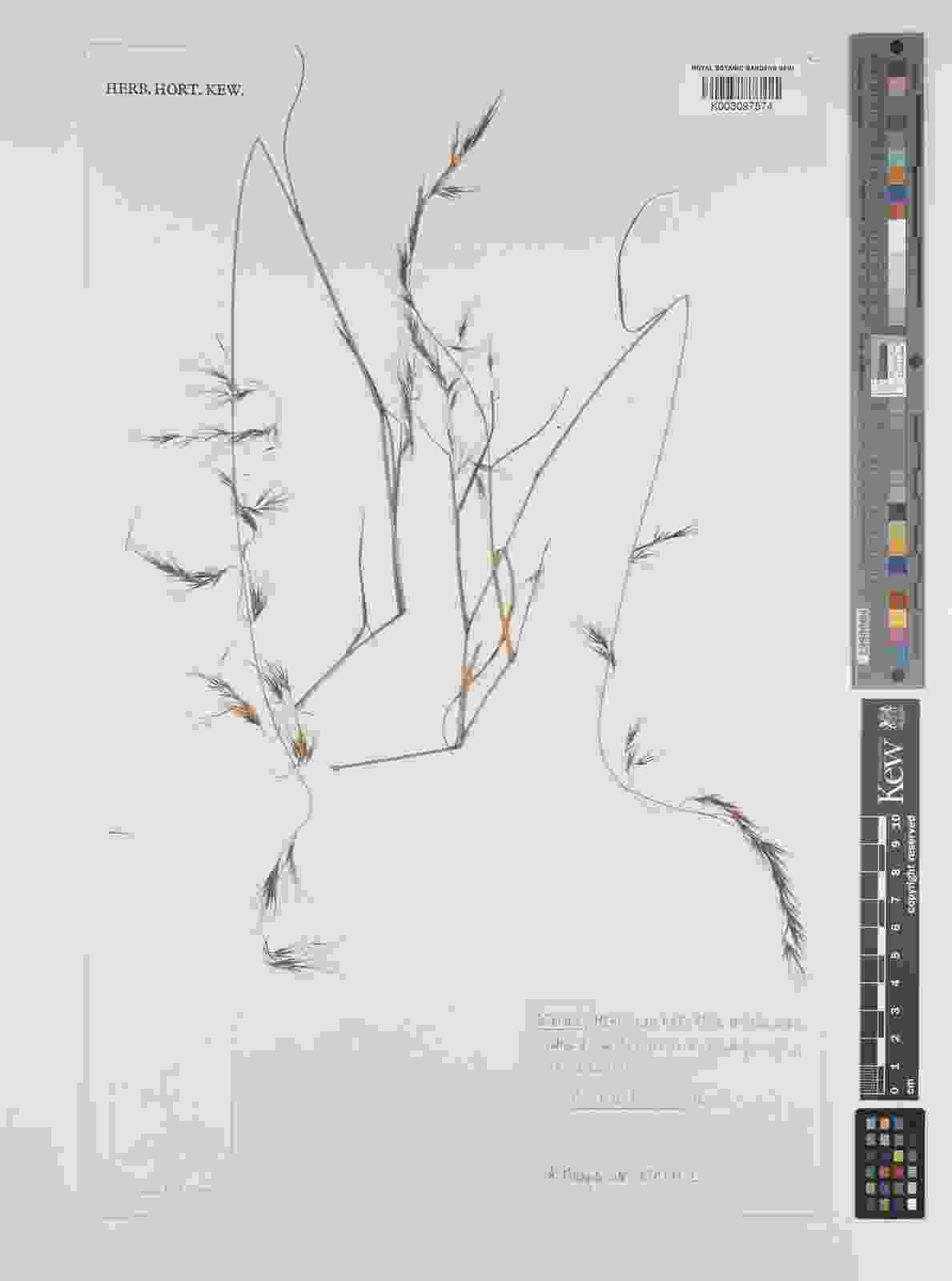

### Aristida_kenyensis_4853811829.jpeg

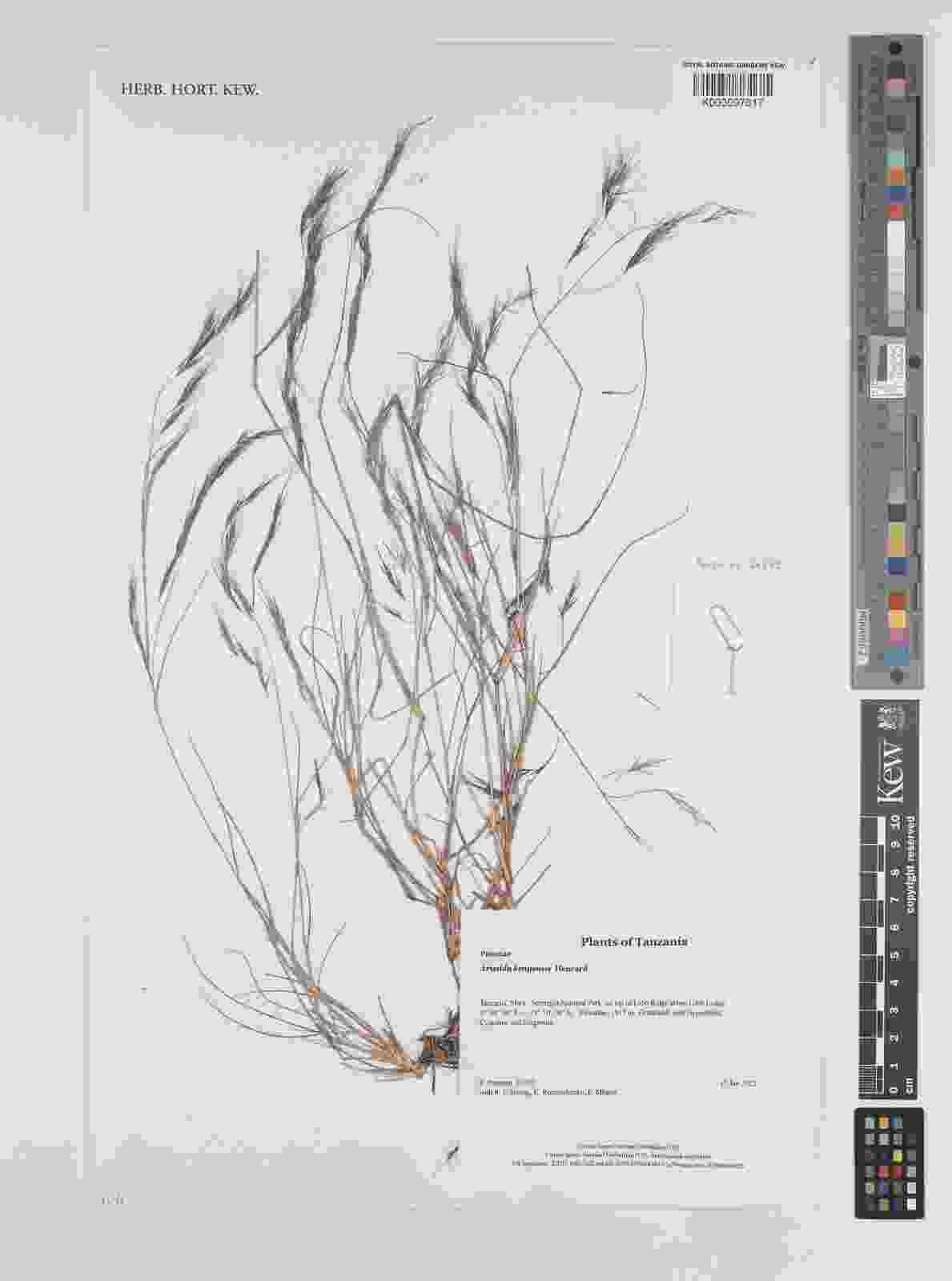

### Aristida_kenyensis_4853811829_1.jpeg

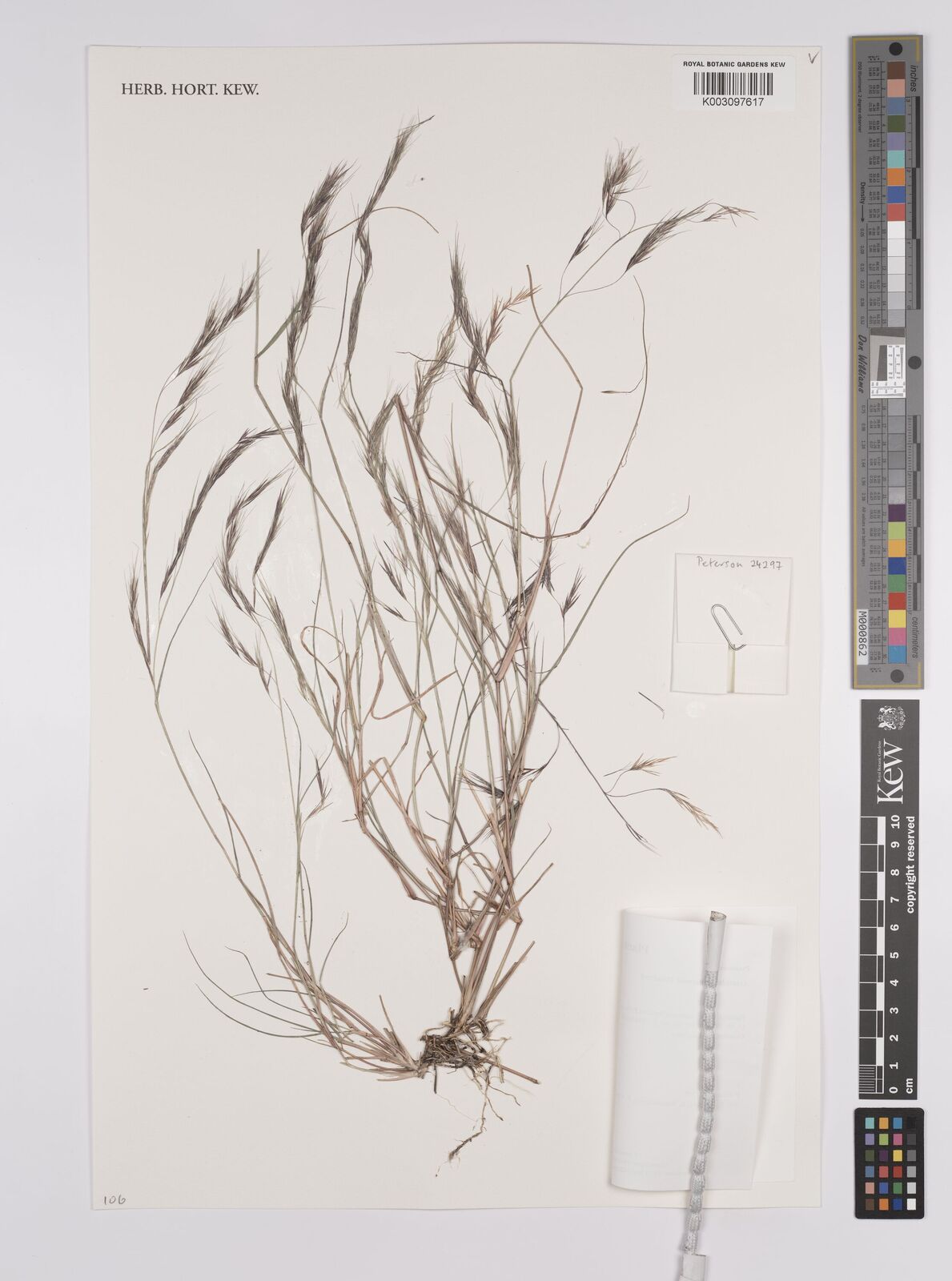

### Astragalus_aretioides_1852148593.jpeg

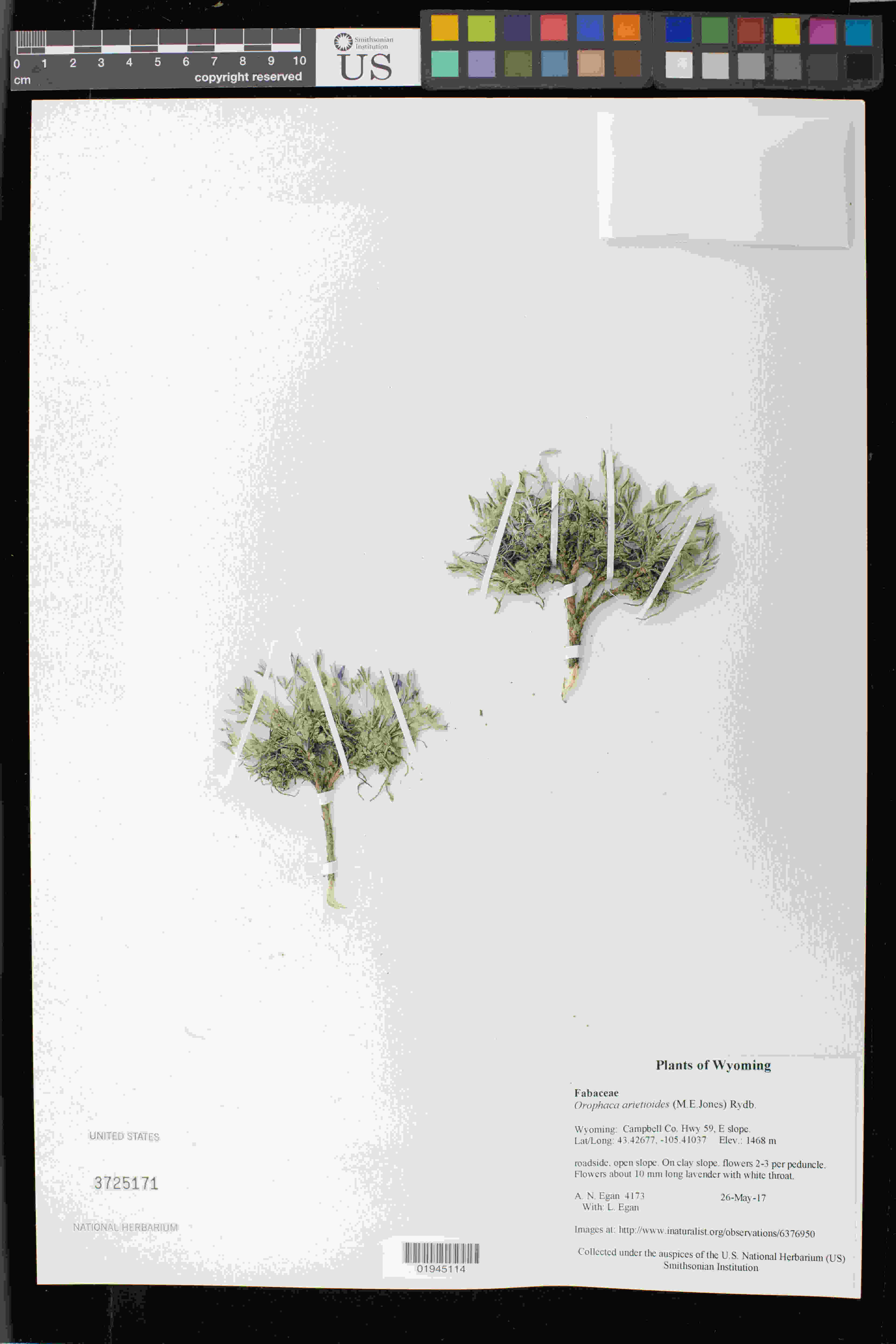

### Astragalus_aretioides_1929430021.jpeg

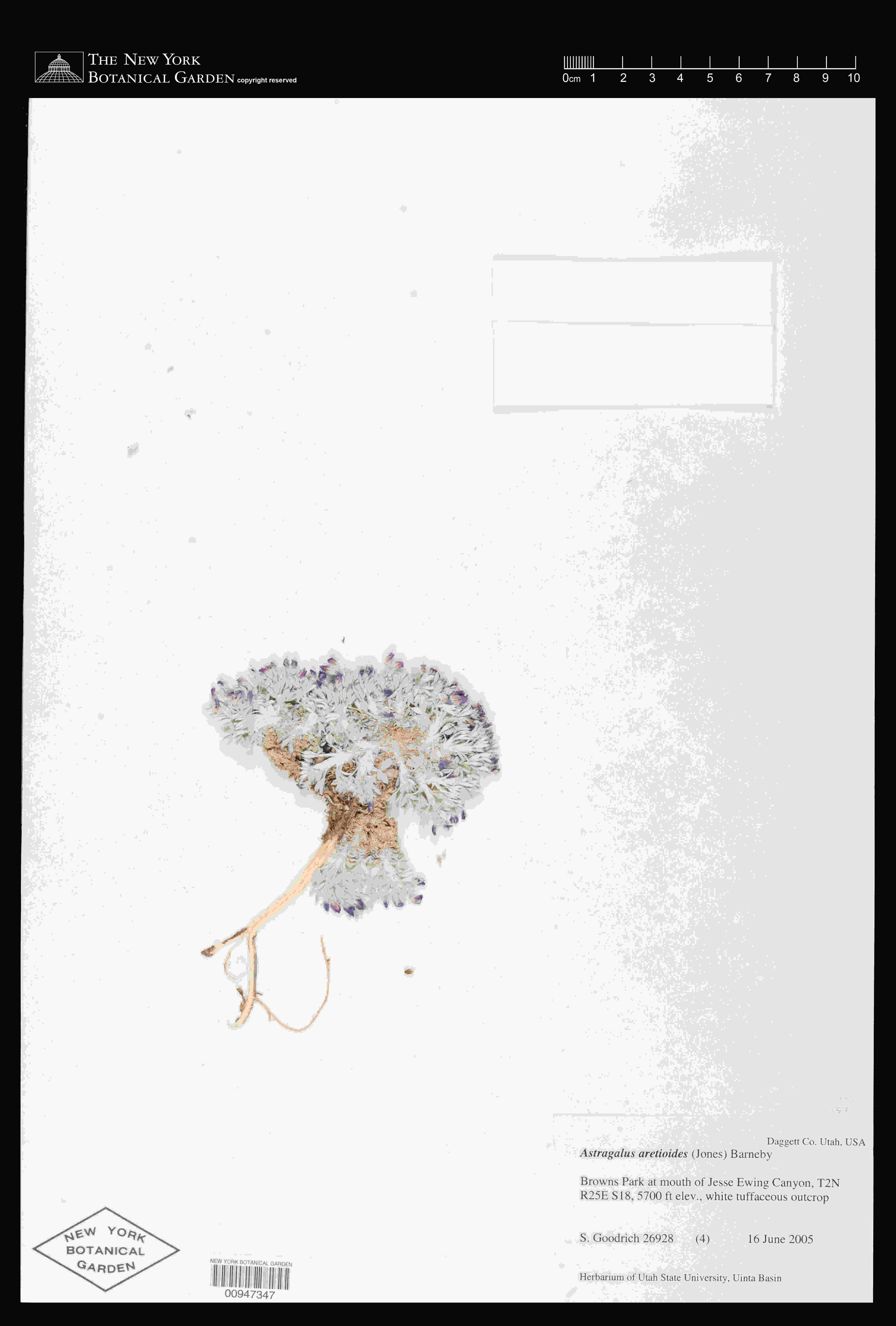

### Astragalus_aretioides_2573390729.jpeg

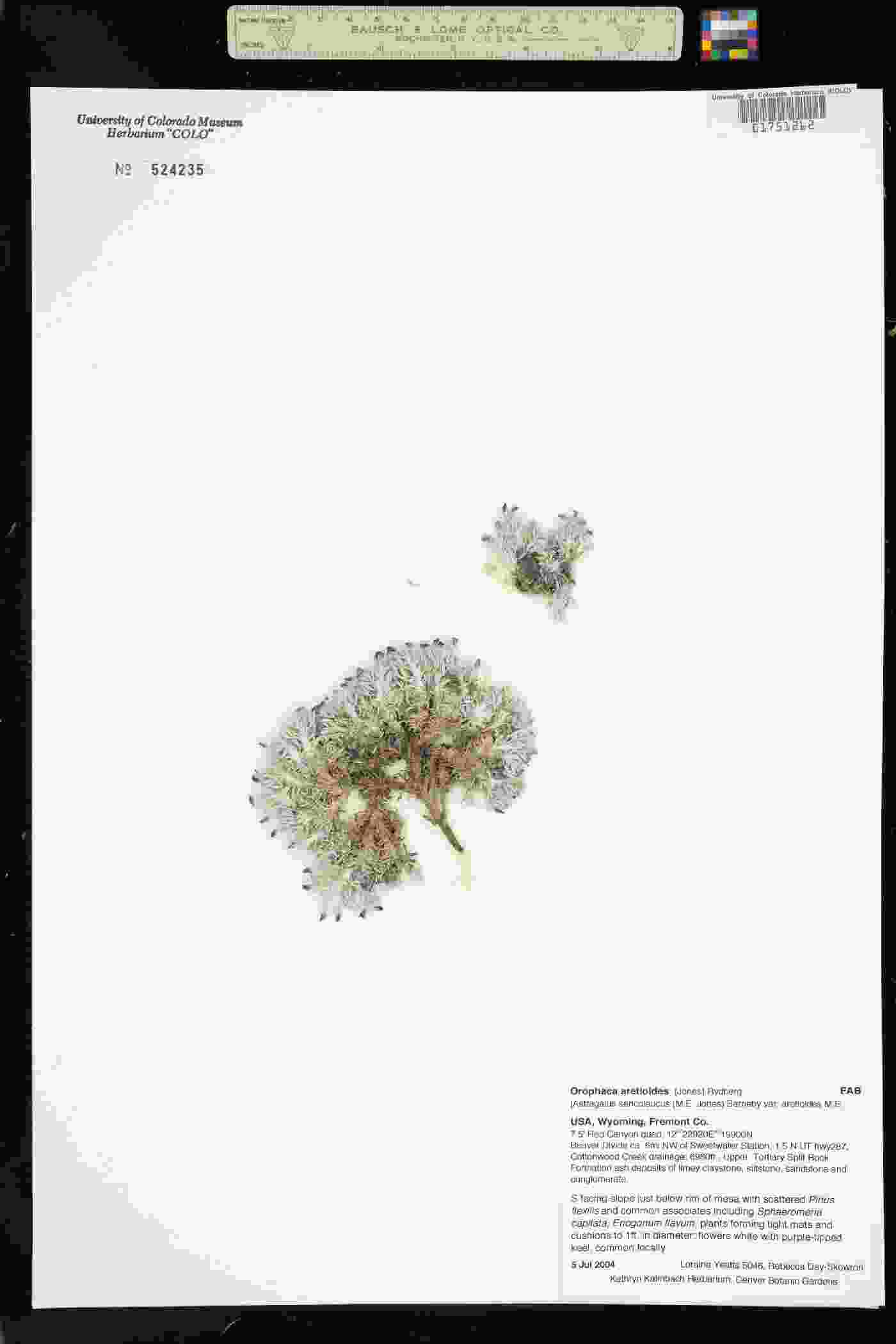

### Astragalus_aretioides_3775729775.jpeg

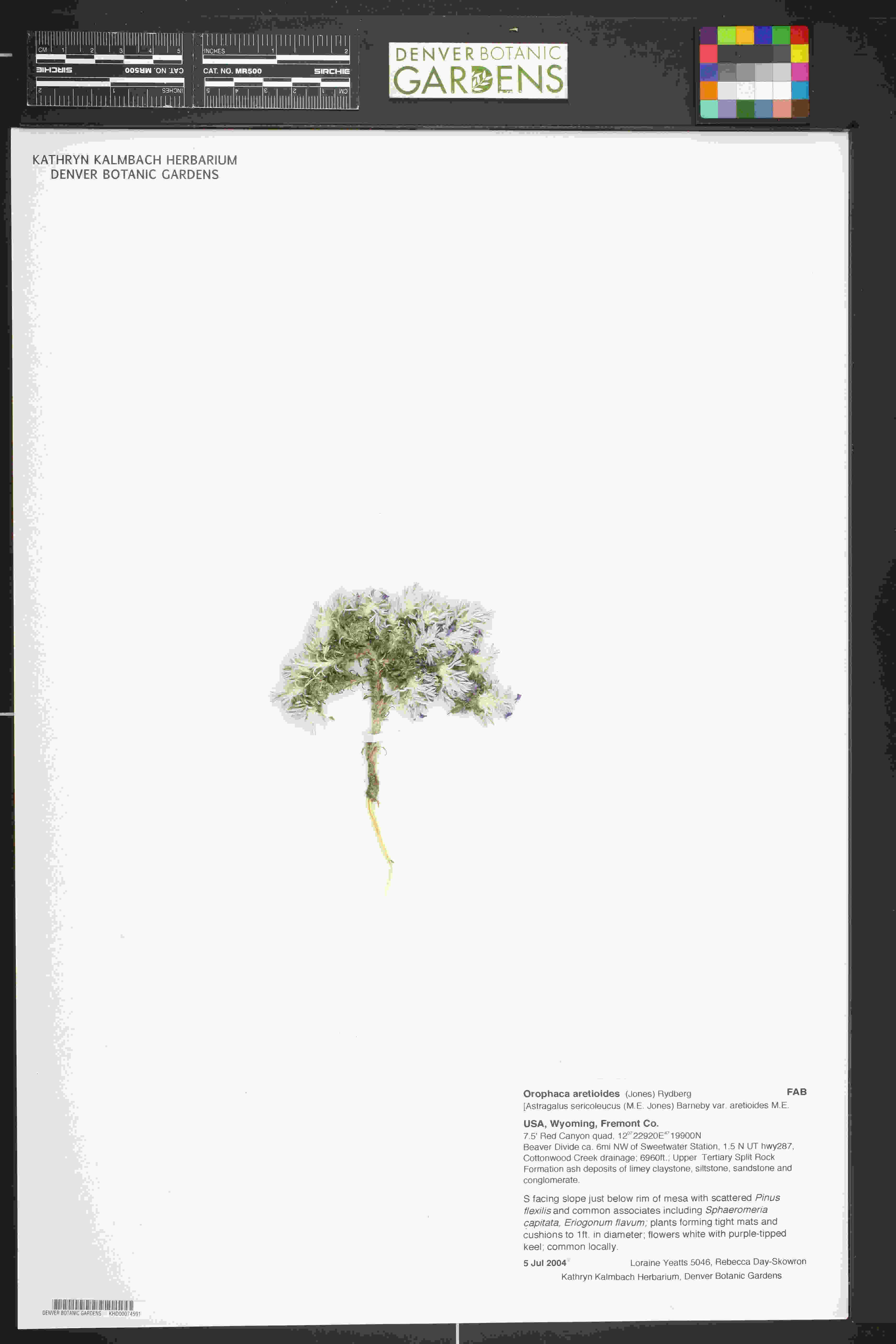

### Astragalus_aretioides_4912130250.jpeg

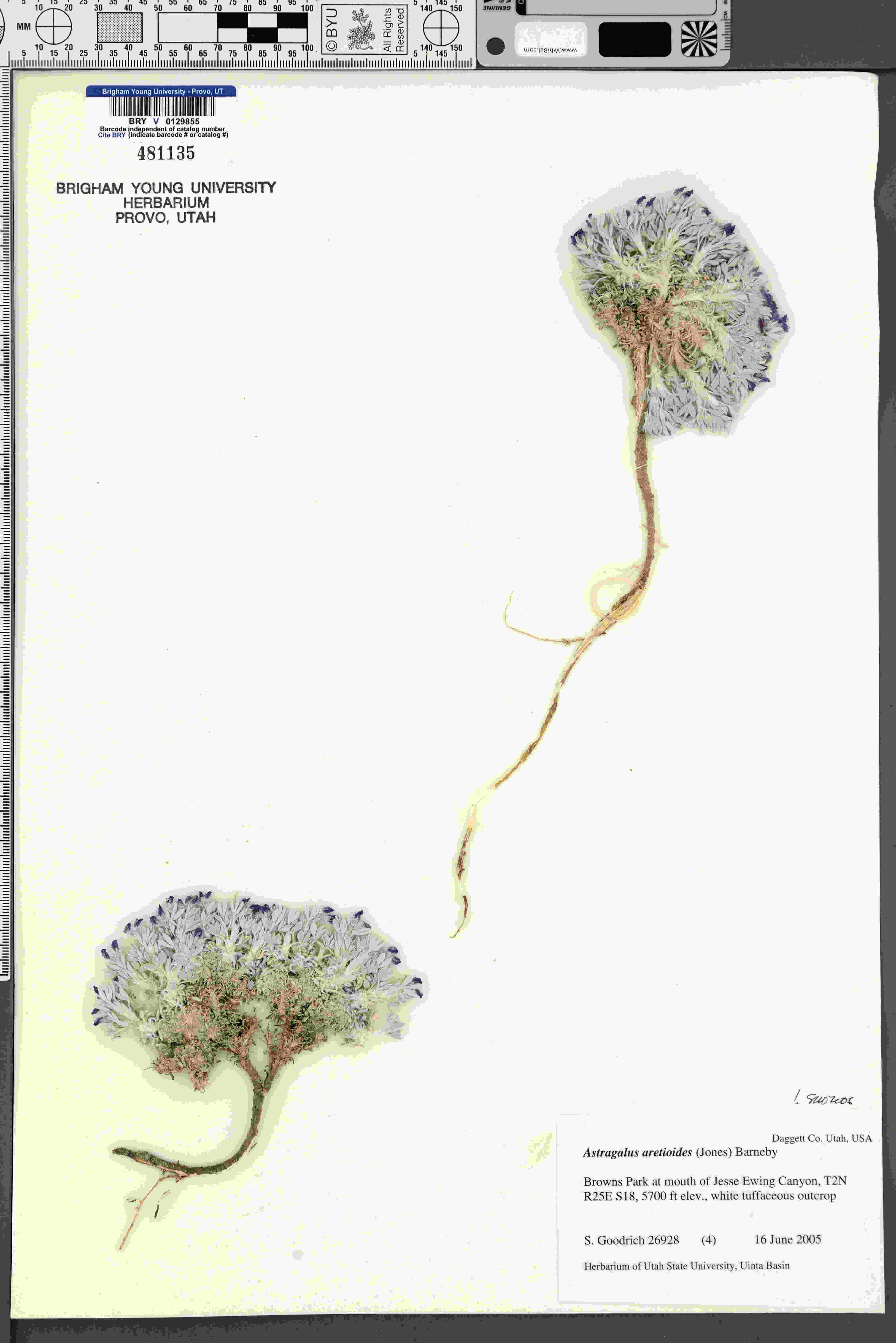

### Bellevalia_eigii_437028838.jpeg

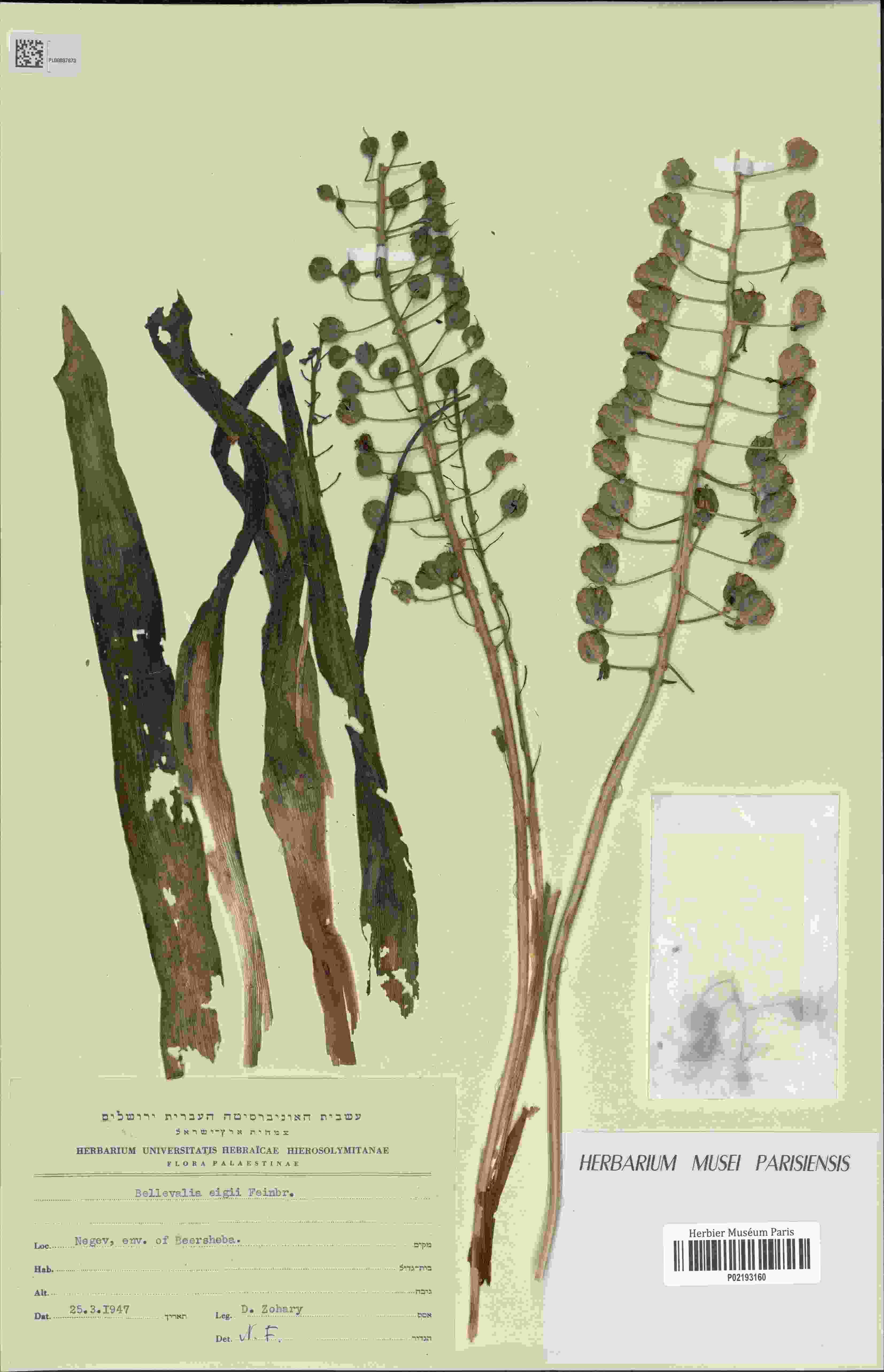

### Bellevalia_eigii_1882642624.jpeg

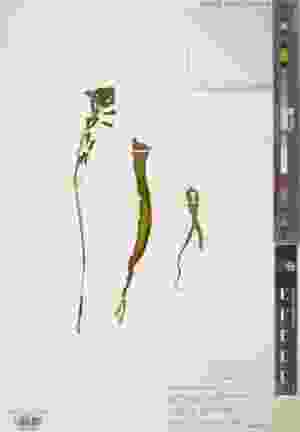

### Bellevalia_eigii_2517397454.jpeg

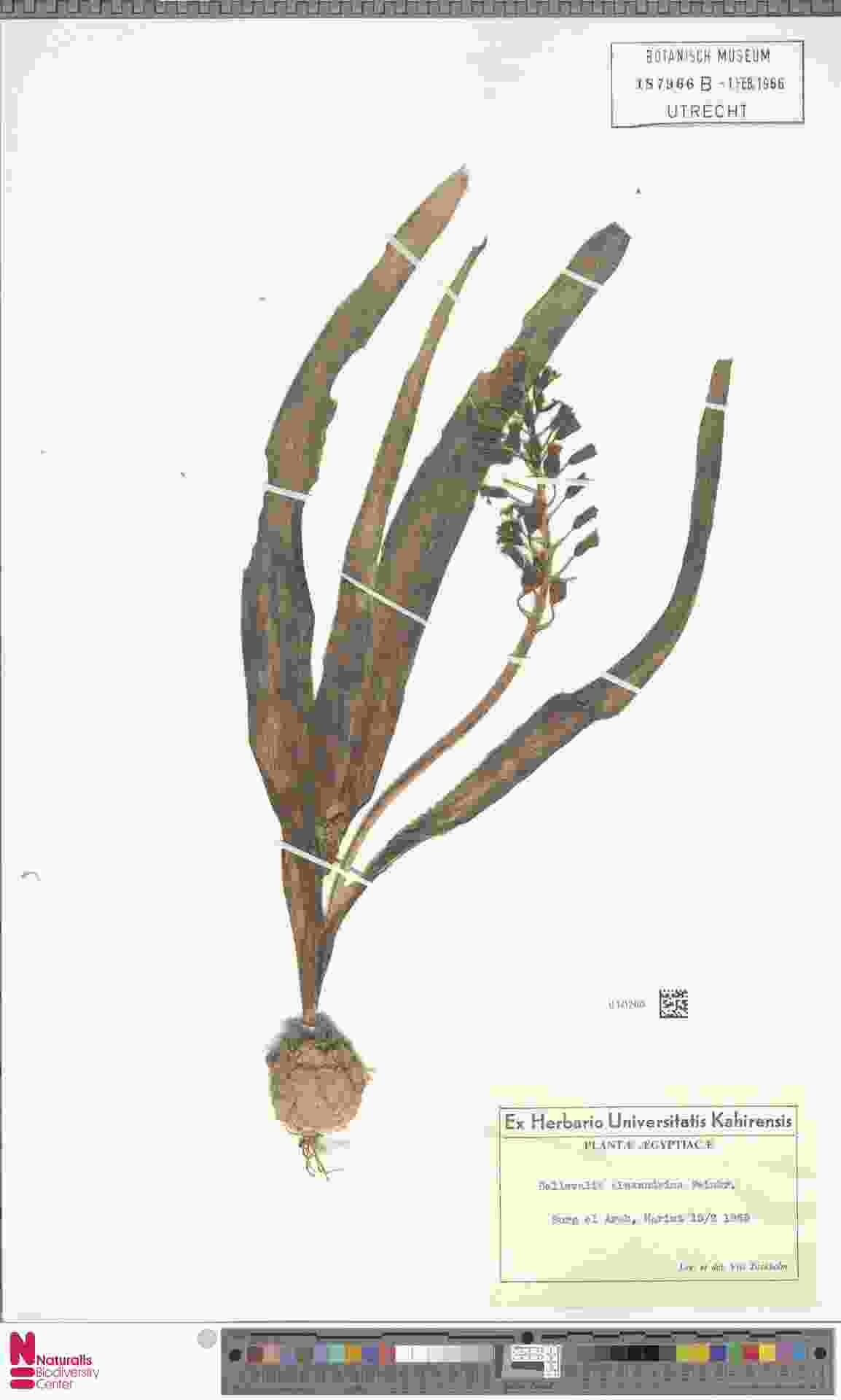

### Bellevalia_eigii_4073252680.jpeg

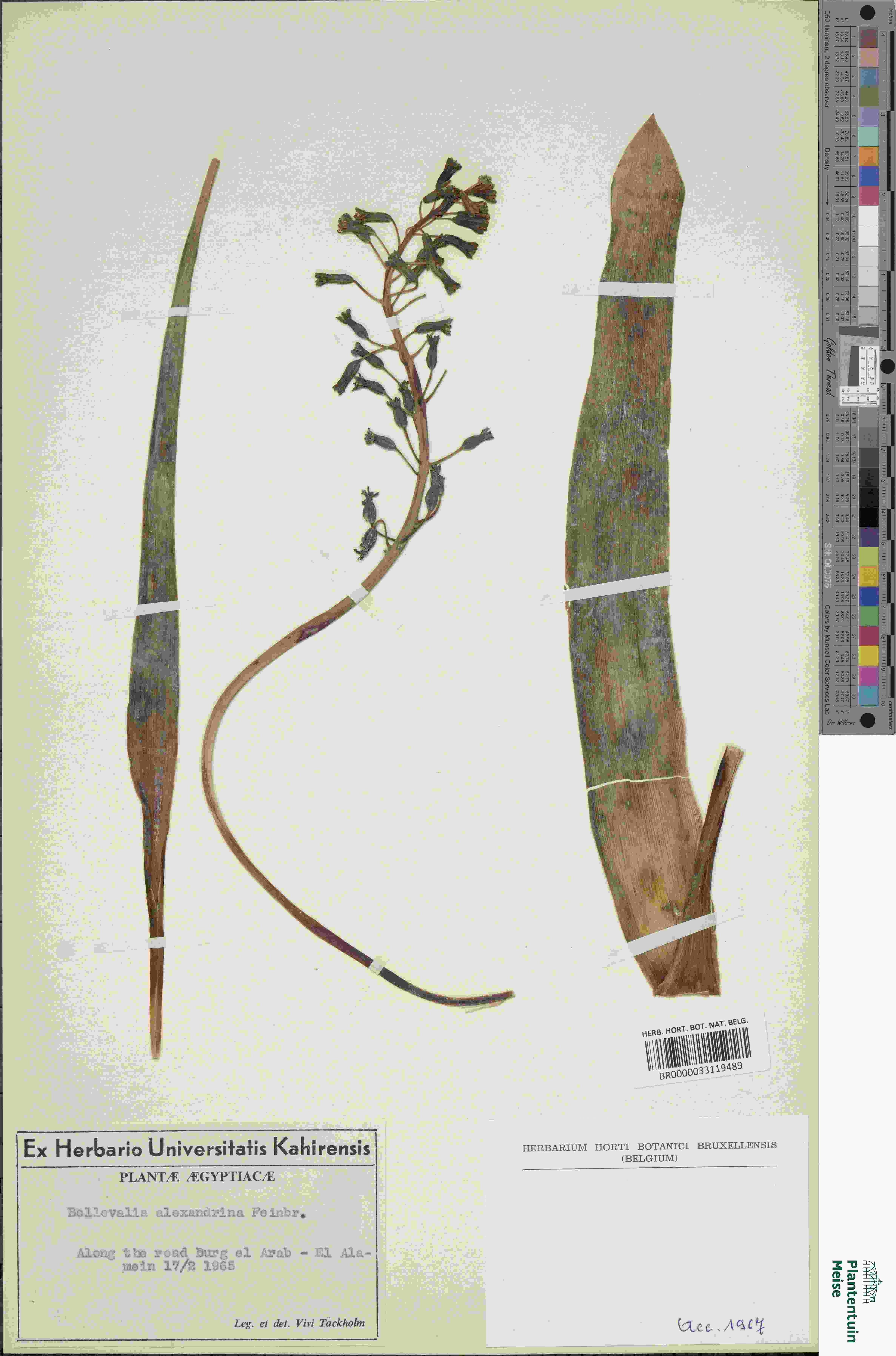

### Bellevalia_eigii_4607950294.jpeg

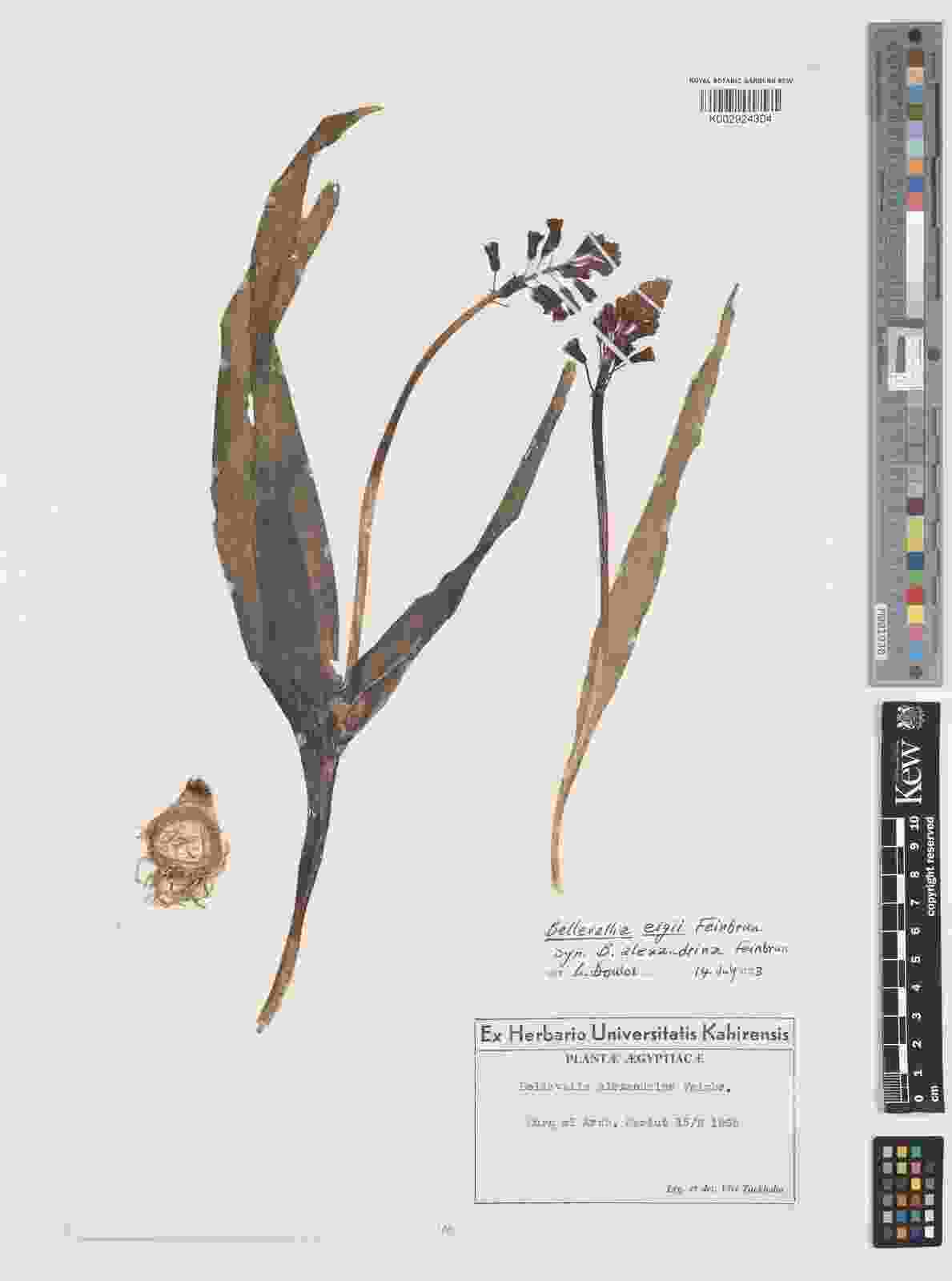

### Blindia_magellanica_4016433369.jpeg

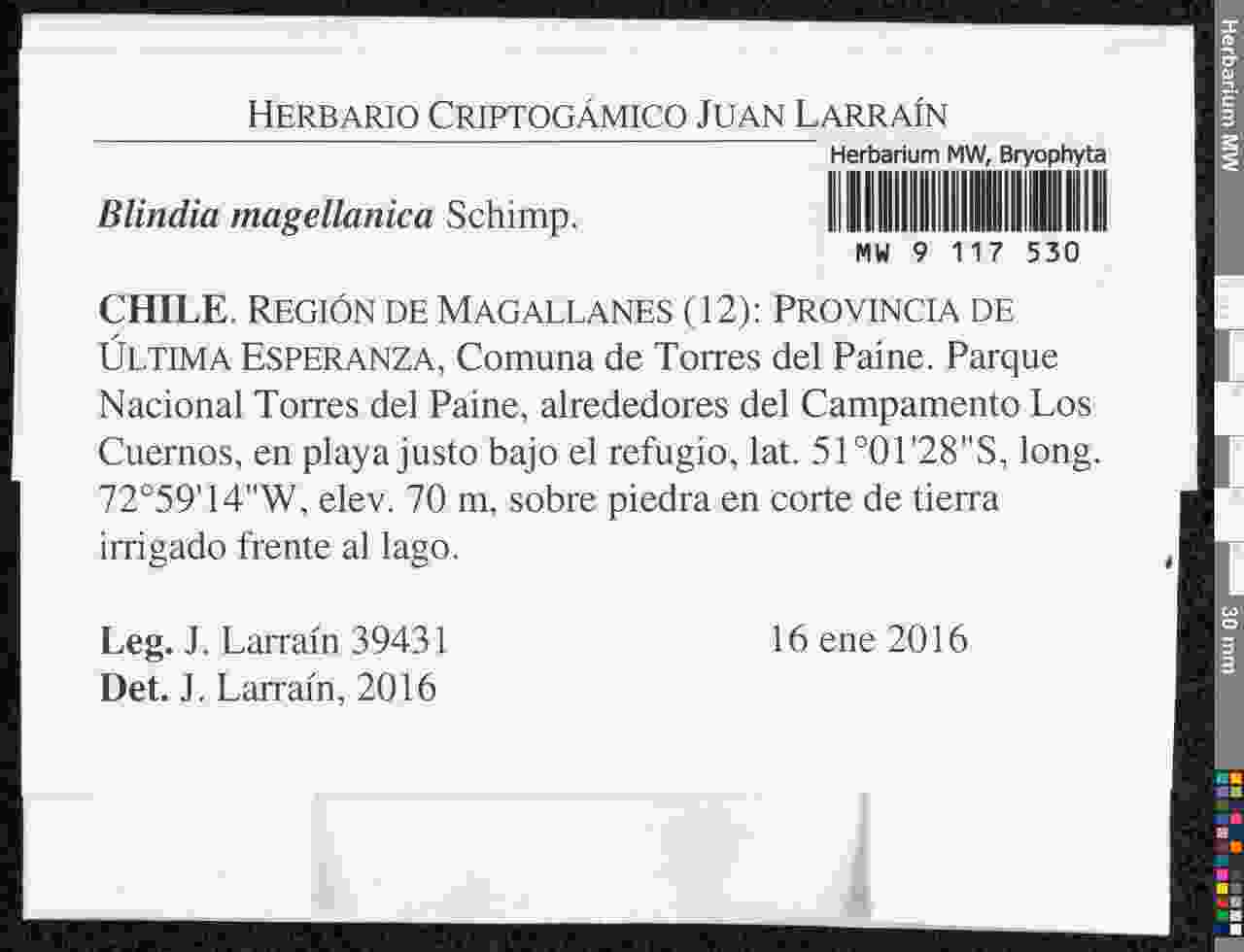

### Blindia_magellanica_4016436363.jpeg

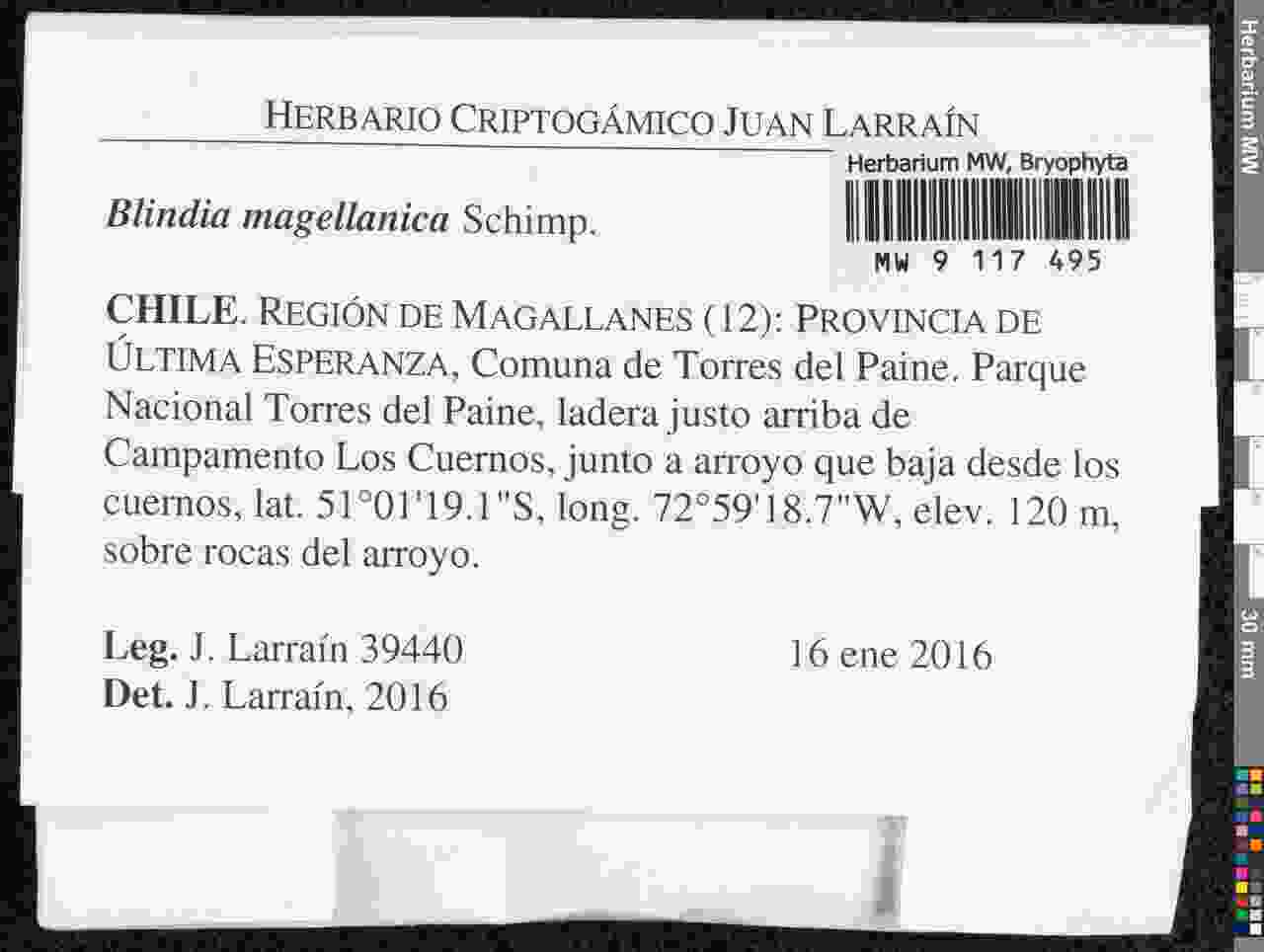

### Calluna_vulgaris_4173254545.jpeg

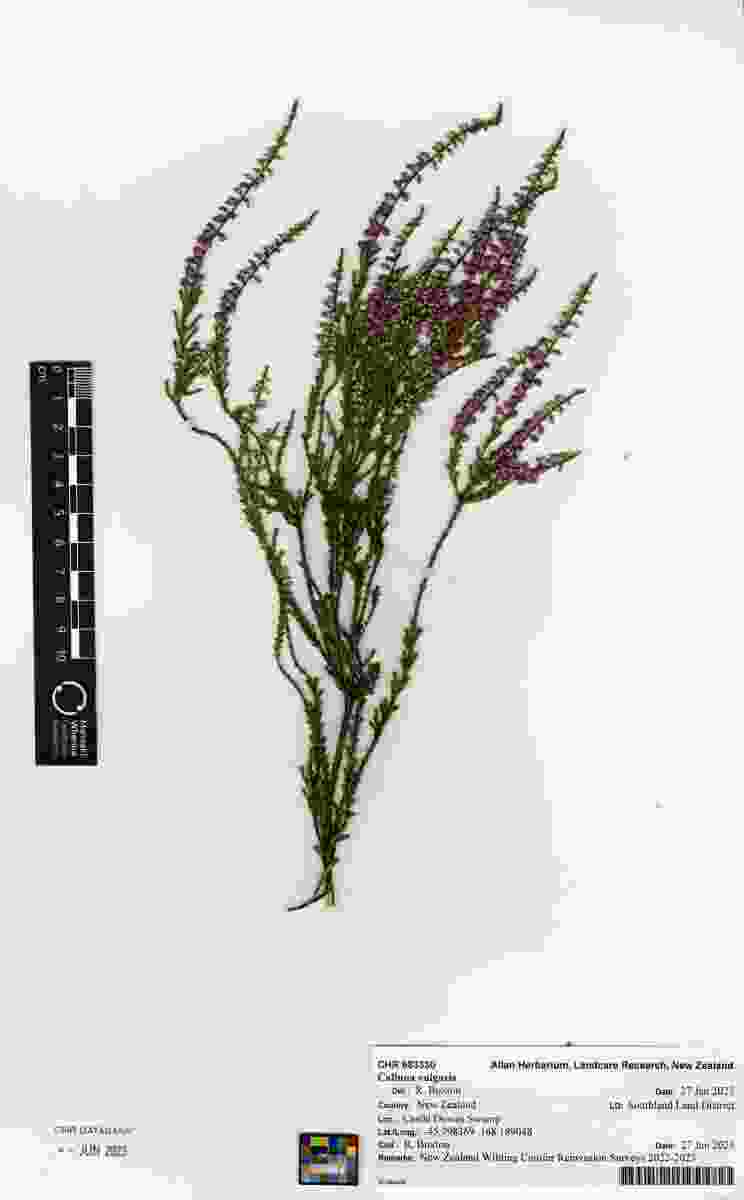

### Carex_cyanea_4435329390.jpeg

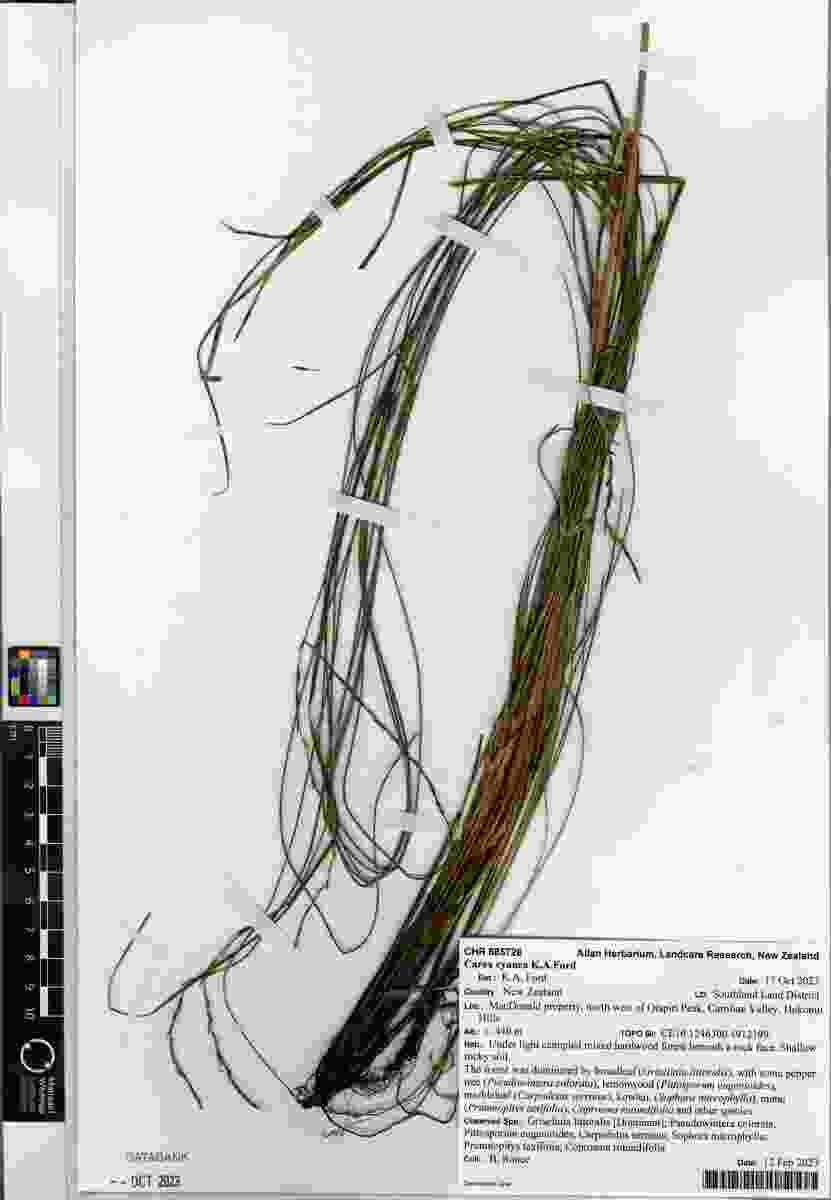

### Cousinia_resinosa_1799060258.jpeg

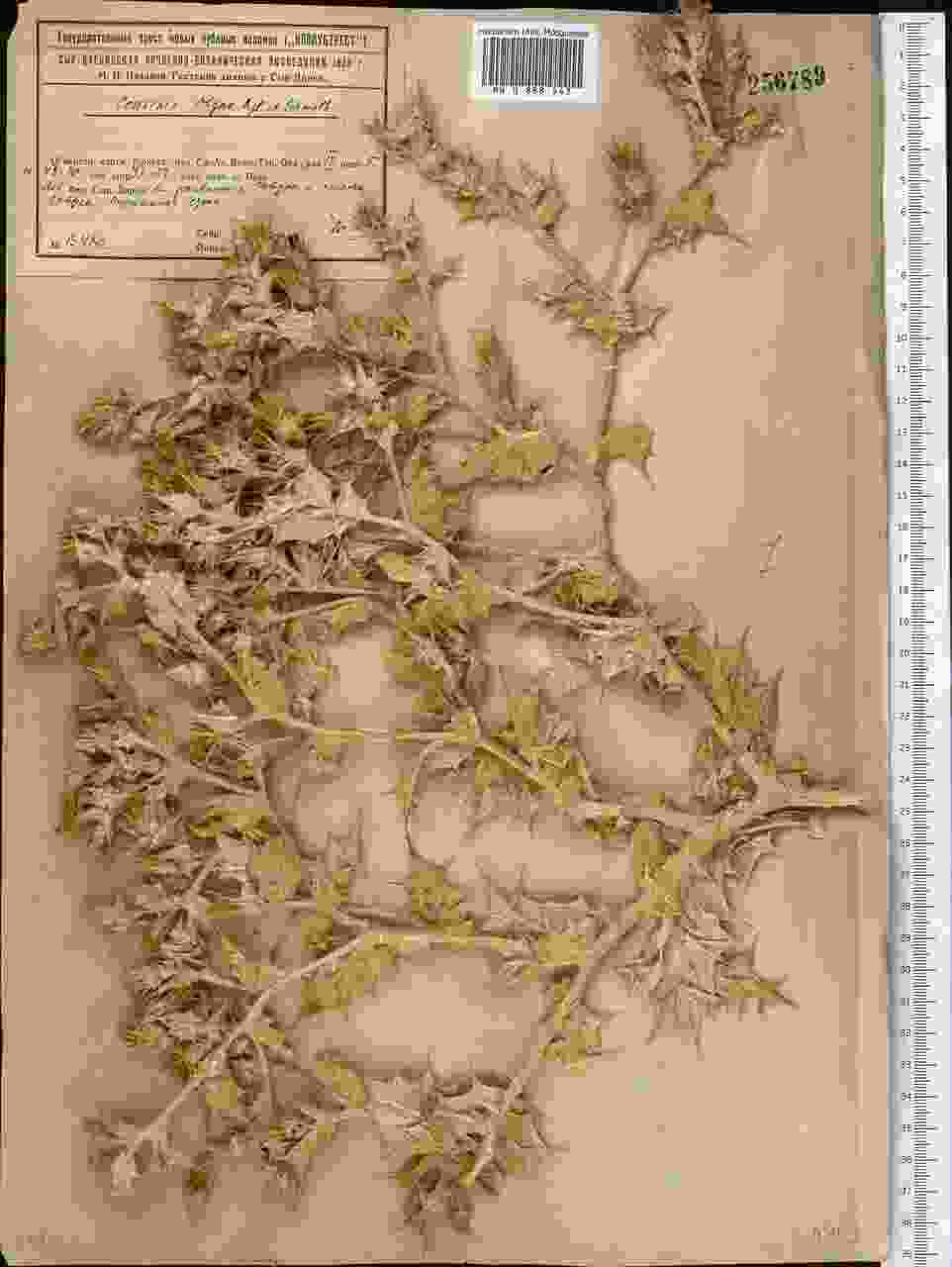

### Cousinia_resinosa_1799060272.jpeg

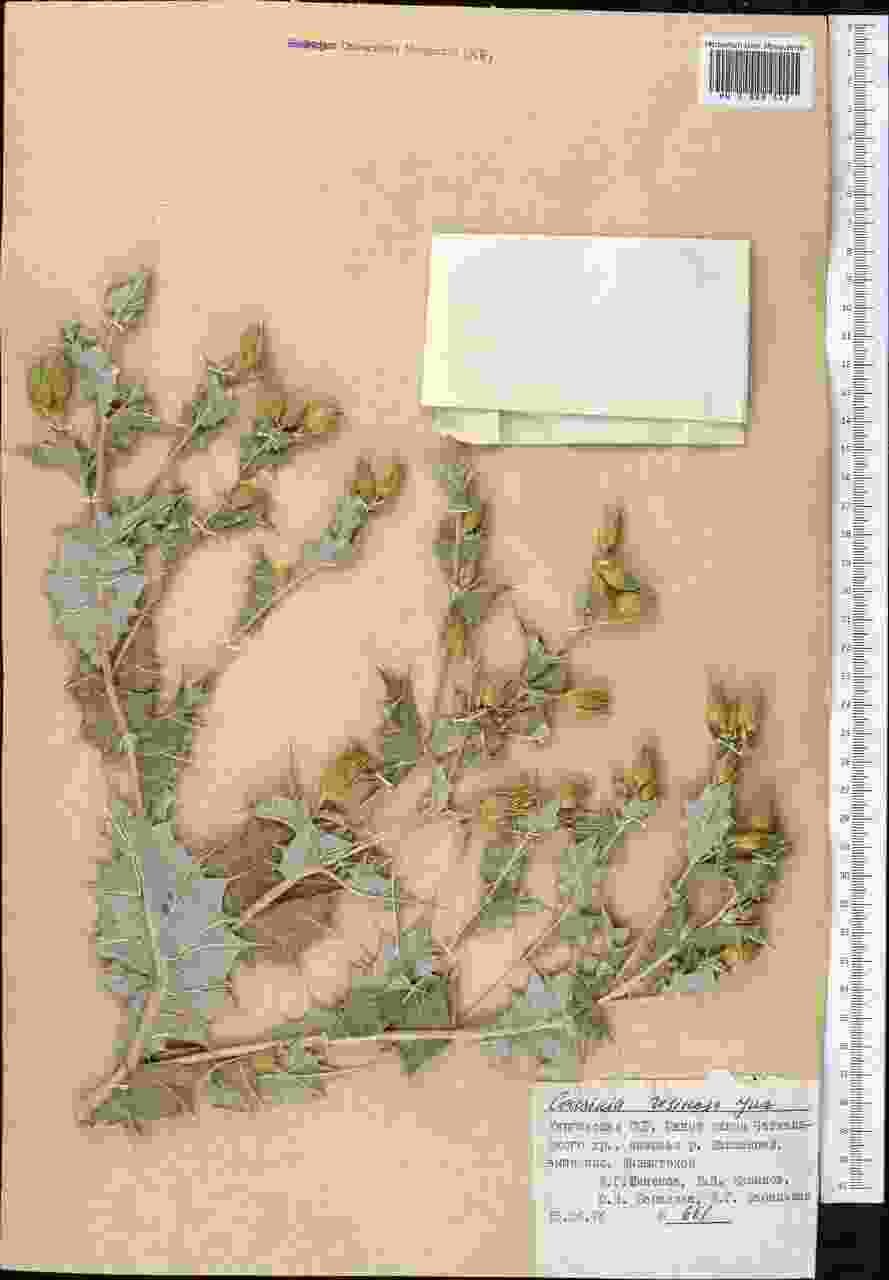

### Cousinia_resinosa_4906864274.jpeg

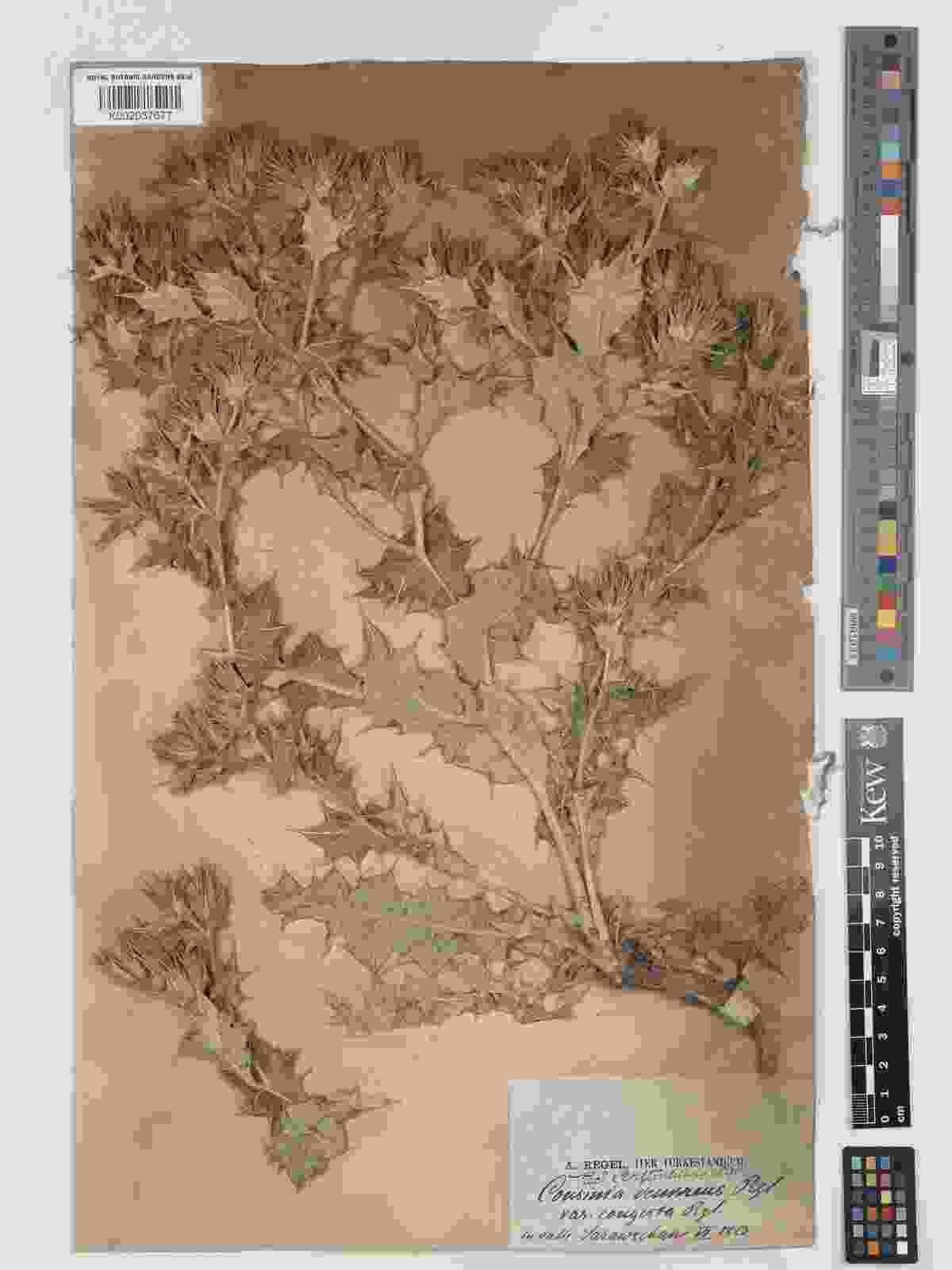

### Cousinia_resinosa_4906868867.jpeg

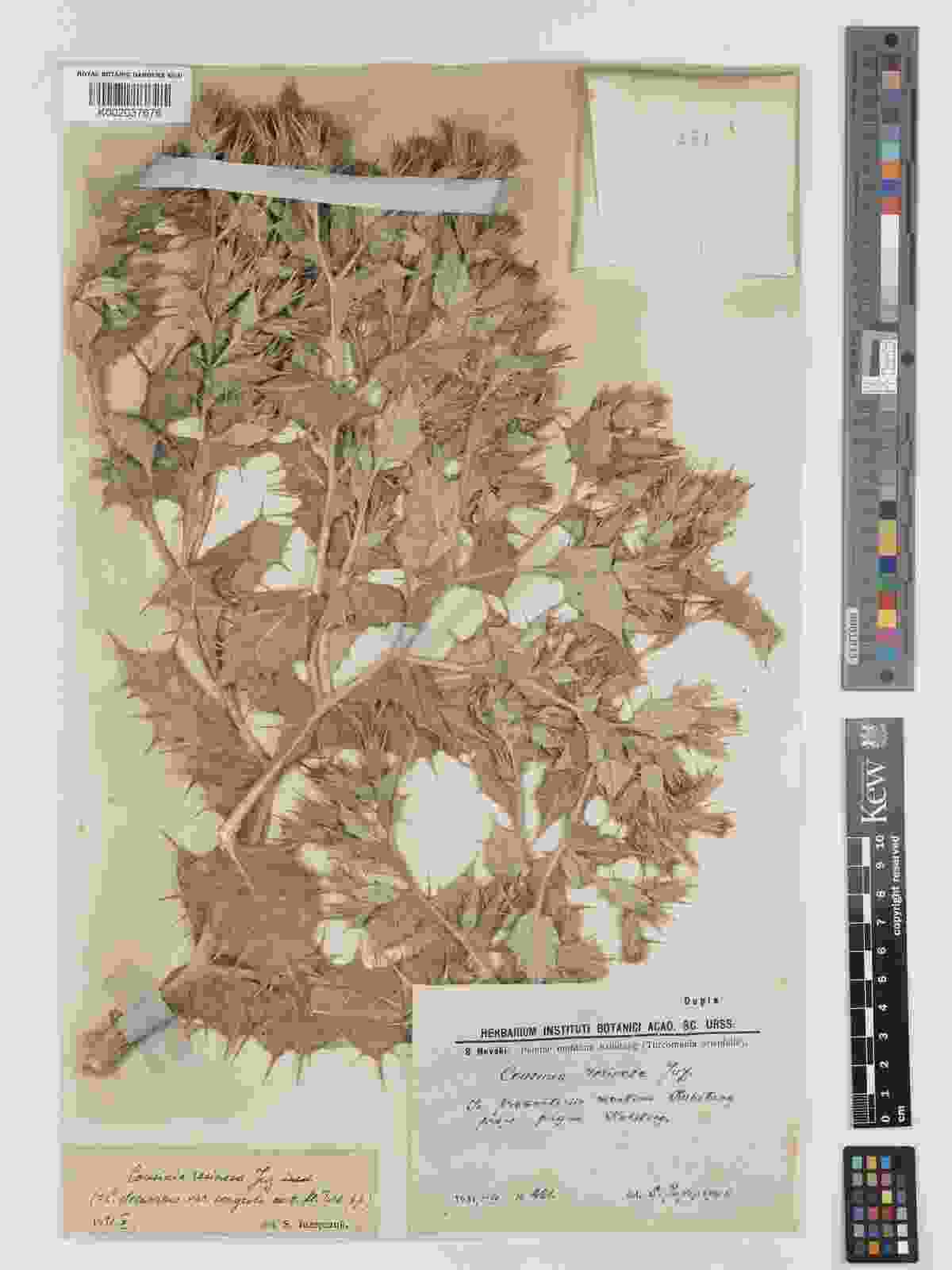

### Dicranella_hookeri_4016437351.jpeg

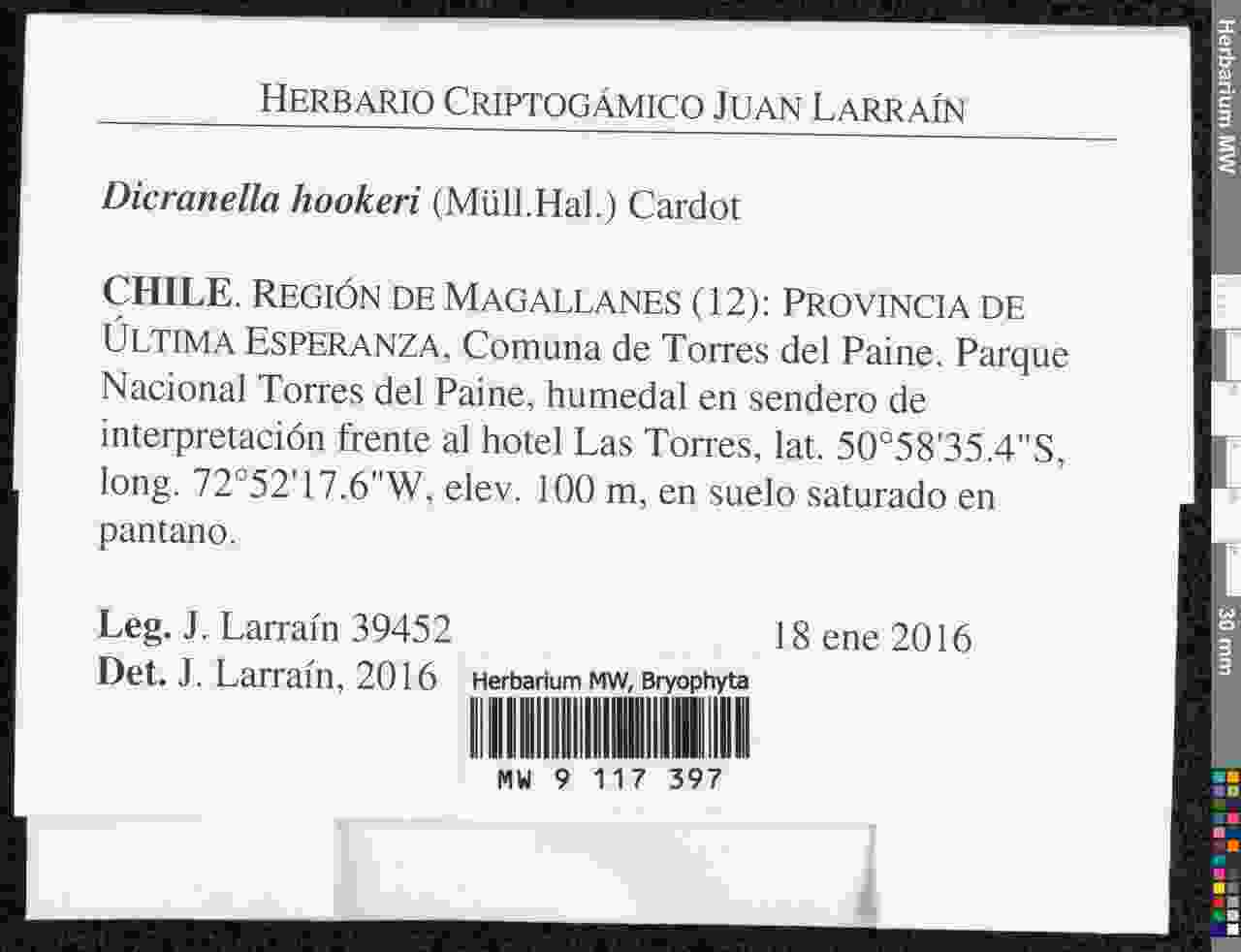

### Eumorphia_davyi_3400971459.jpeg

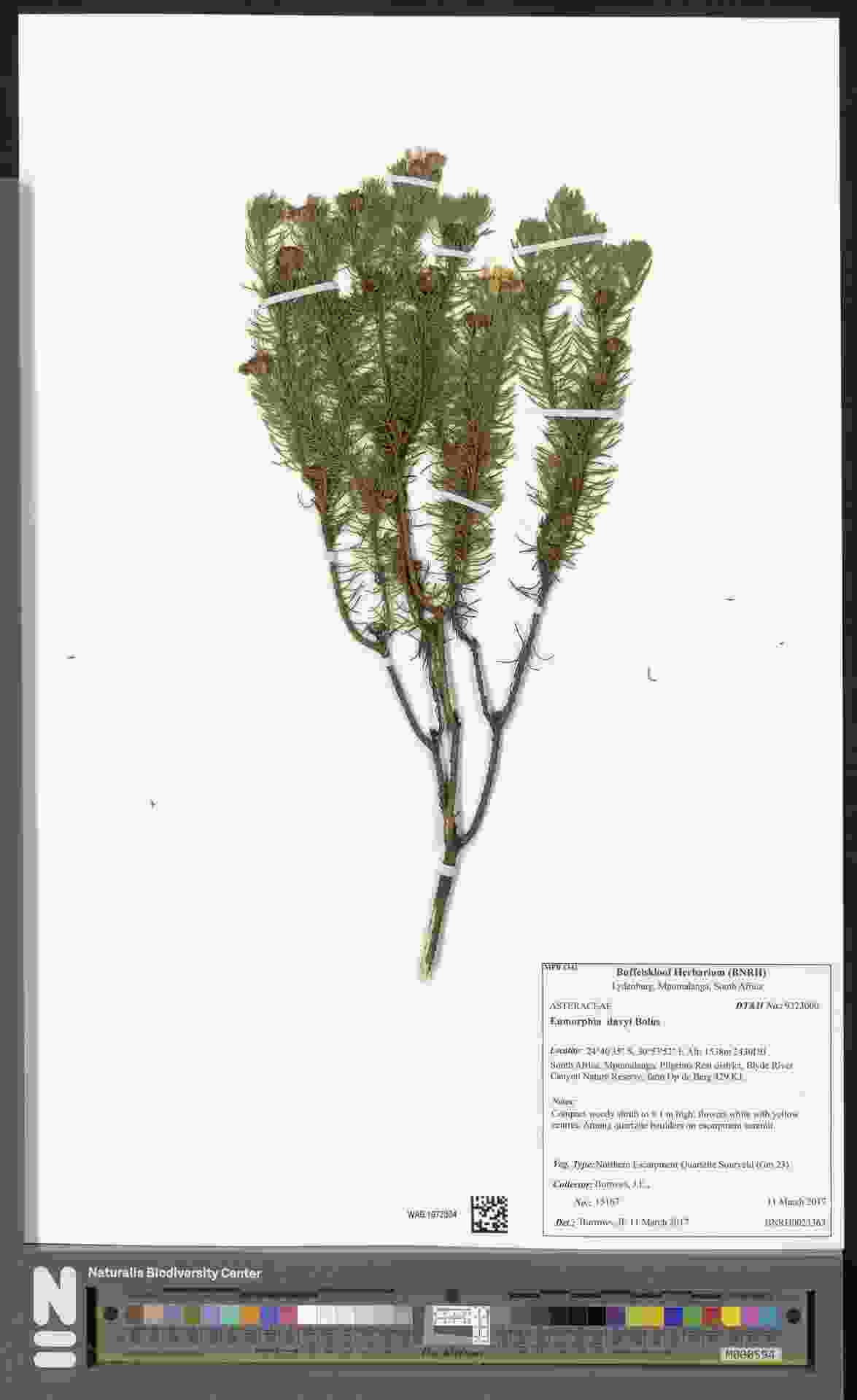

### Gnaphalium_uliginosum_4173253398.jpeg

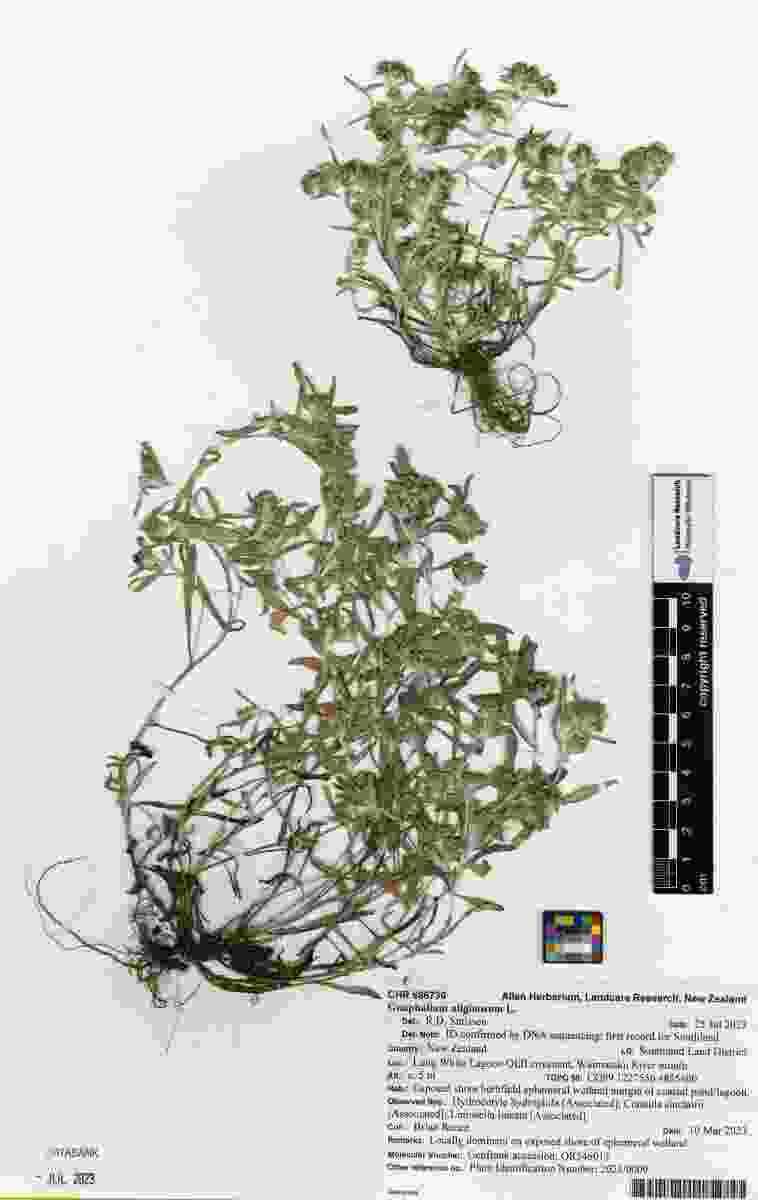

### Hermannia_erodioides_1840197269.jpeg

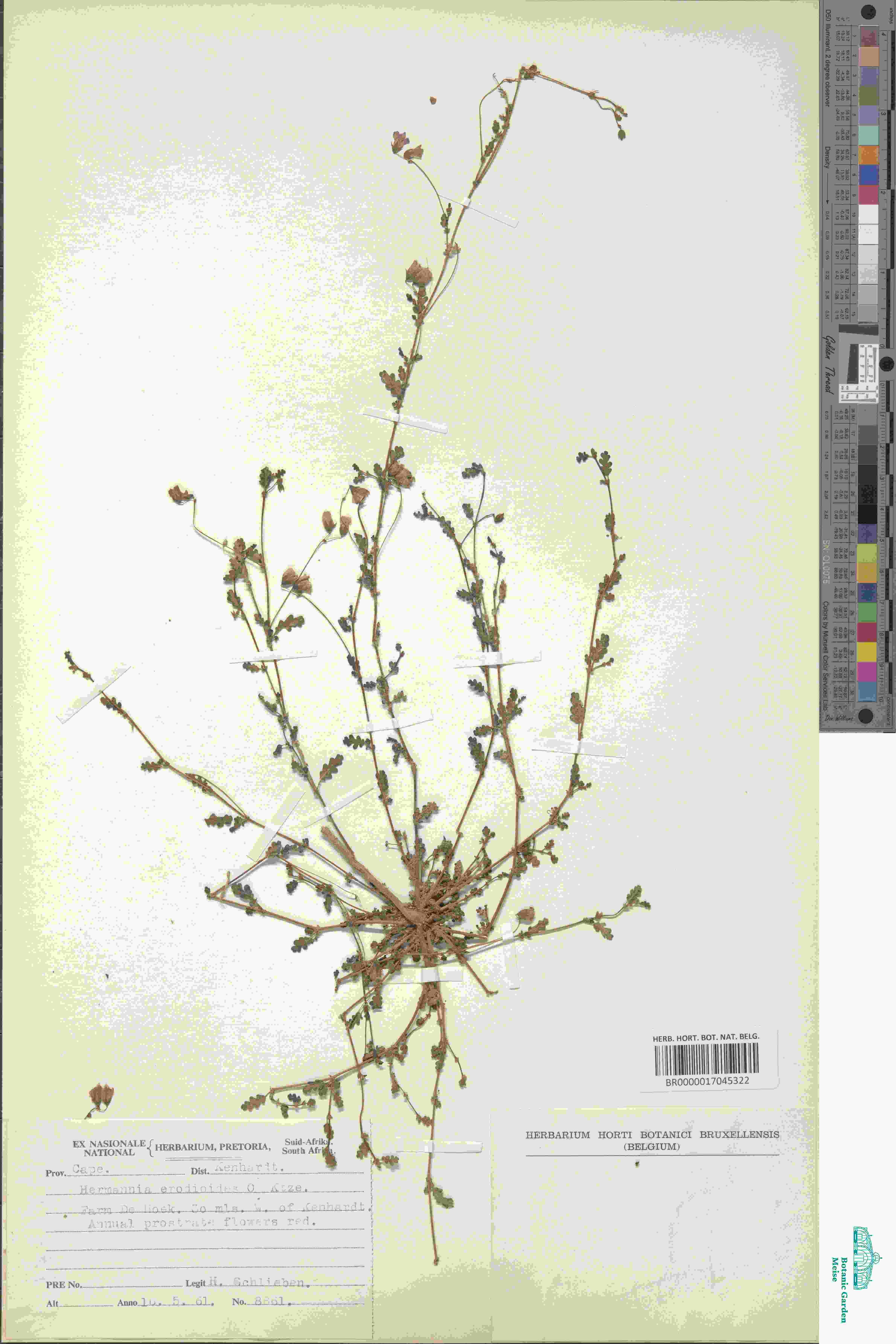

### Hermannia_erodioides_2517185910.jpeg

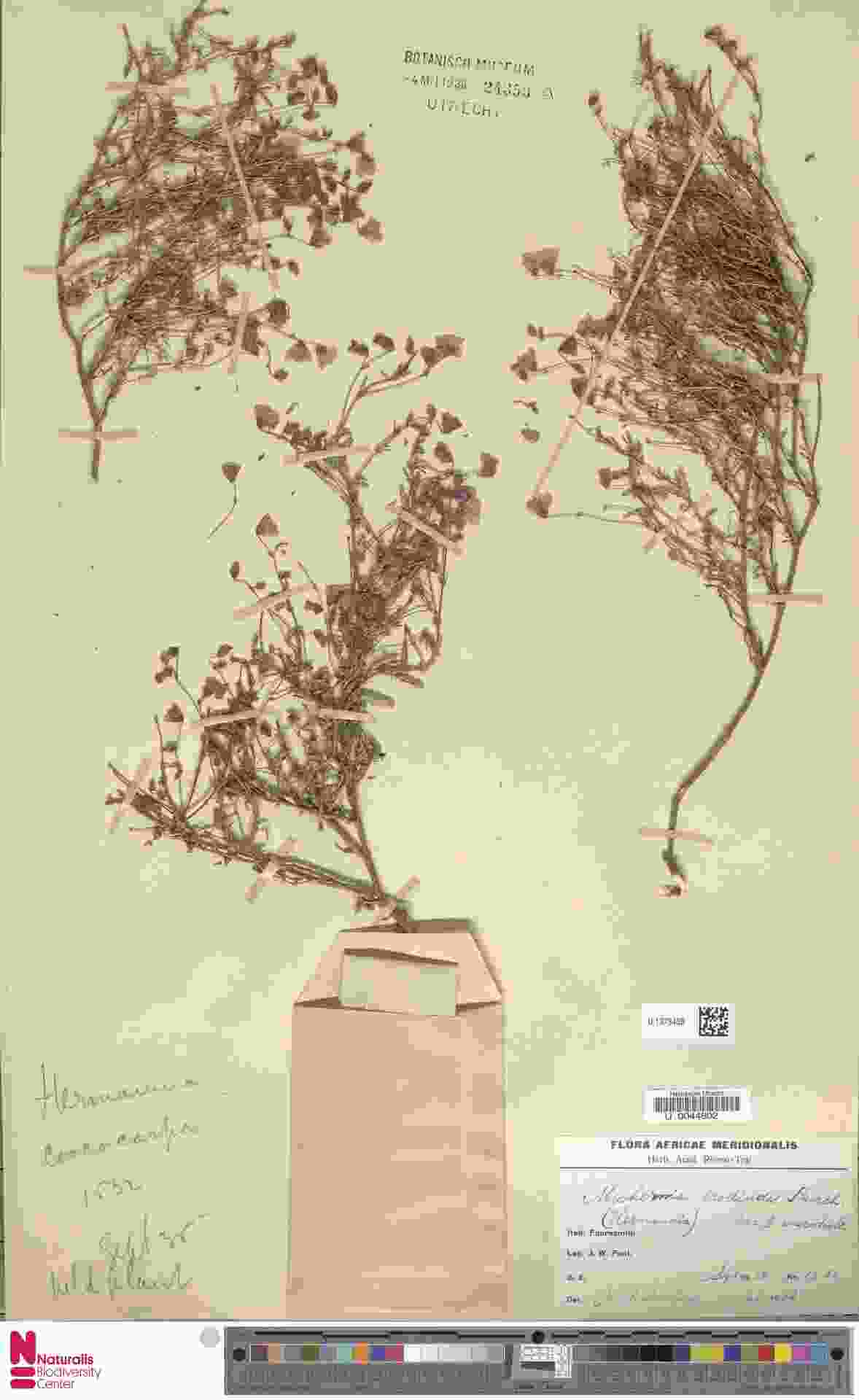

### Hermannia_montana_3400988489.jpeg

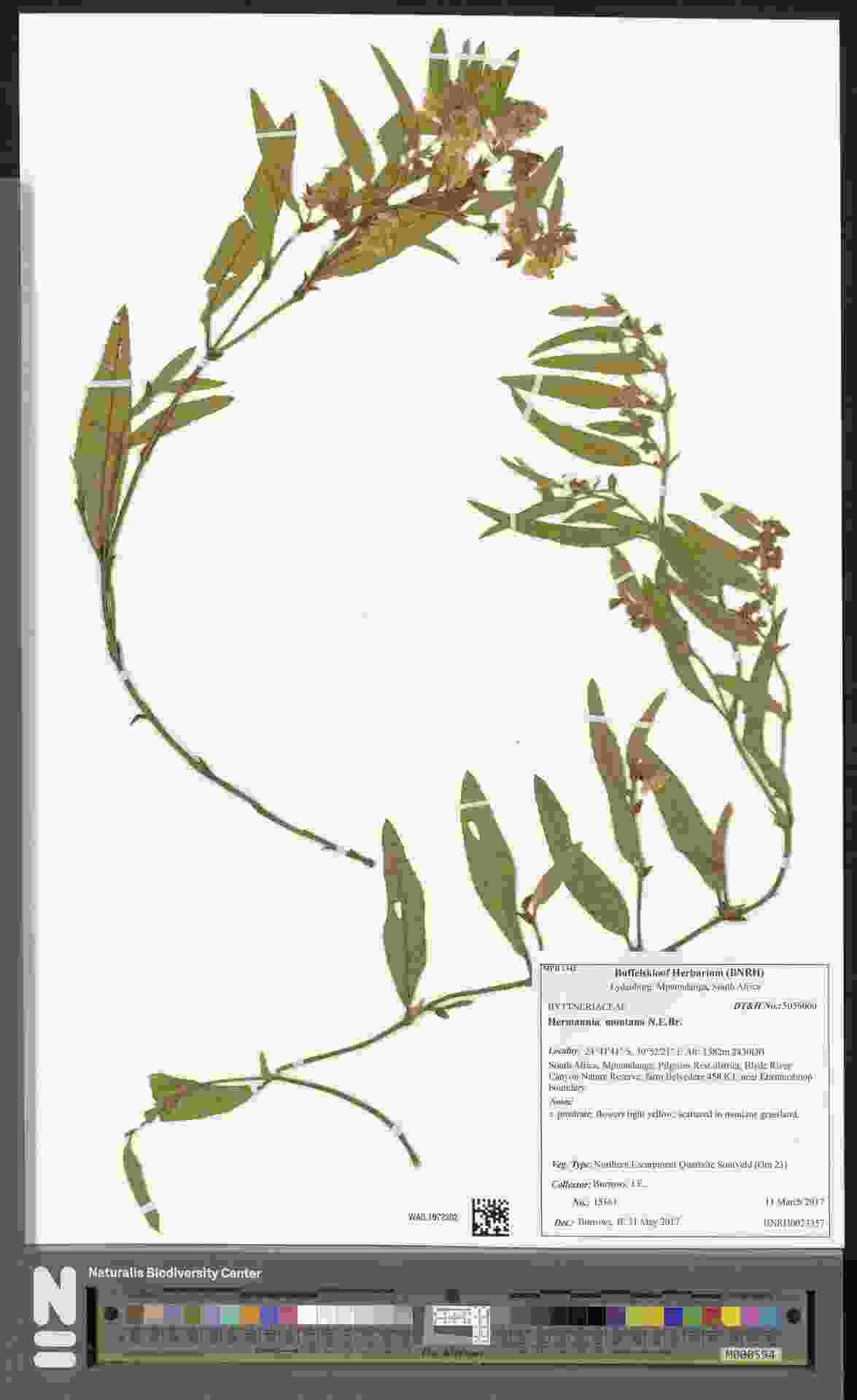

### indet_4016432354.jpeg

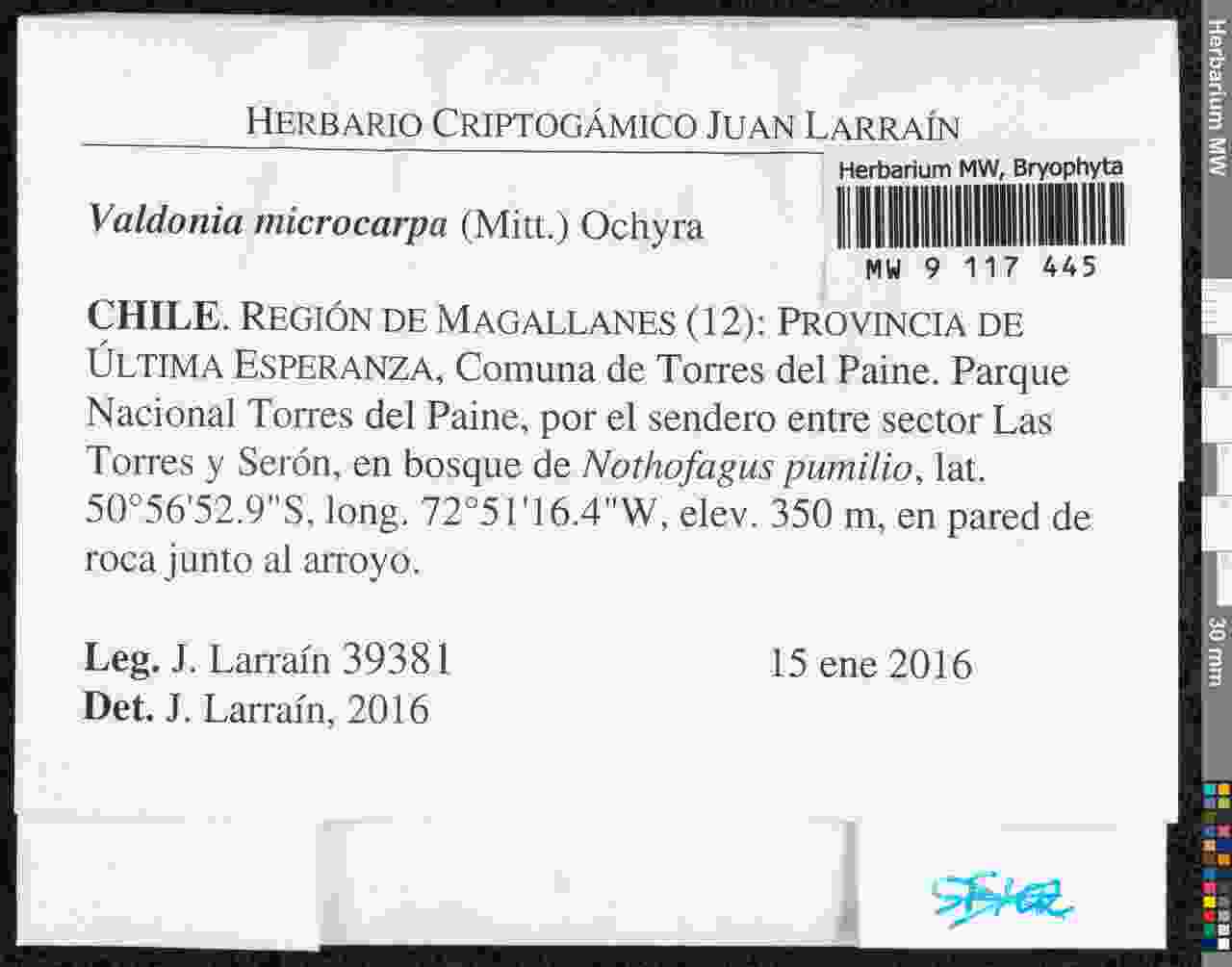

### indet_4016435368.jpeg

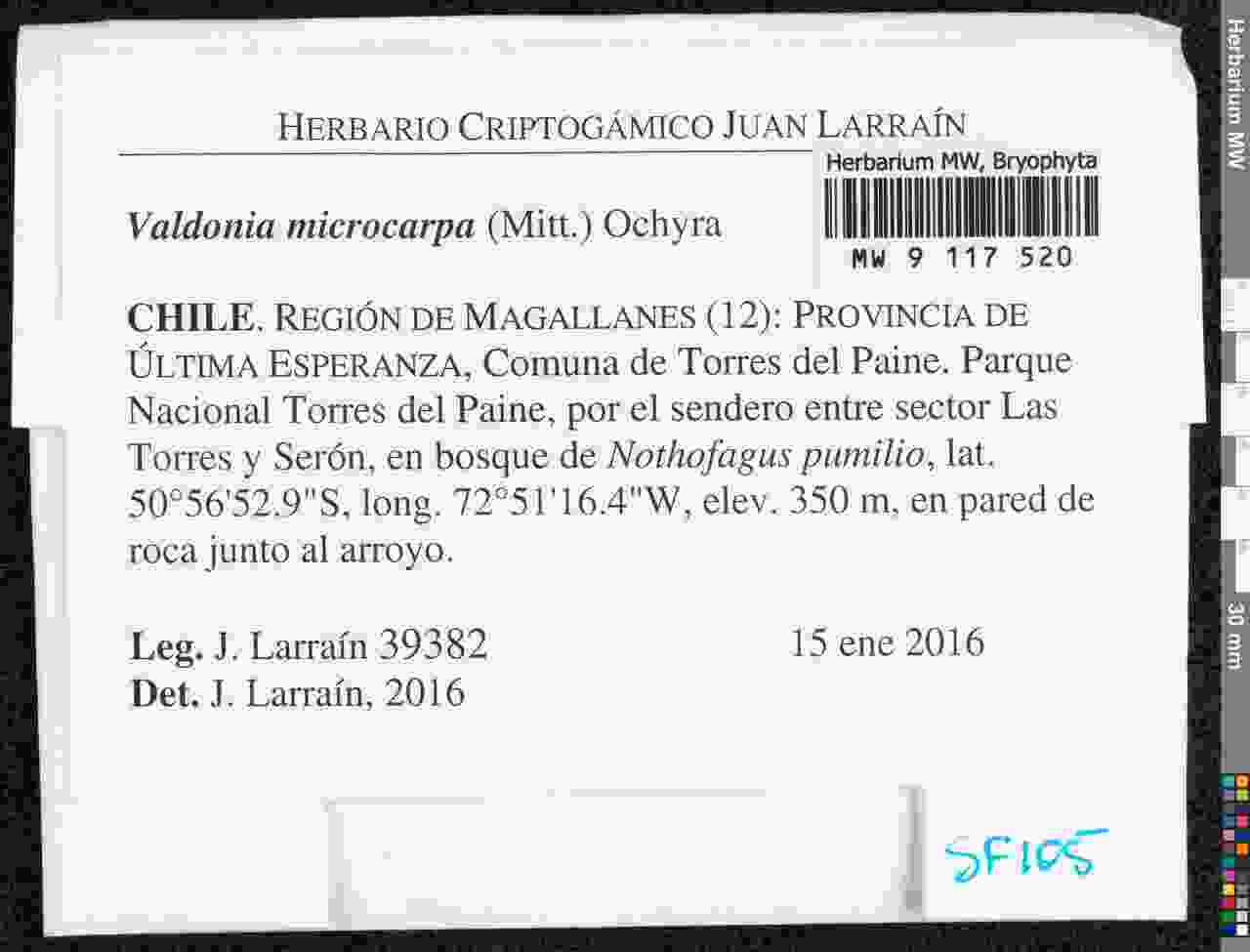

### indet_4435328399.jpeg

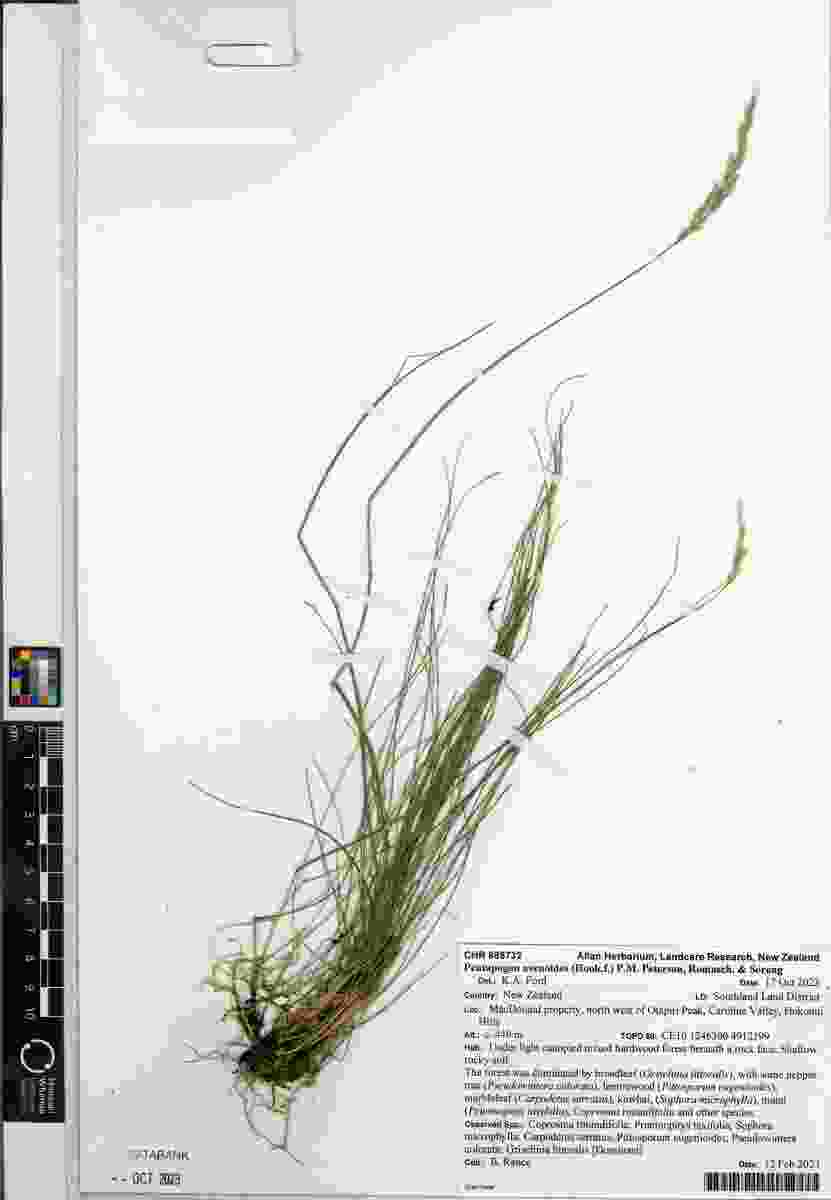

### Isoglossa_delicatula_4020490460.jpeg

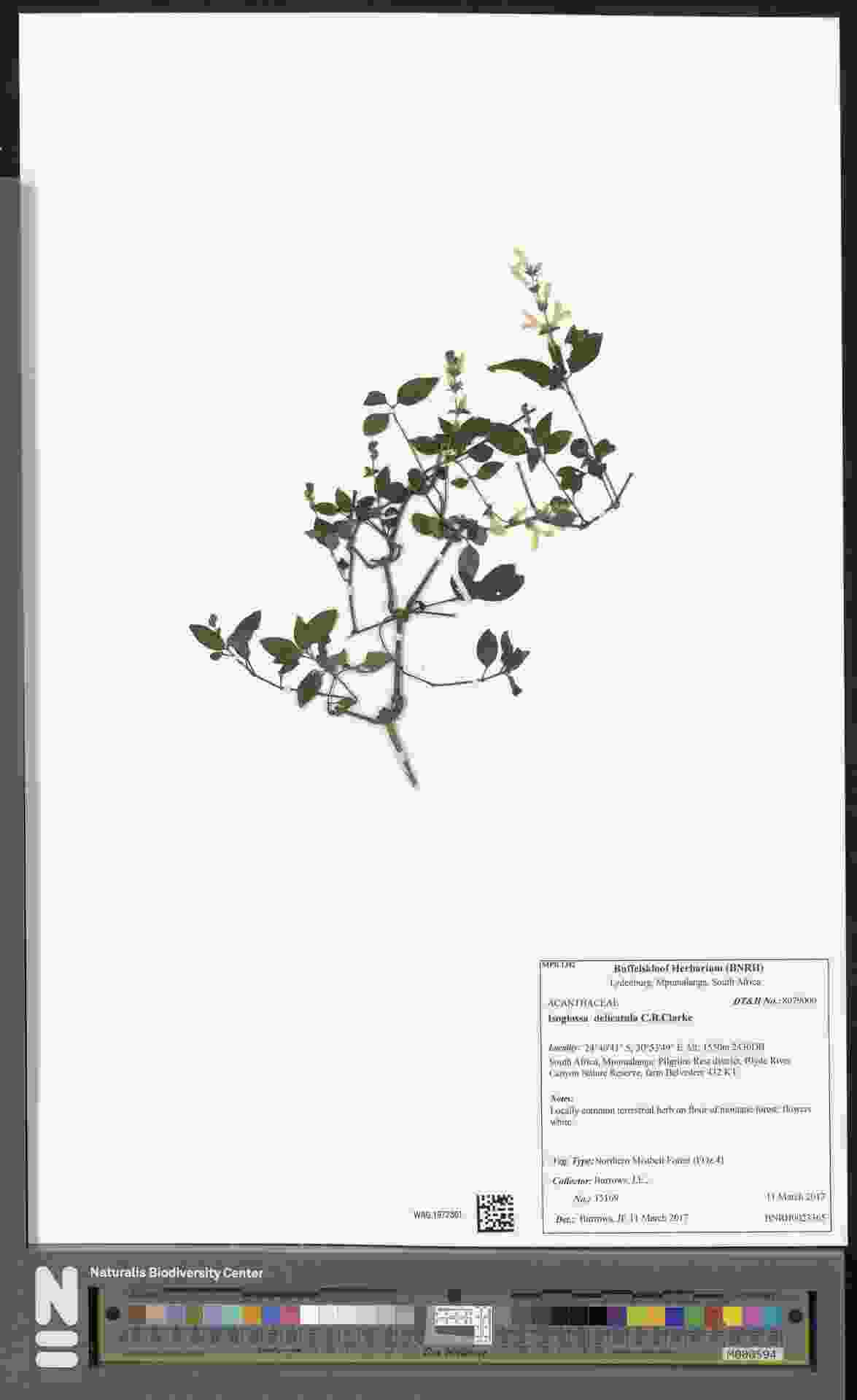

### Lipotriche_scandens_4072653091.jpeg

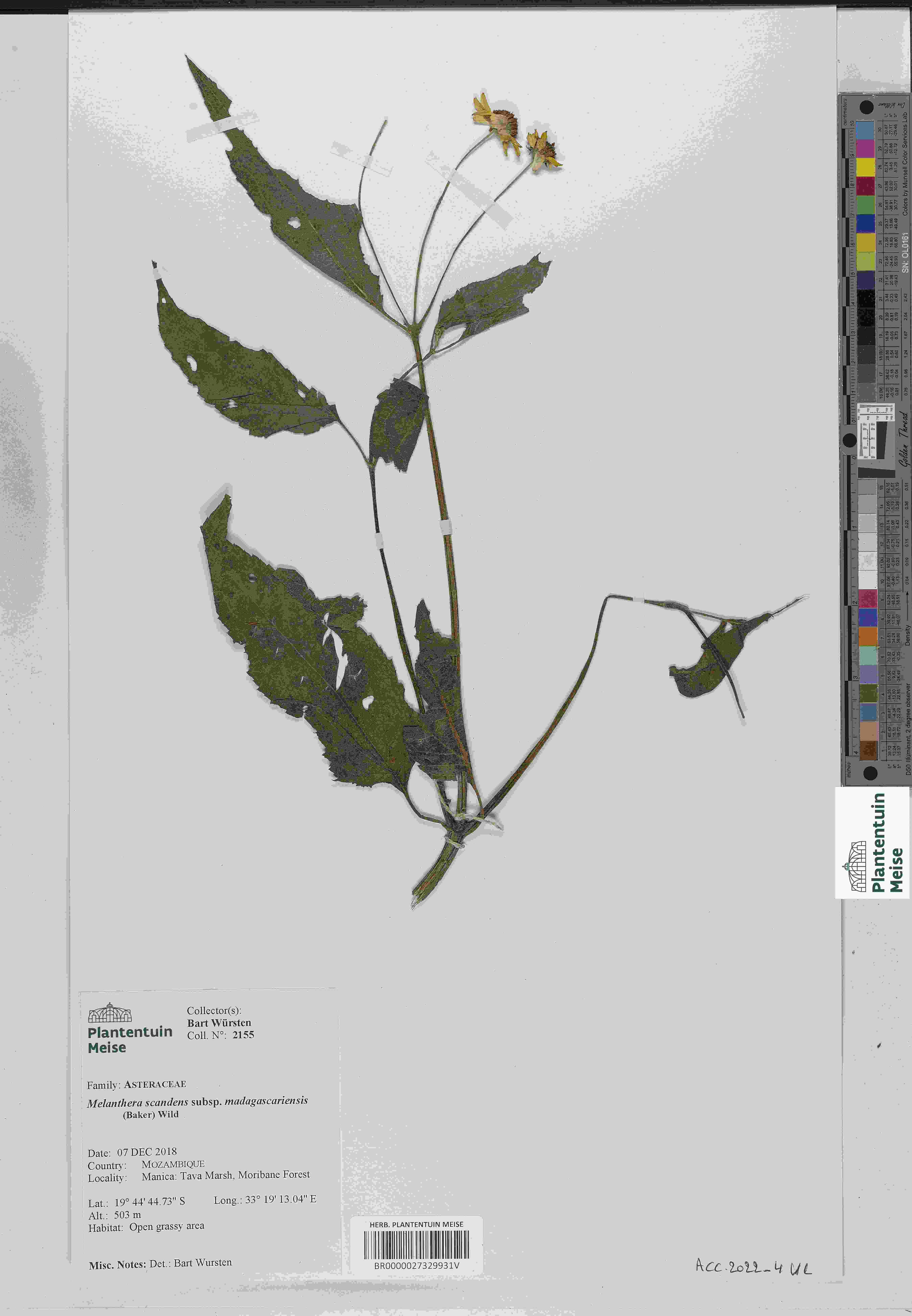

### Lycianthes_testacea_1319345712.jpeg

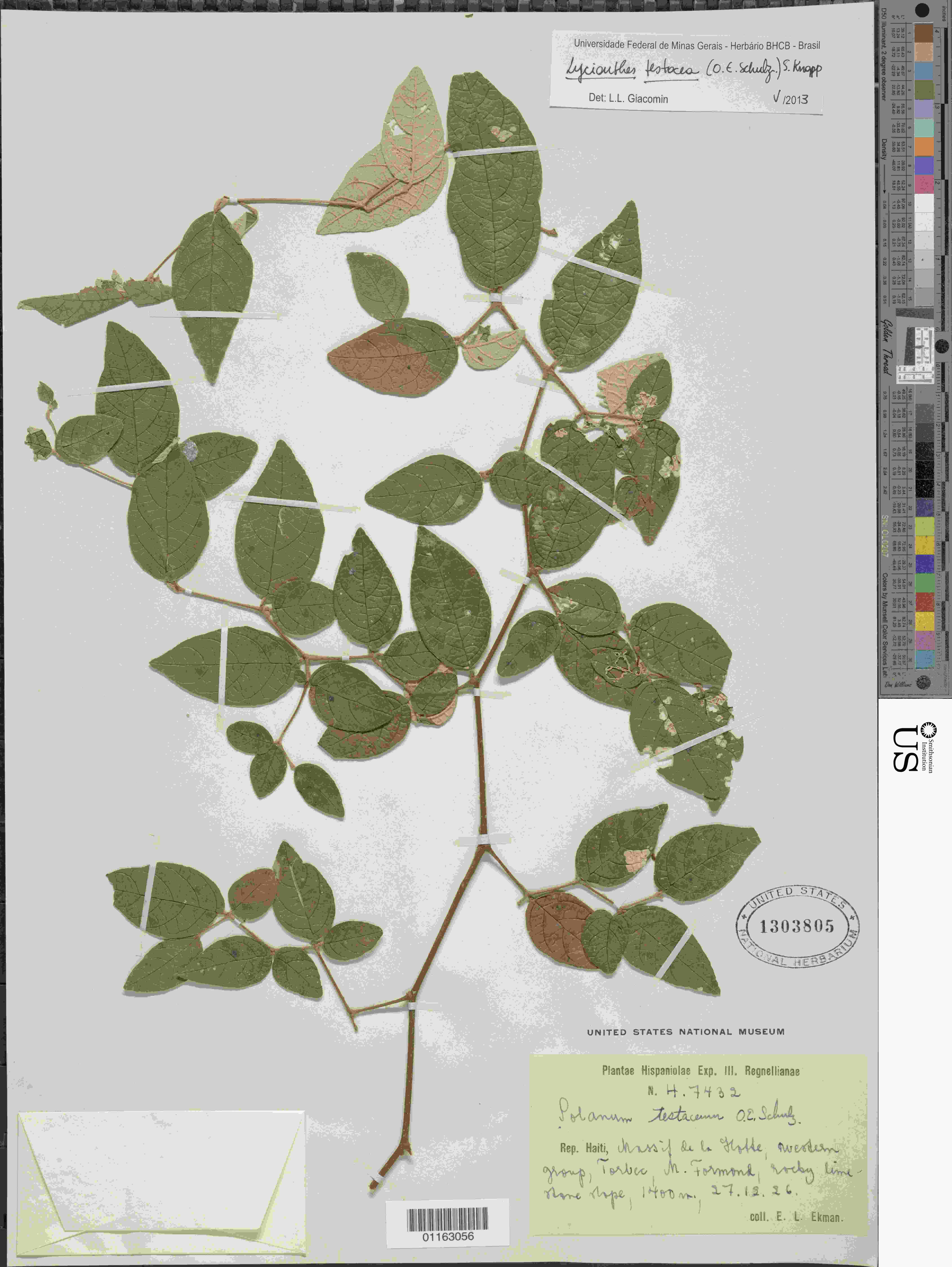

### Lycianthes_testacea_1998442228.jpeg

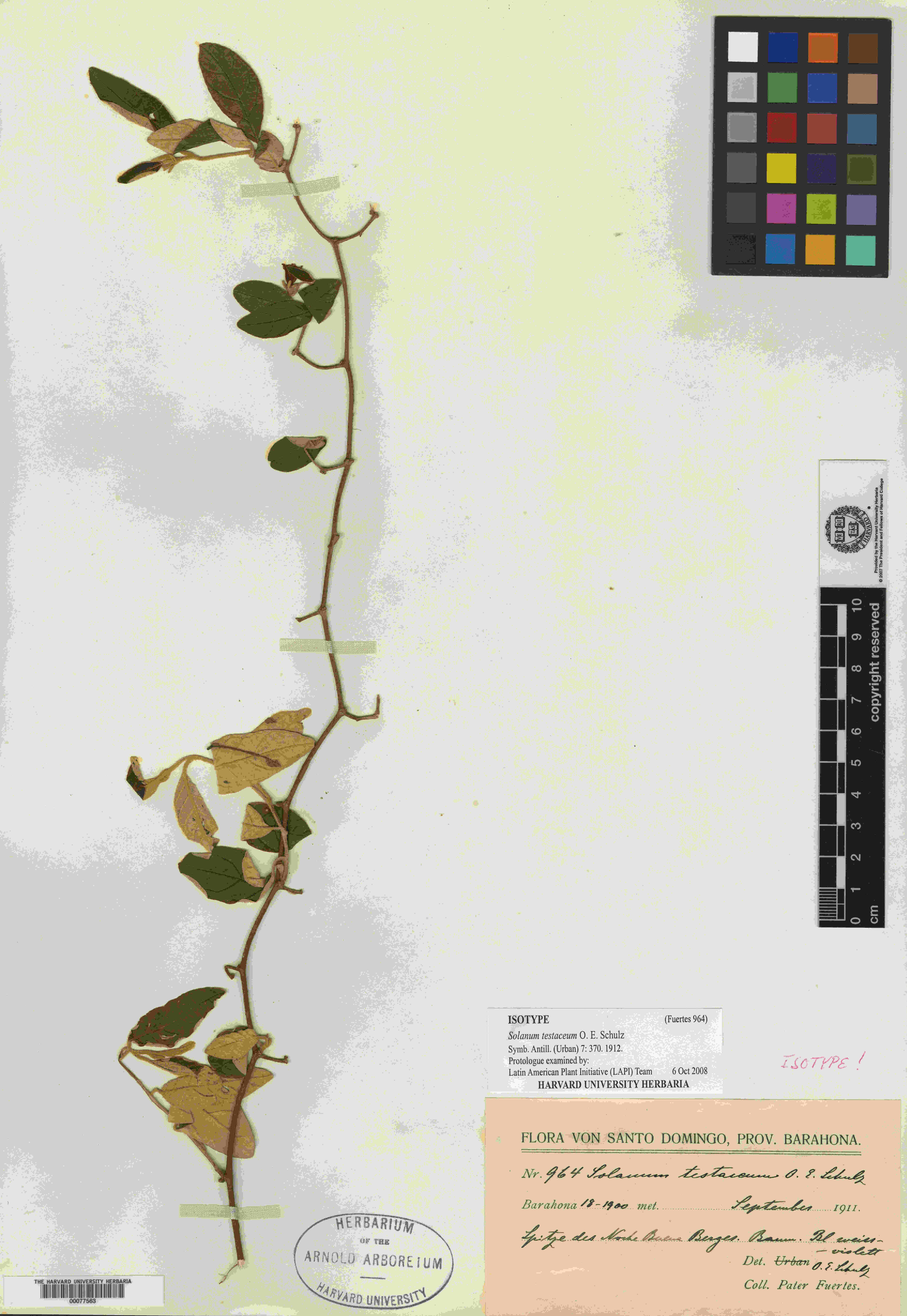

### Lycianthes_testacea_2515682512.jpeg

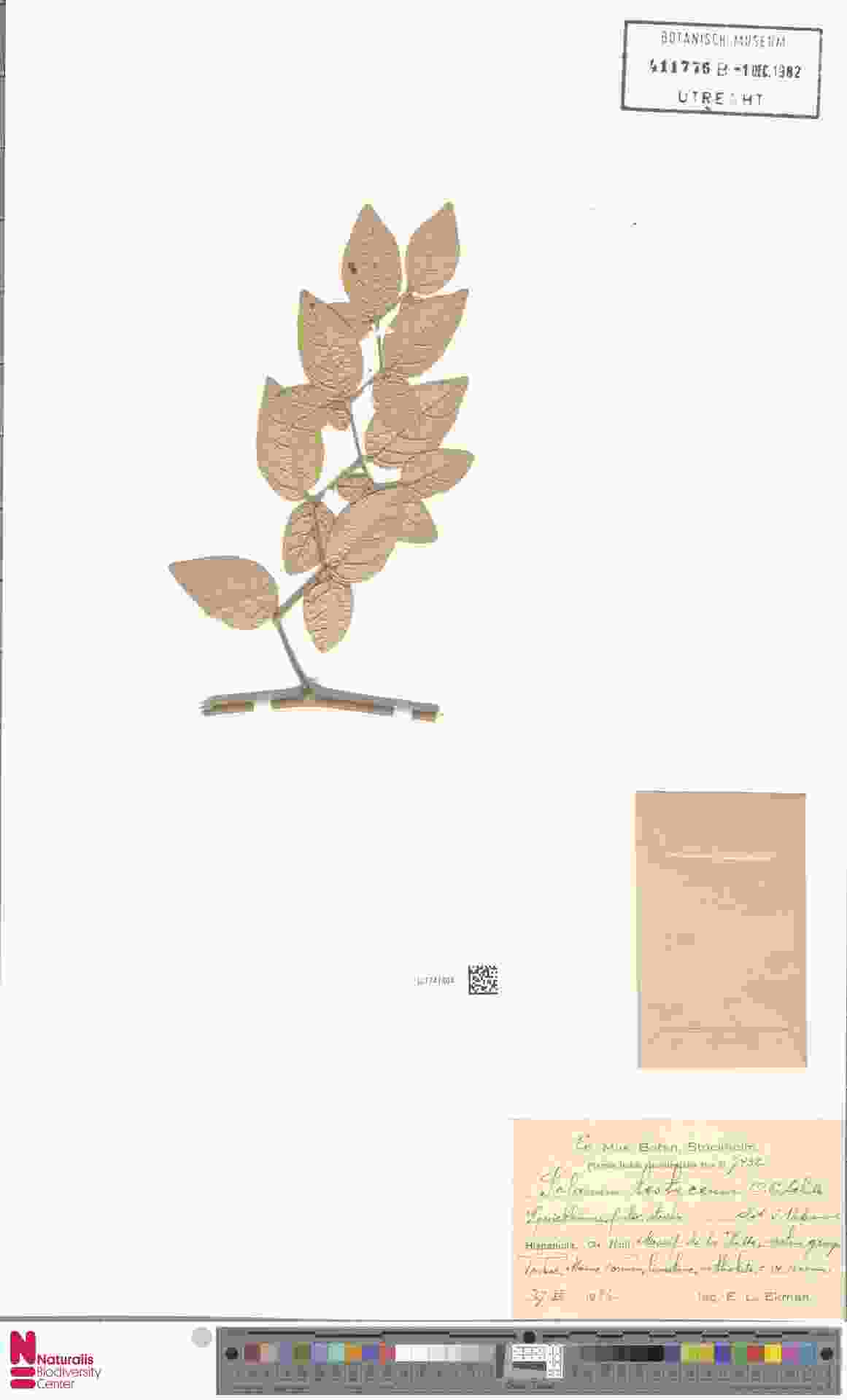

### Monopsis_kowynensis_3400983473.jpeg

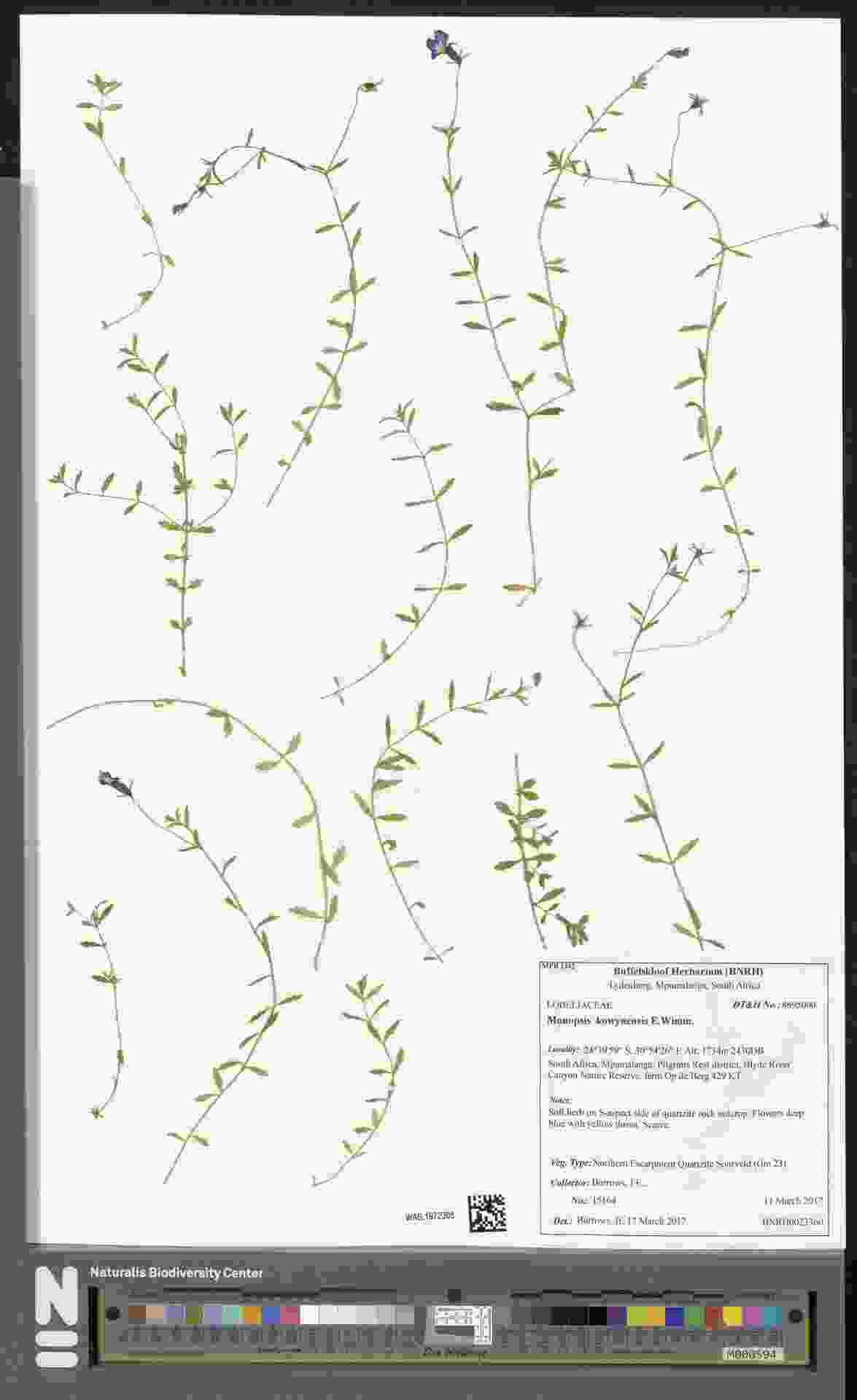

### Pavetta_gardeniifolia_1839819824.jpeg

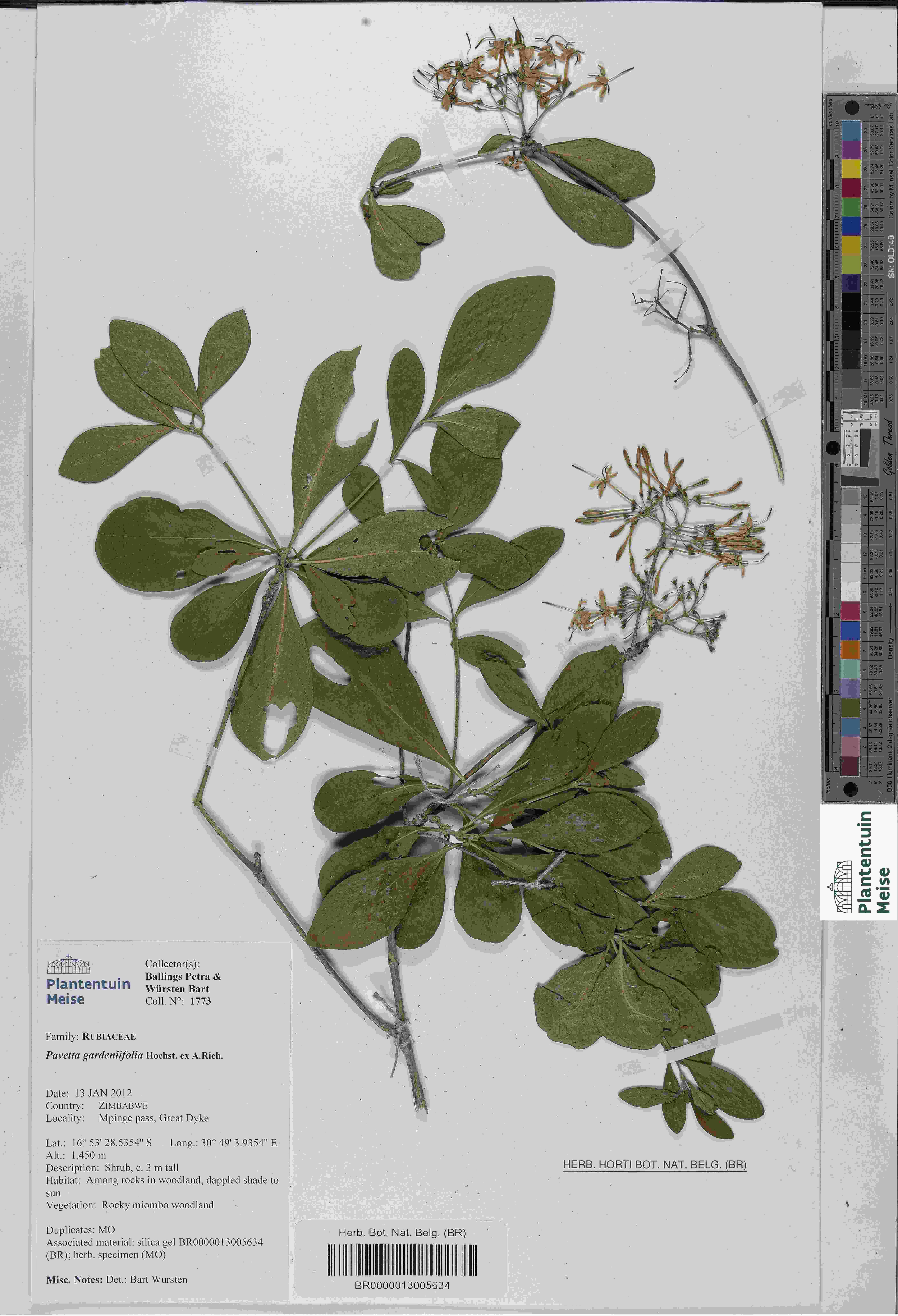

### Pavetta_gardeniifolia_1839836999.jpeg

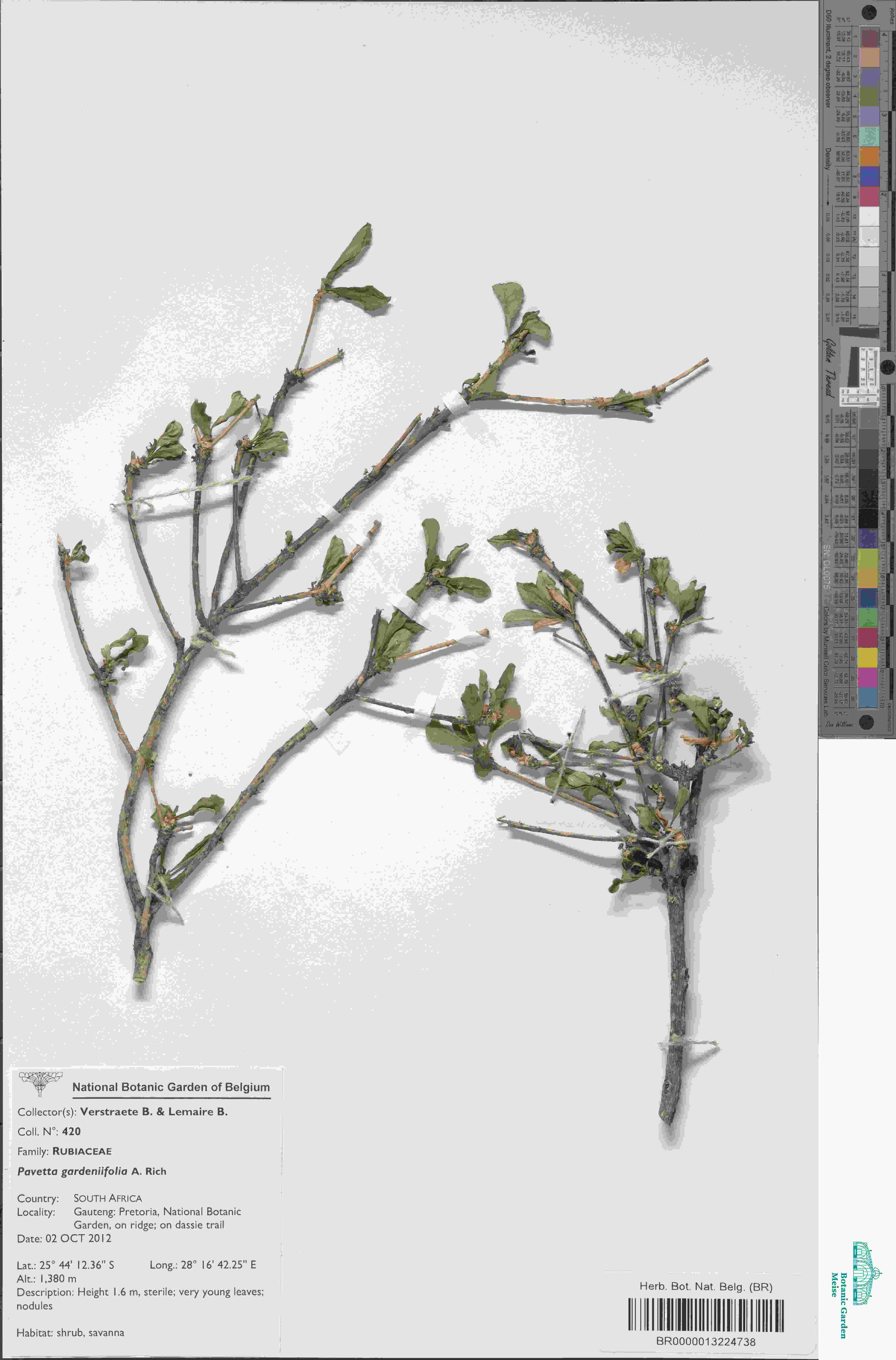

### Pavetta_gardeniifolia_1840214194.jpeg

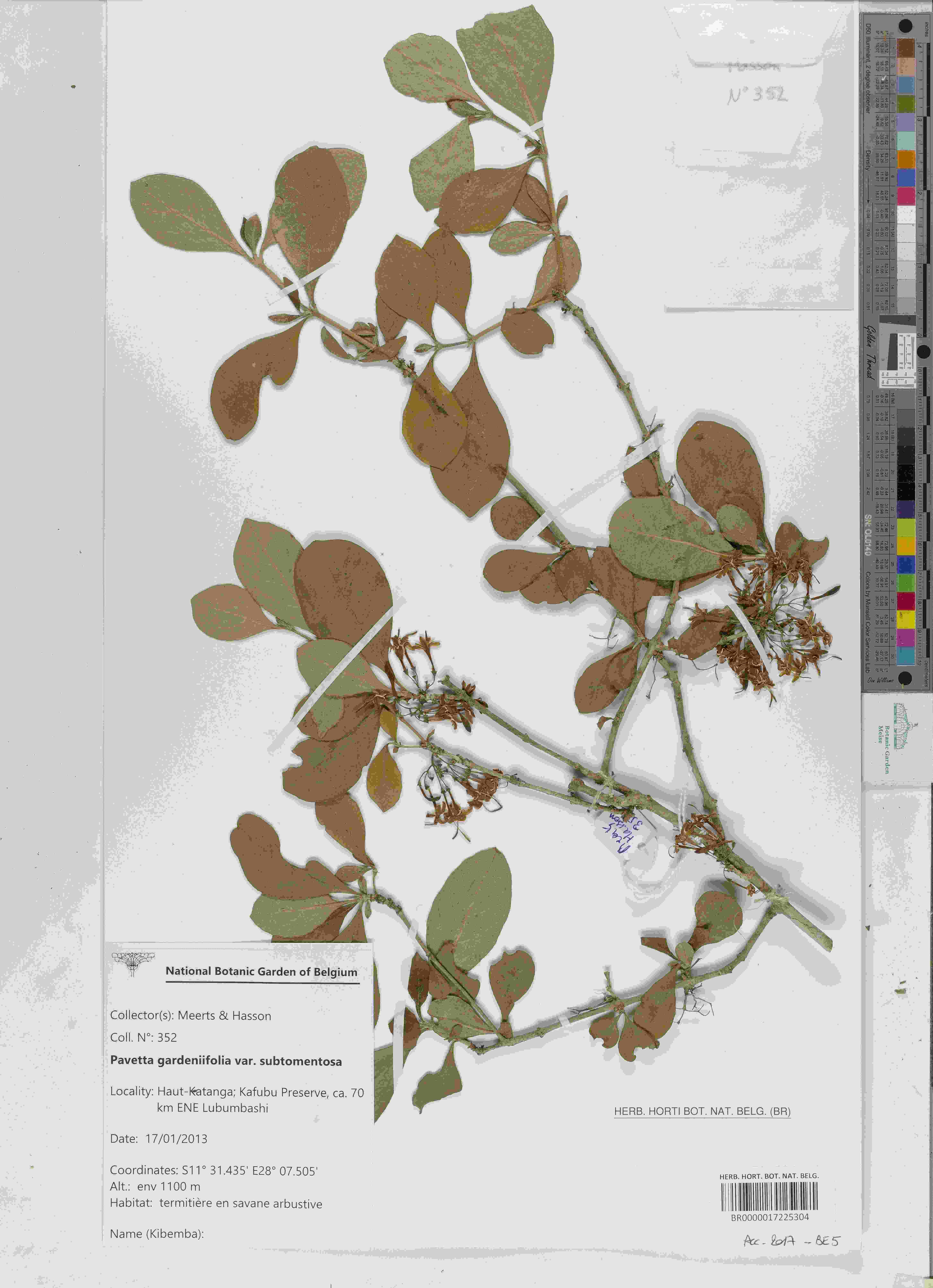

### Pavetta_gardeniifolia_3695659485.jpeg

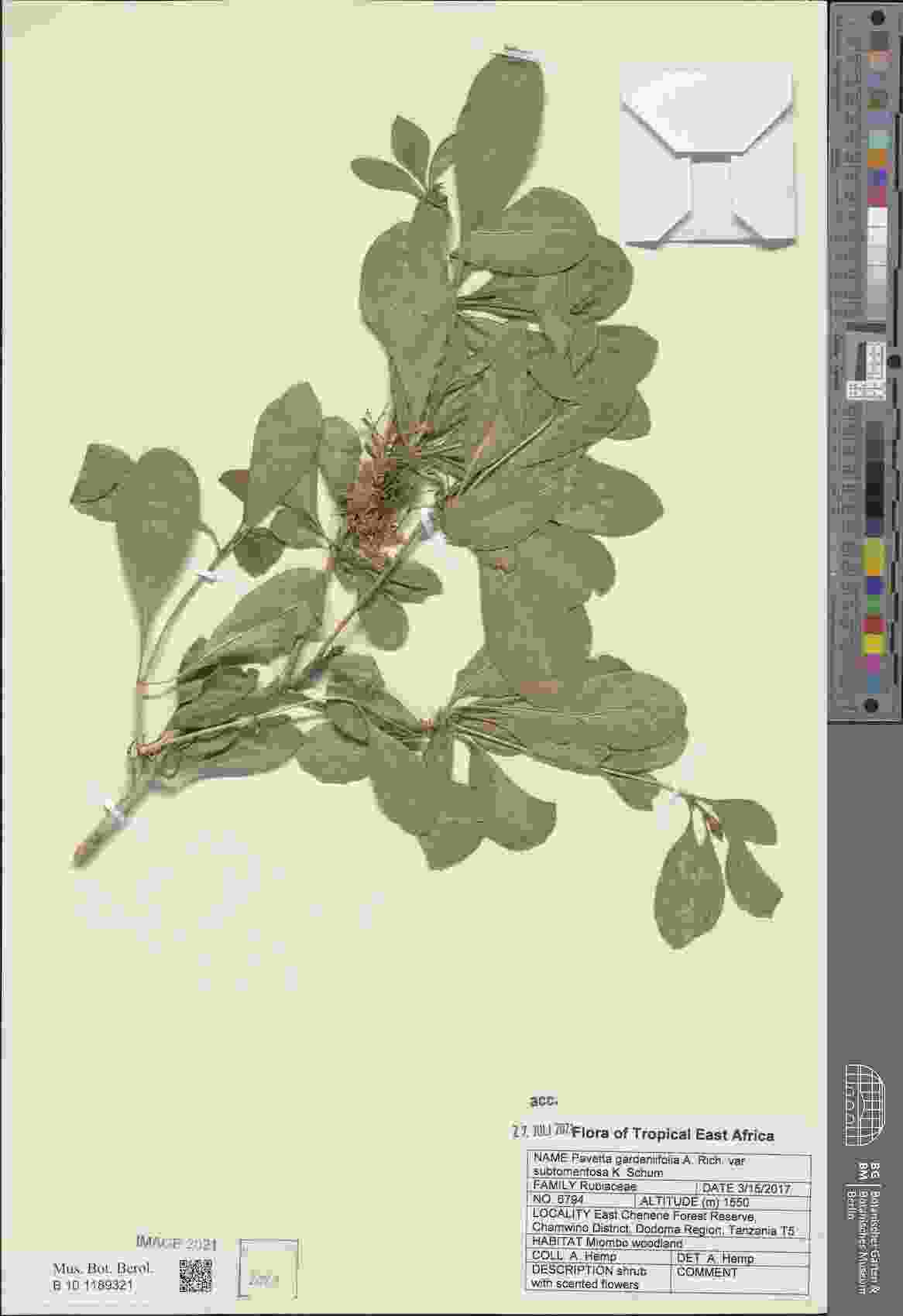

### Pavetta_gardeniifolia_4072354247.jpeg

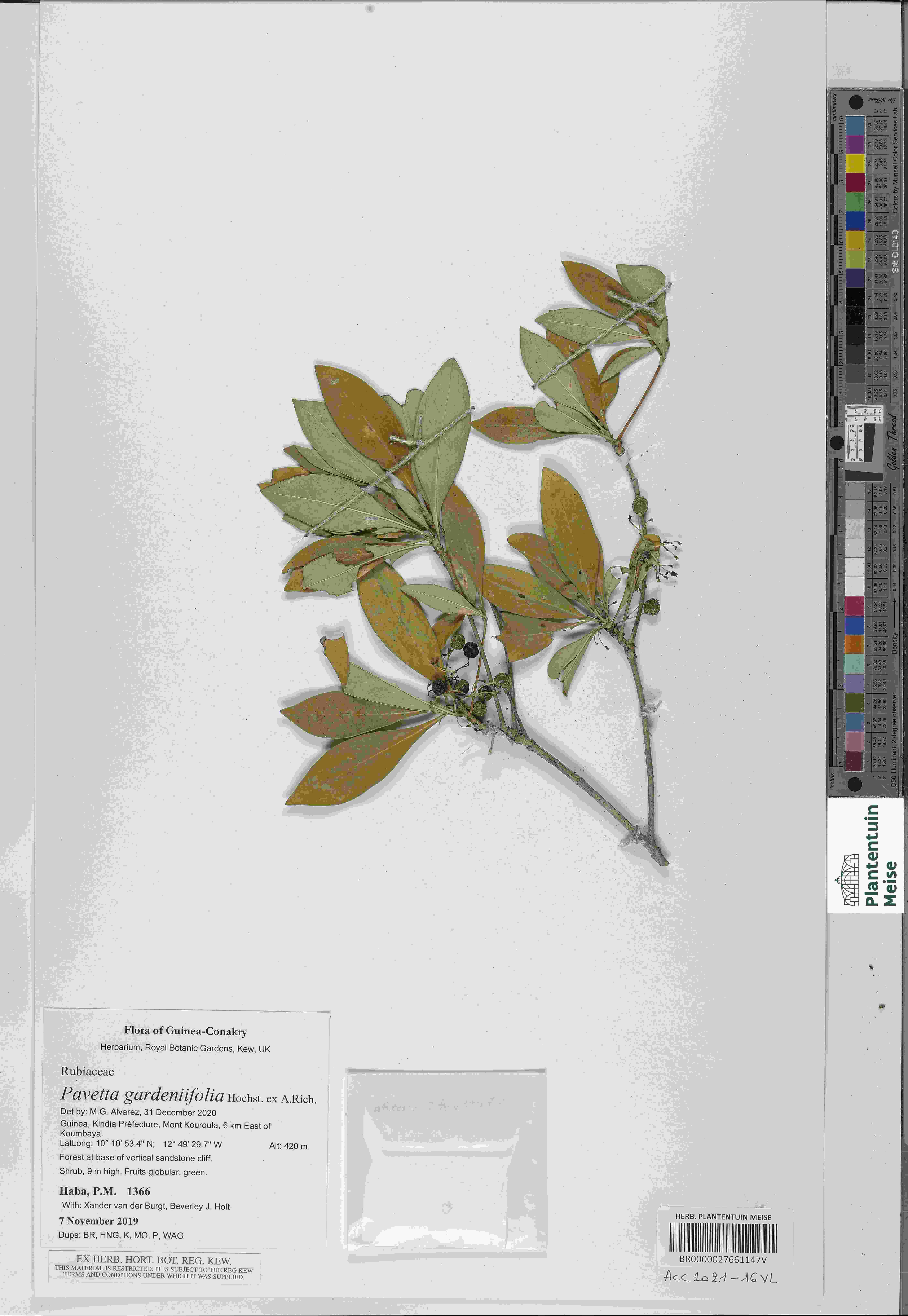

### Phyllanthus_maderaspatensis_3400968503.jpeg

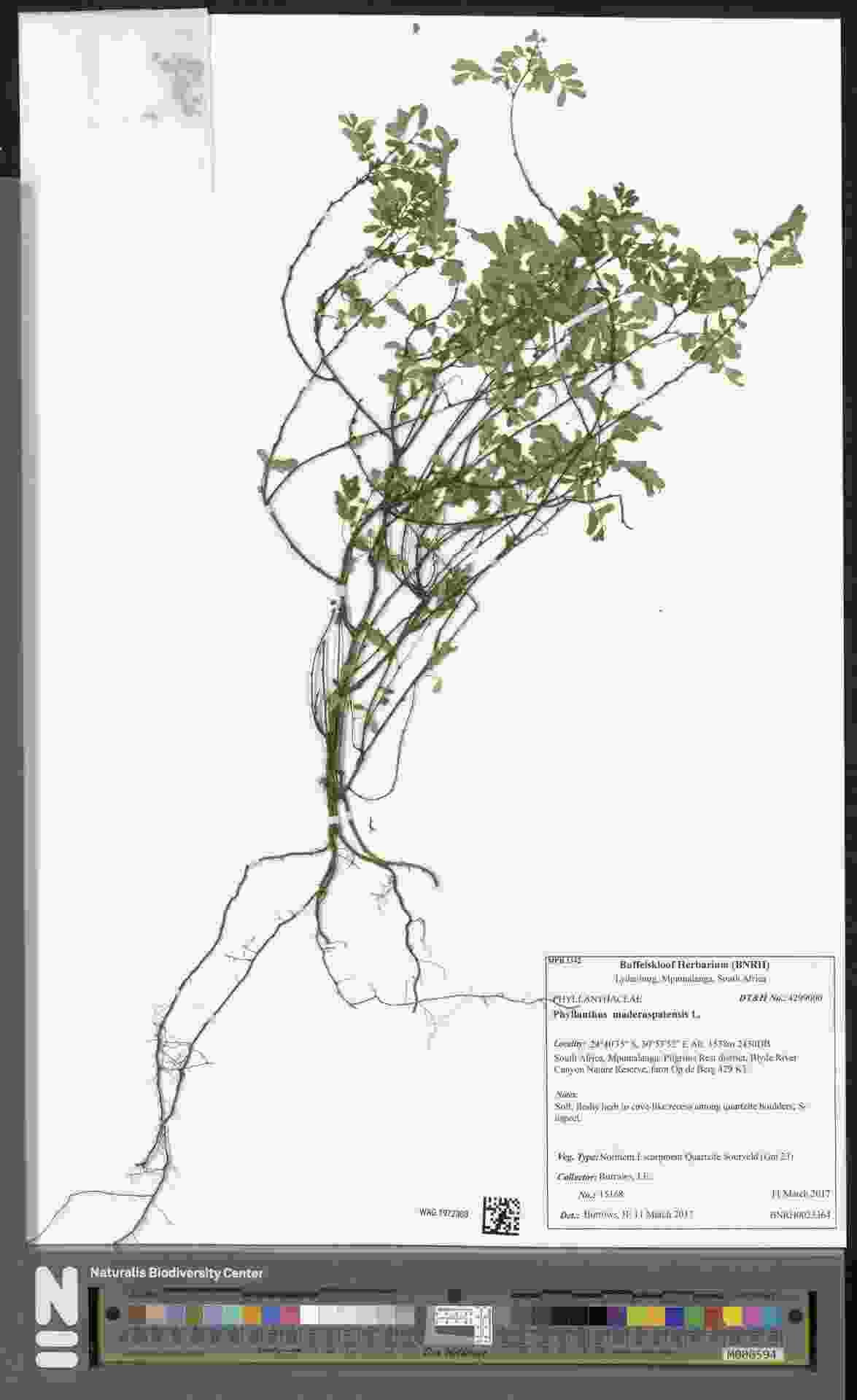

### Pinus_contorta_3991166301.jpeg

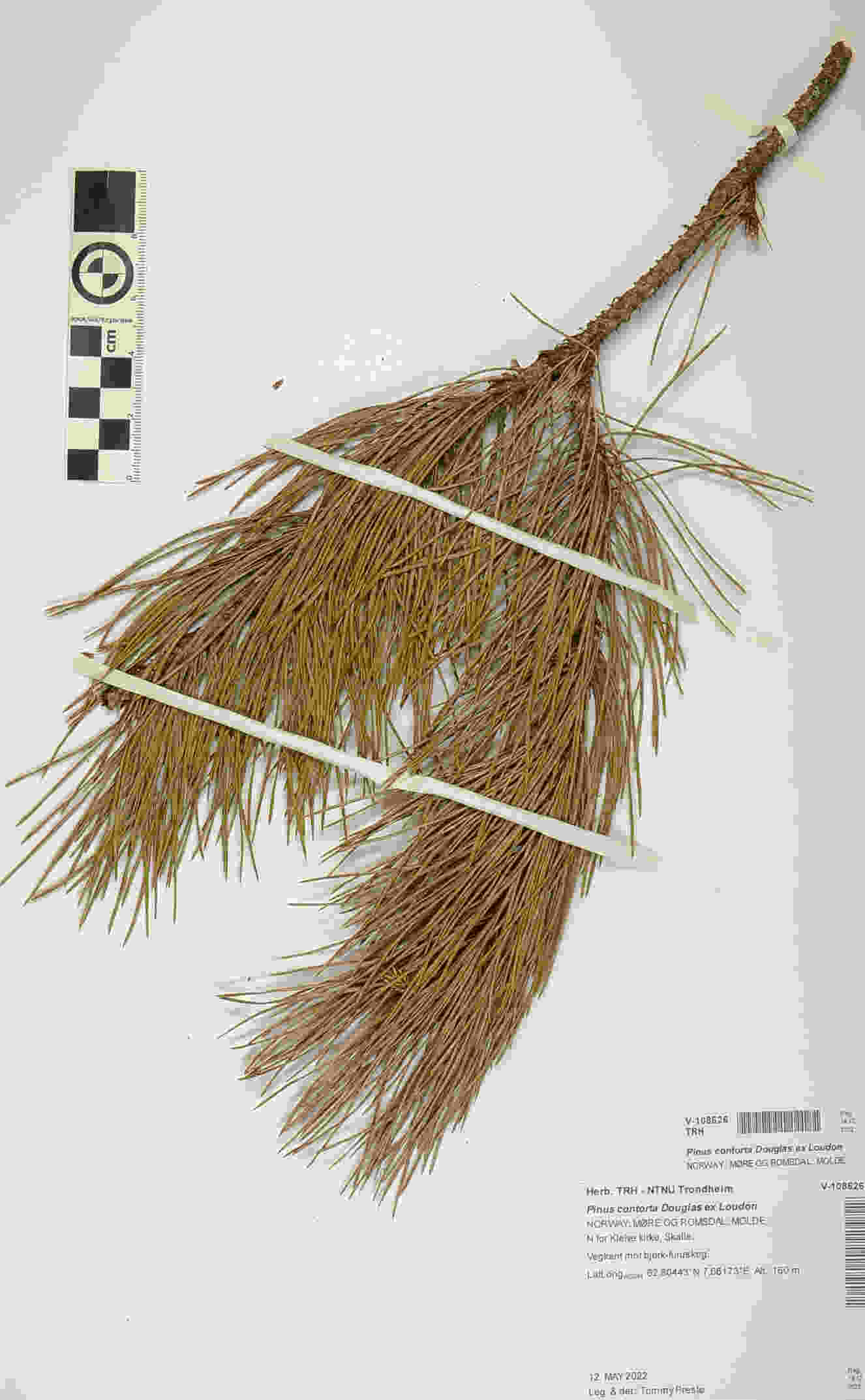

### Pinus_contorta_4608088879.jpeg

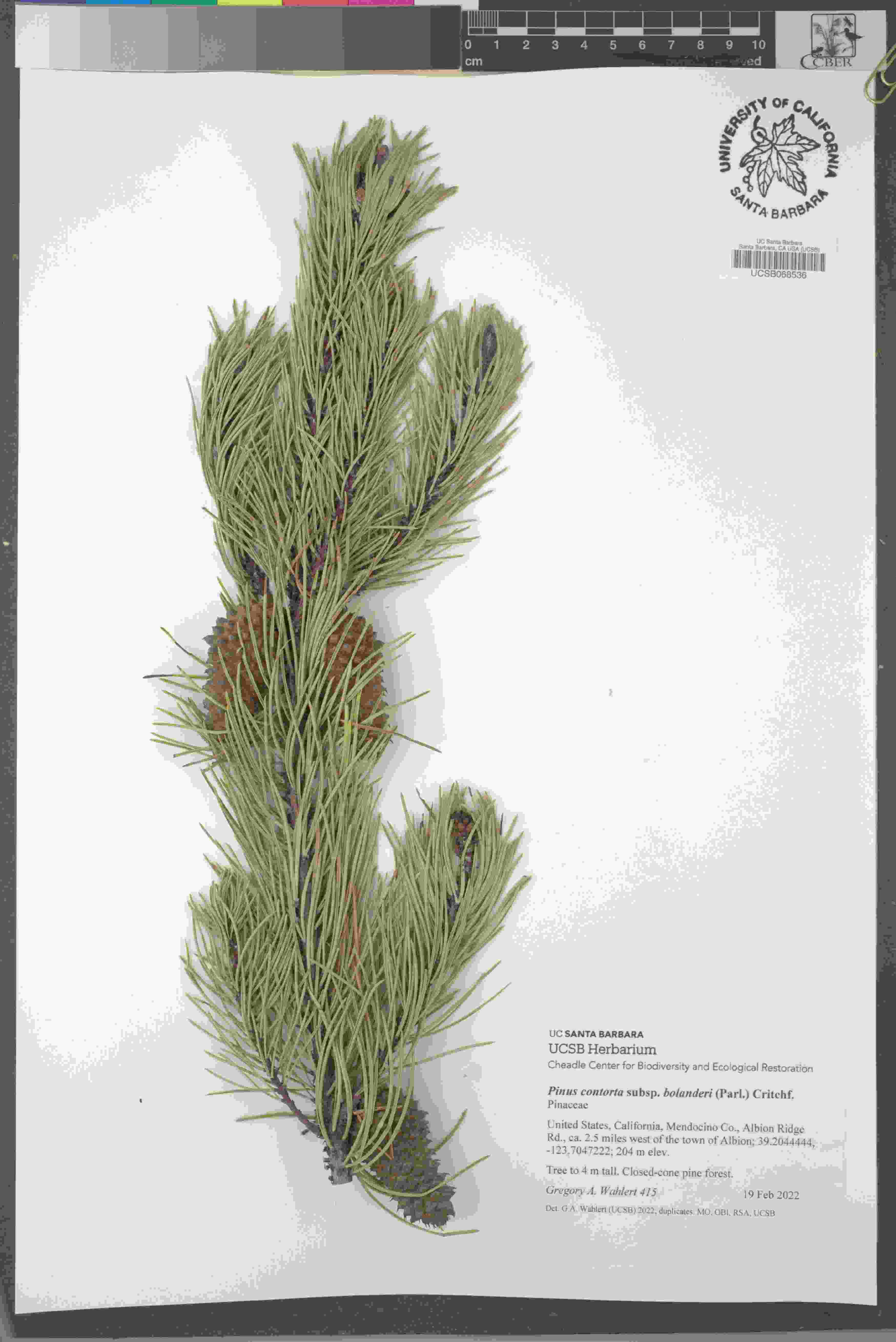

### Pinus_contorta_4851300009.jpeg

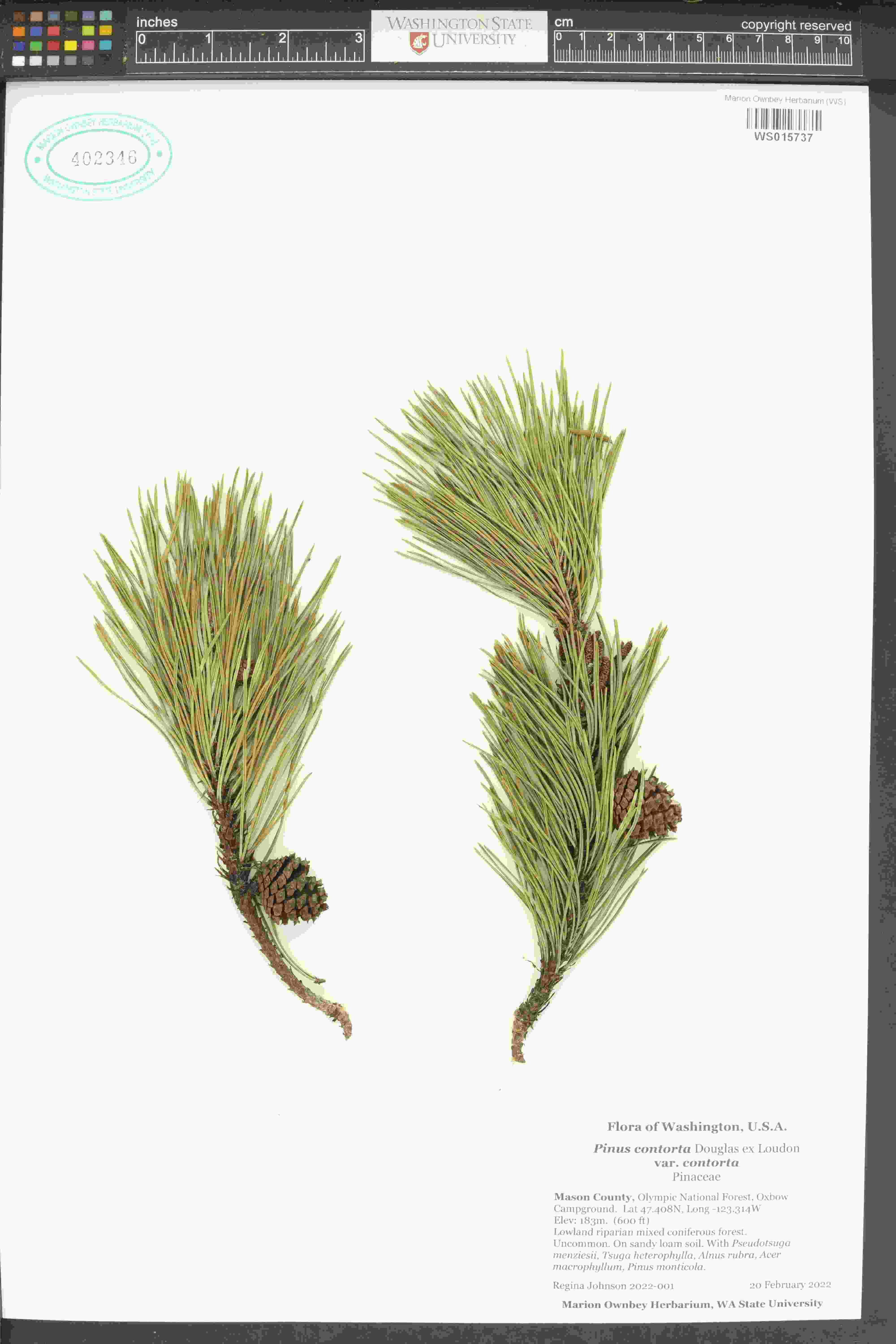

### Pinus_contorta_4871967302.jpeg

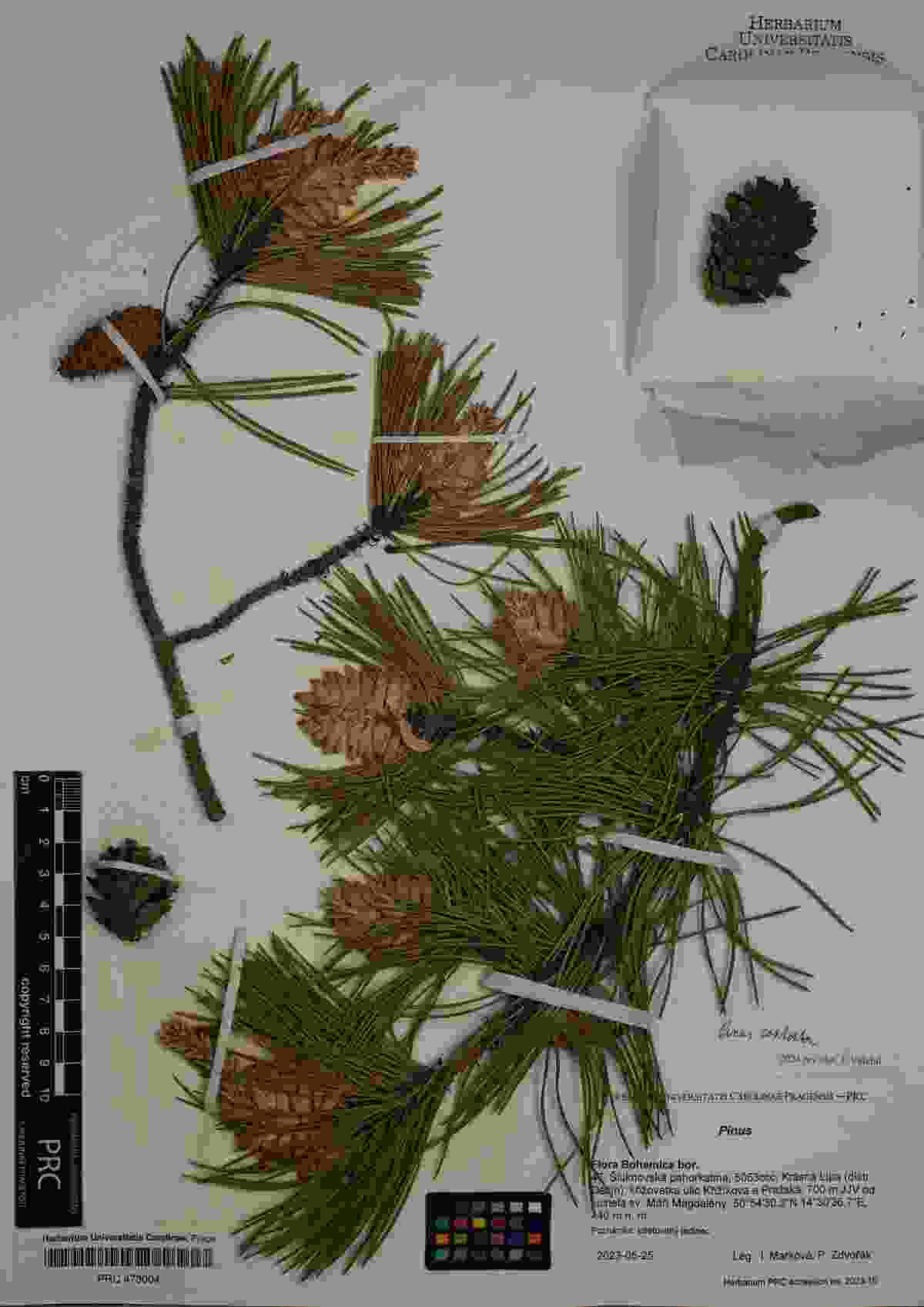

### Pinus_contorta_4912281621.jpeg

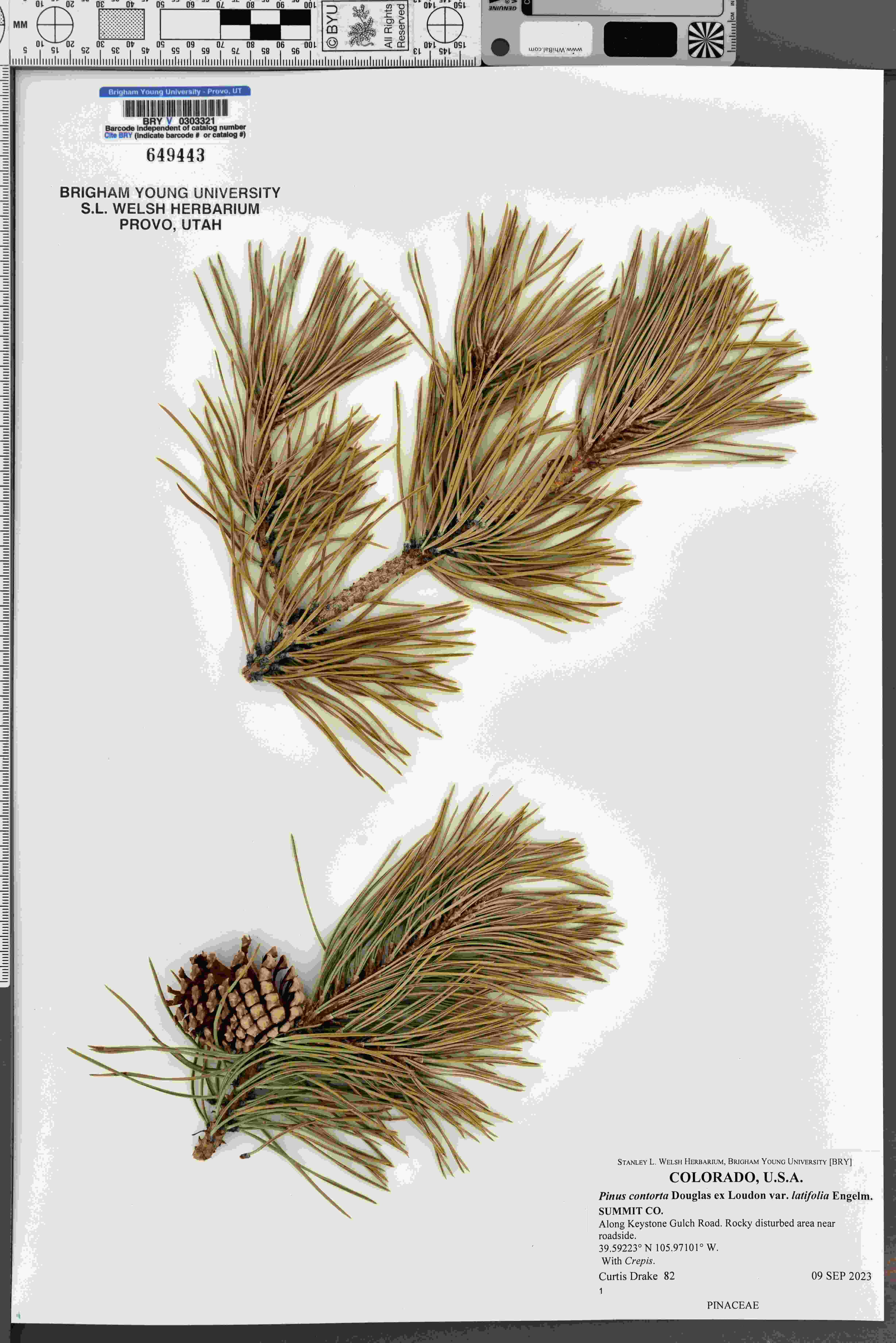

### Pinus_mugo_4903561838.jpeg

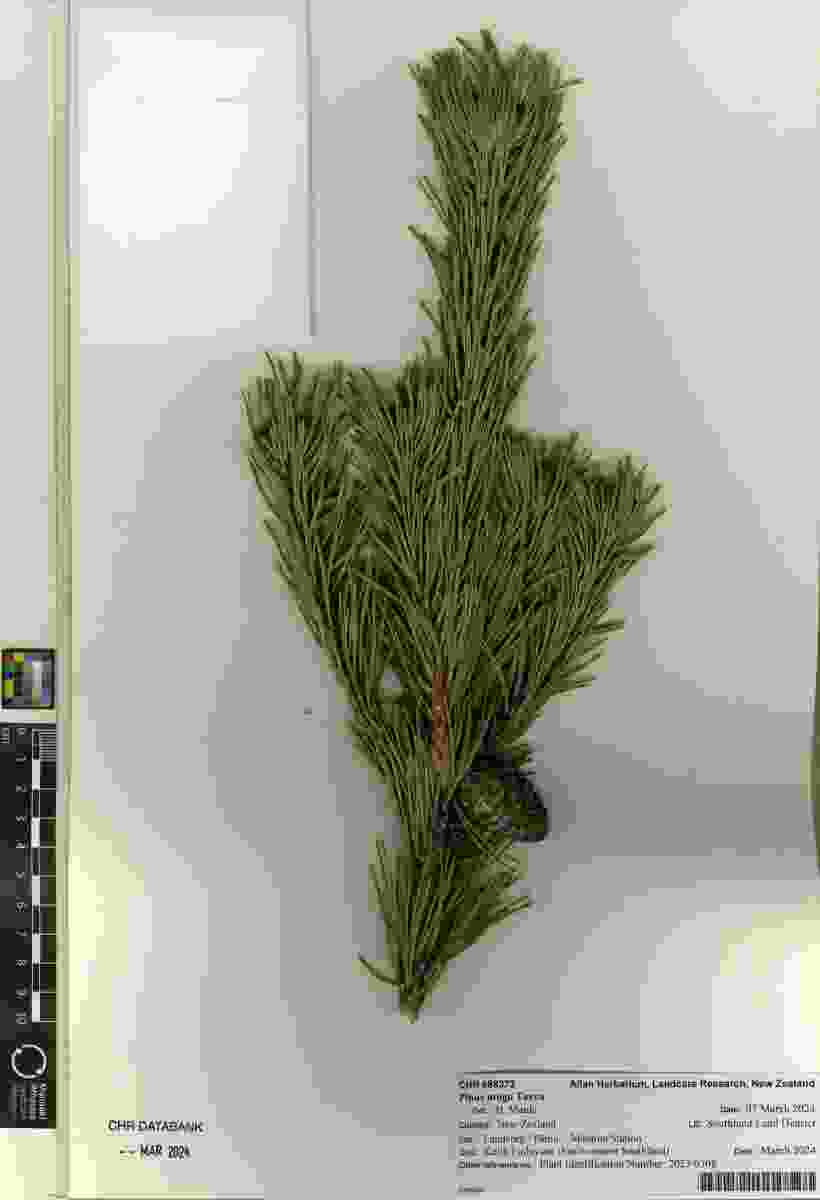

### Piper_staminodiferum_439005812.jpeg

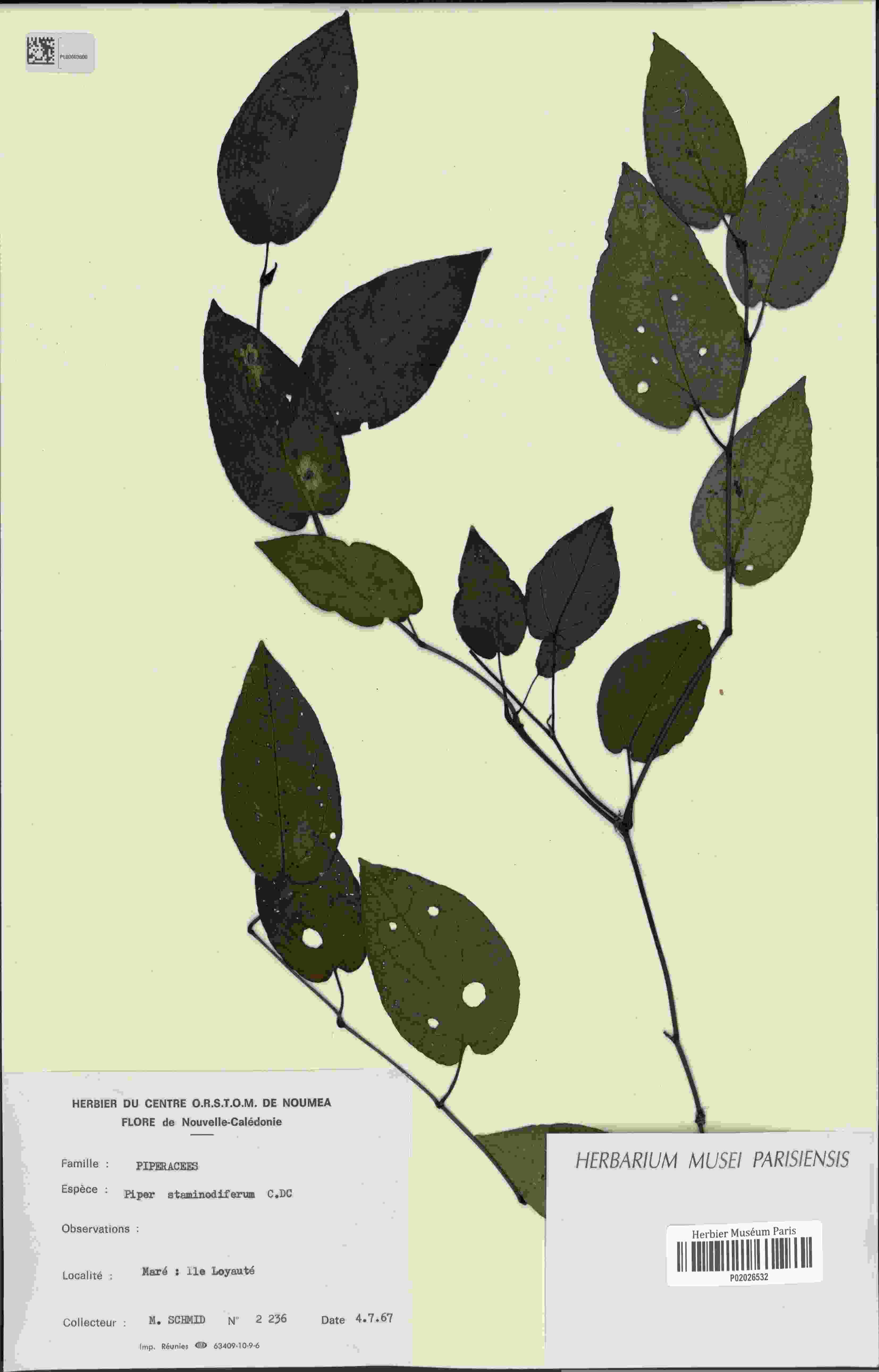

### Pithecellobium_lanceolatum_4441028404.jpeg

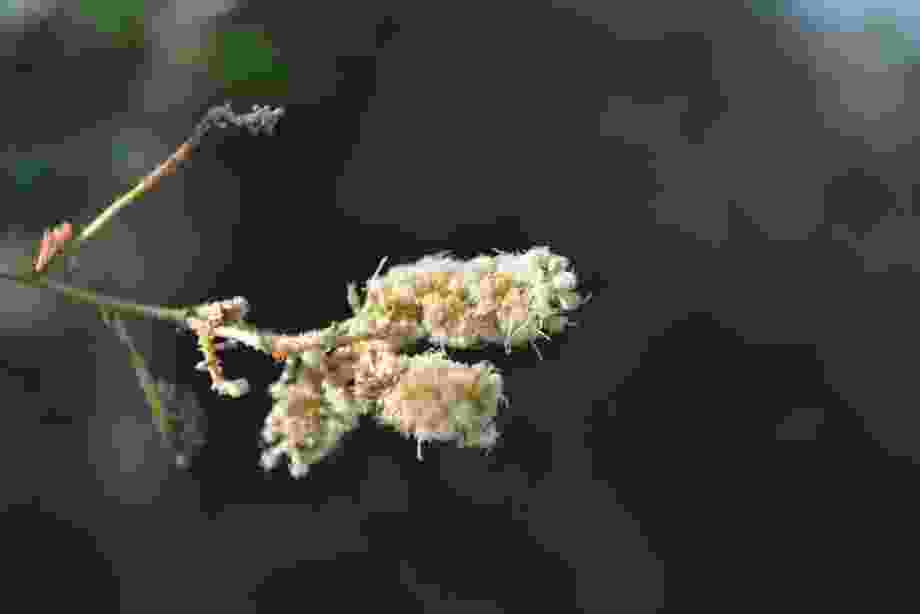

### Pithecellobium_lanceolatum_4441028404_1.jpeg

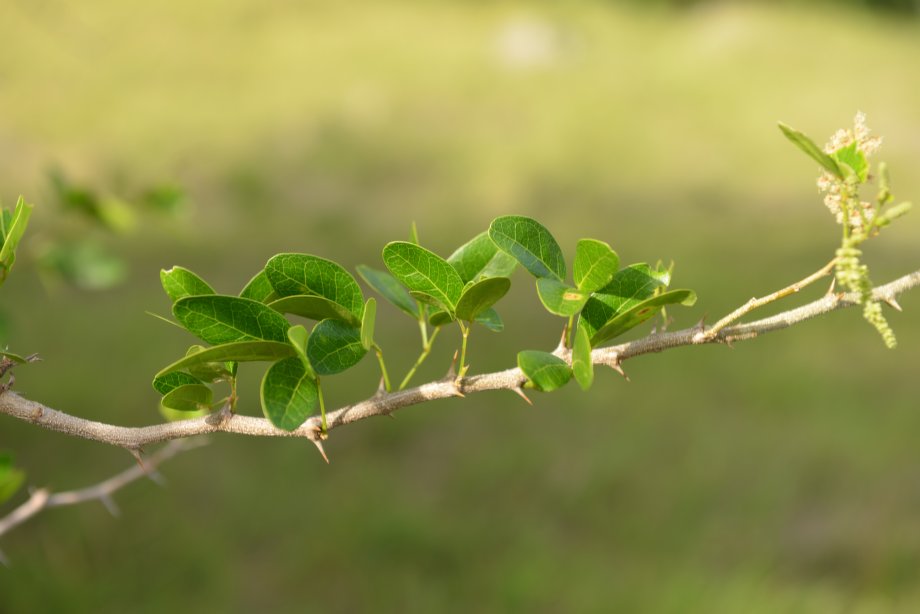

### Pithecellobium_lanceolatum_4441029617_1.jpeg

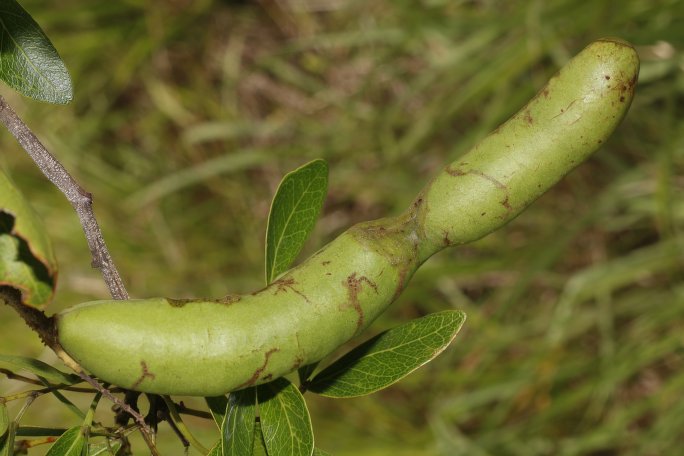

### Pithecellobium_lanceolatum_4441029617_6.jpeg

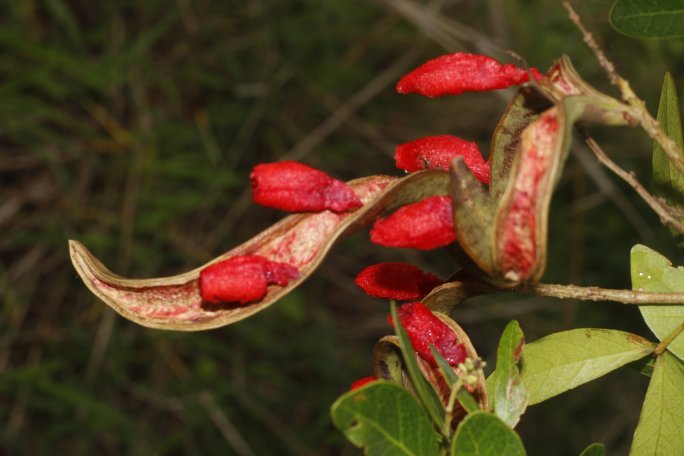

### Pithecellobium_lanceolatum_4441029617_7.jpeg

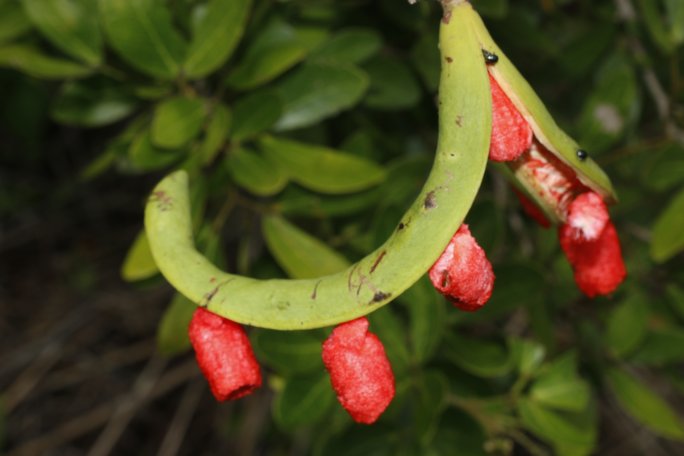

### Pithecellobium_lanceolatum_4441029617_10.jpeg

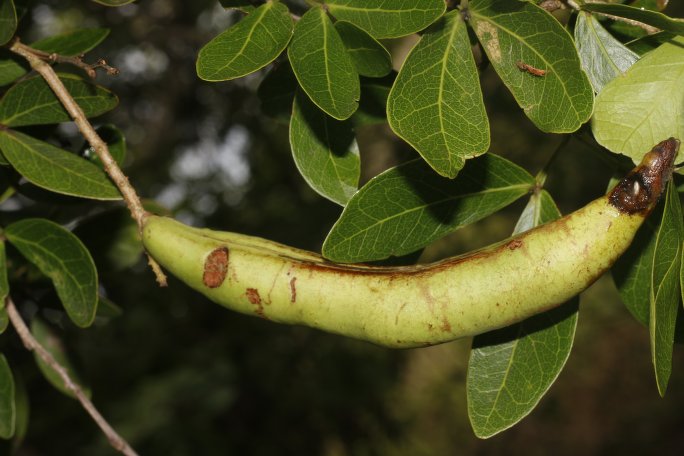

### Pithecellobium_lanceolatum_4441029617_11.jpeg

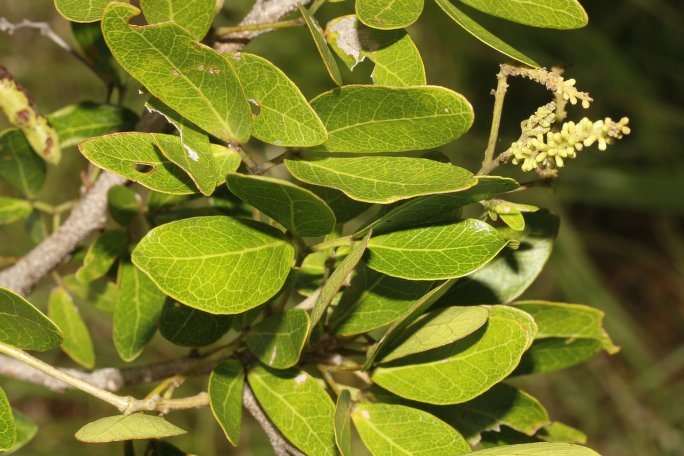

### Pithecellobium_lanceolatum_4441029617_16.jpeg

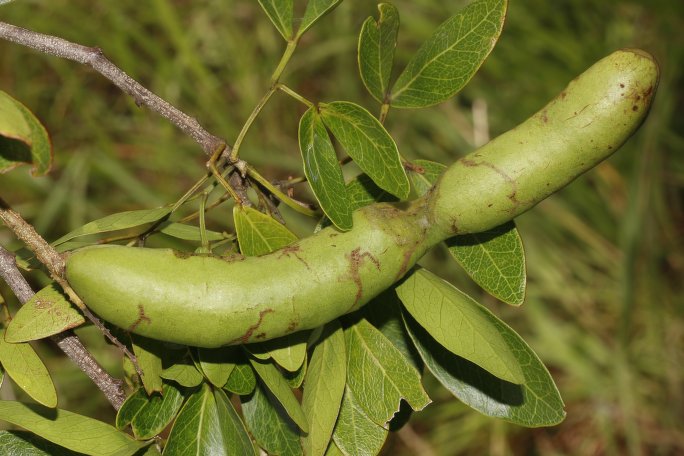

### Pithecellobium_lanceolatum_4441037521_3.jpeg

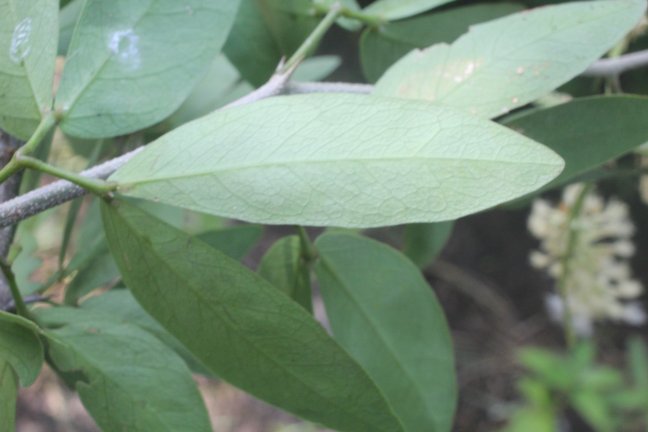

### Pithecellobium_lanceolatum_4441037521_4.jpeg

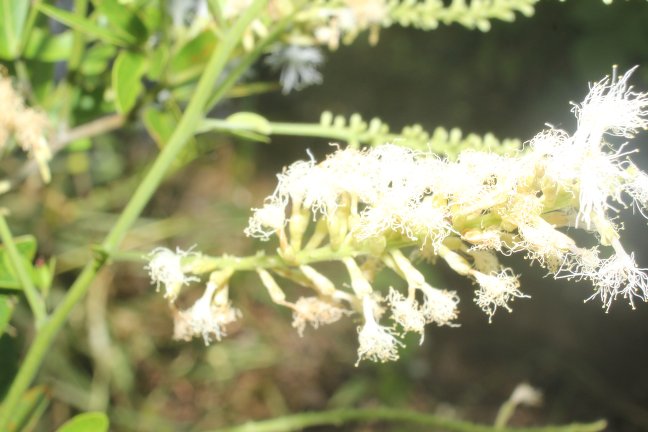

### Pithecellobium_lanceolatum_4441037521_5.jpeg

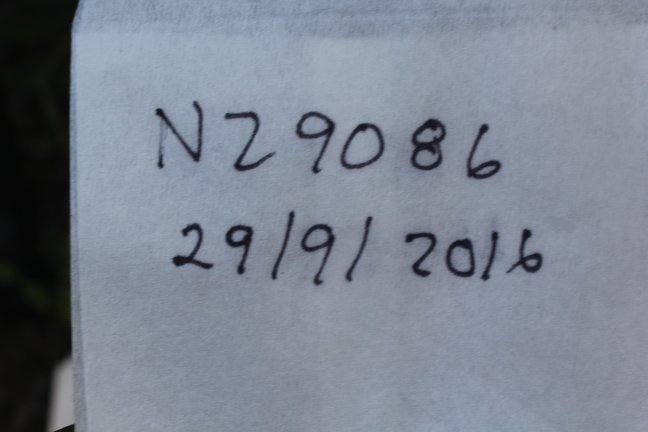

### Pithecellobium_lanceolatum_4441037521_8.jpeg

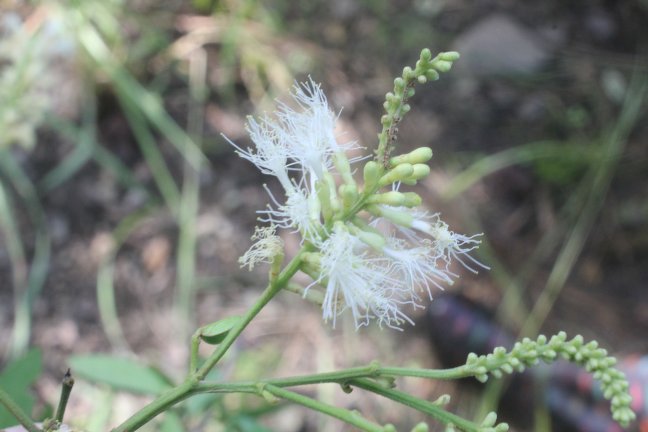

### Pithecellobium_lanceolatum_4441037521_9.jpeg

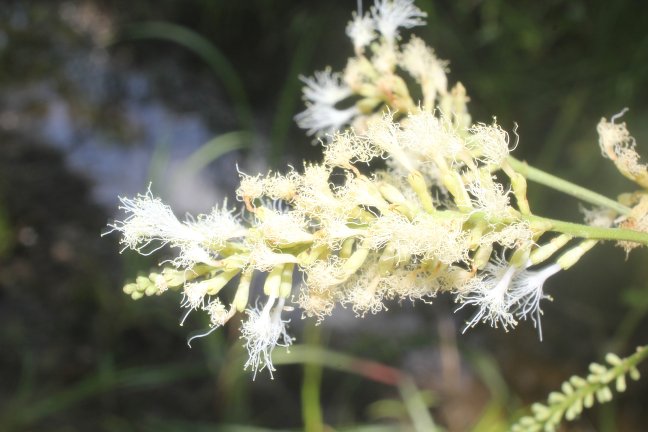

### Pithecellobium_lanceolatum_4441037521_12.jpeg

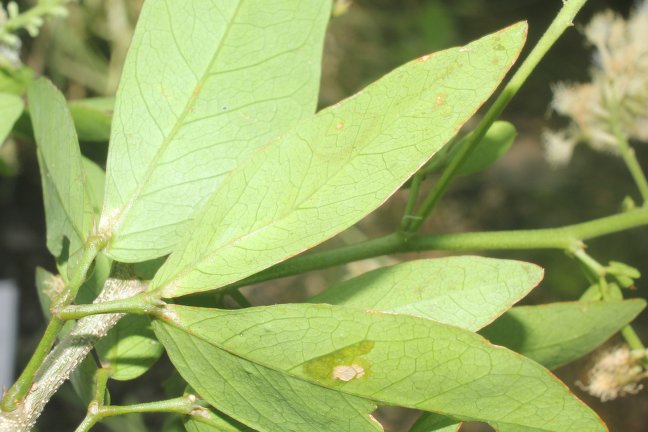

### Pithecellobium_lanceolatum_4441037521_14.jpeg

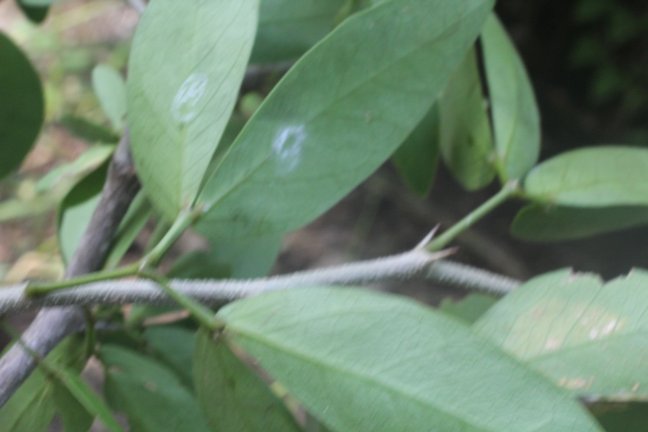

### Pithecellobium_lanceolatum_4441037521_15.jpeg

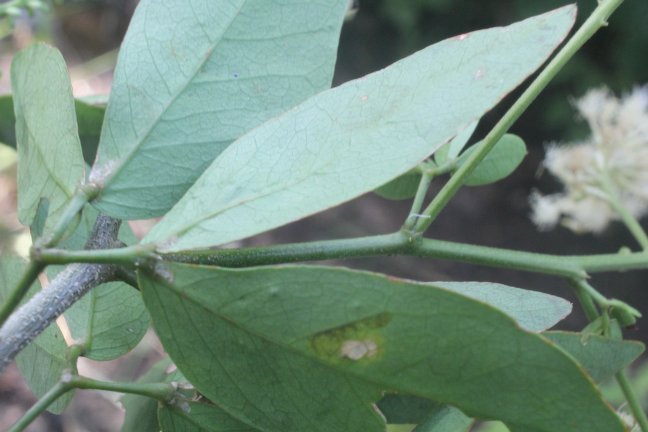

### Pithecellobium_lanceolatum_4441043496.jpeg

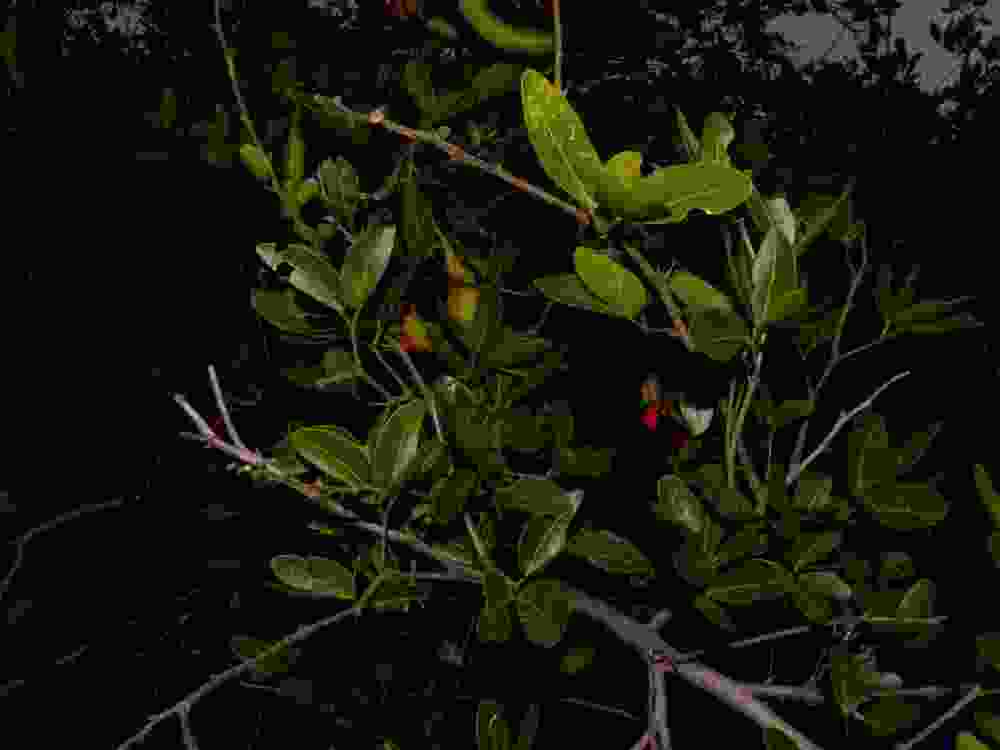

### Pithecellobium_lanceolatum_4441043496_2.jpeg

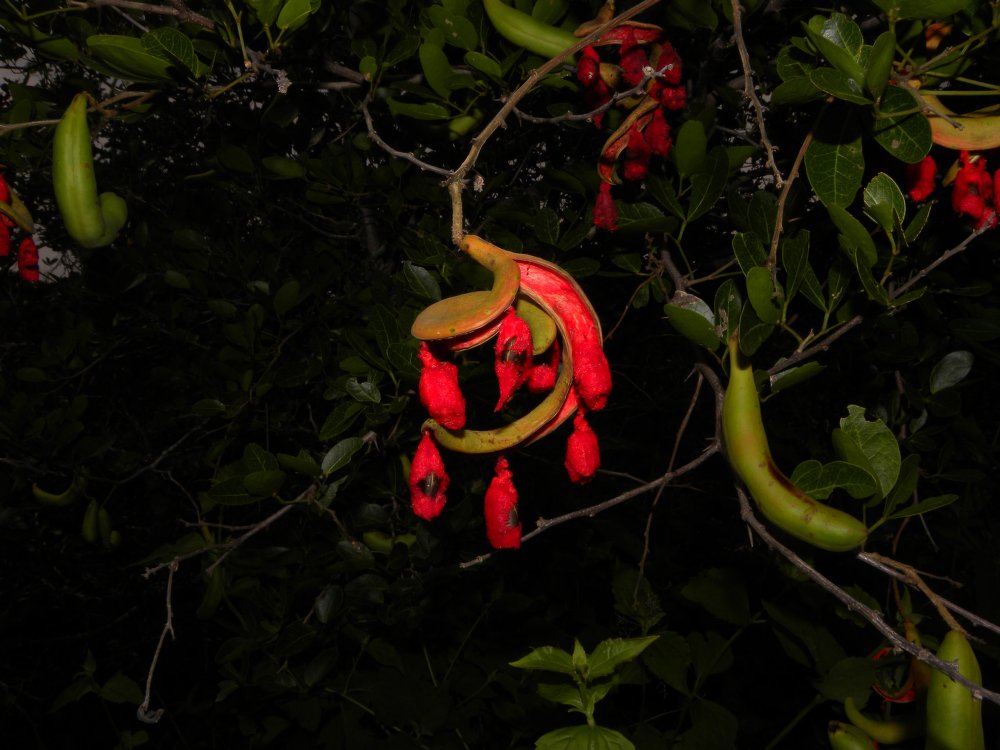

### Pithecellobium_lanceolatum_4441043496_3.jpeg

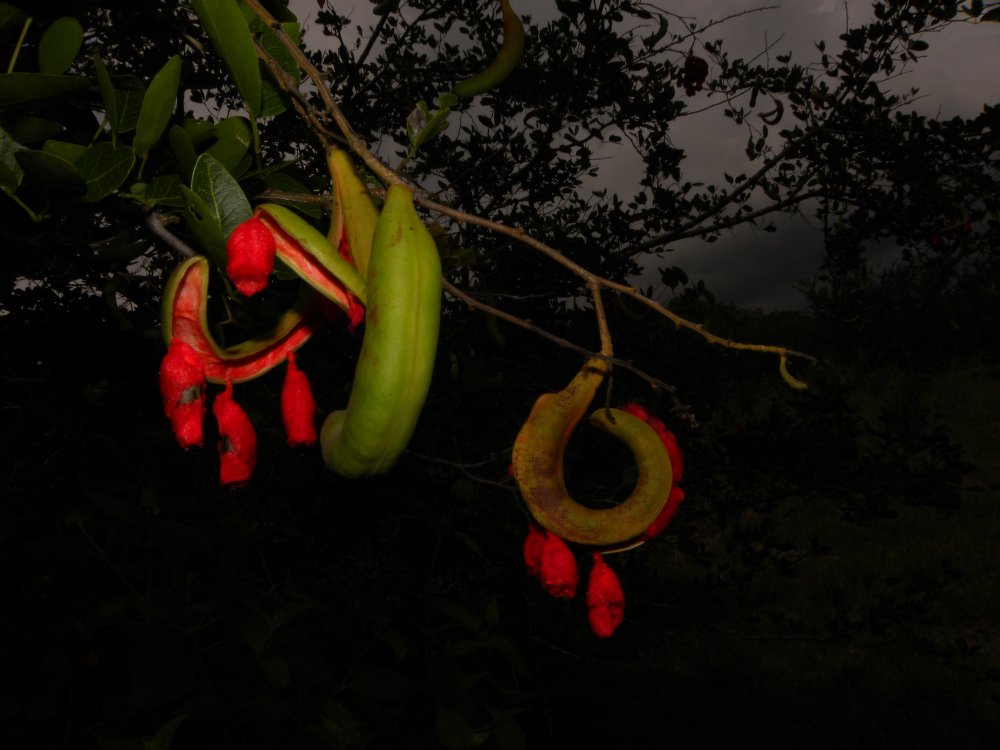

### Pithecellobium_lanceolatum_4441043496_4.jpeg

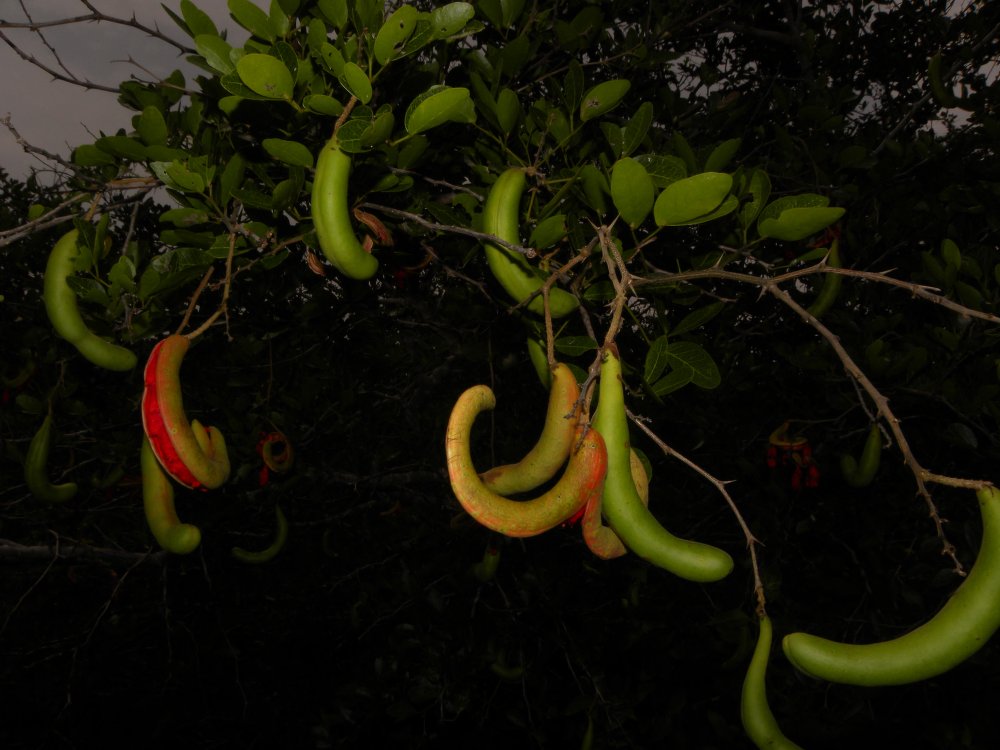

### Scilla_siberica_4132503517.jpeg

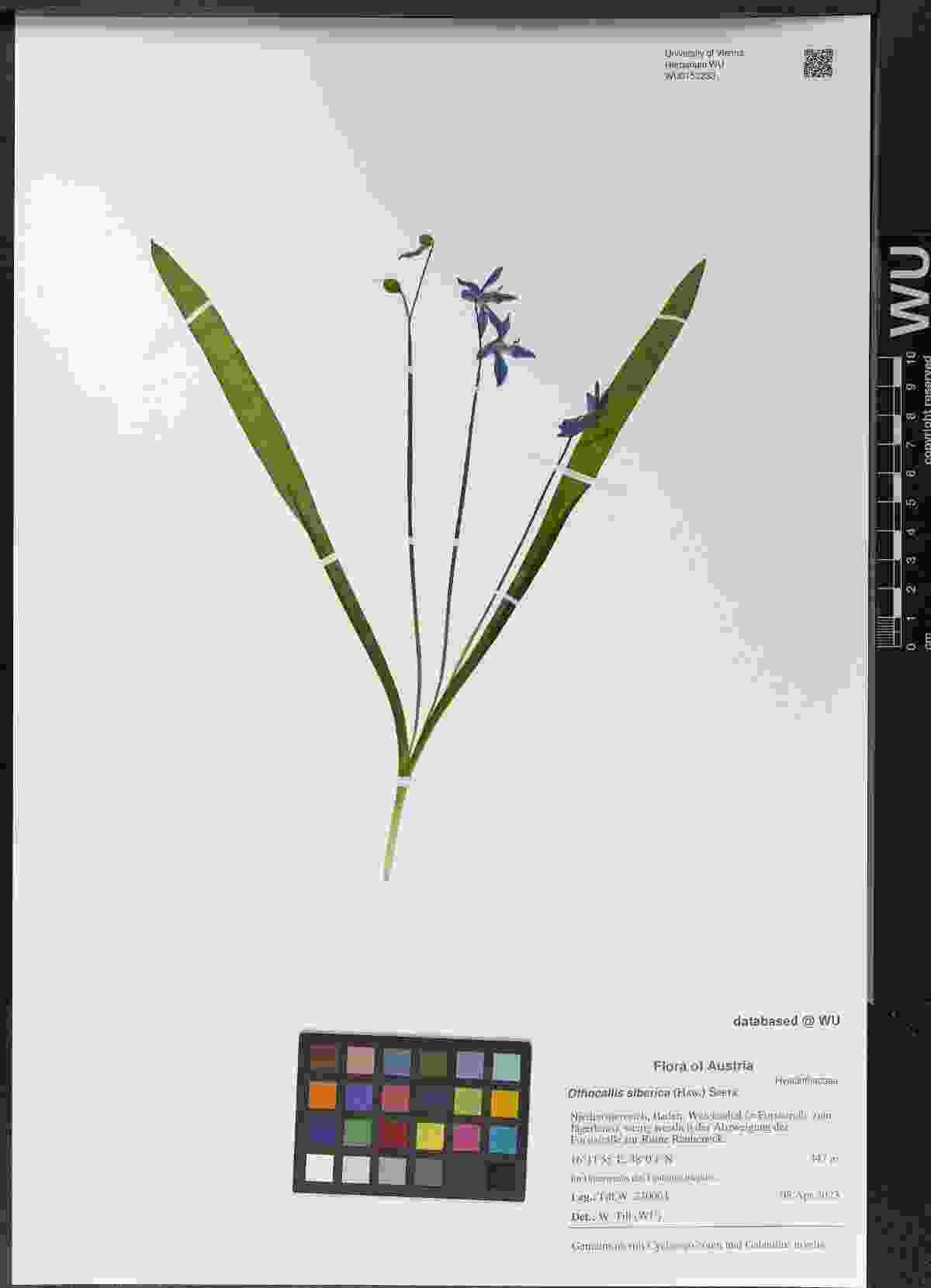

### Scilla_siberica_4134375439.jpeg

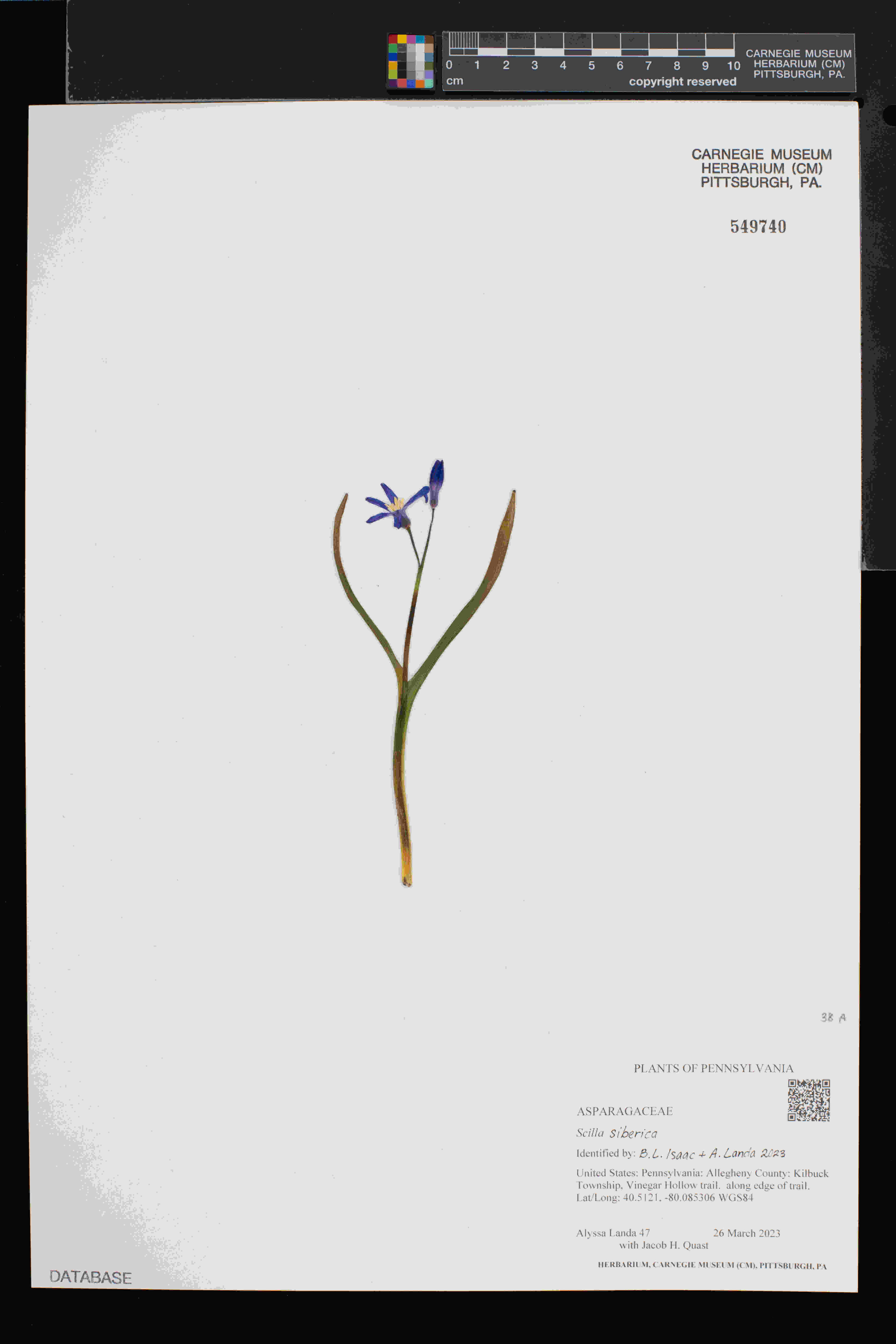

### Scilla_siberica_4856085645_2.jpeg

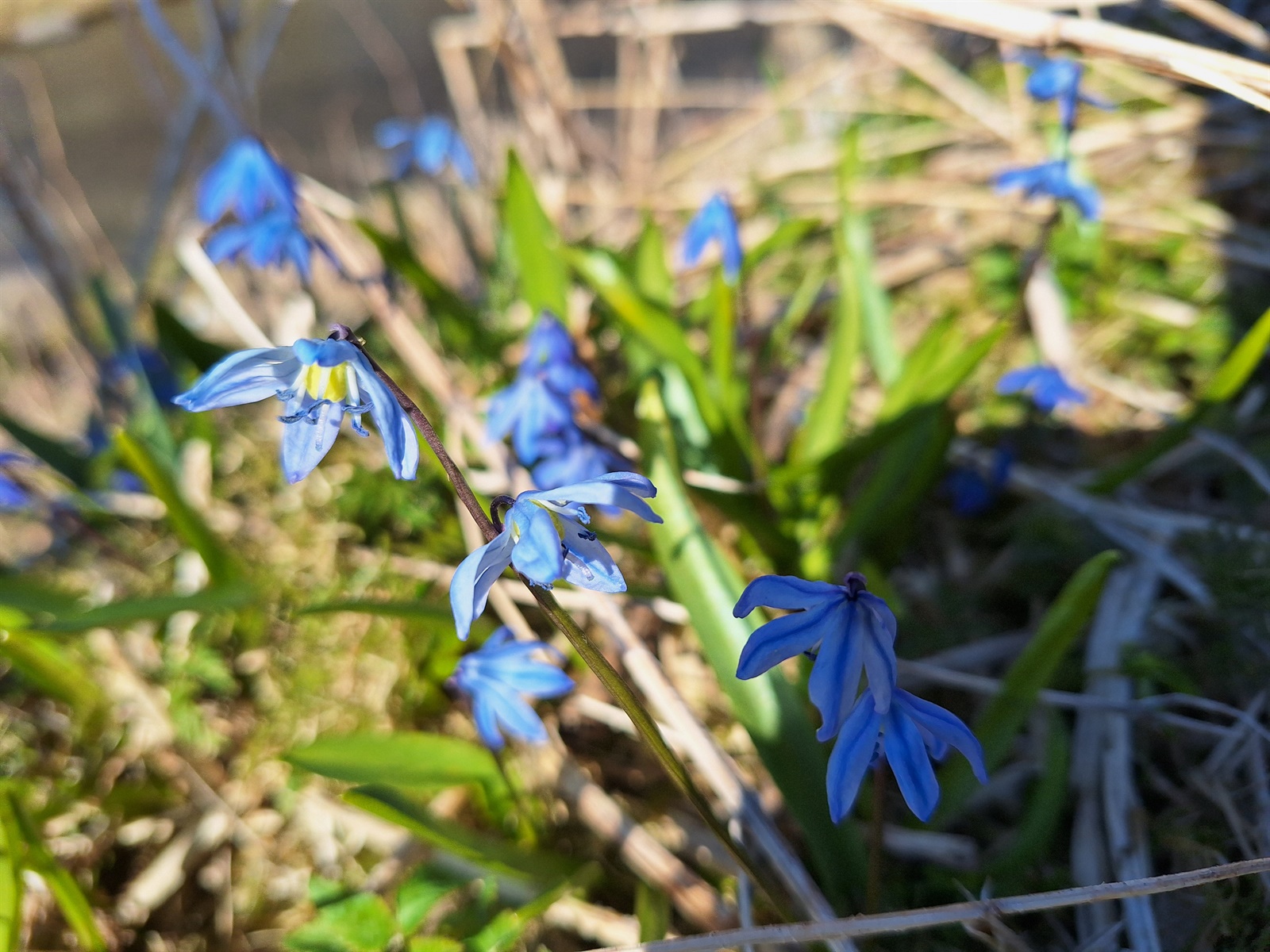

### Senecio_gallicus_2515073250.jpeg

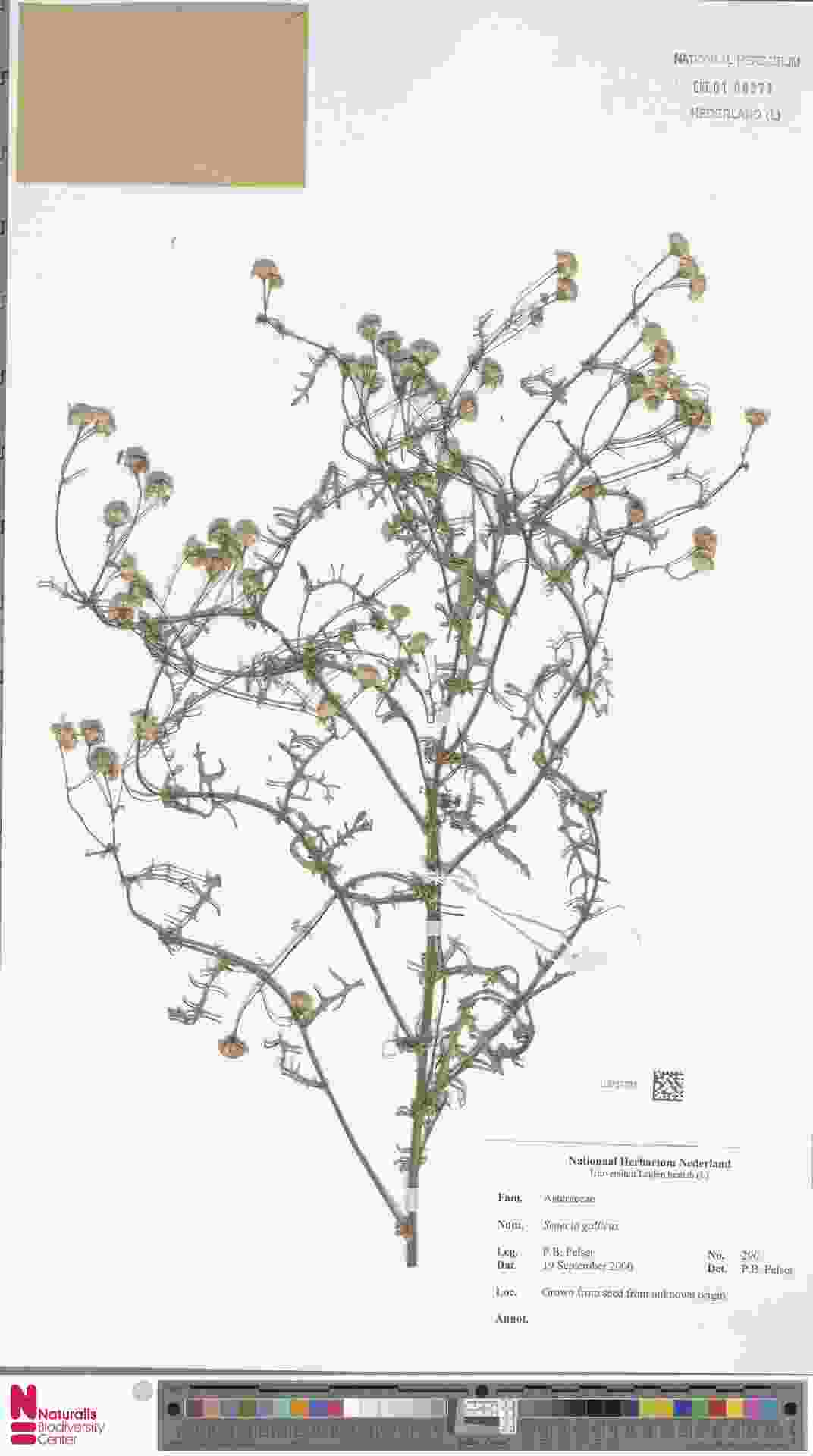

### Senecio_gallicus_2598788914.jpeg

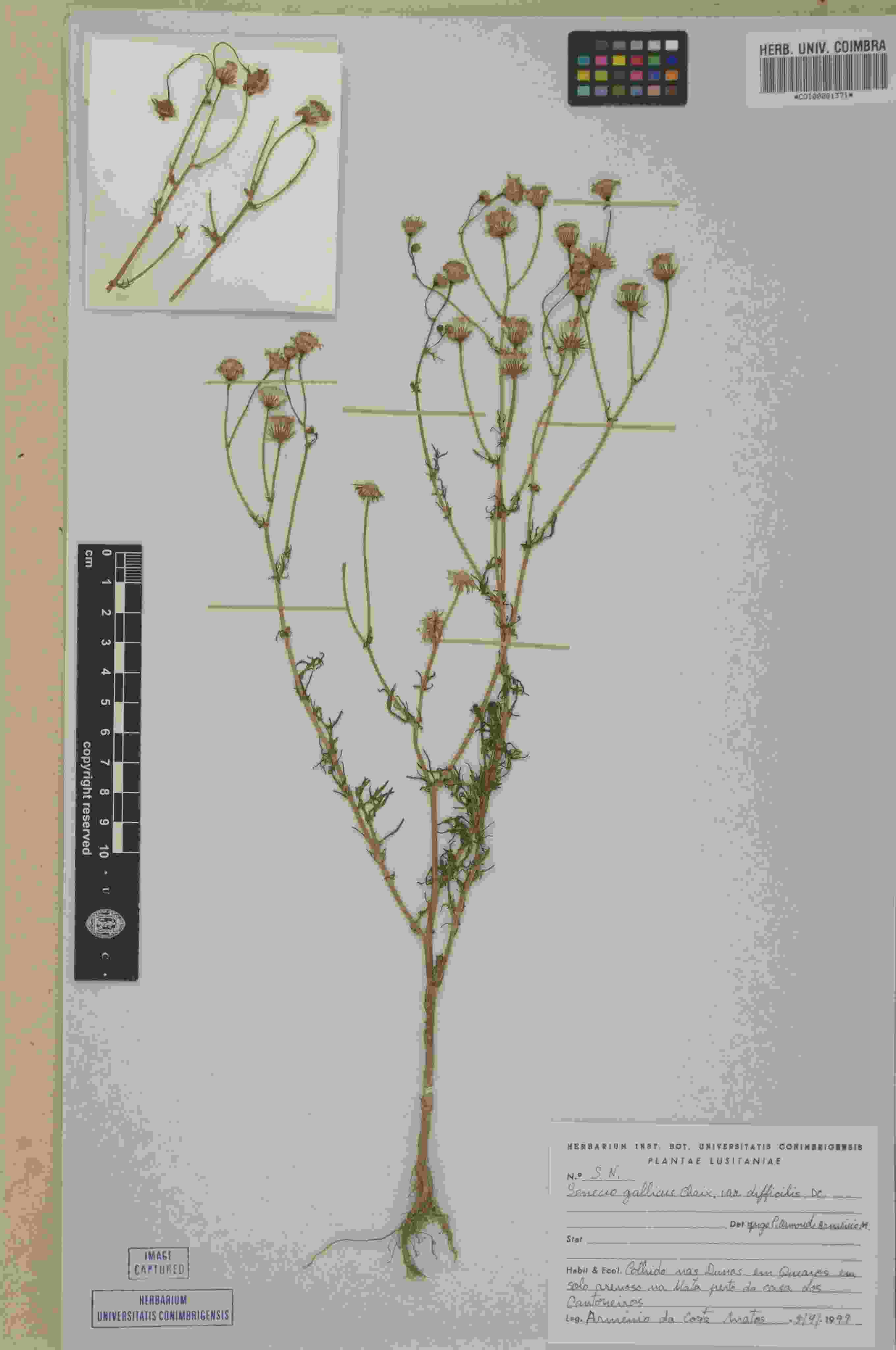

### Senecio_gallicus_2598791000.jpeg

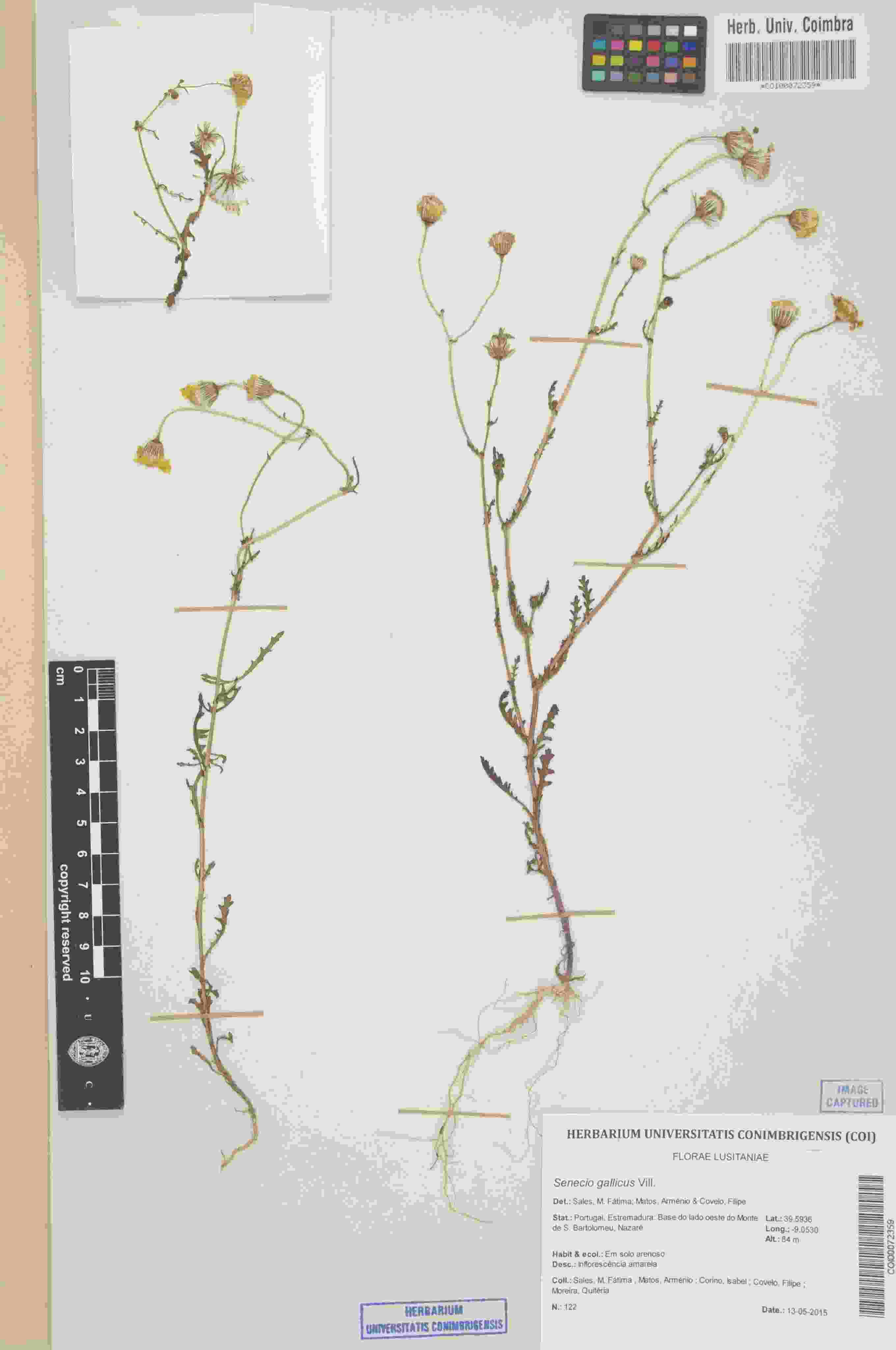

### Silene_kubanensis_1697783672.jpeg

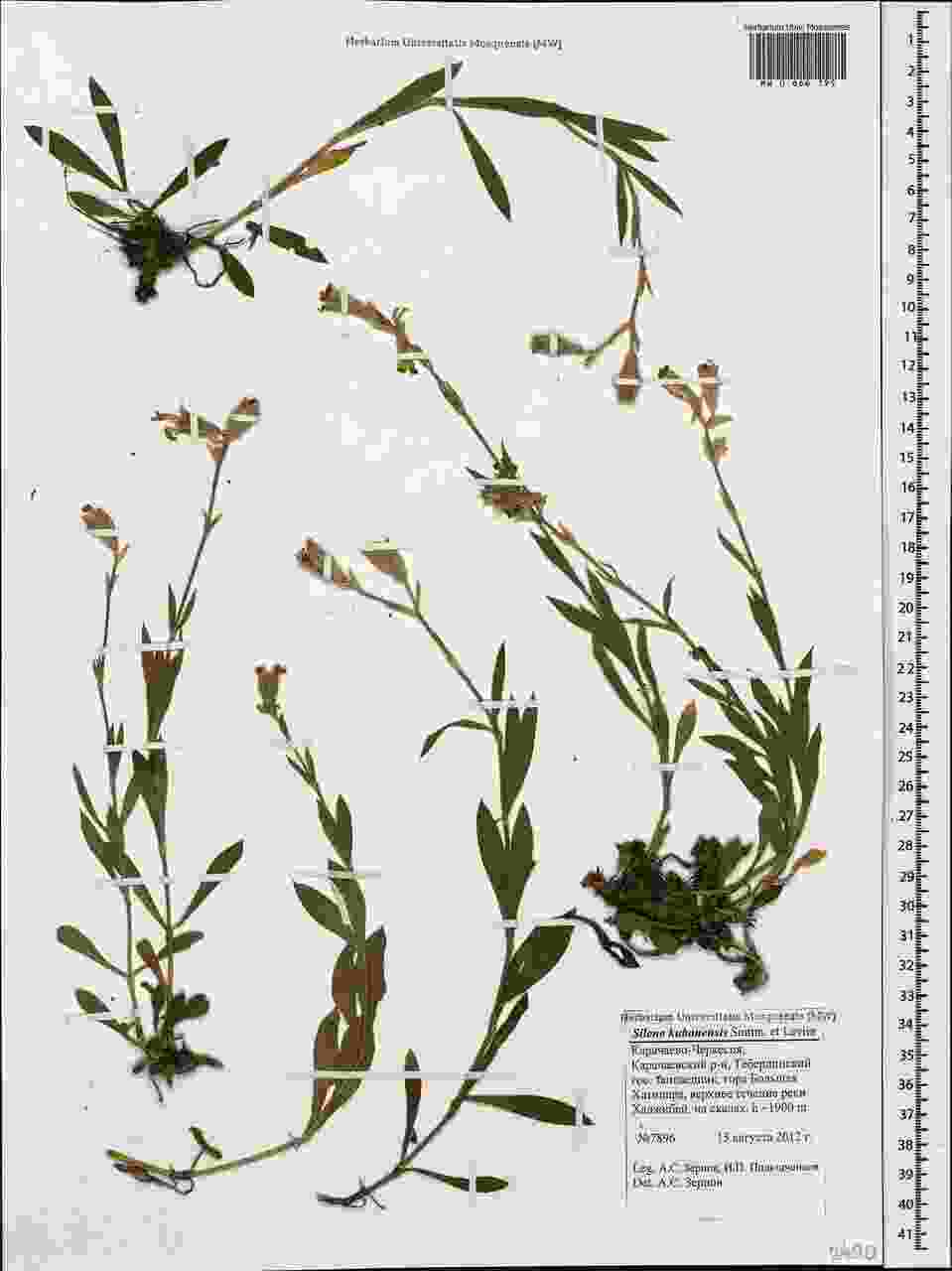

### Silene_kubanensis_1697783683.jpeg

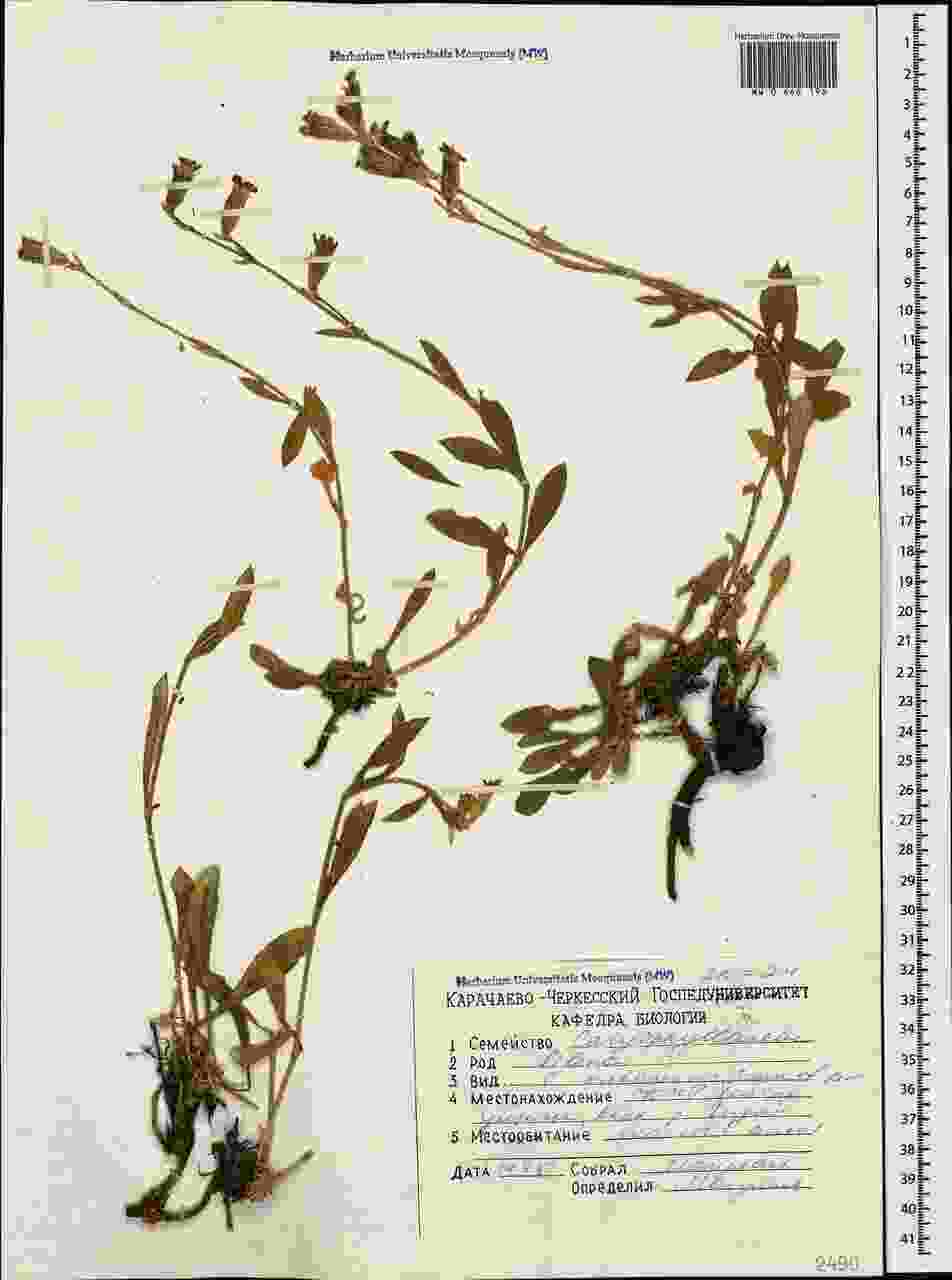

### Silene_kubanensis_1697783687.jpeg

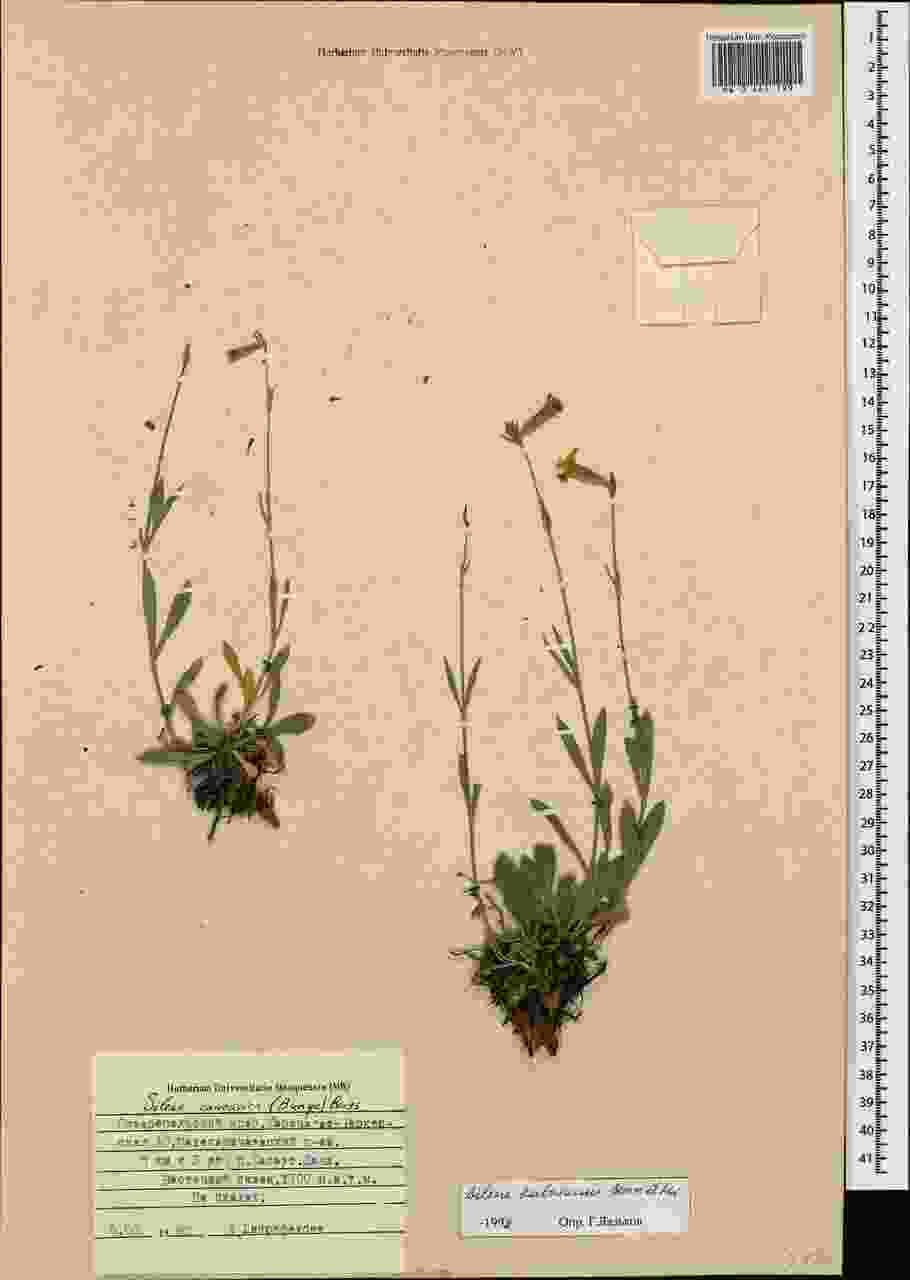

### Silene_kubanensis_3424031472.jpeg

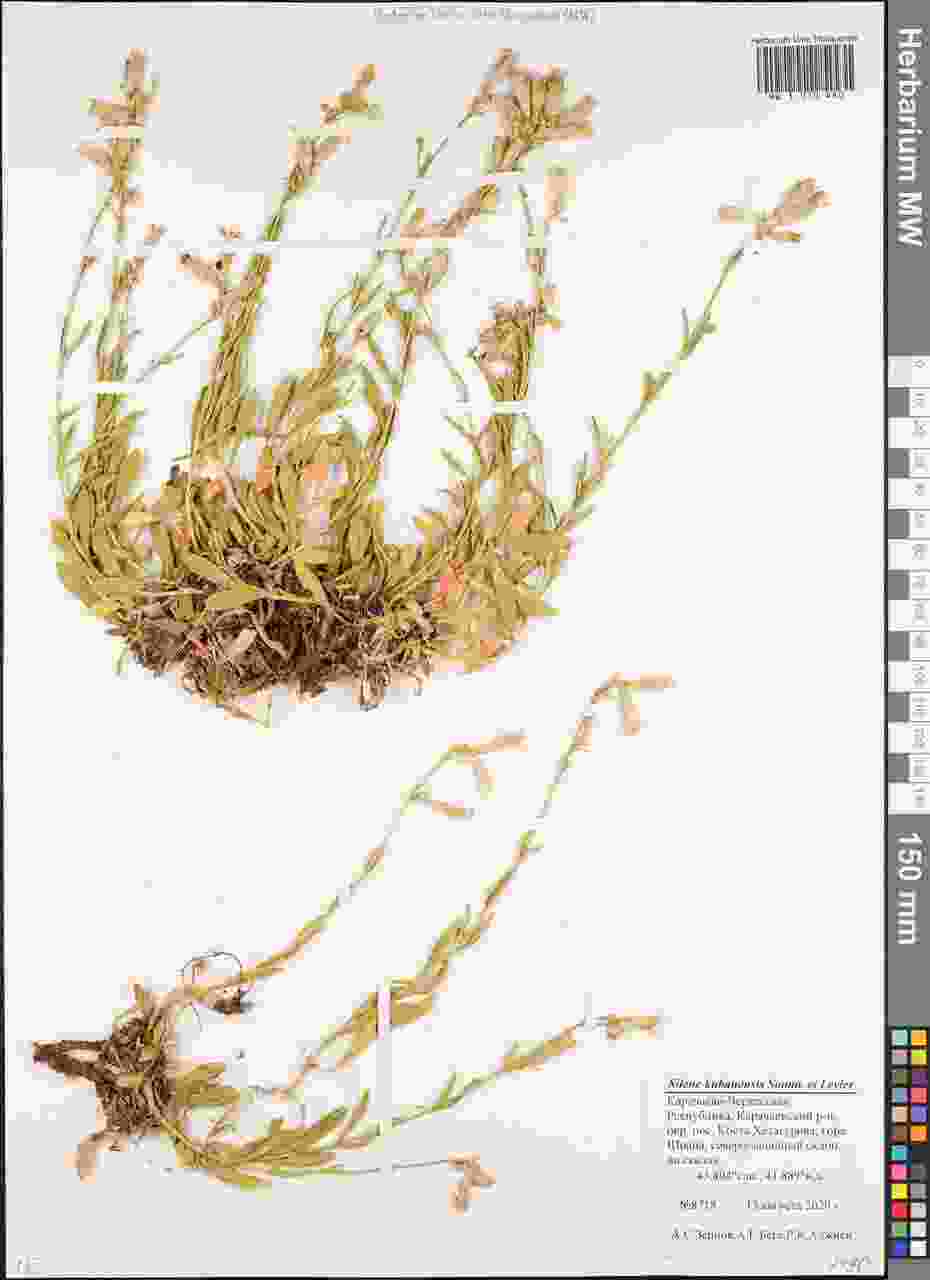

### Silene_kubanensis_3424035468.jpeg

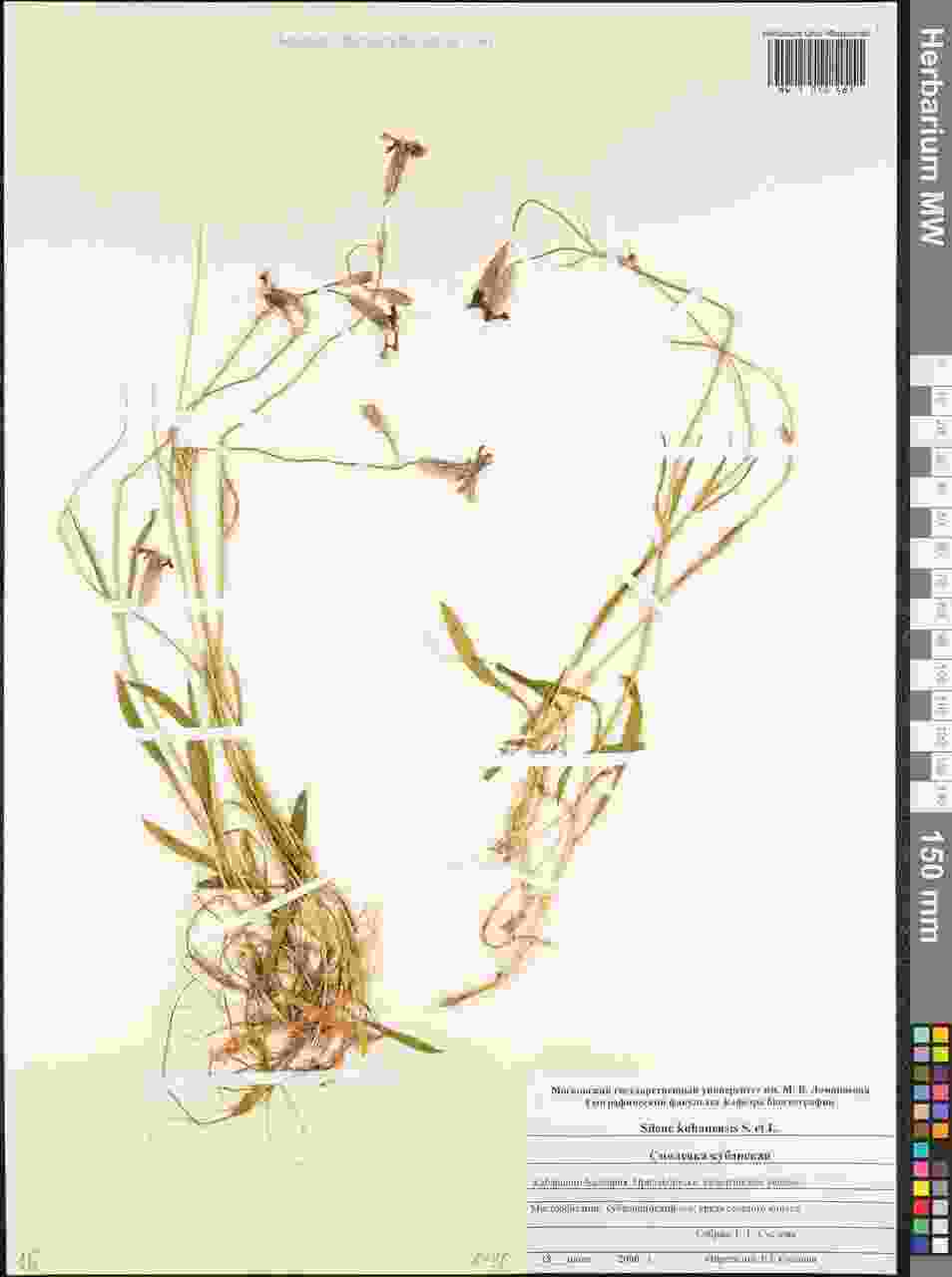

### Trichanthecium_yavitaense_1318930226.jpeg

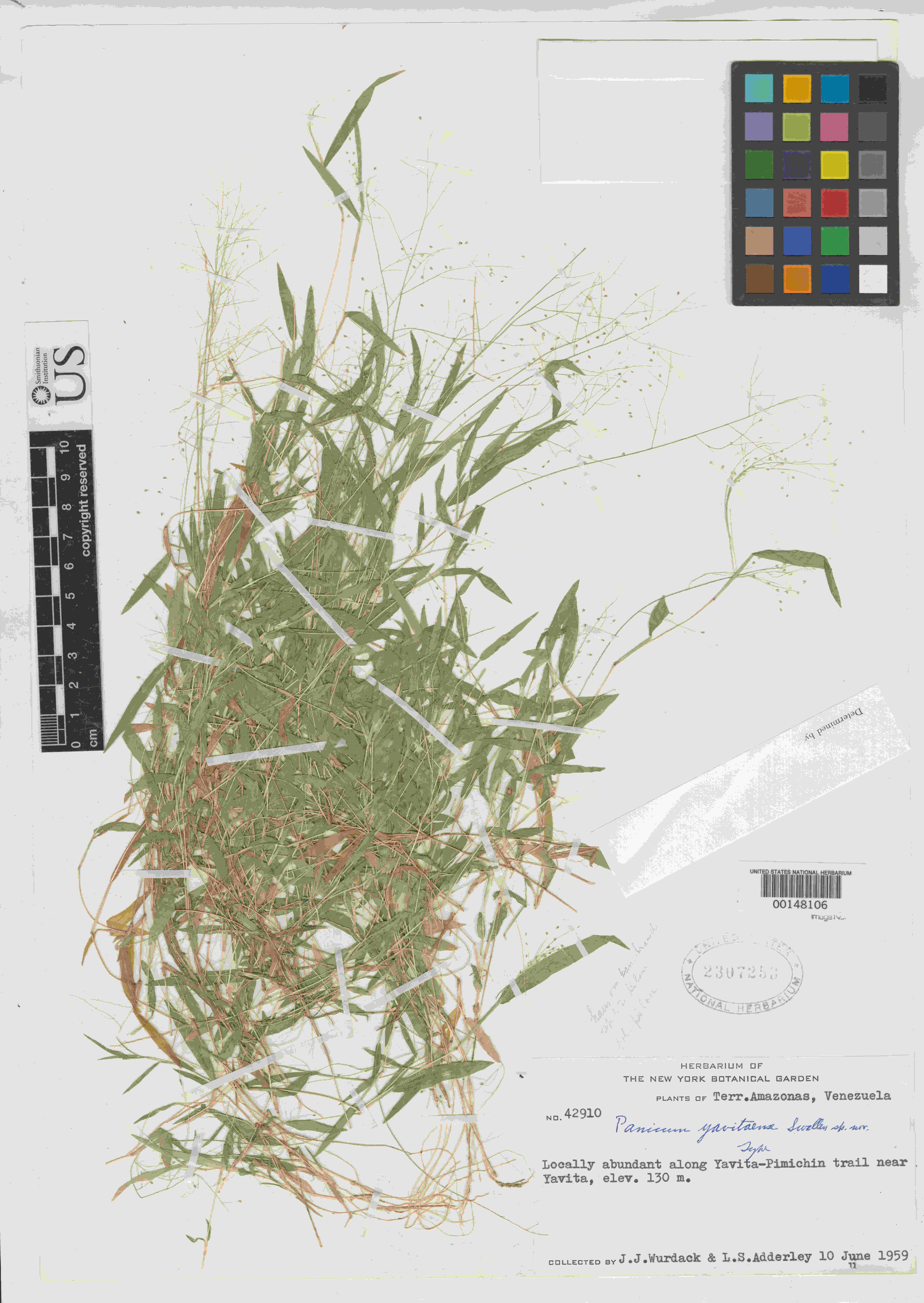

### Trichanthecium_yavitaense_1456316555.jpeg

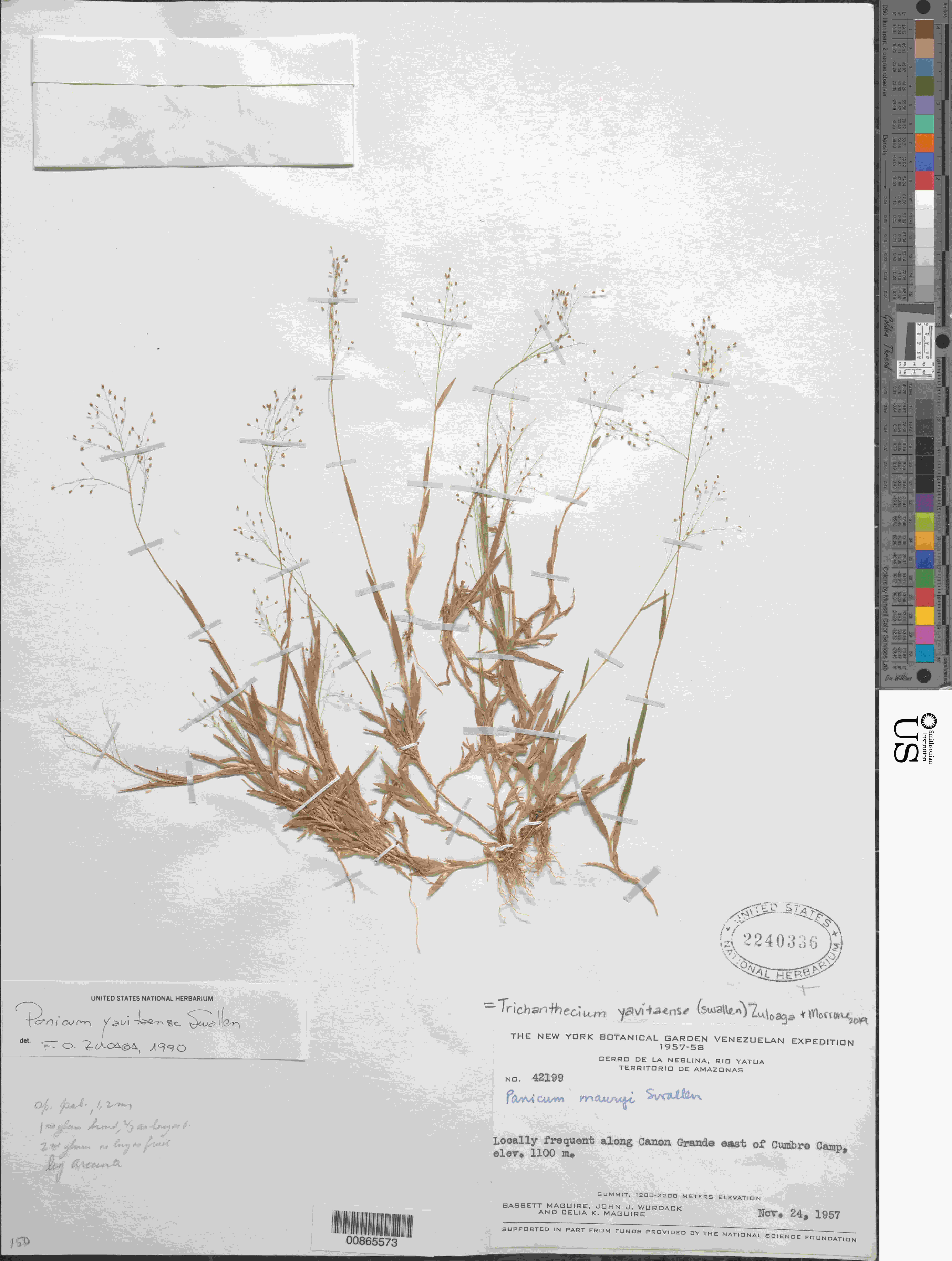

### Trichanthecium_yavitaense_1930180310.jpeg

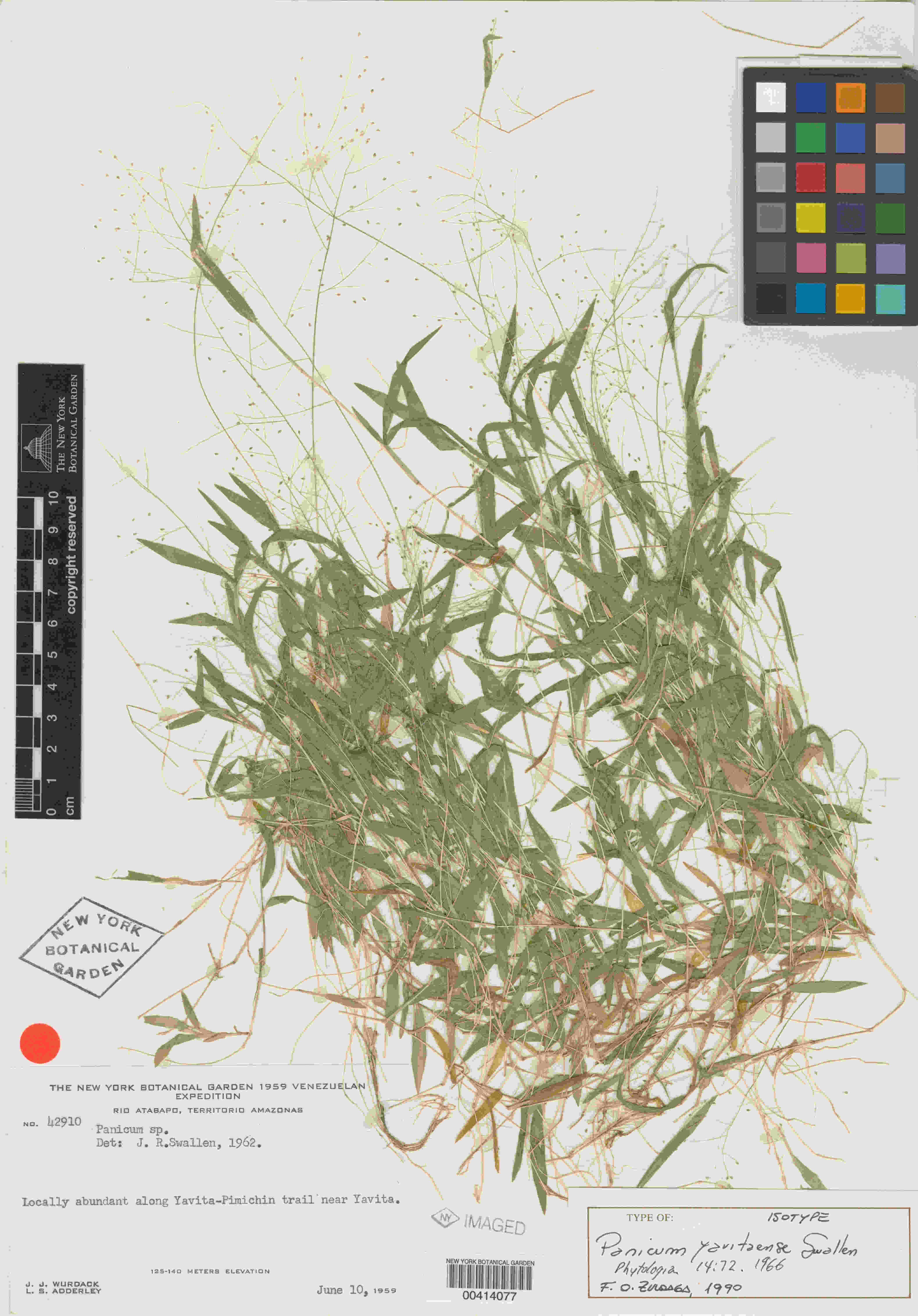

### Trichanthecium_yavitaense_4031773599.jpeg

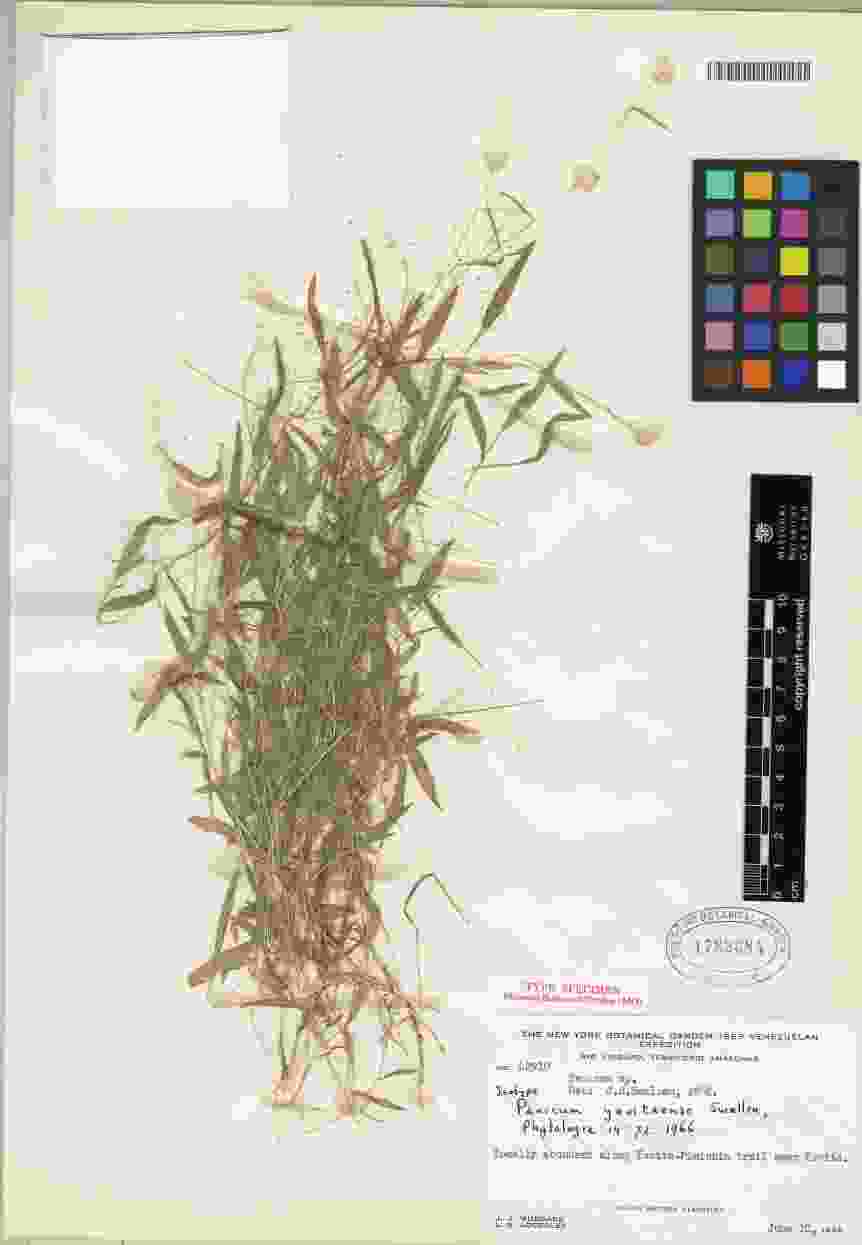

### Trichanthecium_yavitaense_4519428744.jpeg

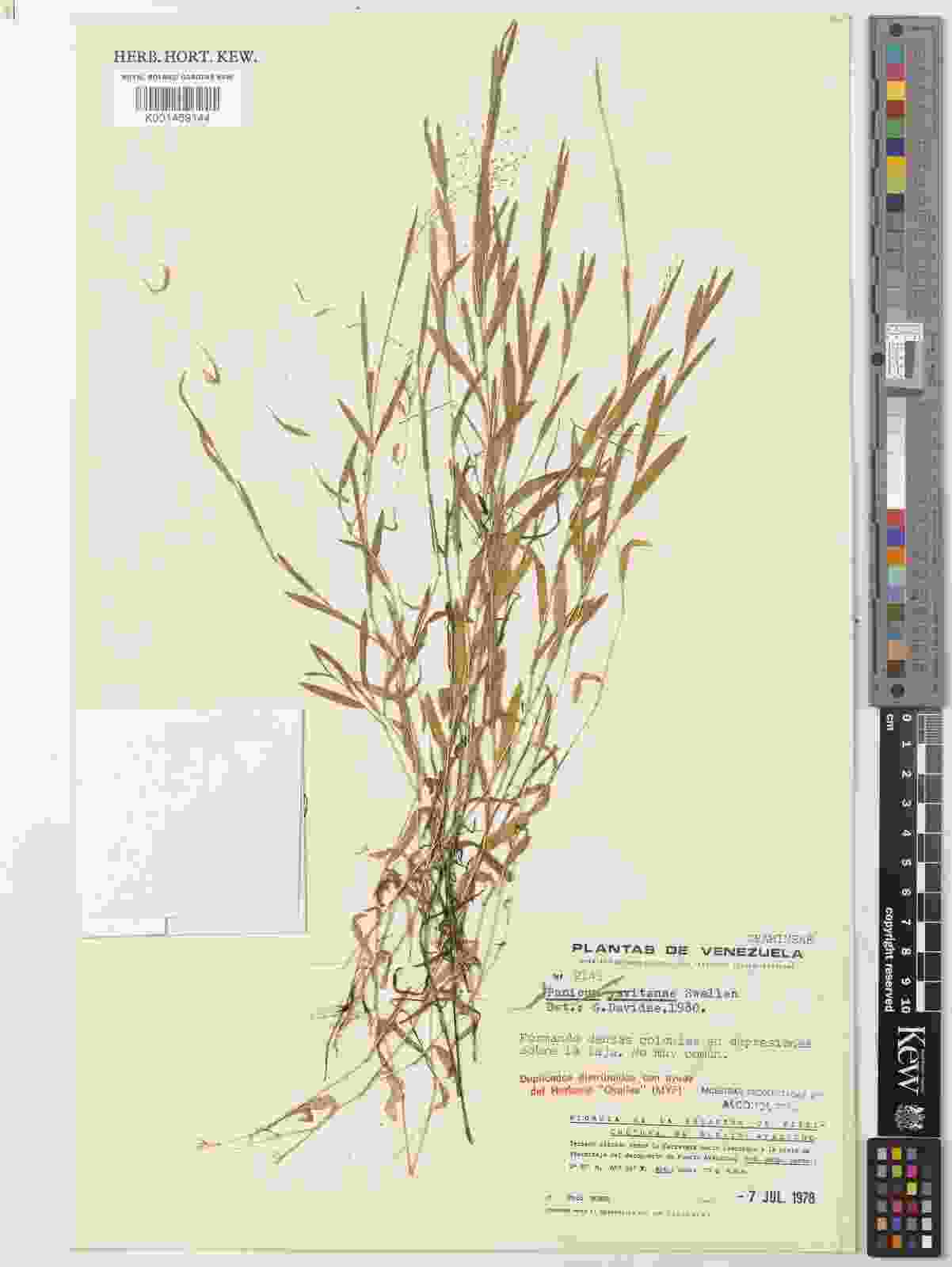

### Tricholepis_tibetica_574825372.jpeg

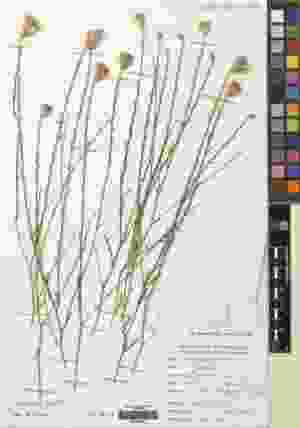

### Tricholepis_tibetica_575034483.jpeg

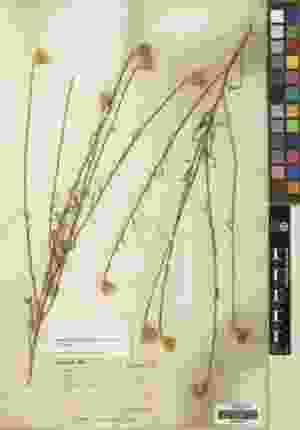

### Tricholepis_tibetica_1456012418.jpeg

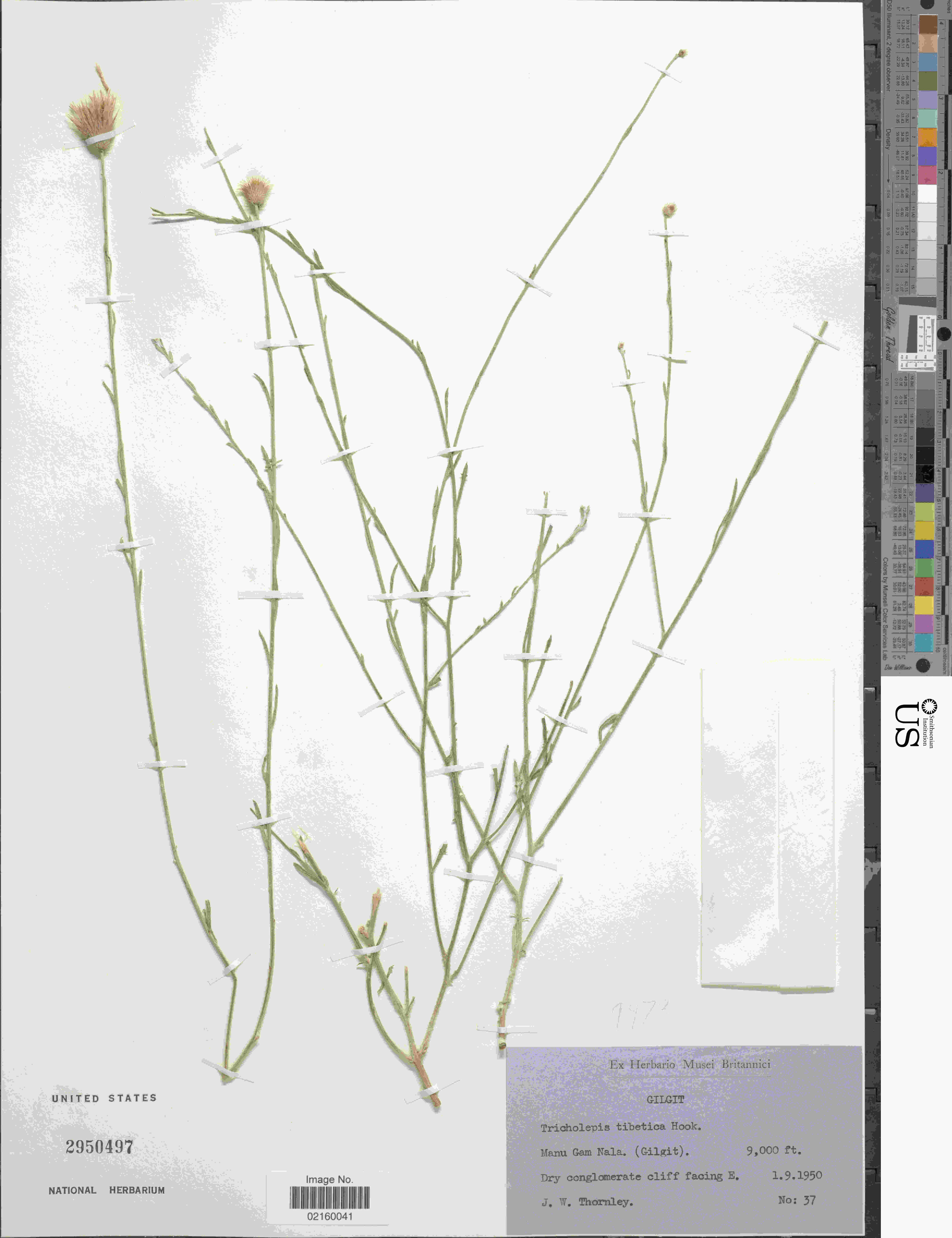

### Tricholepis_tibetica_1456141728.jpeg

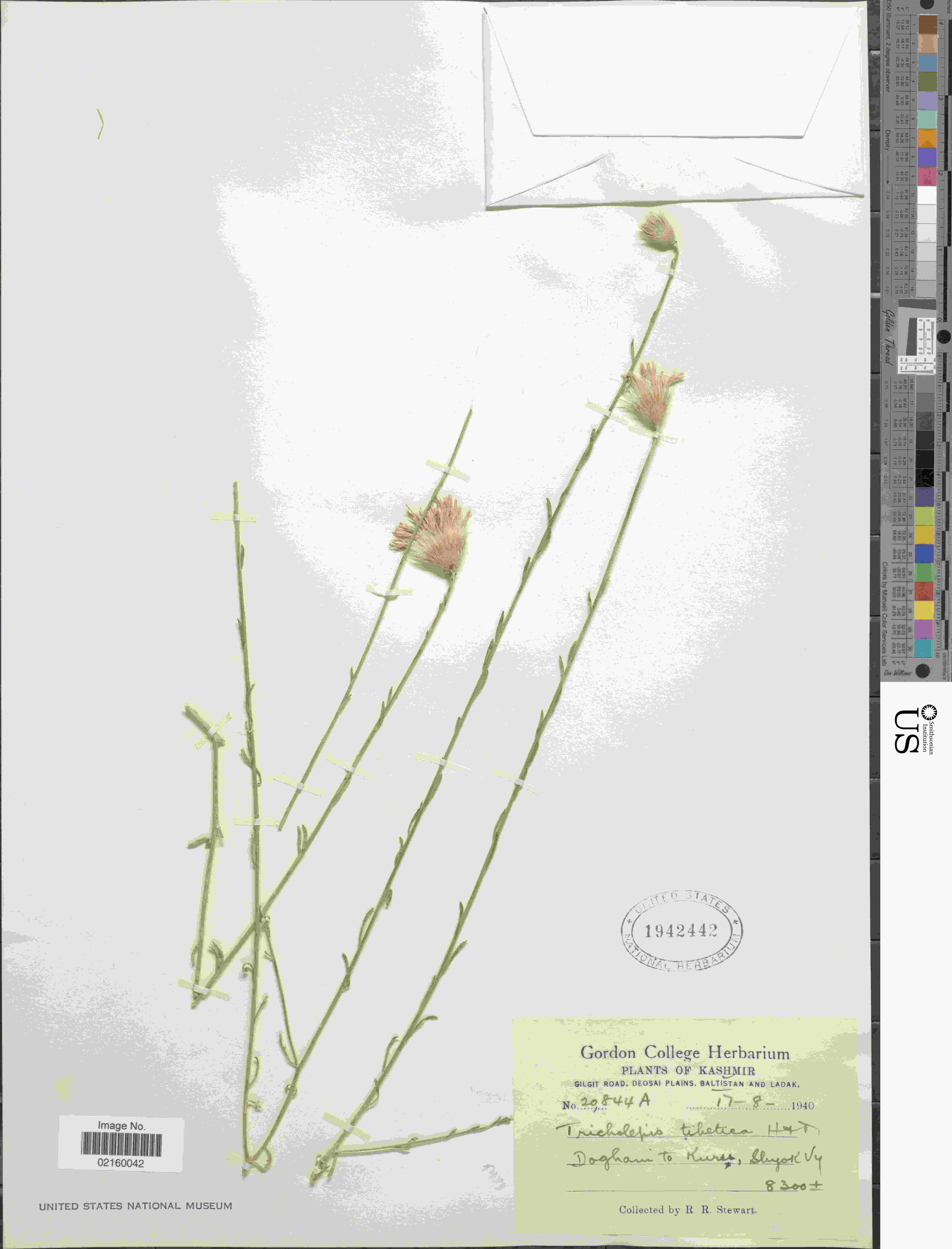

### Tricholepis_tibetica_1988955708.jpeg

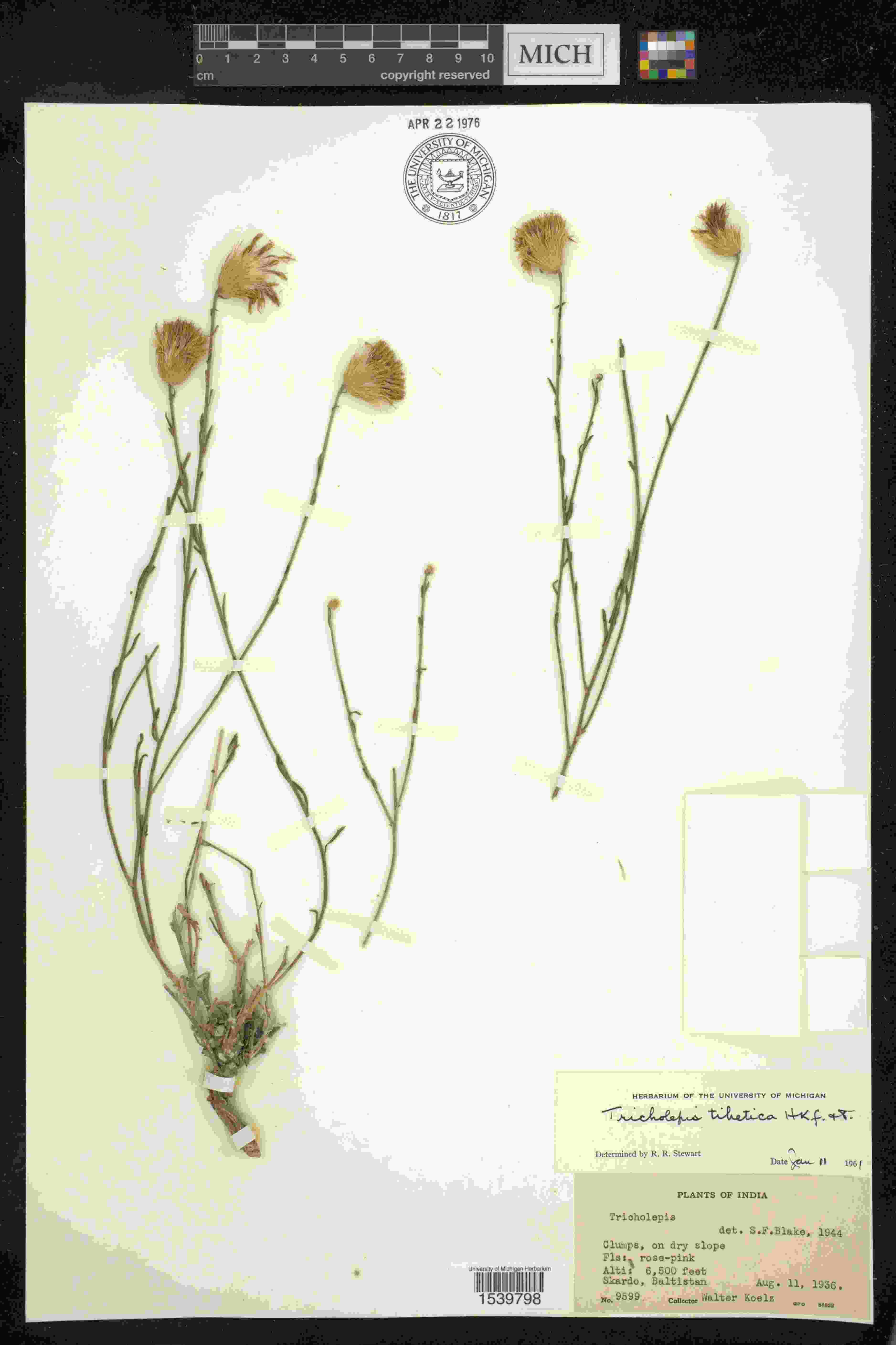

### Tridentopsis_buckleyana_1929262468.jpeg

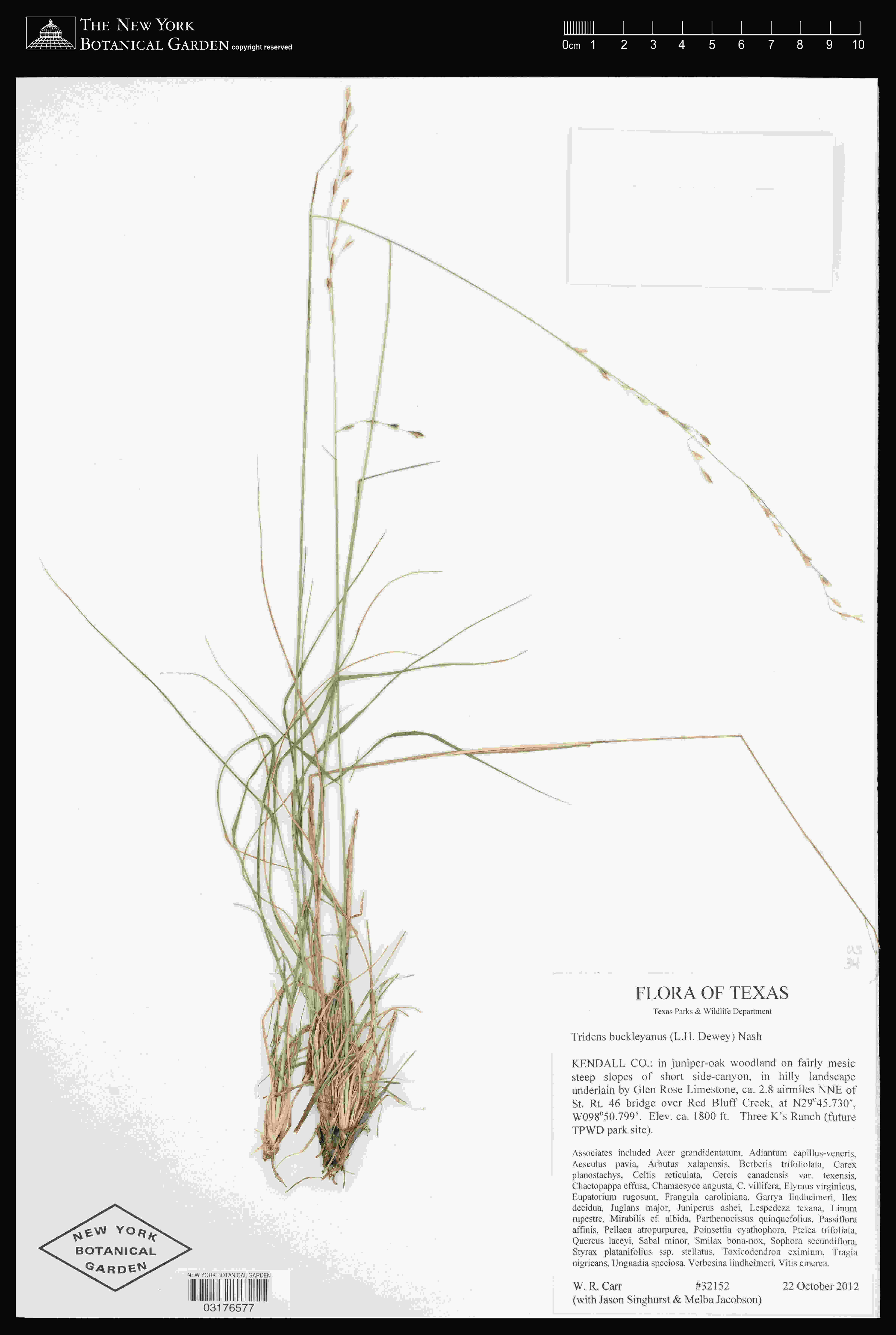

### Tridentopsis_buckleyana_2807560244.jpeg

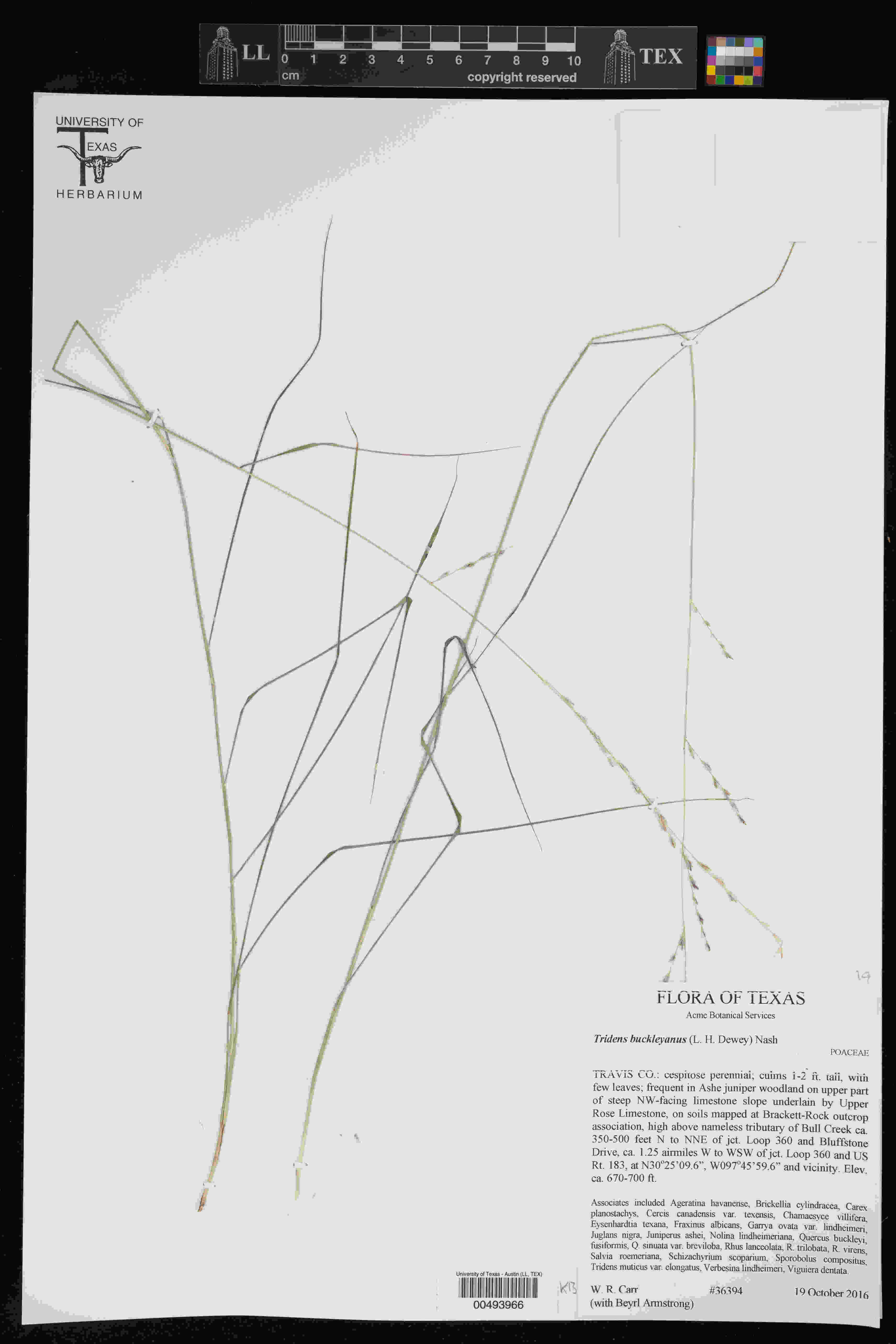

### Tridentopsis_buckleyana_4041438315.jpeg

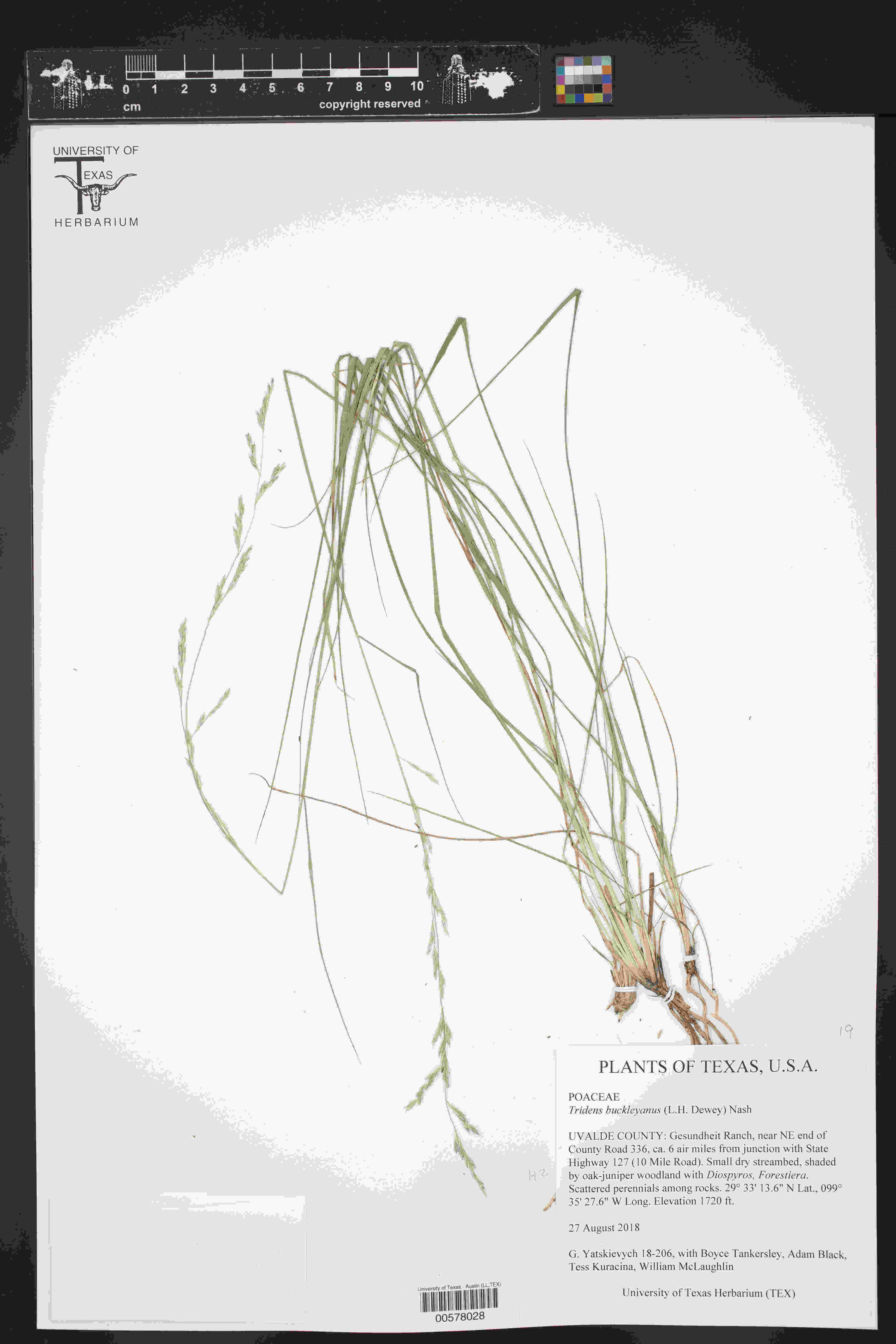

### Verbascum_austroiranicum_4524877318.jpeg

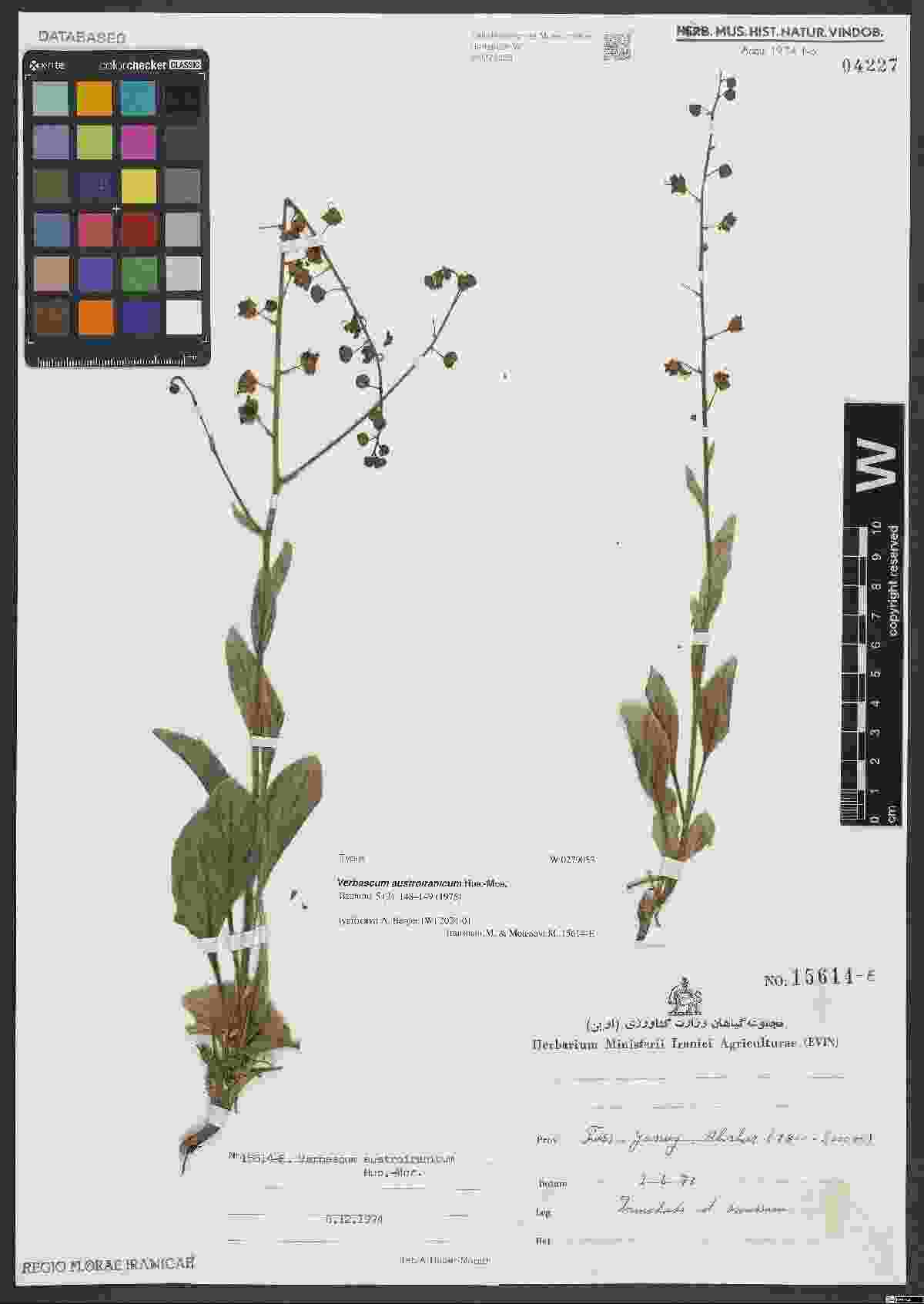

### Viscum_orientale_2515190297.jpeg

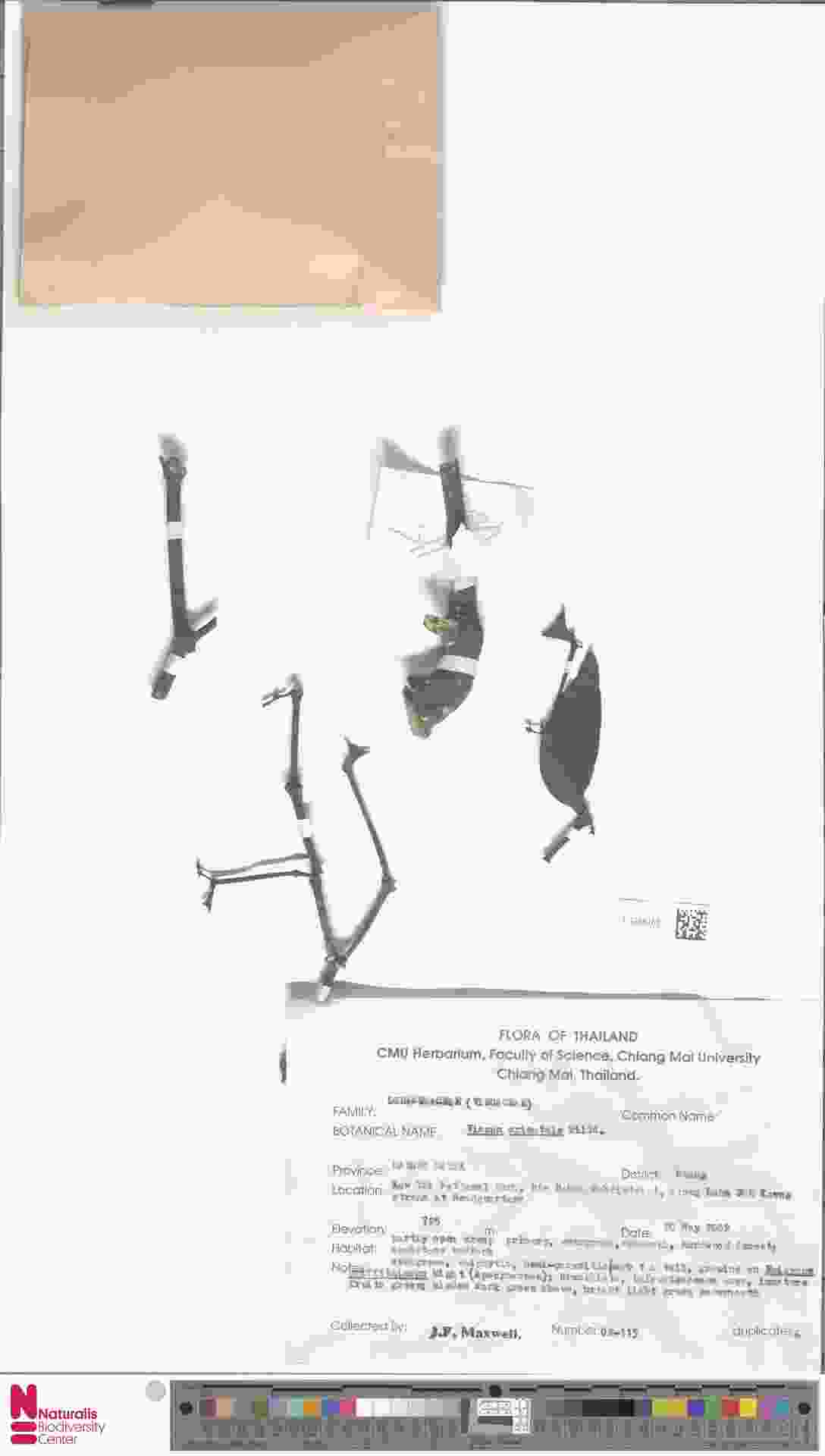

### Viscum_orientale_2516872067.jpeg

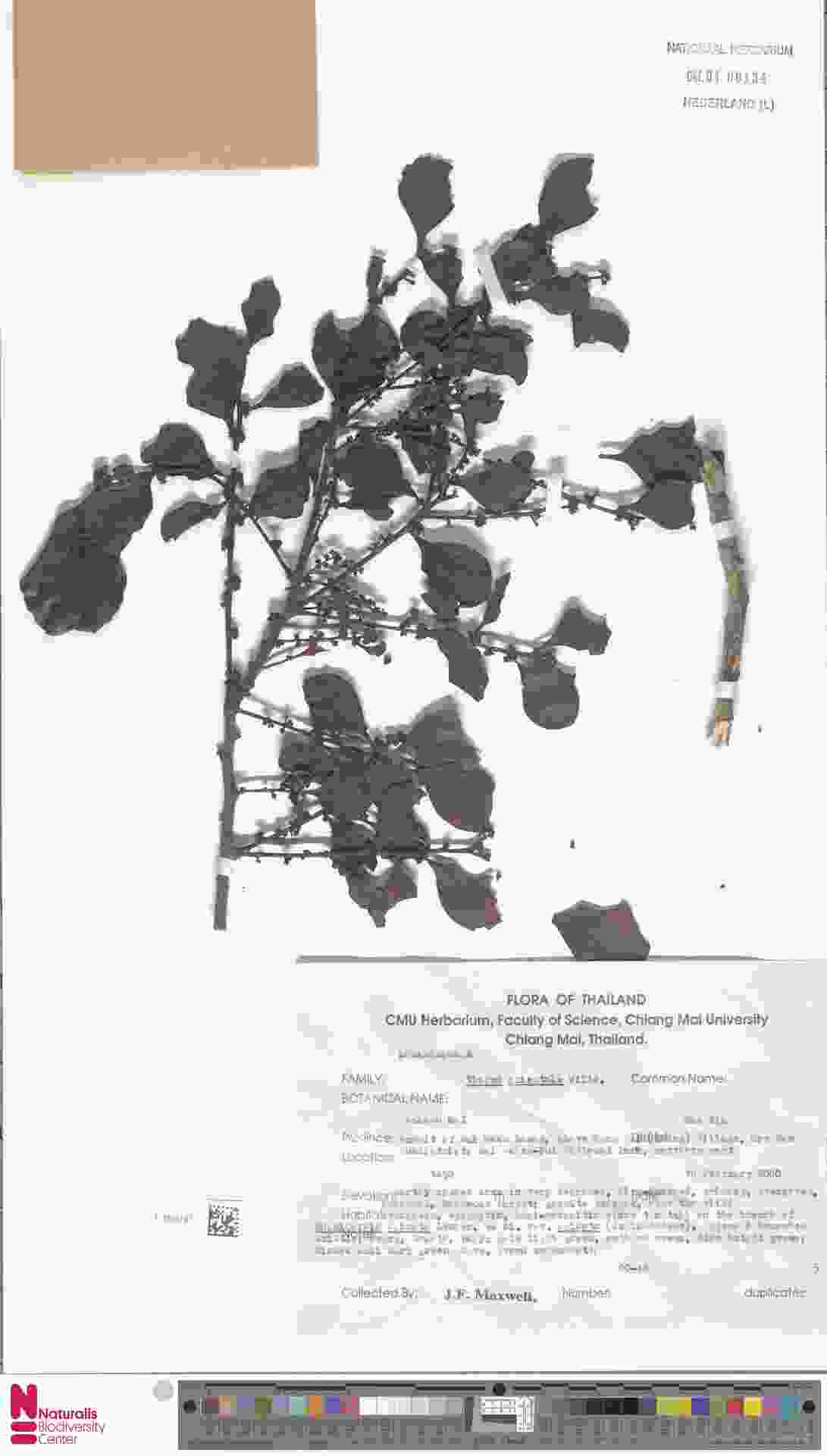

### Viscum_orientale_4514278310.jpeg

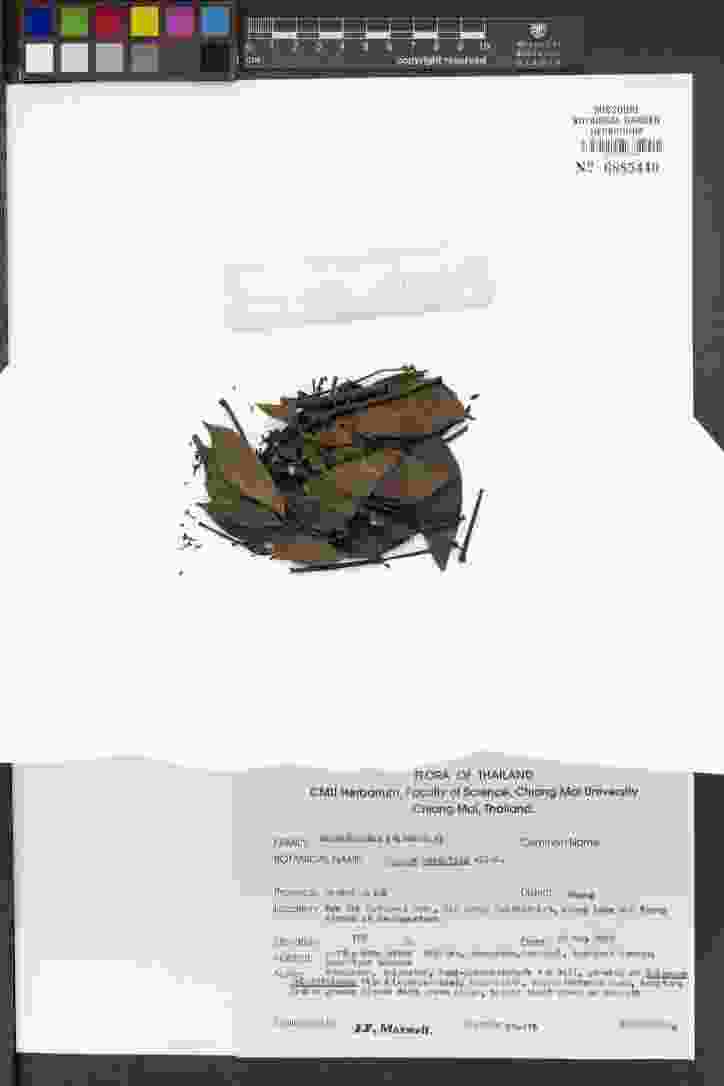
